# Atom-level generative foundation model for molecular interaction with pockets

**DOI:** 10.1101/2024.10.17.618827

**Authors:** Xingang Peng, Fenglin Guo, Ruihan Guo, Jiayu Sun, Jiaqi Guan, Yinjun Jia, Yan Xu, Yanwen Huang, Muhan Zhang, Jian Peng, Xinquan Wang, Chuanhui Han, Zihua Wang, Jianzhu Ma

## Abstract

Understanding molecular interactions is essential to structural biology and drug discovery. Despite the progress of AI models in revealing and exploiting the interaction mechanisms for various applications, they are predominantly tailored to specific tasks without fully exploiting the underlying transferability across molecular data and tasks. Here, we present PocketXMol, an atom-level generative foundation model to decipher fundamental atomic interactions for general protein-pocket-interacting molecular tasks. It adopts a novel unified generative framework with an innovative task prompt mechanism and an exclusive atom-level representation, making it applicable to diverse tasks covering structure prediction and design of small molecules and peptides, without requiring fine-tuning. PocketXMol was compared to 55 baseline models across 13 typical tasks, achieving state-of-the-art performance in 11 tasks and remaining competitive in the others. We successfully utilized PocketXMol to design novel small molecules that inhibit caspase-9 with efficacy comparable to that of commercial pan-caspase inhibitors. Furthermore, we employed PocketXMol to design PD-L1-binding peptides, demonstrating a success rate substantially higher than random library screening. Three representative peptides underwent further experiments, which validated the cellular specificity and confirmed their potential for molecular probing and therapeutics. PocketXMol presents a powerful and versatile tool with promising prospects for future applications and will have a profound impact on AI-aided drug discovery.

## Introduction

Molecular structures and interactions are crucial for their biological functions yet remain challenging to characterize^1, 2^. Recently, artificial intelligence has made significant strides in understanding and deciphering these complexities^3–9^. Molecular structure prediction tools, such as AlphaFold, have demonstrated an unprecedented capacity to accurately construct molecular structures^6–11^. Additionally, generative AI models have found significant applications in various molecular design tasks, including the design of functional proteins^12–14^, binding peptides^15^, and small-molecule drug candidates^11, 16^. These models typically employ specialized algorithms and data representations tailored to specific tasks or molecular types. For instance, protein structure prediction models generally learn the conditional distributions of tertiary structures based on primary sequences, while protein design models focus on the joint distribution of sequences and structures^6, 14^. Another example is that different molecular categories are represented using distinct building blocks, such as amino acids for proteins and peptides, and atoms for small molecules^7, 11^. However, from a chemical perspective, all these tasks can essentially be viewed as learning the underlying atomic interactions based on molecular structure data, albeit from different aspects and scopes. The physical principles governing atomic interactions are broadly applicable across the entire molecular universe^17^. Thus, it is natural to consider the feasibility of unifying all molecular generative tasks related to atomic interactions within one model and treating all molecular types exclusively at the atom level, to leverage the shared fundamental principles and eliminate the need for building numerous task-specific models.

Another motivation for developing such a unified model is the remarkable success of foundation models across various domains, including language understanding^18, 19^, computer vision^20, 21^, network biology^22–24^, and pathology analysis^25, 26^. These models utilize large-scale data to systematically capture the interactions among the basic units of their respective systems, enabling them to generalize across a wide array of tasks^27^. Therefore, we are inspired to unify generative molecular tasks within a foundation model that learns the fundamental atomic interactions from abundant molecular data. We anticipate that the transferability of atomic interactions will lead to superior performance on involved tasks and benefit those tasks that lack sufficient data for constructing tailored models.

However, challenges exist for developing such foundation models. Firstly, it is non-trivial to represent diverse and highly flexible molecular tasks in a precise and unified manner. In natural language processing, tasks are expressed through text prompts, enabling their unification within language models^28^. Although the text prompts can also be used to describe molecular tasks in the molecular models^29–31^, they do not capture the essence of the tasks, i.e., they cannot properly express the relationship among tasks. Secondly, current generative frameworks are not suitable for simultaneously accommodating diverse molecular generative tasks. While the diffusion-based generative frameworks^32, 33^ have demonstrated robust generative power and successful application to individual molecular generative tasks^7, 13, 34–36^, integrating all these tasks within a single diffusion model is challenging due to the distinct noise prior distributions and denoising trajectories that the generative process relies on for different tasks. Lastly, an intrinsic molecular representation is essential for learning the shared underlying principles among molecules. Current molecular models adopt different representations for different molecular types, which can hinder learning unified knowledge across molecules.

To tackle these challenges, we present an atom-level generative foundation model, named PocketXMol, to unify generative tasks related to the interactions among protein pockets, small molecules, and peptides. Firstly, various molecular generative tasks, ranging from docking to design, are uniformly represented through a novel task prompt mechanism, which directly defines the inputs and outputs of tasks at the atom level to encode the internal relationship among tasks. Users can precisely control the model to execute specific generative tasks through the prompt and, more importantly, can leverage the prompt to execute generative tasks that are difficult to express through text. Secondly, to perform generative tasks, PocketXMol takes the idea of denoising from the diffusion models but discards the complicated dependency between data and noise of the diffusion generative framework. Specifically, PocketXMol is built as a simple but effective universal denoiser, which facilitates accommodating distinct task noise distributions into a unified molecular noise space and enables flexible integration of prior knowledge during the generation process. With the unified task prompt and the novel generative framework, all the relevant tasks and structural data are trained together, thereby enabling the model to learn the shared knowledge. PocketXMol is trained by restoring the 3D molecules from the noisy ones and does not require additional fine-tuning for individual tasks. Lastly, PocketXMol adopts the pure atom-level representation for all molecular types to push the model to learn the atomic interactions. Notably, PocketXMol overcomes the conventional need to explicitly model amino acids for proteins and peptides, marking a pioneering achievement in generating amino acids through exclusive atom-level modeling. This results in a completely distinct treatment of peptides with all existing models and exploits the most fundamental transferability between small molecules and peptides.

We assembled a dataset comprising two million 3D structures, including small molecules, protein–peptide complexes, and protein–small-molecule complexes. These structures were collected from PDBBind^37^, Binding MOAD^38^, CrossDocked2020^39^, PepBDB^40^, AlphaFold DB^3^, ChEMBL^41^, ZINC^42^, GEOM-Drug^43^, CREMP^44^, and prior work^45^ (Table S4). Redundant entries overlapping with the test sets were removed before training PocketXMol (Supplementary Notes 6.2.2). We extensively evaluated its performance on 13 generative tasks (small molecule docking, linear peptide docking, cyclic peptide docking, molecular conformation generation, structure-based drug design, 3D molecule generation, fragment linking, PROTAC drug design, fragment growing, molecule optimization, linear peptide design, cyclic peptide design, and peptide inverse folding) in non-redundant test sets and compared them against 55 different baseline methods using 51 different evaluation metrics. PocketXMol surpassed state-of-the-art methods on 11 out of 13 tasks and was competitive in the remaining ones. We also demonstrated its applicability in constrained docking or design with prior knowledge, enzyme substrate screening, and peptide design involving nonstandard amino acids. Next, we used PocketXMol to design caspase-9 inhibitors and validated one lead that suppressed caspase-3 and PARP1 cleavage under ABT-737 treatment, with one of its structurally optimized derivatives from the second design round showing specific inhibition comparable in efficacy to a commercial inhibitor. Finally, we leveraged PocketXMol to design peptides targeting PD-L1 and validated the efficacy of the designed binders. Among the 382 selected peptides, 76 peptides achieved a binding affinity of 10^−7^M and 15 peptides reached 10^−8^M, representing a success rate substantially higher than that achieved by random library screening. We selected three peptides to perform more rigorous validation at the cellular level, each showing effective binding to the cell membrane and distinct binding contrasts between PD-L1-positive and negative cells. Their effectiveness as molecular targeting probes was further proved by tail vein injections in lung tumor mouse models, and the potential for therapy was validated through the ligand inhibition assay.

## Results

### Overview of PocketXMol

As a unified model, the denoiser abstracts any input molecule as a set of atoms and chemical bonds to facilitate learning fundamental atom-level interactions. All generative tasks are directly defined through a task prompt composed of groups of binary indicators to control the generation of the molecules. The backbone of PocketXMol is a versatile molecular denoiser built on geometric neural networks and can produce clean molecules from the input noisy one based on the designated task prompt (Figure 1a). The generation process is repeatedly to add and remove noise on the molecule, as is inspired by the denoising mechanism of diffusion-based generative models^32, 33^. However, PocketXMol has major differences from diffusion models (Figure S7). To incorporate distinct noise prior distributions of different tasks, our model eliminates the input requirement of the time step or noise scale but instead automatically determines the noise type and scale from the noisy molecule with the assistance of the task prompt. Besides, our model is designed to directly predict the clean molecule rather than the previous molecule at the generation trajectory as the diffusion models do. These modifications break the denoising Markov chain imposed by the diffusion framework and thus lead to a flexible noise-adding process. Another extension is the detailed injection of noise down to the atom level to enable different molecular components to be injected with different noise, which has brought tremendous adaptability to support more generative tasks.

**Figure 1.**
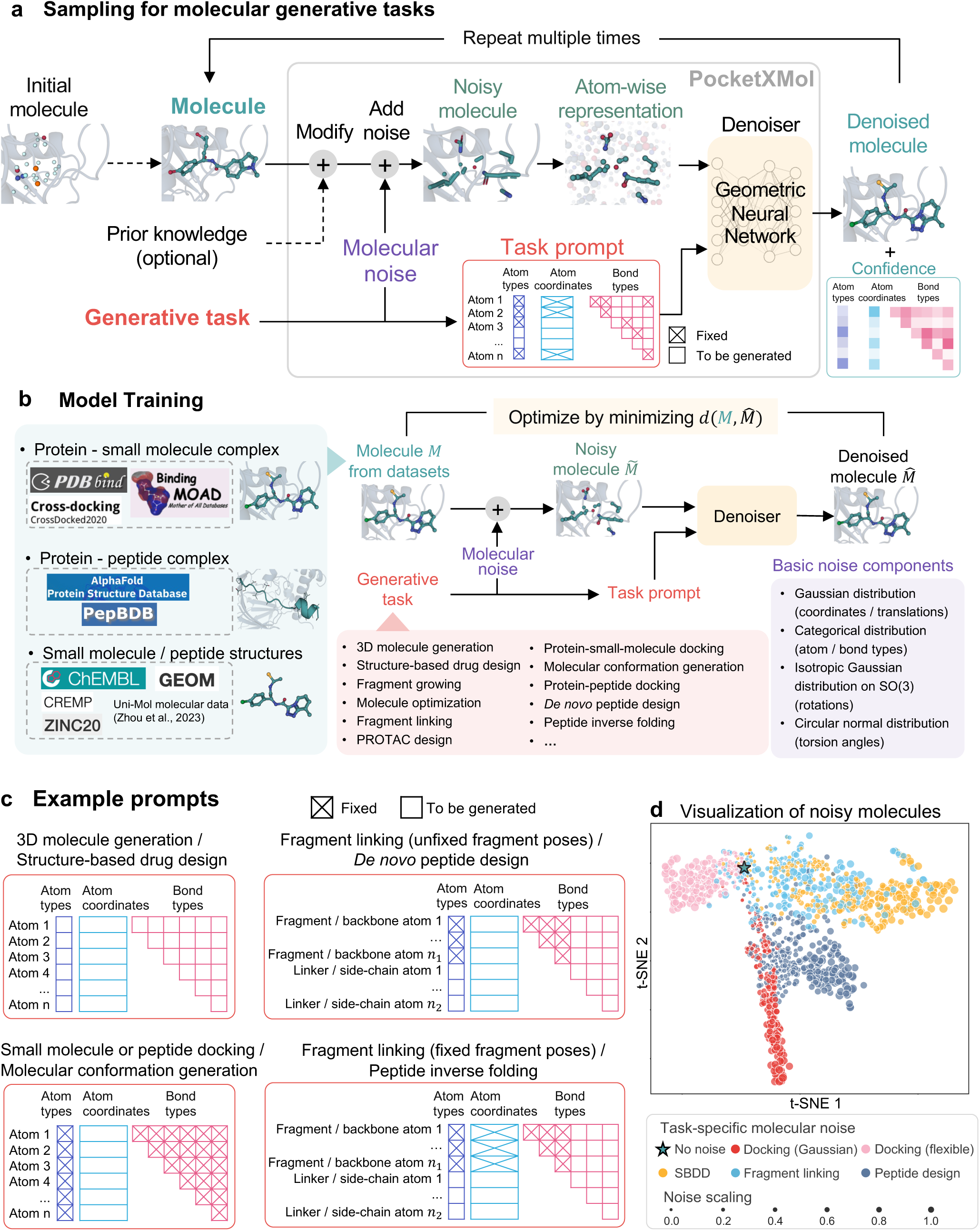
PocketXMol framework. **a**, Schematic representation of the sampling process of PocketXMol. PocketXMol perturbs the molecule using task-specific noise and prior-knowledge-guided modification and subsequently utilizes the denoiser neural network to generate the denoised molecule with the confidence scores. **b**, The training process of PocketXMol. The denoiser was trained to recover the molecules from the noisy ones, with data assembled from multiple 3D molecular structure datasets. **c**, Examples of how the task prompts are designed for different generative tasks. **d**, t-SNE visualization of the noisy molecules with different task-specific molecular noise. Different colors represent noisy molecules after the injection of different types of noise. We selected a protein-peptide complex and added different task-specific noise with random noise scales to the peptide molecule. For each molecule, we calculated the probability distributions of atom pairwise distances and probability distributions of the atom and bond types as the molecular representation. We then used t-SNE to visualize the representations.

To specify a generative task, users can set the task prompt to precisely convey intentions. A task prompt is a group of binary variables indicating whether each atom or bond of the input molecule should be fixed or requires to be generated for a particular generative task (Figure 1c). This task representation accurately captures the requirements of any molecular generative task in a more fundamental way than other representations such as natural language descriptions^29, 31, 46^. As shown in Figure 1b, with the unified molecule and task representation, all generative tasks were jointly trained on a comprehensive dataset covering all relevant structure data (redundancy with test sets was removed). Once trained, PocketXMol can be immediately applied to individual tasks without the requirement of fine-tuning, which is different from many other pre-training models^5, 22–24, 45^. The training scheme of PocketXMol facilitates learning the shared commonality among tasks, as exampled by the distributions of the noisy molecules with different task-specific molecular noise (Figure 1d). For instance, fragment linking, which generates partial molecules from the other parts, is located between the clean molecule and structure-based drug design (SBDD) that generates the complete molecule in the embedding space; Peptide design, which requires generating side-chain atoms and docking backbone atoms, is located between docking with Gaussian noise and the molecule design tasks (SBDD and fragment linking). Docking with flexible noise, which perturbs coordinates using random rigid translation and rotation of torsion angles, deviates from all other tasks where the Gaussian noise is applied to the coordinates (Figure 1d). More details are outlined in Methods and Supplementary Notes.

### Performance on small molecule design

#### Structure-based drug design

Our evaluation began by assessing the performance of PocketXMol on small molecule design. Structure-based drug design (SBDD) with AI models, which involves designing novel small molecules based on the 3D protein pocket, has emerged as a successful approach in drug discovery as it directly models the interactions between amino acids and ligand atoms in the 3D space^16^. One example is that Novartis has successfully designed three WDAC inhibitors using an SBDD model combined with their compound databases^47, 48^. To evaluate the performance of PocketXMol on the SBDD task, we conducted a comprehensive comparison against multiple existing methods, including a VAE-based method liGAN^49^, autoregressive-based methods 3DSBDD^50^ and Pocket2Mol^48^, and diffusion-based methods TargetDiff^35^, DecompDiff^51^, IPDiff^52^, and AliDiff^53^. We designed 100 molecules for each of the 100 protein pockets in the benchmark set used in previous work^35, 39, 48–51^. PocketXMol could employ two different generation strategies: a default refine-based strategy (PocketXMol) and an auto-regressive-like strategy (PocketXMol-AR) (Supplementary Notes 4.3.4). We first evaluated the designed molecules with 14 diverse metrics, spanning molecular properties, the coherence of molecular graph distribution, and the fidelity of the 3D structures of the generated molecules. PocketXMol ranked first in 11 out of 14 metrics with balanced overall performance (Figure 2a and Tables S5 and S6). The performance of the refine-based PocketXMol and the auto-regressive-based PocketXMol(AR) were similar, suggesting that the strength of our model primarily stemmed from the training process rather than the sampling strategy. Following previous work^35^, we utilized AutoDock Vina^54^ to calculate ratios of molecules with better Vina scores than the corresponding molecules in the test set for each pocket to evaluate their binding. We also employed the PoseBusters 3D validity checker^55^ as a filter due to the previous observation that some models tended to sacrifice the intrinsic rationality of the molecules in pursuit of stronger binding interactions with protein pockets^56^. Results showed that the proportion of better-Vina and 3D valid molecules generated by PocketXMol exceeded all baselines with clear margins (Figures 2b and S8a). Notably, in the absence of 3D structural filtering, around half of the molecules generated by liGAN demonstrate better Vina scores, however, almost all generated molecules displayed a certain degree of irrationality, highlighting the importance of applying the 3D valid checker. Furthermore, we found that PocketXMol’s self-confidence scores correlate with Vina scores, as selecting generated molecules with higher confidence consistently led to improved Vina performance. (Figures 2c and S8b).

**Figure 2.**
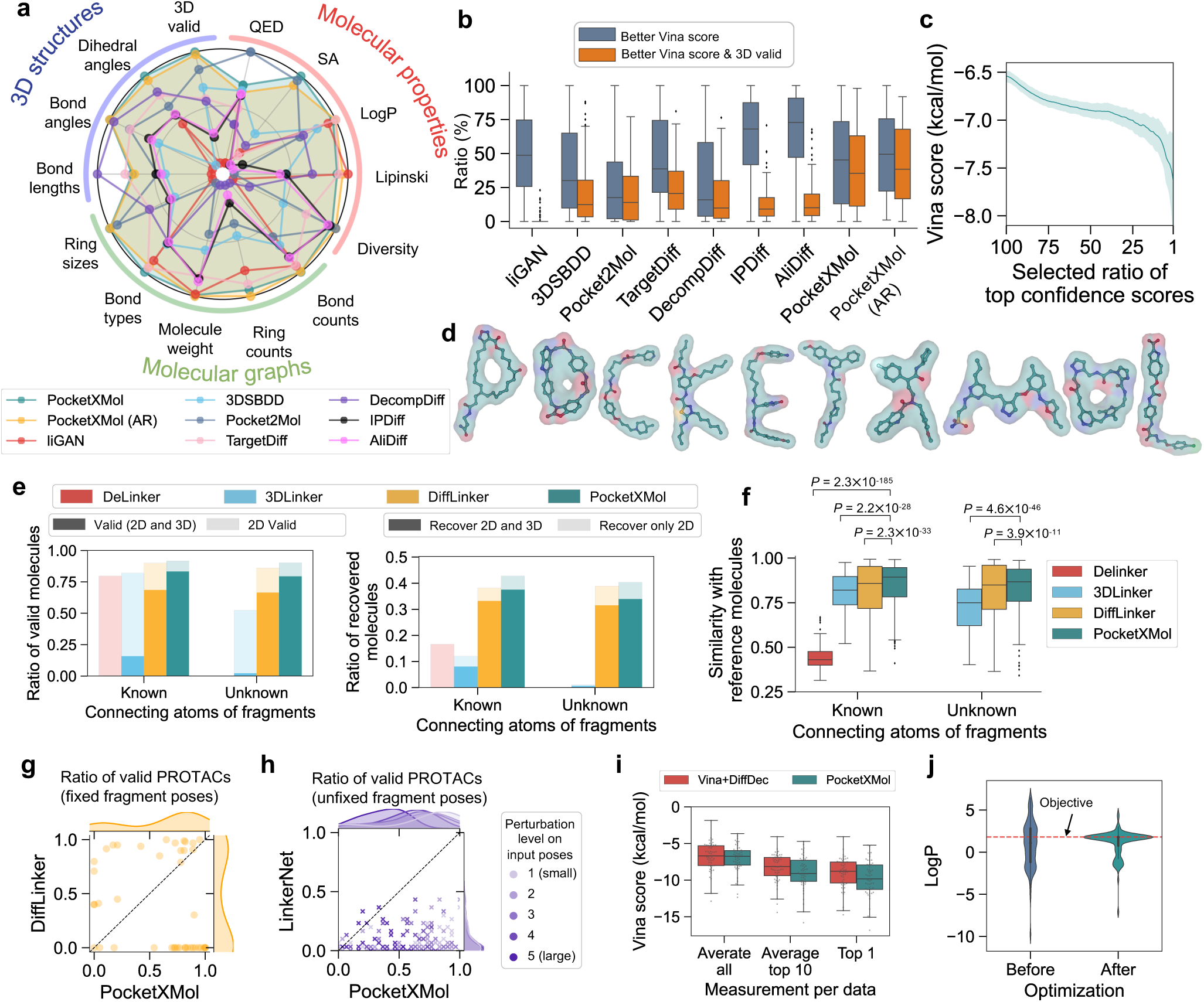
Performance on small molecule design. **a**, Performance on the SBDD task. Each metric on the radar plot was normalized to a scale from 0 to 1, with 1 representing the best method and 0 indicating the worst. The values of the metrics can be found in Tables S5 and S6. The definitions are in Supplementary Notes 6.1. **b**, Ratios of the generated molecules exhibiting both better Vina score and 3D validity for each pocket. Better Vina score is defined as those with Vina scores better than the corresponding reference molecule in the test set. The 3D validity was annotated by the PoseBusters validity checker. *P* values based on the one-sided paired *t*-test (*n* = 100) for ratios of better Vina scores and 3D valid molecules between baselines and PocketXMol are 4.9 × 10^−24^, 1.6 × 10^−12^, 2.2 × 10^−10^, 2.4 × 10^−12^, 1.3 × 10^−13^, 2.3 × 10^−15^ and 1.6 × 10^−16^ from left to right, respectively. The box plot outlines quartiles of the metrics with whiskers spanning up to 1.5 times the interquartile range. **c**, Vina scores of molecules with different ratios of top PocketXMol confidence scores. **d**, Molecules generated by PocketXMol with shape specifications. **e**, Ratios of valid molecules (left) and recovered molecules (right) for fragment linking, with known connecting atoms or not. **f**, Similarities between the generated molecules and the reference molecules in the fragment linking test set. *P* value was calculated using a one-sided paired *t*-test. The sizes for each box from left to right are 282, 283, 413, 415, 255, 409, and 416. The differences in sizes were caused by the fact that no valid molecules were generated for some input pairs by some methods. **g**, Comparison of the ratio of valid molecules between DiffLinker and PocketXMol for PROTAC design with fixed fragment poses. Here the validity included both 2D and 3D validity and the ratios were calculated for each fragment pair. **h**, Comparison of the ratio of valid molecules between LinkerNet and PocketXMol for PROTAC design with unfixed fragment poses. The fragment poses were randomly perturbed with five different noise levels. Here the validity included both 2D and 3D validity and the ratios were calculated for each fragment pair and noise level. **i**, AutoDock Vina scores of generated molecules for fragment growing. The x-axis represents the average of the top 1, top 10, and all Vina scores of the generated molecules for each test sample. **j**, LogP distribution of 100 molecules in the SBDD test set before and after applying the molecule optimization by PocketXMol. The optimization was to modify the molecules to adjust their LogP values towards the objective of 1.8. The box within the violin plot outlined the interquartiles (*n* = 100).

To evaluate PocketXMol’s robustness to variations in input pocket structures, we also generated molecules using pocket structures derived from alternative sources. Specifically, instead of directly using the true binding (holo) protein structures in the test set, we considered the following four types of input structures: (1) predicted by AlphaFold^6, 57^, (2) side-chain repacked by Rosetta^58^, (3) retrieved from the RCSB PDB with identical sequences, (4) apo structures retrieved from the RCSB PDB with identical sequences (Supplementary Notes 6.4). Results showed that most evaluation metrics remained largely consistent across the different types of input pocket structures, particularly for structures predicted by AlphaFold and those retrieved from the RCSB PDB (Figure S9 and Table S7). Structures with side chains repacked by Rosetta showed relatively weaker Vina scores, likely due to repacked side chains partially occluding the binding pockets and resulting in flatter pocket shapes. Apo structures retrieved from the RCSB PDB also exhibited slight shifts in the distribution of molecular properties such as QED, Lipinski metrics, and diversity. This can be attributed to differences in pocket characteristics between apo and holo forms, which may lead to the generation of molecules with distinct physicochemical properties.

#### 3D molecule generation

Compared to the SBDD task, the 3D molecule generation task operates without the constraints imposed by protein pockets. In drug discovery, it acts as an alternative *in silico* approach that first generates 3D molecules and then screens them against the target proteins. We evaluated the performance on this task and compared it with diffusion-based molecular generation models^59–63^ by generating 1000 molecules with sizes akin to drug-like compounds. The molecules generated by PocketXMol exhibited a high ratio of validity and also matched the distributions of real drug-like molecules well (Figure S10).

In order to evaluate the capability of incorporating prior knowledge, we showcased generating 3D molecules under arbitrary volumetric shape specifications. Specifically, we generated 3D molecules conditioned on the shapes of the letters of our model’s name, which is achieved by adjusting the atom coordinates to align with the point clouds of the shapes during the generation process^14^ (Supplementary Notes 4.3.5). The resulting molecules not only exhibited the predefined shape but also featured reasonable topology and structures (Figure 2d and Table S8), demonstrating its designing capacity with prior knowledge conditions.

#### Fragment linking

Having explored PocketXMol’s performance for *de novo* design, we next considered designing based on existing fragments or molecules, including fragment linking, fragment growing, and molecule optimization. These strategies are widely used in drug development to achieve desired properties by modifying or adding certain chemical groups of existing molecules^16, 64, 65^. Fragment linking involves the strategic design of a linker to unite disconnected fragments into one complete and stable drug-like molecule^66^. Such technology can be extended for PROTAC (proteolysis-targeting chimeras) drug development, where a molecular bridge needs to be designed to connect a protein-targeting ligand with an E3 ligase-recruiting ligand^67^.

Due to the atom-level molecular representation, one of the advantages of PocketXMol is its multi-functionality in handling various settings of the fragment linking task, while other methods were proposed to cover only some specific settings (Figure S11a). In a typical fragment linking task, molecular linkers are designed based on fixed fragments within the protein pocket. We conducted comparisons in this setting with models that support fixed-fragment linker design, including DeLinker^66^, 3DLinker^68^, and DiffLinker^69^ on a linker design test set proposed by DiffLinker^69^ from the Binding MOAD dataset^38^. For each model to be compared, we generated 100 3D molecules for each fragment pair in the case of either knowing the connecting atoms of fragments or not. PocketXMol could generate higher proportions of molecules satisfying both 2D and 3D validity than the second-place model DiffLinker (Figure 2e left). Here, 2D and 3D valid molecules indicate molecules with realistic chemical topology examined by RDKit^70^ and valid 3D geometry examined by the PoseBusters validity checker, respectively. PocketXMol also recovered the 2D molecular graphs or the 3D poses of the ground truth 3D linkers for more fragment pairs than baselines (Figure 2e right). Additionally, the molecules sampled by PocketXMol more closely resembled the ground truth molecules than those by baselines, based on the similarity score proposed by DeLinker (Figure 2f).

#### PROTAC linker design

For PROTAC linker design, we considered two scenarios where the poses of the fragments were specified or not. We compared PocketXMol with DiffLinker^69^, the state-of-the-art linker design model that was conditioned on known fragment poses, and LinkerNet^71^ proposed specifically for unknown fragment poses. The test set was constructed by LinkerNet from the PROTAC-DB dataset^72^ with 43 PROTAC molecules. In the scenario where fragment poses are provided, we generated 100 linkers for each pair of fragments (a pair of warhead and E3 ligand). PocketXMol produced 60% valid (both 2D and 3D) PROTAC molecules on average, much higher than 34% achieved by DiffLinker (Figure 2g). PocketXMol also generated molecules exhibiting higher similarities with the real PROTAC molecules (Figure S11b). In the scenario where fragment poses were unknown, the models needed to generate the linker and determine the poses of fragments simultaneously. We thus perturbed the input fragment poses randomly at five distinct noise levels and generated 30 molecules for each pair of fragments and noise levels. In comparison to LinkerNet, PocketXMol successfully designed more valid (both 2D and 3D) PROTAC molecules (Figure 2h). The PROTACs designed by PocketXMol also demonstrated certain similarities with the real ones (Figure S11c).

#### Fragment growing

Fragment growing is another drug development strategy that starts from target-binding molecular fragments with relatively small sizes and applies chemical modifications to design complete molecules with improved binding affinity and other properties. To evaluate PocketXMol’s capability for fragment growing, we constructed a test set using the data in the SBDD test set by segmenting molecules into multiple fragments using BRICS decomposition^73^ and selected connecting fragments as the initial fragments with similar sizes. For each pair of the protein pocket and the fragment graph (without 3D poses), we invoked PocketXMol to generate 100 molecules with the same sizes as the original molecules in the test set. We compared PocketXMol with a baseline method DiffDec but had to first apply Vina to dock the fragments as the inputs for DiffDec because DiffDec cannot determine the fragment poses by itself. Results showed that PocketXMol generated molecules with better completeness and more similarities with the ground truth molecules than the baseline (Figure S12a,b). The molecules designed by PocketXMol showed better bindings in terms of the average of top 1, top 10, and all Vina scores of the generated ones for each test sample (Figure 2i). Moreover, the molecules by PocketXMol with the top 1 or top 10 Vina scores displayed significantly improved Vina scores compared to the corresponding reference molecules containing the same initial fragments in the test set (Figure S12c).

#### Molecule optimization

The aim of molecule optimization is to modify input molecules to improve specific properties while preserving structural similarity. For this task, PocketXMol uses the same prompt as in the SBDD task, where all ligand variables are unfixed and subject to generation. However, unlike SBDD, which begins from a randomly sampled molecule with high initial noise (noise scale 1.0), molecule optimization starts directly from the input reference molecule with a lower noise scale. This enables PocketXMol to generate new molecules that partially retain the structure of the input. The degree of similarity is implicitly controlled by the noise level: lower noise leads to higher similarity. Then, generated molecules with improved properties are then selected and can be used as new starting points for next optimization rounds (Supplementary Notes 4.3.9).

We demonstrated this setup by optimizing the LogP values of 100 molecules in the SBDD test set toward a target value of 1.8, which is commonly considered favorable in many drug design scenarios^74^. Molecules with LogP values closer to 1.8 were considered better. In each round, PocketXMol generated multiple candidates, selected those with LogP values closest to the target value, and used them as new starting points for the next round. After three rounds, the LogP values of the optimized molecules rapidly converged to the target, exhibiting significantly smaller errors to 1.8 compared to those before optimization (one-sided paired *t*-test, *P* = 6.12 × 10^−20^*, n* = 100; Figures 2j and S13).

#### Caspase-9 inhibitor design

Mitochondrial permeabilization could release cytochrome c into the cytosol to form the apoptosome and initiate intrinsic apoptosis by activating caspase-9 (CASP9); caspase-9 thereafter cleaves and activates caspase-3/7 (CASP7/3), the central executioner of apoptosis, to further cleave downstream substrates, such as PARP1, DFFA, and lamin A, and promote cell death^75^ (Figure 3a). Caspase-9 mediated intrinsic apoptosis is a major barrier of tumor endogenous cyclic GMP–AMP synthase (cGAS)–stimulator of interferon genes (STING) pathway, and inhibition of caspase-9 could significantly provoke antitumor immunity to synergize with radiation and immunotherapy^75^. However, the current first-in-class caspase inhibitor, emricasan, is a pan-caspases inhibitor that could inhibit caspase-9/3/7 medicated intrinsic apoptosis, as well as caspase-8 mediated extrinsic apoptosis and caspase-1 mediated inflammasome activation. Notably, the inflammasome pathway is also critical to antitumor immunity^76, 77^; therefore, Emricasan might also target caspase-1 to suppress antitumor immunity. To investigate whether PocketXMol could help *de novo* design specific small molecule inhibitors, we applied it to design inhibitors targeting caspase-9 (Supplementary Notes 8.1) and validated them experimentally. We synthesized 16 designed molecules (Figure S14) and one of them (ID: 84663, Figure 3b) could effectively inhibit ABT-737, a BCL-2 inhibitor triggering mitochondria-mediated intrinsic apoptosis, mediated caspase-9 and caspase-3 activation and cleavage of PARP1, a target of activated caspase-3 (Figure 3c and Figure S15). This small molecule was completely novel and had low similarities with existing caspase inhibitors (Figure S16). There were no similar molecules found in the ZINC20 database^42^ and the ChemDiv Library (www.chemdiv.com), and the most similar molecules in the ChEMBL^41^ and PubChem^78^ databases also had massive differences with the molecule 84663 (Figure S17). Based on the predicted structure of this lead molecule (Figure 3b), we then optimized it for improved binding (Supplementary Notes 8.1.2) and synthesized 15 optimized molecules (Figure S18), among which four (IDs: D12, D13, D18, and D19; Figure 3d) showed remarkable effects. As shown in Figure 3e, treatment with ABT-737 could enhance the cleavage of caspase-3 that was diminished after QVD (Q-OVD-oph, a powerful pan-caspase inhibitor) treatment; notably, the designed molecules could also suppress the activation of caspases-9 and caspases-3 with efficacy comparable to QVD.

**Figure 3.**
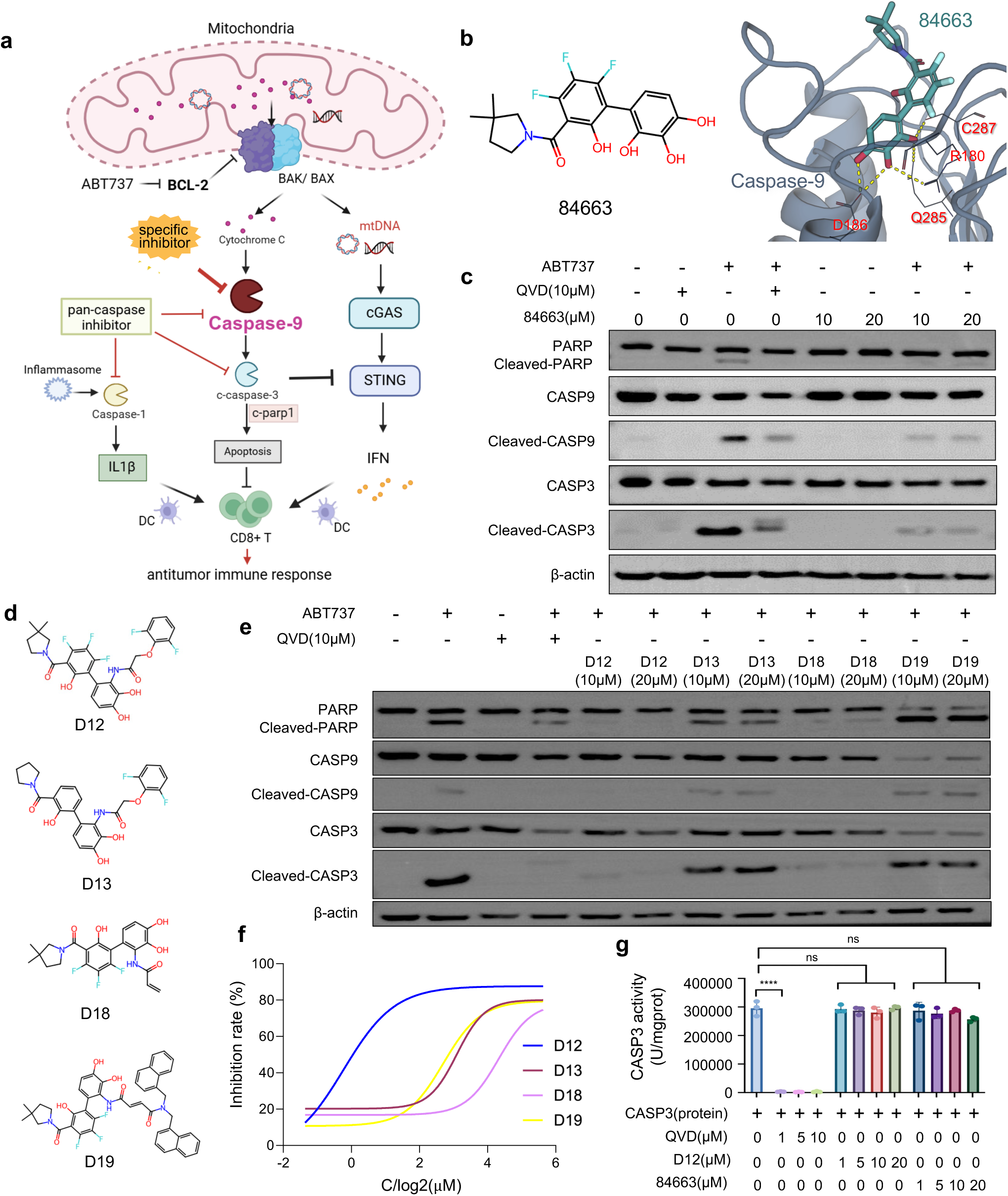
Caspase-9 inhibitor design. **a**, The diagram of the caspase-9 related pathway. **b**, The designed molecule 84663 that was validated to bind with caspase-9 (left) and the generated binding pose by PocketXMol (right). The yellow dash is the polar contact identified by PyMOL. The contact residues are annotated with residue names and indices. **c**, Western blot analysis of the expression of PARP and caspase-9/3. MC38 cells were treated with QVD (10µM and 20µM) or 84663 (10µM and 20 µM), combined with ABT-737 (10 µM) or not. The term “CASP” is the gene name of “caspase” **d**, The second-round molecules validated to bind with caspase-9 (D12, D13, D18, and D19). **e**, Western blot analysis of the expression of PARP and caspase-9/3. MC38 cells were treated with different caspase inhibitors (10µM and 20µM) combined with ABT-737 or not. **f**, Caspase-3 activity assay was done to determine the EC_50_ values of the molecules for suppressing the ABT-737 induced caspase-9 mediated caspase-3 activation. **g**, Analysis of the activity of caspase-3 from the mix of caspase-3 protein (0.5ng/ml) and caspase inhibitors by caspase-3 assay kit.

Using the caspase-3 function assay, we found that the four molecules could inhibit ABT-737-induced, caspase-9-mediated caspase-3 activation with EC_50_ values ranging from 0.85 to 8.56 µM (Figure 3f). Among these, the molecule D12 demonstrated an EC_50_ much better than 84663, and were even comparable to Z-LEHD-FMK TFA, a commercial caspase-9 inhibitor (Figure S19). To further verify that the designed molecules exert their effects by targeting caspase-9 rather than caspase-3, we conducted an in vitro caspase-3 enzymatic activity assay (Figure 3g). The results showed that, despite their upstream inhibitory effects, neither D12 nor 84663 directly inhibited the catalytic activity of recombinant active caspase-3. This contrasts with the pan-caspase inhibitor QVD, which completely abolished caspase-3 enzymatic activity. These findings suggest that the designed molecules likely inhibit apoptosis by directly binding to caspase-9, thereby blocking its apoptosome-mediated activation and preventing subsequent cleavage of caspase-3, a mechanism distinct from direct caspase-3 inhibition. To characterize the direct binding interactions between caspase-9 and the novel inhibitors D12 and 84663, we employed surface plasmon resonance (SPR) biosensor analysis, as co-crystallization attempts were unsuccessful due to time-dependent proteolytic degradation of caspase-9^79^. Based on the predicted binding structures (Figure 3b), we introduced alanine substitutions at key contact residues (C287, R180, Q285, and D186) to assess changes in binding affinity (Table S9). With the exception of D186A, which failed to express, all other mutants exhibited reduced binding affinities compared to the wild-type protein, supporting the involvement of these residues in ligand interaction. Notably, C287, the catalytic residue of caspase-9, was essential for ligand binding; its alanine mutant (C287A) showed a thirty-fold decrease in binding affinity to 84663, confirming the binding site and explaining the compound’s inhibitory activity.

### Performance on peptide design

With the atom-level representation of PocketXMol, peptide design exhibits no inherent distinction from the small molecule design. The primary challenge lies in how to precisely control the model to generate chains of amino acids instead of arbitrary molecules. Based on the fact that the backbone atoms remained consistent across peptides with differences primarily stemming from their side-chain atoms, PocketXMol can regard peptide generation as a special case of fragment growing, wherein the side-chain atoms are generated based on the peptide backbone and the binding protein pocket. Specifically, given the length of the peptide, PocketXMol generates new atoms (and bonds) based on the backbone fragment and the binding pocket to form a complete molecule, and these newly generated atoms constitute the side chains, from which the amino acid types can be annotated (Figure 4a). This turns the design of peptides’ structures and sequences into the end-to-end design of molecular fragments, which distinguishes PocketXMol fundamentally from existing peptide design methods where they explicitly model the amino acid types during designing. Additionally, PocketXMol simultaneously generated all atom positions, eliminating the need for additional docking or side-chain packing using separated post-processing steps.

**Figure 4.**
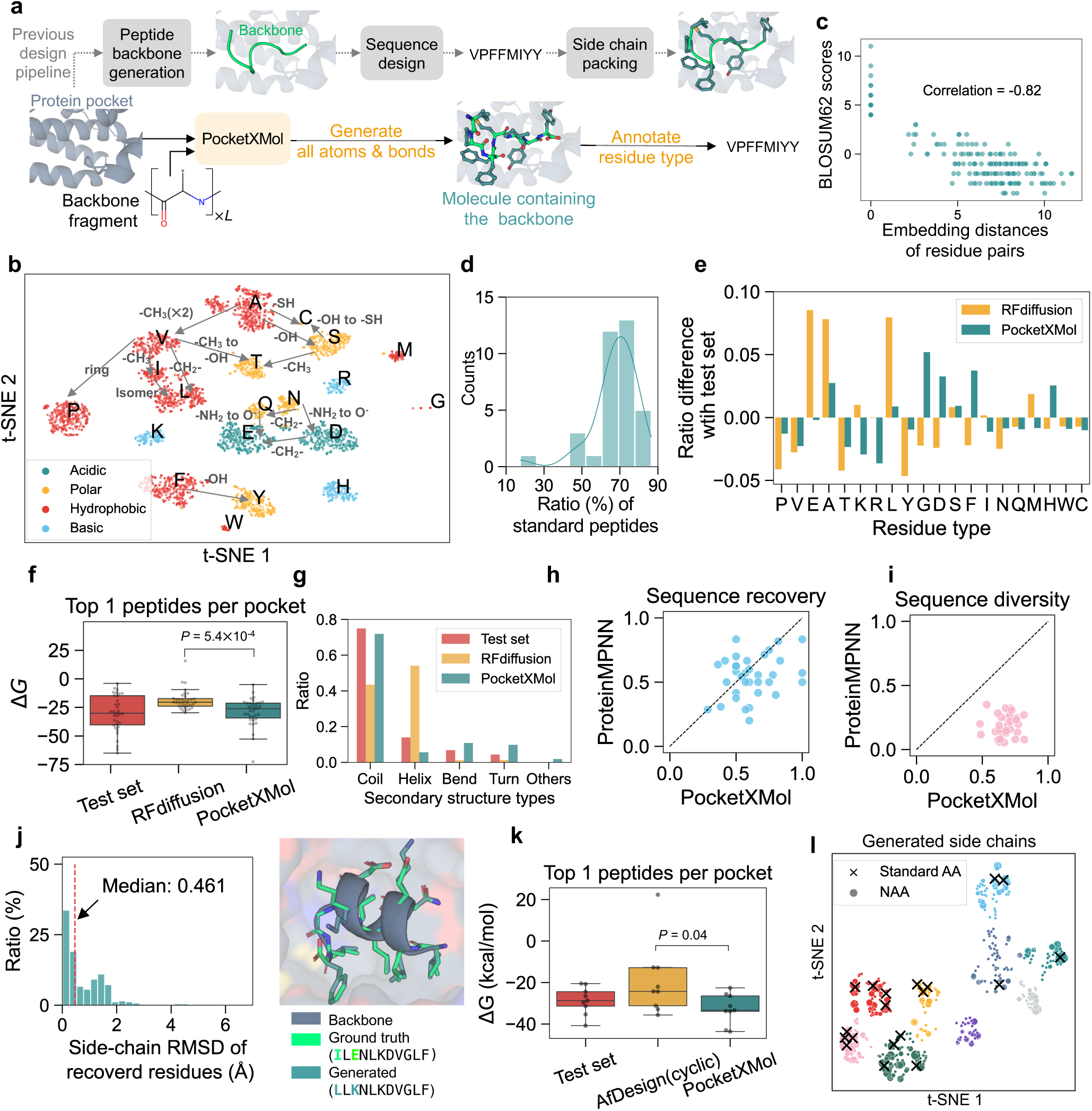
Performance on peptide design. **a**, Differences between PocketXMol and the previous peptide design pipeline. **b**, Relationship between BLOSUM62 substitution scores and embedding distances for each pair of amino acids. **c**, t-SNE visualization of the amino acid embeddings. The embeddings were calculated as the average of the side-chain atoms’ embeddings at the last layer of the denoiser neural network (embeddings of glycine were set as all-zero vectors since their side chains do not contain heavy atoms). The differences in the side chains between similar amino acids were annotated. **d**, Ratio distribution of standard peptides (i.e., peptides containing only standard amino acid side-chains) generated by PocketXMol for each pocket in the test set. **e**, Ratio differences of amino acid types between the generated peptides and the test set. **f**, Distributions of the Rosetta binding energy for the test set, RFdiffusion, and PocketXMol. The binding energy was calculated using the Rosetta Interface Analyzer, and the peptides with the lowest Rosetta energies for each pocket were selected for comparison (one-sided paired *t*-test with *n* = 35). **g**, Ratio of secondary structures in the peptides from the test set, RFdiffusion and PocketXMol. **h-i**, Sequence recovery rate and sequence diversity for ProteinMPNN and PocketXMol on the inverse folding task. For both methods, the peptides with the highest sequence recovery among the 100 generated samples per pocket were selected for comparison. The sequence diversity was calculated as *1-average sequence similarity* among the generated peptides for each pocket. **j**, RMSD distribution of the side-chain atoms whose amino acid types were recovered by PocketXMol in inverse folding (left), and an example showing the structure and the sequence of the generated peptide (right). **k**, Distribution of the Rosetta binding energy for the test set, AfDesign(cyclic), and PocketXMol for cyclic peptide design. The binding energy was calculated using the Rosetta Interface Analyzer, and the peptides with the lowest Rosetta energies for each pocket were selected for comparison (one-sided paired *t*-test with *n* = 9). **l**, t-SNE visualization of side chains of all generated amino acids by PocketXMol for the peptide design task. The topological fingerprints were calculated for all side-chain fragments extracted from generated peptides. Then t-SNE was applied to the fingerprints, followed by K-means clustering (*K* = 9) on the t-SNE space. The cross markers represent the standard amino acids, and the circle markers are generated NAAs. The size of the circles represents the counts, and the color represents the cluster.

#### Linear peptide design

To validate this novel peptide design strategy, we built a peptide design test set by collecting 35 protein-peptide complexes from the PepBDB docking test set^40^ and utilized PocketXMol to design peptides for each protein pocket (Supplementary Notes 4.3.10). Firstly, to ascertain whether PocketXMol learned that atoms could form amino acids, we retrieved the average embeddings of the side-chain atoms of the generated peptides and projected the amino acid embeddings into two-dimensional space using t-SNE (Figure 4c). The amino acid clusters correspond precisely to the types of amino acids with an Adjusted Rand Score of 0.813. The learned embeddings reflect chemical similarities among side-chain atoms, positioning amino acids with similar side chains in closer proximity. Moreover, the learned embeddings contain the structured semantic information of side chains. In the embedding space, the difference between glutamic acid (E) and glutamine (Q) is approximately equal to the difference between aspartic acid (D) and asparagine (N), both of which correspond to replacing an amine group -NH_2_ with an oxhydryl group -OH. Additionally, the distances between the amino acid embeddings exhibited a strong correlation with the BLOSUM62 substitution scores, with a Pearson correlation of -0.82 (Figure 4b). Since the amino acid types were inferred from the side chains, a potential concern is that the side-chain atoms may not always constitute standard amino acids. However, in practice, we observed that the ratios of generated peptides containing all standard side chains (referred to as standard peptides) are acceptable for various pockets and peptide lengths (Figure 4d). For the subsequent experiments, we only considered and retained these standard peptides.

Next, we compared PocketXMol with the state-of-the-art protein/peptide design pipeline, which utilizes RFdiffusion^13^ to generate backbones, followed by the sequence generation by ProteinMPNN^12^ and side-chain packing by Rosetta^58^ (referred to as RFdiffusion pipeline). For both RFdiffusion and PocketXMol, we generated at most 100 standard peptides for each protein target in the test set and assessed the quality of the generated peptides. First, we analyzed the amino acid types and observed that the distribution of types generated by PocketXMol was much closer to the distribution of the test set than the RFdiffusion pipeline, with a Jensen–Shannon divergence of 0.16 for PocketXMol and 0.23 for the RFdiffusion pipeline (Figure 4e). The average sequence recovery rate of PocketXMol reached 37.2%, better than 32.1% of RFdiffusion (Figure S20a). For 3D structural consistency, we calculated the backbone RMSD of each generated peptide to the reference peptide in the test set. PocketXMol generated peptides with smaller backbone RMSDs than RFdiffusion (Figure S20b). Moreover, we calculated the binding energy Δ*G* between the proteins and peptides using Rosetta^58^ and found that PocketXMol generated peptides with superior Rosetta binding energy compared to RFdiffusion, which can be attributed to directly modeling interactions at the atom level instead of the interactions between amino acids (Figure 4f). We also observed that the RFdiffusion pipeline generated peptides predominantly featuring helix structures, while PocketXMol produced mostly coils with a minority of helices, bends, and turns, which more closely resembled the distribution of peptides’ secondary structures in the test set (Figure 4g). Finally, we employed MolProbity, a widely used tool for diagnosing issues in 3D polymer structures, to assess the quality of generated structures of peptide containing all standard amino acids^80^. PocketXMol produced a high proportion of peptides that satisfied most key MolProbity quality metrics (Table S10), and its self-confidence scores showed correlation with MolProbity evaluations (Table S11), suggesting that the confidence scores provided by PocketXMol can serve as a reliable indicator for identifying peptides with physically realistic conformations. All these results demonstrate the strong performance of PocketXMol in *de novo* peptide design. To further support this conclusion, we conducted similar analyses on the Q-BioLiP peptide test set, which comprised 57 protein–peptide pairs with low similarity to those in the PepBDB test set (Figure S21). The results remained consistent, further confirming the effectiveness and generalizability of PocketXMol(Figure S22).

Similar to the SBDD task, we evaluated PocketXMol’s robustness to variations in input pocket structures by generating peptides using pocket conformations from alternative sources. The results showed that most evaluation metrics remained generally consistent with those obtained using the experimentally determined holo structures (Figure S23). The peptide backbone RMSD values were slightly higher when using AlphaFold predicted structures, PDB-retrieved structures, or apo PDB-retrieved structures compared to using the experimental holo structures. However, as this metric relies on aligning the pocket backbones, it is inherently influenced by variations in the input pocket structure itself. The predicted Rosetta binding energies were slightly less favorable when using apo structures, potentially reflecting differences in surface properties compared to ligand-bound (holo) structures.

#### Peptide inverse folding

Peptide inverse folding is a generative task involving the design of peptide sequences based on the structures of the peptide backbone and protein pocket^12^. To assess PocketXMol’s performance on this task, we removed the side chains of peptides in the PepBDB test set while retaining the peptide backbone structures and protein pockets and employed PocketXMol and the baseline ProteinMPNN^12^ to design 100 standard peptides for each pocket, respectively. PocketXMol demonstrated a significantly higher sequence recovery rate of 61.7% than 54.4% for ProteinMPNN (one-sided paired *t*-test, *P* = 0.02*, n* = 35), indicating the effectiveness of directly generating and placing the side-chain atoms within the pocket, as opposed to merely classifying amino acid types based on the backbone structures as ProteinMPNN did (Figure 4h). PocketXMol also exhibited significantly higher sequence diversity compared to ProteinMPNN (one-sided paired *t*-test, *P* = 3.18 × 10^−24^*, n* = 35), due to the fact that predicting side-chain atoms is less prone to overfitting compared to merely predicting amino acid types during training (Figure 4i). Moreover, the side-chain conformations were simultaneously determined by PocketXMol during the generation process. The atoms of the residues recovered by PocketXMol are consistent with the reference structure, indicating its robust ability to generate accurate complete peptide structures (Figures 4j and S20c).

#### Cyclic peptide design

Thanks to its atom-level modeling, PocketXMol can be directly applied to cyclic peptide design without explicit training on such peptides. To evaluate this capability, we curated a test set of 26 protein–cyclic-peptide complexes and clustered them into nine groups based on target sequence identity. One representative pocket from each cluster was selected for evaluation. For each pocket, we generated up to 100 cyclic peptides and compared PocketXMol with AfDesign(cyclic)^81^, a hallucination-based baseline method that adapted AlphaFold for cyclic peptide design. We assessed sequence recovery, backbone RMSD, Rosetta binding energy (ΔG), and secondary structure composition. PocketXMol achieved significantly better performance in terms of top-1 and top-10 Rosetta binding energies per pocket and performed comparably on other metrics (Figures 4k and S24, and Table S12). This advantage stems from PocketXMol’s ability to transfer atom-level interaction knowledge from small molecule and linear peptide data to the cyclic peptide setting, underscoring the strength of its atom-level modeling.

#### Design peptides with non-standard amino acids

Since PocketXMol directly generates side-chain atoms without explicitly predicting amino acid types, it naturally supports the generation of non-standard amino acids (NAAs) beyond the standard 20 ones. During the *de novo* linear peptide design task, we observed that PocketXMol generated both standard and non-standard amino acids (Figure 2d), and we now revisited the results to analyze the NAAs in detail. We extracted all peptides containing NAAs and identified 454 unique NAA side chains, varying in size, frequency, novelty, and physicochemical properties^82^ (Figure S25a and S27), revealing a wide spectrum from subtle modifications to entirely novel structures (Figures 4l and S26). We hypothesize that PocketXMol generates NAAs when alternative atomic arrangements enable more favorable interactions within the target pocket than standard residues. To investigate this, we mapped frequent NAAs to their most structurally similar standard amino acids and compared the Rosetta energies of the original NAA-containing peptides with their standard analogs. The NAA peptides exhibited comparable or better energies (Figure S25b,c), supporting their effectiveness and highlighting PocketXMol’s potential for designing peptides with non-standard amino acids.

#### PD-L1 binder design

To rigorously validate PocketXMol’s design capability, we employed it to design peptides targeting PD-L1 and experimentally confirmed their binding affinities and specificities. The peptide design pipeline involved generating a large number of candidate sequences using PocketXMol followed by a filtering process to identify top-ranked peptides. Specifically, peptides were generated for three PD-L1 pocket structures (PDB IDs: 3BIK, 4ZQK, and 5IUS), followed by thorough filtering and ranking using several scoring tools, including self-confidence, FoldX^83^, Rosetta^58^, and AlphaFold^6, 57^ (Supplementary Notes 8.2). Finally, 382 peptides were selected for validation of their binding with PD-L1 using SPRi experiments. The results revealed that 15 peptides achieved the dissociation constant (*K*_D_) of 10^−8^M and 76 peptides achieved *K*_D_ of 10^−7^M (Figure 5a). As a baseline, we constructed a random peptide library with a capacity of 10^7^ to experimentally screen peptides targeting PD-L1, which only resulted in 8 positive peptides with *K*_D_ values of or close to 10^−8^M. This sampling-and-filtering pipeline based on PocketXMol demonstrated a significantly higher success rate than the baseline screening approach. To further assess PocketXMol’s intrinsic sampling capability independent of filtering, we experimentally validated an additional set of 382 peptides generated without any post-generation ranking. Among these, 9 peptides exhibited high binding affinity (*K*_D_ of 10^−8^M), confirming PocketXMol’s strong ability to directly sample high-quality and target-specific peptides (Table S13).

**Figure 5.**
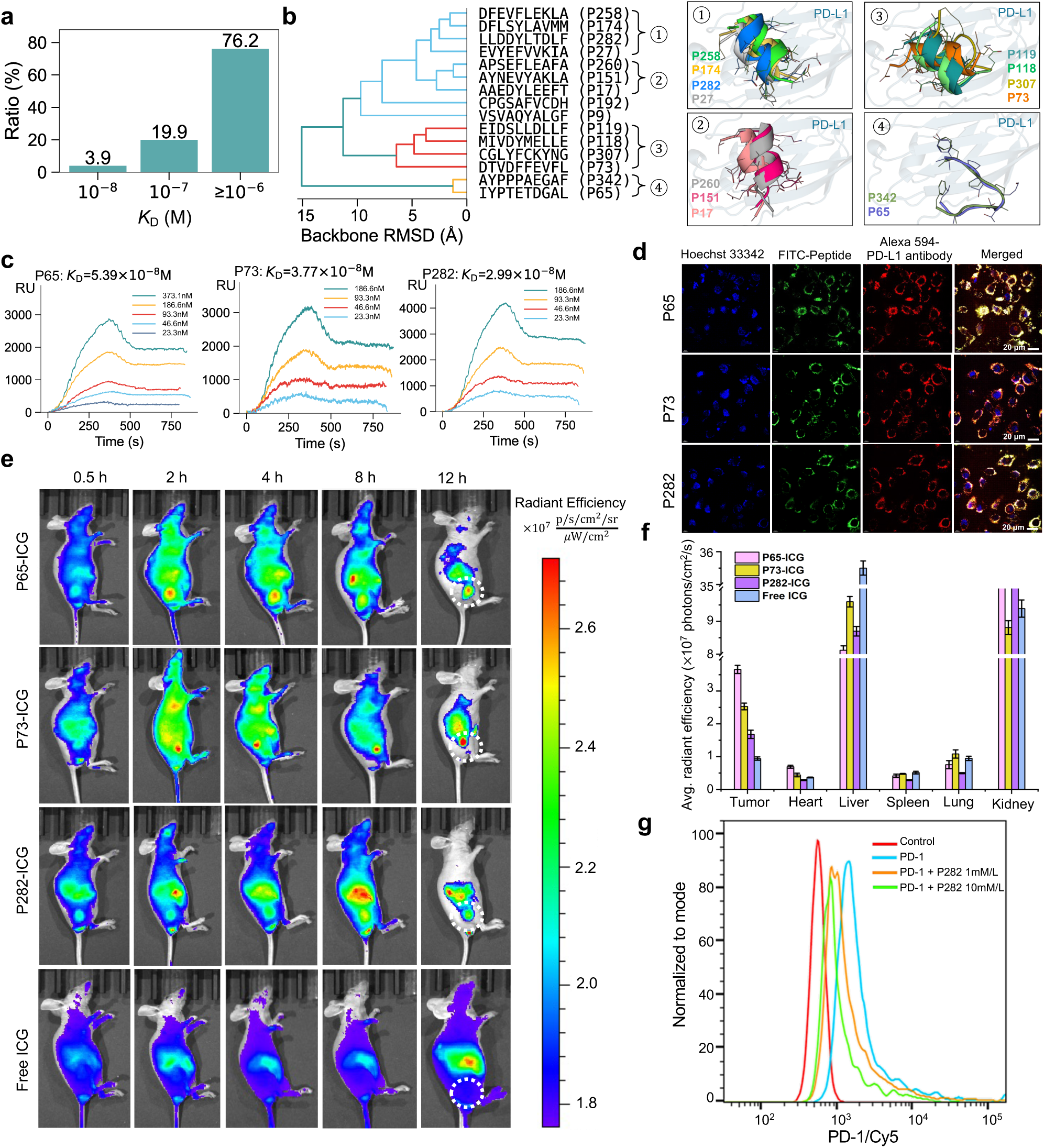
PD-L1 binding peptide design. **a**, Ratio of designed peptides with different disassociation constant *K*_D_ ranges with PD-L1. The dissociation constants were obtained using the SPRi experiments. **b**, Structure clusters of the 15 generated peptides with *K*_D_ of 10^−8^M, with the structures of peptides in four major clusters. The hierarchical clusters were constructed based on the backbone RMSDs among peptides and used the maximum distances between two sets as the linkage criterion. **c**, The SPRi binding curves of the generated peptides selected for further experiments (P65, P73, and P282) with PD-L1. **d**, Confocal image of FITC labeled candidate peptides binding toward the positive cell H1975. The negative cells 293T are shown in Figure S28. Scale bar was 20 µm. Hoechst 33342 represents the nucleus, blue; FITC represents the peptide, green; Alexa Fluor 594 represents the PD-L1 protein, red. **e**, In vivo fluorescence imaging was conducted on H1975 xenograft model mice at various time points after intravenous injection of P65-ICG, P73-ICG, P282-ICG, or free ICG, each administered at a dose of 50 µM. The tumors are indicated by white circles. **f**, The average fluorescence signal intensity of isolated organs at 12 hours post-injection. Bars with extremely high values are truncated. **g**, Flow cytometry analysis to investigate the PD-1/PD-L1 interaction using different concentrations of P282 peptide.

For the 15 peptides with *K*_D_ of 10^−8^M obtained from the sampling-and-filtering pipeline, we clustered the generated peptide structures (Figure 5b), which demonstrated both the diversity and the consistency of the peptides. We presented three examples of the peptide binding curves and the interaction profiles in Figures 5c and S32a, which depicted the rich interactions between the PD-L1 pocket and the designed peptides.

Next, we selected three peptides with high affinity and representative structures (P65, P73, and P282) and conjugated them with FITC (Fluorescein isothiocyanate) fluorescent dye, aiming to validate their affinity and specificity at the cellular level. After co-incubating the three fluorescent-labeled peptides with PD-L1 positive cells H1975, the peptides effectively bound to the cell membrane, emitting strong green fluorescence. Co-localization staining with PD-L1 antibodies further confirmed that the peptides were specifically bound to PD-L1 proteins on the cell membrane surface (Figure 5d). In contrast, when these peptides were co-incubated with PD-L1 negative 293T cells, there was negligible green fluorescence observed on the cell membranes (Figure S28), which demonstrated the robust affinity and specificity of the designed peptides targeting PD-L1.

To further assess the in vivo targeting efficacy of the PD-L1 peptides, we conducted in vivo imaging experiments using indocyanine green (ICG), an FDA-approved dye commonly used for fluorescence imaging in vivo. The peptides conjugated with ICG were administered at a concentration of 50 µM via the tail vein in a PD-L1 positive tumor model with the non-small cell lung tumor. As depicted in Figure 5e, P65, P73, and P282 exhibited significant accumulation at the tumor sites within 0.5h post-injection, with the fluorescence signal gradually increasing over time and peaking at 4h. In contrast, no signal was observed at the tumor site in the control group injected with free ICG, indicating that peptides modified with ICG can accurately target tumors expressing high levels of PD-L1. After 12 hours, we removed the heart, liver, spleen, lungs, kidneys, and tumor tissues from each group of mice for ex vivo imaging. As shown in Figures 5f and S29, there was still a strong signal at the tumor site 12 hours after injection in the peptide probe group, while there was almost no signal at the tumor site in the control group. The liver and kidneys of each group show higher non-specific accumulation due to peptide degradation and free ICG accumulation during circulation and metabolism through the liver and kidneys. As a comparison, we also conducted the same experiments for two positive peptides obtained through library screening (Figures S30 and S31). These peptides failed to sustain durable in vivo tumor targeting and were therefore excluded from subsequent quantitative analyses. These results demonstrate that the peptides designed by PocketXMol have excellent targeting and selectivity in vivo and can be used for targeted tumor imaging and diagnosis.

To determine whether the designed peptides can inhibit the interaction between PD-1 and PD-L1 and exert therapeutic effects, we conducted a ligand inhibition assay for the peptide P282 using flow cytometry analysis (Figure 5g). In the absence of the peptide, PD-1 could effectively bind to PD-L1. However, in the presence of the peptide, the binding of PD-1 was reduced. As the concentration of the peptide increased, the PD-1-Cy5 fluorescence signal further diminished. These results indicate that the peptide can inhibit the interaction between PD-1 and PD-L1 to some extent and has the potential in cancer immunotherapy.

To validate the predicted binding structures of peptides P65, P73, and P282, we performed alanine-scanning mutagenesis on key PD-L1 residues identified from PocketXMol-generated structures and measured binding affinities via SPR (Figure S32). For P65 and P73, most or several mutations caused over a 10-fold drop in affinity, supporting the predicted binding interfaces. For P282, one critical mutation showed a strong effect, while others had limited impact, likely due to the structural stability of the alpha-helix conformation of P282. These results support the structural validity of the PocketXMol-designed complexes.

### Performance on molecular structure generation

#### Small molecule docking

We evaluate the performance of PocketXMol on small molecules docking using the PoseBusters benchmark set^55^ that contains 428 protein-small-molecule complex structures. The setting is to dock the flexible ligand into the rigid and fixed protein pocket, consistent with the common docking scenarios^55^. Following the PoseBusters benchmark setting, all baselines provided one docked pose per protein according to their default settings^7, 11, 34, 54, 55, 84–89^. PocketXMol also provided one pose for each pocket, by selecting the top one pose from 100 generated docked poses sorted by the ranking scores (details in Supplementary Notes 4.3.1). The ranking scores were mainly based on the confidence scores produced together with the generated poses by PocketXMol (self ranking), and we also provided a confidence predictor tuned from the original PocketXMol to facilitate better ranking (tuned ranking). We calculated the atom RMSDs of the docked molecular poses and measured the ratio of RMSD *<* 2°A among 428 proteins. PocketXMol exhibited accurate docking performance, with 82.5% and 83.4% of poses achieving RMSD *<* 2°A with self ranking and tuned ranking, only second to the recent AlphaFold 3 with defined pockets (Figure 6a and Table S14). The RMSDs of PocketXMol exhibited correlations with the ranking scores (Figures 6b and S33a) and increasing the number of selected poses for each pocket also consistently enhanced the performance (Figure S33b). Moreover, if the best poses among the generated poses per protein could be precisely selected, i.e., with oracle ranking scores, the performance can be boosted to 96.5%, underscoring the robust generation capability and substantial room for improvement with more advanced ranking methodologies. We also observed no correlation between the RMSDs and molecular properties, confirming no bias toward specific molecule types (Figure S34).

**Figure 6.**
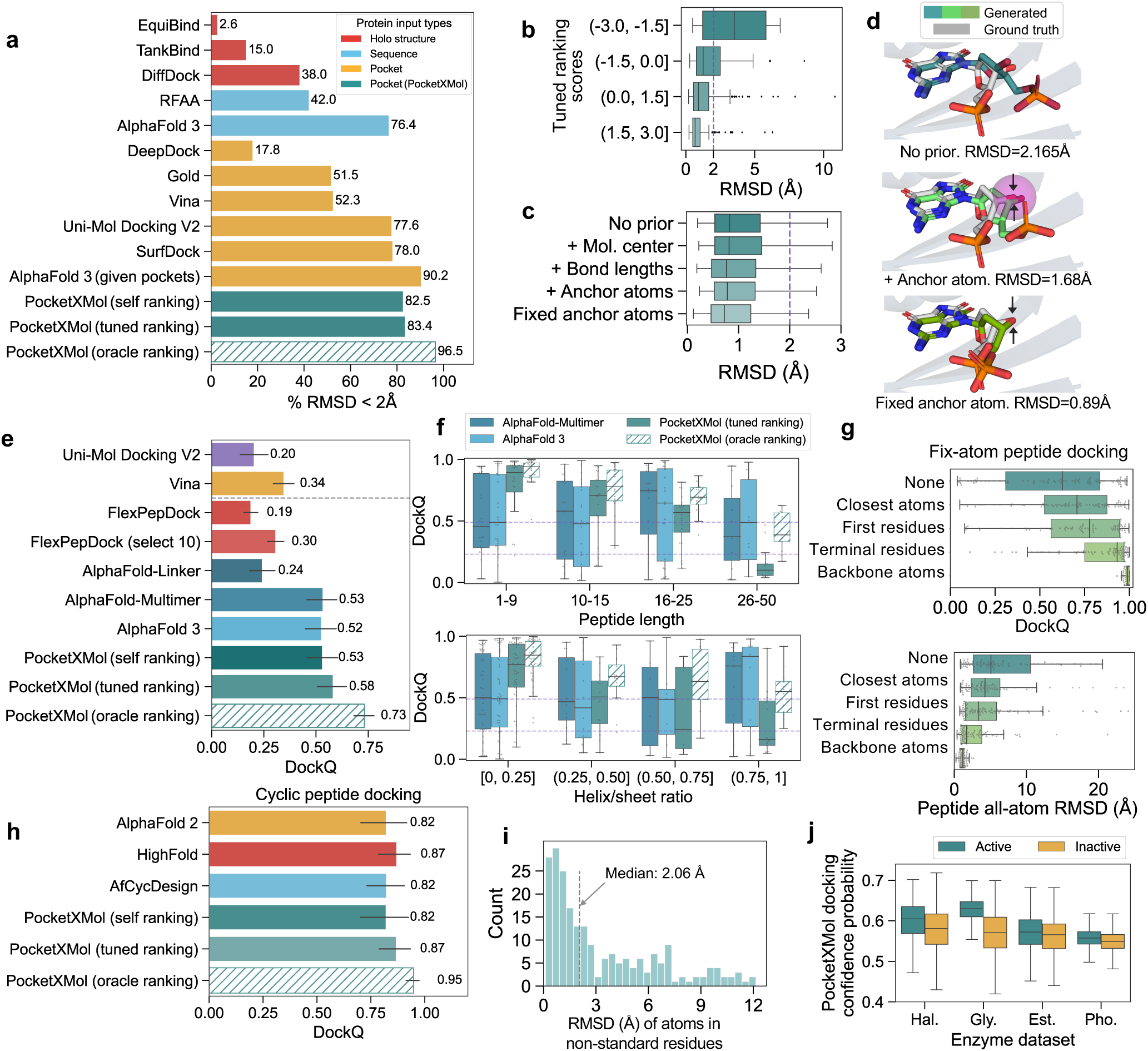
Performance on molecular structure generation. **a**, Ratio of docked poses with RMSD *<* 2°A by different models for protein-small-molecule docking. The numbers on the bars are the ratio values. **b**, The relationship between PocketXMol’s tuned ranking scores and the RMSDs for the docked poses. **c**, The RMSD values by incorporating different types of prior knowledge using PocketXMol for protein-small-molecule docking. **d**, An example showing that docking can be improved by incorporating prior knowledge. The top, middle, and bottom structures represent the generated docking poses with no prior information, with the approximated location of the anchor atom, and with the exact coordinate of the anchor atom, respectively. The arrows indicate the anchor atoms and the purple sphere represents the approximated locations of the anchor atom. **e**, DockQ of protein-peptide docking by different models. Error bars represent 95% confidence intervals (*n* = 79). **f**, Relationship between DockQ for peptide docking and the peptide length (left) or the ratio of secondary structures helix and sheet (right). The box plot outlines quartiles of the metrics with whiskers spanning up to 1.5 times the interquartile range (*n* = 79). Here we only displayed the best four methods ranked in panel E. **g**, DockQ and RMSD of protein-peptide docking by PocketXMol with different types of prior knowledge. The box plot outlines quartiles of the metrics with whiskers spanning up to 1.5 times the interquartile range (*n* = 79). **h**, DockQ scores for cyclic peptide docking across different models. For each cluster, DockQ values were averaged over all samples within the cluster, and the final values represent the mean across clusters. **i**, RMSD for atoms in non-standard amino acids. **j**, Distributions of PocketXMol docking confidence scores for enzyme-substrate pairs stratified by whether they are active or not. The x-axis is the abbreviation of the enzyme family, where Hal., Gly., Est., and Pho. are Halogenase, Glycosyltransferase, Esterase, and Phosphatase, respectively. *P* values of one-sided unpaired *t*-test from left to right are 9.8 × 10^−13^, 2.9 × 10^−150^, 1.5 × 10^−26^, and 3.0 × 10^−67^, respectively, with *n* from left to right being 323, 2281, 837, 3460, 2786, 9886, 5424, and 30546, respectively.

For a more rigorous evaluation, we employed the PoseBusters test suite^55^ as a docking validity checker. It employs a series of criteria to assess both the chemical and geometric coherence of 3D molecular structures, categorizing poses as PB-valid only when all the criteria are satisfied. We measured the ratio of poses satisfying both RMSDs *<* 2°A and being PB-valid. PocketXMol also achieved the best performance with 78.5% and 79.4% poses for self-ranking and tuned ranking, respectively, among the 428 samples in the PoseBusters set (Figure S33c and S35). PocketXMol was compared with AlphaFold 3 on the PoseBusters Version 2 set (containing 308 test samples) and achieved 84.7% and 84.4% for self ranking and ranker scores, respectively, comparable to 84.4% of AlphaFold 3 with defined pockets as reported in its paper (Figure S33d).

To evaluate PocketXMol’s robustness to variations in pocket structures, we performed docking using pocket conformations from alternative sources. By design, PocketXMol strictly adheres to the input pocket coordinates when placing ligand atoms and does not adjust for inaccuracies in pocket atom positions. For example, if pocket atoms occupy the space where the true ligand should reside, PocketXMol avoids placing ligand atoms there due to steric clashes. To enhance tolerance to input structure variations, we finetuned the original PocketXMol on perturbed pocket structures for docking tasks (Supplementary Notes 5.4). We referred to this finetuned version as PocketXMol-PF (PocketXMol with Pocket-Flexible adaptation) and evaluated it for docking tasks using structure variants. For pose selection of PocketXMol-PF, we relied on the model’s self-confidence score, avoiding the need for finetuning an additional ranking model. when provided with either the true experimental structures or Rosetta-repacked structures, PocketXMol-PF achieved performance comparable to the original PocketXMol with true structures (Figure S36b,c), although the original PocketXMol with Rosetta-repacked structures achieved weaker performance than original PocketXMol with true structures (Figure S36a). This indicates that the PocketXMol-PF can reliably predict the docking poses with side-chain variations. When tested on input structures derived from AlphaFold predictions, alternative PDB-retrieved crystal structures, or PDB-retrieved apo structures, the resulting docked ligand RMSDs must be interpreted alongside pocket alignment accuracy. Since ligand RMSD is computed after aligning the docked pocket to the true structure’s backbone, it captures both ligand pose errors and discrepancies in backbone alignment. To disentangle these effects, we plotted ligand RMSDs against the corresponding pocket backbone RMSDs (Figures S37 and S38). The results showed that PocketXMol-PF maintained high docking accuracy with alternative pocket structures, provided the backbone deviations are moderate. In cases with larger pocket backbone RMSDs, higher ligand RMSDs are observed, but much of this increase may stem from backbone alignment discrepancies and does not necessarily reflect a failure in the docking prediction itself.

#### Small molecule docking with prior knowledge

It is common to incorporate prior knowledge into the docking process during the drug design. For instance, when docking a new molecule extended from a fragment hit, it is usually assumed that the fragment pose remains unchanged in the new molecule^64^, which necessitates fixing the fragment positions throughout the docking process. Our flexible framework can efficiently integrate prior knowledge by either editing the molecules before adding noise at each generation step or directly employing the task prompt to express the constraints. Here, we showcased docking with different prior knowledge, including known molecular center positions, bond lengths, and approximate or exact coordinates of anchor atoms. The findings highlighted the effectiveness of various forms of prior knowledge in decreasing the docking RMSD (Figure 6c). Notably, utilizing the exact coordinates of anchor atoms yielded the most performance improvement, attributed to the substantial reduction in freedom of molecular poses (Figure 6d).

#### Application on enzyme-substrate interactions

We applied PocketXMol to dock small-molecule substrates into enzyme proteins (Table S18) and investigated whether its self-confidence scores could discriminate active from inactive enzyme–substrate pairs (Supplementary Notes 7). PocketXMol’s docking confidence scores exhibited strong discriminative power across multiple enzyme families (Figure 6f), significantly outperforming AlphaFold 3 in most cases (Figures S48 and S49). Furthermore, by extracting structural representations from PocketXMol and training simple logistic regression classifiers, we further enhanced activity prediction accuracy (Figure S50). Finally, PocketXMol generated enzyme–substrate complex structures in which the substrates, especially their reactive groups, were more frequently positioned near catalytic residues, suggesting that PocketXMol produced catalytically plausible docking poses (Figure S51).

#### Molecular conformation generation

PocketXMol can also generate valid 3D conformations for small molecules without the consideration of protein pockets. Following the test settings of previous works^45, 90, 91^, PocketXMol was benchmarked with software specifically designed for 3D conformation prediction and exhibited comparable performance to most of these tailored methods (Table S15). An example is shown in the Figure S39, demonstrating that PocketXMol can generate almost identical conformations to the low-energy ones.

#### Linear peptide docking

Current peptide docking methods are primarily constructed at the amino acid level^92, 93^. In contrast, PocketXMol leverages atom-level representation and does not distinguish between peptides and small molecules, regarding the peptide docking exactly the same as small-molecule docking. We evaluated PocketXMol’s performance against peptide docking approaches, including the AlphaFold series^7, 57, 94^ and the energy-based method FlexPepDock^92^, on 79 protein-peptide complexes compiled from a previous peptide docking benchmark set^93, 95^. We also included two representative small-molecule docking tools, Uni-Mol Docking V2^87^ and Vina^54^ as baselines by treating peptides as small molecules. The setting is to dock the peptide into the rigid and fixed protein pocket. Similar to the small molecule docking, PocketXMol selected one docked peptide structure per protein pocket from 100 generated ones according to the ranking scores. PocketXMol achieved satisfying performance with an average DockQ^96^ of 0.53 and 0.58 for self ranking and tuned ranking, respectively (Figure 6e and Table S16). The performance of small-molecule docking tools was much lower than PocketXMol (Figure S41), indicating that current docking tools did not have the transferability between small molecules and peptides like PocketXMol did. We observed that the accuracy of PocketXMol can also be consistently enhanced by selecting more structures from the generated ones (Figure S40a), and the average DockQ score could reach 0.73 if the oracle ranking scores were available, highlighting considerable opportunities for enhancement through more advanced ranking strategies. When considering the peptide properties, PocketXMol can achieve much higher accuracy on peptides with shorter lengths or lower proportions of helix/sheet secondary structures (Figure 6f), which is explained by the fact that these peptides exhibited more flexibility and were more similar to small molecules. Notably, peptides longer than 15 were not included in our training dataset, but they could still be docked with modest accuracy, highlighting PocketXMol’s extrapolation and generalization abilities. Additionally, we also benchmarked with AlphaFold-Multimer^57^ and AlphaFold 3^7^ on another test set comprised of the latest protein-peptide complexes collected from the Q-BioLiP database^97, 98^, and the conclusions were the same as the above ones (Figure S42).

Furthermore, owing to the atom-level representation, PocketXMol can naturally model peptides with modified amino acids. There are 13 peptides containing nonstandard amino acids in the test set, and all of them were successfully docked with good accuracy (DockQ≥0.49) and showed small RMSDs for those atoms in the nonstandard amino acids (Figure 6i and S40c). As a comparison, AlphaFold-Multimer used the closest standard amino acids to replace these nonstandard ones, and only six among the 13 peptides achieved DockQ≥0.49 (Figure S40b). PocketXMol offers a more expansive scope for real biological applications because modifications are commonly utilized in peptide drug design to enhance their stability and prolong the half-life^15^.

Similar to small molecule docking, we also assessed PocketXMol’s performance on peptide docking to pocket structures from alternative sources, using both the original PocketXMol and the PocketXMol-PF model. Both models exhibited consistent performance, demonstrating robustness to input structure variations (Figures S43 and S44). Notably, PocketXMol-PF showed a slight performance advantage over the original PocketXMol for longer or more rigid peptides (Figures S45, S46, and S47).

#### Peptide docking with prior knowledge

For peptide docking, there are also cases when the positions of certain atoms or residues of the peptides are known in advance^99–101^, necessitating integrating such prior knowledge into the docking process. Here, we showcased docking peptides while fixing certain atoms at various positions, including 1) the atom that was the closest to the protein (named anchor atom), 2) atoms in the first residue (N-terminus), 3) atoms in the two terminal residues (N- and C-terminus), and 4) all backbone atoms. We observed that the performance on docking could be enhanced to different extents with prior knowledge (Figure 6i). In particular, fixing only one anchor atom could yield an improvement of the median DockQ from 0.62 to 0.71. Fixing the atoms at the N-terminus and both N- and C-terminus could greatly enhance the performance to 0.78 and 0.93, respectively. These findings underscored the potential applicability of PocketXMol in protein loop modeling tasks, such as predicting the structures of antigen-binding antibody complementarity-determining regions (CDRs), which can be conceptualized as flexible peptides, with terminal amino acids linked to a relatively rigid framework^102^. A special case is fixing all the backbone atoms of the peptide and only determining the side-chain atoms, which is similar to side-chain packing. The RMSDs of the peptide side-chain atoms between the generated and the ground truth were small, with a median RMSD of 1.1 °A (Figure 6i) for all side-chain atoms.

#### Cyclic peptide docking

To evaluate PocketXMol’s performance on cyclic peptide docking, we compared PocketXMol against AlphaFold 2^6^, HighFold^103^, and AfCycDesign^104^, on the cyclic peptide test set that contained 26 protein-cyclic-peptide complexes of nine cluster groups based on target sequence similarity (Supplementary Notes 6.1). Following the same procedure of linear-peptide docking, we generated 100 docking poses for each pair of protein-peptide using PocketXMol, and we selected the pose ranked highest based on our ranking scores. The baseline was run using their default models and hyperparameters. We calculated DockQ of docked peptides and averaged them within each cluster. PocketXMol demonstrated performance comparable to the specialized baselines (Figure 6h and Table S17), despite not being specifically intended or explicitly trained for protein–cyclic peptide complexes.

## Discussion

To elucidate the fundamental atomic interactions and achieve the unification of molecular generative tasks, PocketXMol has introduced several methodological innovations with potentially profound impacts on the field. PocketXMol exclusively employs atom-level representation, eschewing advanced biological entities such as amino acids. This approach aligns with the core concept of molecular interactions, where all molecules are composed of atoms that interact through universally applicable mechanisms. Our experiments have demonstrated the feasibility of modeling molecules at the atomic level, showcasing the transferability among different molecular types. While some current molecular models can generate atom-level structures, they often treat different molecules using distinct entities. For example, AlphaFold 3^7^ represented proteins as amino acids, nuclear acids as nucleotides, and other molecules (e.g., modified amino acids and small molecules) as atoms and learned the interactions among these entities. However, a modified amino acid and its standard form may only differ in several atoms but are treated as entirely different representations in the model, which may hinder the model from learning the shared knowledge among different molecules. Another advantage of the atom-level representation is that any molecule can be directly modeled, avoiding the need for additional entity-specific representations, which is particularly beneficial for uncommon molecular types, such as macrocycles, peptoids, D-peptides, and modified peptides.

Secondly, we introduced a novel task representation that unifies diverse molecular tasks into a single computational framework, directly reflecting the relationships among tasks. The task representation bridges the gap among molecular tasks, especially for structure prediction and molecular designing. Recent advances in structure prediction models like AlphaFold 3^7^ and RFAA^11^ have made significant progress. Equipped with such a task representation, these successes can be directly extended to molecular designing without requiring additional fine-tuning or re-folding as current models do^13, 105^. Finally, our new generative framework is capable of handling multiple tasks with different prior noise distributions. It allows the denoiser to automatically learn noise characteristics, which is equivalent to performing task identification during training. While initially proposed for molecular tasks, this generative framework can be applied to other fields, enabling mixed noise types and tasks within a single model. For instance, in image generation, different perturbation strategies, such as simple Gaussian noise, heat dissipation, blurring, masking, and pixelating, have proven effective in single diffusion models^106–108^. It is potential to combine all these perturbations into a unified model with appropriate prompt mechanisms to enhance the generative model’s learning capabilities.

These innovations enable PocketXMol to treat peptides in a fundamentally different manner from previous models, providing a new paradigm for the field. By considering peptides as a special type of small molecule, PocketXMol docks peptides in the same way as small molecules and designs peptides through fragment growing. However, this does not imply that existing small-molecule docking or fragment-growing tools can be easily extended to peptides. Our results showed that small-molecule docking tools like AutoDock Vina^54^ and Uni-Mol Docking V2^87^ have limited performance on peptides. Designing peptides through fragment growing requires docking backbone fragments while generating side-chain atoms of standard amino acids, a capability that current fragment-growing tools lack. PocketXMol not only learns the commonalities between small molecules and peptides but also models their subtle differences.

## Limitations of the Study

While PocketXMol demonstrates strong performance across a broad range of pocket-interacting molecular tasks, several limitations should be acknowledged. First, PocketXMol is specifically designed for pocket-interacting tasks and does not extend to more general molecular interaction problems, such as protein folding or protein–nucleic acid docking, due to scalability constraints. Larger molecular systems, such as full proteins, contain far more atoms than the systems considered in this study, and the quadratic complexity of PocketXMol’s architecture with respect to atom count renders such applications currently infeasible. Second, the model requires predefined pocket structures as input and lacks the capability to jointly infer binding pockets along with the ligand, which may limit its applicability in scenarios where pocket definition is ambiguous or needs to be co-determined. Although the model has demonstrated robustness on predicted and non-native pocket structures, a more natural extension would be the ability to generate or refine pocket components, thereby enabling broader applications such as flexible-pocket docking and pocket design. Third, due to limited training data, the model does not support ligands containing rare element types or metal ions, restricting its chemical space. Lastly, further experimental validation, such as the design and testing of more complex ligands (e.g., cyclic peptides) and structural determination of binding poses via X-ray crystallography, would be valuable for substantiating the model’s generative capabilities and expanding its applicability.

## Author contributions

X.P. and J.M. conceived and designed the generative model with contributions from R.G. and J.G. X.P., R.G., J.G, and Y.J. processed the data. X.P., R.G., and J.G. conducted computational experiments. F.G, J.S, Y.X., Y.H, X.W, C.H, and Z.W designed and conducted the wet-lab experiments. X.P., F.G, Y.X, Y.H, M.Z., C.H, Z.W., C.H., and J.M. analyzed the results. X.P., C.H., Z.W., and J.M. wrote the manuscript with feedback and contributions from all authors.

## Methods

Ultimately, any generative molecular task can be understood as a process of generating the atom types, coordinates, and chemical bonds for a set of atoms conditioned on another set of atoms. In this work, we enhanced the capabilities of atom-level models beyond traditional modeling approaches through multiple new directions including constructing a universal denoiser, new task representations, new molecular representations, and a new training framework.

### The universal molecular denoiser

The core of PocketXMol is a universal denoiser parameterized by an E(3)-equivariant geometric neural network, which is applied to detect and eliminate noise from a 3D molecule. Let M represent a molecule, composed of a set of atoms with types and coordinates and chemical bonds with types, P be the task prompt, and *ξ* be the task-specific molecular noise. Assume Φ be a noise-adding process, a noisy molecule M̃ is produced as:

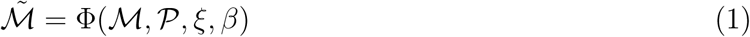

where *β* ∈ [0, 1] is a parameter controlling the noise scale. The universal denoiser *F*_Θ_ parameterized by Θ is designed to remove the noise from the noisy molecule, conditioned on the task prompt P and the protein pocket atom set K:

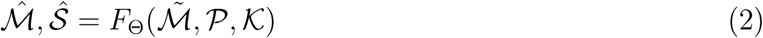

where M^^^ is the denoised molecule and S^^^ is the confidence scores for all variables of M^^^ . In contrast to diffusion models, Eq. 1 introduces additional terms P and *ξ*, enabling the generation to be controlled by the task prompt. Without the need to know the noise distribution *ξ* or scale *β* beforehand, the denoiser *F*_Θ_ removes the noise based on the prompt P and conditioned on pocket K, resulting in the creation of new molecules. These two functions work together to steer an initial molecule towards the one that conforms to the requirements specified by the prompt P and follows the distributions of real molecules.

### Atom-level model architecture

For a molecule (small molecule or peptide) with *N* atoms, the molecule M is composed of atom types **A**, atom coordinates **X** ∈ R*^N^*^×3^, and bond types **B**, i.e. M = {**A**, **X**, **B**}. The atom types **A** are *N* one-hot vectors of atom types including C, N, O, F, P, S, Cl, B, Br, I, and Se. The bond types **B** are *N* × *N* one-hot vectors representing bond types of atom pairs, including single, double, triple, aromatic bond, and none-bond indicating no covalent bond between the two atoms. For the protein pocket, denoted as K, it is also represented as sets of atoms with types and coordinates as above, and PocketXMol does not explicitly model amino acids.

A molecule encoder is first applied to process the features of the noisy molecule M̃ = {**A**, **X**, **B**}, the task prompt P and the pocket K. It builds a graph based on all atoms and utilizes graph neural networks to learn the latent features of atoms and atom pairs of the molecule, which are denoted as node and edge features in the neural networks, respectively:

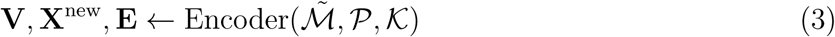

where **V** and **E** are the learned latent features of nodes (atoms) and edges (atom pairs) of the molecule. **X**^new^ are the updated molecular node coordinates. Based on the outputs of the encoder, three decoders were applied to predict the probabilities of atom types **A**^raw^, the coordinates **X**^raw^, and the probabilities of bond types **B**^raw^ of the denoised molecules, in company with the confidence scores for individual predicted variables, as follows,

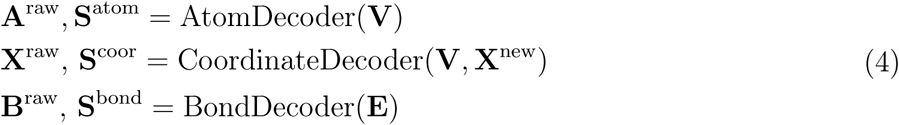

where **S**^atom^ ∈ [0, 1]*^N^ ,* **S**^coor^ ∈ [0, 1]*^N^* , and **S**^bond^ ∈ [0, 1]*^N^*^×^*^N^* are the confidence scores for the predicted atom types, atom coordinates and bond types, respectively, and the complete confidence scores are denoted as S = {**S**^atom^, **S**^coor^, **S**^bond^}.

For the generation process, an additional module, named M-Projector, is designed to derive the atom and bond types from the predicted probabilities and adapt the predicted variables to align with the prompt, as follows,

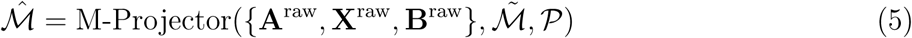

Detail implementations of the architectures can be found in Supplementary Notes 1.

### Task prompt

The task prompt is defined as a group of binary indicators to represent whether the individual variables of the input molecule should be preserved as provided or generated by the model. Instead of setting a series of hard rules, we incorporated prompts as part of the model inputs to let the model automatically follow the requirements specified by the prompt. The advantages of designing such prompts include: 1) Diverse molecular generative tasks can be unified under the single computational framework; 2) Generation demands can be more accurately represented than using natural language; 3) Our prompt can also act as a task embedding, precisely measuring the similarity between tasks, allowing PocketXMol to extend to new molecule tasks in a more effective way. Moreover, PocketXMol is highly flexible and can also support many atypical generative tasks, such as growing an existing molecular fragment, in the meanwhile docking the molecule with the protein. Users can customize their own prompts by leveraging the tool we developed. The complete list of prompts for each task can be found in the Supplementary Notes 2.

### Molecular noise

An important factor enabling PocketXMol to support an unprecedented number of generative tasks is the incorporation of various types of noise. First, Gaussian noise was introduced to perturb individual atom coordinates in 3D space or as random coordinate translation. Second, the discrete variables such as atom and bond types are perturbed using noise following the categorical distribution. Third, random rotations were applied to a set of atom coordinates by multiplying the coordinates with rotation matrices sampled from an isotropic Gaussian distribution in SO(3)^109^. Last, we also introduced noise following the circular normal distribution to perturb the torsion angles of rotatable bonds as an indirect perturbation to atom coordinates. A generative task typically utilizes one or multiple types of these noise distributions. For instance, for structure-based drug design, the noise associated with atom coordinates follows the Gaussian distribution, whereas the distribution of atom and bond types follows the categorical distribution. The definition of the noise distributions and specification for tasks can be found in Supplementary Notes 3.

### Generation process

To perform a generative task, we need to define the corresponding task prompt P and choose the task-specific molecular noise *ξ*. We repeatedly perturbed the present molecule with scaled noise using the perturbation function Φ (Eq. 1) and removed the noise using the denoiser function *F*_Θ_ (Eq. 2). Formally, the generation process is defined as,

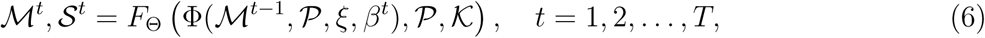

where M*^t^* and S*^t^* are the denoised molecule and the confidence scores at the *t*th step, respectively. *T* is the total number of steps (*T* =100 in our implementation). *β^t^* ∈ [0, 1] is the noise scale at the *t*th step and gradually decays to zero as the step *t* increases. The initial molecule M^0^ is either provided by the user or randomly sampled from the task noise distribution *ξ*. The molecule M*^T^* represents the final generated molecule.

This generation process resembles the diffusion-based generative models, where the generation is also a series of denoising steps^32, 33^, but they have several distinctions. First, instead of defining a Markov chain, in PocketXMol, each perturbing-denoising step can operate independently, which provides enhanced flexibility during the generation process. As explained in the next subsection, our framework can integrate prior knowledge more easily compared to diffusion models by leveraging this mechanism. Second, PocketXMol does not require explicit information regarding the scale or types of the noise as input. In order to incorporate distinct tasks, PocketXMol is designed to automatically identify the noise characteristics from the input molecules and remove them based on the task specification. Additionally, the denoiser also produced confidence scores ranging from 0 to 1 for each generated variable, which can be utilized for molecule selection and adjustment of the noise scale. Last, one diffusion model can only process a single type of noise, while PocketXMol is capable of processing a variety of molecular noises defined for different molecular components, enabling it to concurrently learn and adapt to different molecular generative tasks. Details related to the implementation of the generation process can be found in the Supplementary Notes 4.

### Integration of prior knowledge

An advantage of PocketXMol is the flexibility to integrate prior knowledge during the generation process by either defining the task prompt or adjusting the perturbation process. Certain prior knowledge can be directly expressed through the prompt. For instance, we might already know the exact coordinates of a set of atoms for molecular docking^64^. In this case, we can set the prompt of the associated atom coordinates as fixed and drive the model to only predict the coordinates of other atoms. Other more general prior knowledge can be injected by modifying molecules towards the prior knowledge before adding noise at each perturbation step. For example, when we know a set of atoms are located within a certain area, we can move the atom coordinates if they are located outside the area at the perturbation steps during the generation process. Details of how we modified the molecules for different prior knowledge can be found in Supplementary Notes 4.

### Model training

The neural networks in the denoiser were trained by restoring the molecules in the training dataset from the noisy ones conditional on the task-specific noise and prompt. For each molecule in the training dataset D, we randomly sampled the task-specific prompt and molecular noise from the possible task sets P, and noise scale *β* from the interval (0, 1), and optimized the following loss function L,

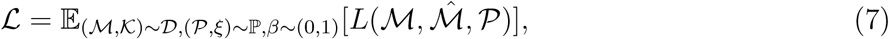

where M^^^ is the output of the denoiser:

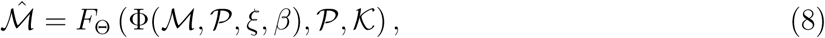

and *L*(M, M^^^ , P) is defined as a weighted summation of the losses of individual variables:

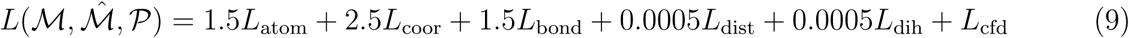

where *L*_atom_ and *L*_bond_ are cross-entropy loss functions for the atom types and bond types, respectively, *L*_coor_ is the square error loss function for atom coordinates. *L*_dist_ and *L*_dih_ are pairwise distance loss function and dihedral angle loss function, respectively, which are only valid when random rigid transformation and rotation of torsion angles are applied. *L*_cfd_ is the predicted confidence. The weights are determined based on the importance of these variables. We trained the model by optimizing the loss function L using the AdamW optimizer^110^, with a batch size (per GPU) of 40 and a base learning rate of 0.001, on eight 80G A100 GPUs for 180,000 steps (around 54 hours). More details related to the definition of each loss function and the training process can be found in Supplementary Notes 5.

### Molecular generative tasks

Here we briefly introduce the task prompt P and molecular noise *ξ* for individual molecular generative tasks.

#### Molecular docking

The molecular docking task involves docking small molecules or peptides to the protein pockets. The prompts and noise distributions for these two tasks are identical because our atom-level model does not differentiate between small molecules and proteins. Molecular docking requires generating 3D coordinates for the atoms with all the atom and bond types provided. The prompt indicators associated with atom coordinates are set as zero (to be generated), and those associated with atom and bond types are set as one (fixed). There exist two kinds of molecular noise for docking and either of them can be used: 1) Gaussian noise for atom coordinates and 2) flexible noise, i.e., a combination of the random translation sampled from Gaussian distribution, the molecule rotation sampled from the isotropic Gaussian distribution in SO(3), and rotation of torsion angles sampled from the circular normal distribution. Details can be found in Supplementary Notes 2.2.1 and 3.4.1.

#### Molecular conformation prediction

This task is to predict the 3D coordinates of each atom given the 2D molecular graph, without the constraints of protein pocket. Therefore, the task prompt and molecular noise for molecular conformation prediction tasks are identical to those of molecular docking. Details can be found in Supplementary Notes 2.2.2 and 3.4.2.

#### Structure-based drug design (SBDD)

The object of SBDD is to generate a 3D molecule that can bind to the 3D protein pocket. It involves generating the types and 3D coordinates of all the atoms and bonds for a small molecule. The indicators of the task prompt were all set as unfixed. The Gaussian noise was introduced to atom coordinates and noise following categorical distributions was introduced to both atom and bond types. Details can be found in Supplementary Notes 2.2.3 and 3.4.3.

#### 3D molecular generation

This task is defined as generating novel small molecules without the consideration of protein pockets. Therefore, the task prompt and molecular noise are identical to those of SBDD. Detail algorithms can be found in Supplementary Notes 2.2.4 and 3.4.4.

#### Molecule optimization

Molecular optimization is a special case of SBDD, where the model takes a protein-molecule 3D complex as input, aiming to generate another 3D molecule with improved properties. The task prompt and molecular noise are identical to SBDD, except that the initial molecule is provided instead of a random one. The noise scale is initialized as 0.3 instead of 1 as SBDD. Details can be found in Supplementary Notes 2.2.5 and 3.4.5.

#### Fragment linking

We utilized the fragment linking benchmark, Binding MOAD dataset^38^ in which each ligand is divided into two fragments based on certain criteria^69^. The docking positions between fragments and pockets are kept the same as the original complex structure. The prompt indicators associated with the atoms and bonds of the fragments were set as fixed and the rest were set as unfixed. For the unfixed part of the molecule, we added Gaussian noise to the atom coordinates and noise following the categorical distribution to atom and bond types, respectively. A common setting in fragment linking is to specify the atoms of fragments that connect to the linker. In this case, we set the prompt indicators as fixed for all the chemical bonds between the non-connecting atoms of the fragments and all atoms of the linkers. In this way, the model can only choose to form chemical bonds via the connecting atoms of the fragments with the linkers to form a complete molecule. Details can be found in Supplementary Notes 2.2.6 and 3.4.6.

#### PROTAC design

The PROTAC drug consists of two linked domains: one that binds to the target protein and another that binds to an E3 ligase. Once the PROTAC molecule brings the target protein into close proximity with the E3 ligase, the target protein is tagged with ubiquitin, a small protein that signals for degradation. The task prompt and the molecular noise are almost the same as those of fragment linking except the atom coordinates of the fragments are unfixed and accept Gaussian noise. More details can be found in Supplementary Notes 2.2.7 and 3.4.7.

#### Fragment growing

The fragment growing task is similar to fragment linking but differs in that medicinal chemists need to pre-define a fragment that binds to the protein pocket and then design a complete molecule starting from this fragment. The initial fragment is typically much smaller than typical drug-like molecules. To evaluate our model, we first selected a molecule from the test set and divided it into multiple pieces using BRICS decomposition^73^. We selected the largest piece and extended it to connect other fragments until five atoms were achieved. The task prompt and noise for fragment growing are similar to fragment linking. Details can be found in Supplementary Notes 2.2.8 and 3.4.8.

#### De novo peptide design

The challenge of peptide design is how to prompt the model to generate a chain of amino acids within the pocket rather than any arbitrary 3D small molecules since PocketXMol does not distinguish between amino acids and small molecules. Our solution is to provide the peptide backbone atoms as input because the backbone atoms of all amino acids are identical, allowing their atom types to be pre-defined, and only the side chain atoms need to be generated. The prompts and the noise are similar to the fragment growing task, regarding the backbone atoms as the fragment part and the side-chain atoms as the growing part. In addition, the backbone atoms capable of connecting to side chains must be the alpha-carbon atoms or the nitrogen atoms (for proline), and hence the prompt indicators for bonds related to the non-connecting atoms (i.e., the oxygen atoms and carbon atoms) of backbones are fixed. Details can be found in Supplementary Notes 2.2.10 and 3.4.9.

#### Peptide inverse folding

Protein inverse folding involves predicting the amino acid sequence that will fold into a specific 3D protein structure Traditional methods typically define the inverse folding problem as a classification task that predicts one of the twenty types of amino acids for each position based on the 3D complex structure. Under our framework, we only need to generate the atom types and coordinates for the side chains, conditioned on the fixed backbone atoms and the protein pocket. Therefore, the only difference between peptide inverse folding and peptide design is that the atom coordinates of the peptide’s backbone are fixed. Details can be found in Supplementary Notes 2.2.11 and 3.4.10.

### Western blot of designed small molecules

Extract total protein from cells and quantify protein concentration using the BSA method. Ensure that the loading amount of each sample is consistent based on the protein concentration. Perform SDS-PAGE electrophoresis, then transfer the electrophoresed protein to a PVDF membrane. Block with non-fat milk, incubate with antibodies specific to the protein of interest based on molecular weight, and after incubating with secondary antibodies, use a chemiluminescent reagent kit for detection. Analyze the expression of specific proteins in each sample. The antibodies of caspase-9 (#9508), cleaved caspase-3 (Asp175) (#9664), *β*-Actin Rabbit mAb(#4970), caspase-3 (#9662) and PARP (#9542) were purchased from Cell Signaling Technology.

### Caspase-3 activity analysis

Mix 50 ng of caspase-3 with cell lysis buffer to a total volume of 50 µL, then add to a 96-well plate containing 45 µL of reaction buffer. After adding Ac-DEVD-pNA (Elabscience, E-CK-A383), mix well and incubate at 37°C for 2 hours. Measure the OD405 and calculate the caspase 3 enzyme activity in each sample based on a pre-made standard curve.

### EC_50_ analysis

MC38 cells were seeded into a 6-well plate, and incubated overnight. Cells were treated with ABT-737(10µM) and/or a two-fold serial dilution of caspase inhibitor. After 12 hours of incubation, cells were collected and lysed. The cell lysate was used to detect the enzyme activity of caspase-3. The EC_50_ were calculated with GraphPad software, determining the concentration of caspase inhibitor at which 50% enzyme activity inhibition is observed.

### SPR analysis of caspase-9 binding

Surface plasmon resonance (SPR) analysis was performed to evaluate the binding of small molecules to wild-type and mutant caspase-9 proteins. Compounds including D12 and 84663 were tested at concentrations ranging from 10 to 10,000 nM. Dimeric caspase-9 or its mutants were immobilized on a CM5 sensor chip, and binding measurements were conducted at 25°C using a 1×PBST buffer containing 5% DMSO. Sensorgrams were recorded at a constant flow rate of 30 µL/min to assess dose-dependent binding profiles.

### SPRi analysis of PD-L1 and peptides

The surface plasmon resonance imaging (SPRi) analysis was conducted on a Plexera PlexArray HT system (Plexera LLC, Bothell, WA) using bare gold SPRi chips (Nanocapture gold chips). A total of 384 peptides (382 designed peptides with two positive and negative controls) were synthesized via solid-phase synthesis, purified, dissolved in dimethyl sulfoxide, and then dispensed into a 384-well plate. The peptides were transferred onto the surface of the bare gold SPRi chip using an Echo550 Liquid Handler and incubated overnight at 4°C in a humidified chamber. After being washed three times with phosphate buffer saline (PBS), the SPRi chip was blocked with 5 % (w/v) non-fat milk for 8 hours, followed by another three washes with PBS, before the chip was inserted into the instrument for detection. The flowing phase contained PD-L1 protein in five concentrations: 23.3, 46.6, 93.3, 186.6, and 373.1 nM, in a PBST buffer. The regeneration solution was 0.5 % (v/v) H_3_PO_4_. Real-time binding signals were recorded, and dissociation constant (*K*_D_) values were analyzed using the PlexArray HT system.

### In vitro fluorescent imaging

We selected the lung cancer cell line H1975 as PD-L1 positive cells, and human embryonic kidney cells, HEK293T, as PD-L1 negative cells. Cells were cultured in confocal dishes using Dulbecco’s modified Eagle’s medium (DMEM) or RPMI-1640 (HyClone) supplemented with 10% fetal bovine serum (Corning), 100 U/mL penicillin, and 100 µg/mL streptomycin, in a 37°C incubator with 5% CO2 for 24 hours. Subsequently, FITC-labeled peptide (20 µg/mL) and Hoechst 33342 solution (10 µg/mL) were added to each sample well. The samples were then incubated for 20 minutes at 37°C, protected from light. Additionally, the cells were treated with Alexa Fluor 594 Anti-PD-L1 antibody (ab213360, 1:1000 dilution) for colocalization analysis. Finally, the cells were gently washed three times with prechilled PBS. Fluorescence imaging of the samples was performed using a laser confocal microscope. Hoechst 33342 represents the nucleus, blue; FITC represents the peptide, green; Alexa Fluor 594 represents the PD-L1 protein, red.

### In vivo fluorescence imaging of tumor

The xenografted tumors were established by subcutaneously injecting 1 × 10^6^ H1975 cells into the right hind leg of 8-week-old female BALB/c nude mice. Once the tumor size increased to 80 mm^3^, the mice were used for in vivo imaging experiments. The peptides P65, P73, and P282, labeled with indocyanine green (ICG), and free ICG dye as a control, were injected into the mice via the tail vein in a volume of 150 µL (n=3). The mice were anesthetized using a gas anesthesia method with 2 L/min oxygen mixed with 2.5 % isoflurane. Imaging was performed at 0.5 h, 2 h, 4 h, 8 h, and 12 h after injection using a small animal 3D in vivo imaging system (IVIS Spectrum). After 12 hours, the mice were euthanized, and the heart, liver, spleen, lung, kidney, and tumor tissues were excised for ex vivo fluorescence imaging and quantitative analysis. All animal protocols were approved by the Institutional Animal Care and Use Committee of the National Center for Nanoscience and Technology.

### Peptide chemical synthesis library

In order to compare with AI-designed peptides, we constructed a random peptide library with a capacity of 10^7^ to search peptides targeting PD-L1. Tenta Gel resin was used as the solid-phase support, and the Fmoc solid-phase synthesis strategy was employed to progressively synthesize X_1_X_2_X_3_X_4_X_5_X_6_X_7_X_8_X_9_X_10_M. The synthesis process employed a “mix and split” approach, where each synthetic product from each step was evenly divided, and different amino acids were randomly added for the next reaction. The products were thoroughly mixed and divided again, and the next round of amino acids was added for reaction. This cycle continued until the final round of amino acid reaction was completed. The entire reaction process took place in a peptide synthesizer. In each coupling reaction, the following steps were carried out: Deprotection (20 % piperidine), Washing (N, N-dimethylformamide (DMF) solution), Coupling (Fmoc-protected amino acid dissolved in a 4 % N-methyl morpholine DMF solution with an equivalent amount of 2-(1H-benzotriazol-1-yl)-1,1,3,3-tetramethyluronium hexafluorophosphate (HBTU)). After the complete reaction, deprotection and washing were performed, followed by thorough mixing of the resin. After completion of the side chain deprotection reaction by 95 % trifluoroacetate (TFA), the resin was washed several times with methanol and vacuum dried. The peptide library was then incubated with the PD-L1 protein, and positive peptide beads were screened using the microfluidic chip method established in our previous work^111^.

### Data availability

The raw data are available in public databases. To train the model, we assembled a large data set and precisely removed redundancy to all test sets. We used three types of data from multiple databases or datasets: 1) protein-small-molecule complexes, including PDBBind^37^, Binding MOAD^38^, and the CrossDocked2020 dataset^39^; 2) protein-peptide complexes including PepBDB^40^ and a synthetic dataset of protein-binding loop structures from AlphaFoldDB^3^; 3) small molecule or peptide structures, including the GEOM dataset^43^ (GEOM-Drug and GEOM-QM9), CREMP dataset^44^, and a large molecular data dataset proposed by Uni-Mol^45^ which contained small molecules from ZINC^42^, ChEMBL^41^ and other purchasable molecular databases.

We cleaned and filtered them using some criteria. To avoid data leakage, we then removed pocket-based data within MMSeqs2^112^ clusters of 30% sequence identity of any data in any test sets and pocket-free data with the same molecular graphs to any data in any test sets. We randomly divided the data into training and validation sets in certain ratios. For details related to data processing, please refer to Supplementary Notes 6 and 6.2. We also provided the raw data and the processing steps in our code base at https://github.com/pengxingang/PocketXMol.

### Code availability

The source code, along with detailed instructions for training, sampling, and usage, is available at https://github.com/pengxingang/PocketXMol. Colab notebooks for easy experimentation can also be found in the `notebooks` directory: https://github.com/pengxingang/PocketXMol/tree/master/notebooks.

