## Supplementary Materials for "Atom-level generative foundation model for molecular interaction with pockets"

#### Contents

Supplementary Notes 9

1 Model architecture 10
- 1.1 Encoder architectures 10
  - 1.1.1 PocketEncoder architecture 10
  - 1.1.2 MoleculeEncoder architecture 11
- 1.2 Decoder architectures 19
- 1.3 M-Projector 20

**2 Task prompt** **24**

**References** **87**

**Supplementary Tables** **91**

**Supplementary Figures** **101**

#### List of Algorithms

1 **Encoder:** learn the representations of atoms and atom pairs for the molecule . . . 10
2 **PocketEncoder:** learn atom representations for a protein pocket . . . 11
3 **MoleculeEncoder:** update the node and edge feature and node coordinate of the molecule . . . 12

4 **DistanceBlock:** calculate distance features of edges . . . 13
5 **NodeBlock**: update molecular node features . . . 13
6 **EdgeBlock**: update molecular edge features . . . 15
7 **PosBlock:** update molecular node coordinates in $\mathcal{G}_{\text{mol}}$ . . . 16
8 **PosBlockFromPocket:** update molecular node coordinates in $\mathcal{G}_{\text{mol-poc}}$ . . . 18
9 **AtomDecoder:** predict atom types . . . 19
10 **CoordinateDecoder:** predict atom coordinates . . . 20
11 **BondDecoder:** predict bond types . . . 20
12 **AdaptCoordinateFixedDist:** adapt the 3D coordinates with fixed distances . . . 21
13 **CalcTorsionAngleChanges:** calculate torsion angle changes between two sets of molecular coordinates . . . 22
14 **CalcDihedralAngle:** calculate a dihedral angle . . . 22
15 **RotBonds:** rotate the bonds . . . 23
16 **AlignStructure:** get the rigid transformation to align two sets of coordinates . . . 24
17 **GetDistPromptFlexNoise:** get the distance prompt indicators using flexible noise 26
18 **GetDistPromptSpecifyRot:** get the distance prompt indicators with specified rotatable bonds . . . 30
19 **PertCoordinateFlex:** $\Phi^{\text{flex}}$, perturb coordinates with flexible noise . . . 37
20 **OrthogonalRotBonds:** rotate rotatable bonds orthogonally to rigid transformation. 37
21 **GeneralPert:** general perturbation step . . . 38
22 **PertDockSmallMolGauss:** perturbation for molecular docking using Gaussian noise. . . . 39
23 **PertDockSmallMolFlex:** perturbation for molecular docking using flexible noise. 39
24 **PertSBDD:** perturbation for structure-based drug design . . . 40
25 **PertFragLinking:** perturbation for fragment linking . . . 42
26 **PertPepDesign:** perturbation for *de novo* peptide design . . . 43
27 **PertPepInvFolding:** perturbation for peptide inverse folding . . . 44
28 **GenerationProcess:** general generation process of PocketXMol . . . 45
29 **SimpleGenDockSmallMol:** simple generation process for protein-small molecule docking . . . 46
30 **PrepareInitCoordinates:** prepare the initial coordinates for docking . . . 47
31 **GenDockSmallMol:** generation process for protein-small-molecule docking . . . 47
32 **GenSelfRankingScore:** get self ranking scores . . . 48
33 **GenSelDockSmallMol:** generation and selection for protein-small molecule docking 48
34 **GenTunedRankingScore:** get tuned ranking scores . . . 49
35 **GenFlexRankingScore:** get confidence scores using flexible noise . . . 49
36 **ModifyCenter:** modify the molecule to align the reference center position . . . 50
37 **ModifyBondLength:** modify the molecule to align the bond lengths . . . 51
38 **ModifyAtomCoordinate:** modify the molecule to align the reference atom coordinate . . . 51
39 **GenConf:** generation process for molecular conformation generation . . . 52
40 **SimpleGenSBDD:** simple generation process for structure-based drug design . . . 53
41 **PrepareInitMolecule:** prepare initial molecule for molecule design . . . 53
42 **GetMolSizeFromPocket:** sample number of molecular atoms based on pocket size 54
43 **GetMolSizeFromRef:** sample number of molecular atoms based on reference size 54
44 **GenSBDD:** generation process with refining using confidence scores for SBDD . . 55

45 **PrepareRegen:** prepare for re-generation round of SBDD . . . . . . . . . . . . . . . 56
46 **PrepareRegenInitMolecule:** prepare initial molecule for re-generation . . . . . . 56
47 **DefineRegenNoise:** define molecular noise and noise scale for re-generation . . . . 57
48 **GenSBDDAR:** generation process using confidence scores for SBDD in an auto-regressive-like manner . . . . . . . . . . . . . . . . . . . . . . . . . . . . . . . . . . 57
49 **PrepareAR:** prepare for auto-regressive-like generation . . . . . . . . . . . . . . . . 58
50 **ModifyShape:** modify the molecule to align with shape . . . . . . . . . . . . . . . . . 59
51 **GenFragLinking:** generation process for fragment linking . . . . . . . . . . . . . . . 60
52 **PrepareInitMoleculeFrag:** prepare initial molecule with fragment . . . . . . . . . . 61
53 **GenFragGrowing:** generation process for fragment growing . . . . . . . . . . . . . . 62
54 **GenOptLogP:** generation process for optimizing the LogP of molecule . . . . . . . 62
55 **GenSimilarMol:** generation process for generating a similar molecule . . . . . . . 63
56 **GenPepDesign:** generation process for *de novo* peptide design . . . . . . . . . . . 63
57 **GetBackboneFrag:** get the backbone fragment for linear peptide generation . . . 64
58 **GetNumAtomsPep:** sample the number of atoms for peptide generation . . . . . 64
59 **GetBackboneFragCyclic:** get the backbone fragment for cyclic peptide generation 65
60 **GenPepInvFolding:** generation process for peptide inverse folding . . . . . . . . . 65
61 **ModelTraining:** train the denoiser neural networks . . . . . . . . . . . . . . . . . . 66
62 **CalcLoss:** calculate the loss for a denoised molecule . . . . . . . . . . . . . . . . . 69
63 **CalcAtomLoss:** calculate the loss for predicted atom types . . . . . . . . . . . . . . 70
64 **CalcCoordinateLoss:** calculate the loss for predicted atom coordinates . . . . . . 70
65 **CalcBondLoss:** calculate the loss for predicted bond types . . . . . . . . . . . . . . 70
66 **CalcDistanceLoss:** calculate the loss for predicted atom distances . . . . . . . . . 71
67 **CalcDihedralLoss:** calculate the loss for predicted dihedral angles . . . . . . . . . 71
68 **GetFlexDihedralAngleIndices**: get the indices of the active dihedral angles in the flexible noise . . . . . . . . . . . . . . . . . . . . . . . . . . . . . . . . . . . . . . . 72
69 **CalcConfidenceLoss:** calculate the loss for predicted confidence scores . . . . . . 73

S24 The Rosetta binding energies and the distributions of secondary structure for the cyclic peptides in the datasets or generated by different models. . 117
S25 Properties and Rosetta energy of NAAs. . . . . . . . . . . . . . . . . . . . . . . 118
S26 Side chains of standard amino acids and some frequent NAAs generated by PocketXMol. . . . . . . . . . . . . . . . . . . . . . . . . . . . . . . . . . . 119
S27 Some physical and chemical properties of the standard amino acids and NAAs calculated by AAindexNC. . . . . . . . . . . . . . . . . . . . . . . . 120
S28 Confocal image of FITC labeled peptides binding toward negative 293T cells. . . . . . . . . . . . . . . . . . . . . . . . . . . . . . . . . . . . . . . 121
S29 Ex vivo images of tumors and main organs (heart, liver, spleen, lung, kidney) that were taken 12 hours after post-injection. . . . . . . . . . . . . 121
S30 The SPRi binding curves and the confocal image of FITC labeled peptides binding with cells for the two positive peptides obtained through library screening. . . . . . . . . . . . . . . . . . . . . . . . . . . . . . . . . . . . 122
S31 The in vivo experiments for the two positive peptides (P1 and P5) obtained through library screening. . . . . . . . . . . . . . . . . . . . . . . . . . . 123
S32 Binding structures generated by PocketXMol and measured $K_D$ changes for binding-residue mutants of three designed peptides (P65, P73, and P282). . . . . . . . . . . . . . . . . . . . . . . . . . . . . . . . . . . . . . 124
S33 Extended results for small molecule docking. . . . . . . . . . . . . . . . . . . 125
S34 Relationships between docking performance and different molecular properties. . . . . . . . . . . . . . . . . . . . . . . . . . . . . . . . . . . . . . 126
S35 The number of molecules that satisfy the 3D structural screening criteria proposed by Posebusters. . . . . . . . . . . . . . . . . . . . . . . . . . . . 127
S36 Small molecular docking with different PocketXMol and input pocket structures. . . . . . . . . . . . . . . . . . . . . . . . . . . . . . . . . . . . . 128
S37 Relationship between the performance of small molecular docking and the pocket RMSDs for PocketXMol (self ranking) with different input structures. . . . . . . . . . . . . . . . . . . . . . . . . . . . . . . . . . . . 129
S38 Relationship between the performance of small molecular docking and the pocket RMSDs for PocketXMol (oracle ranking) with different input structures. . . . . . . . . . . . . . . . . . . . . . . . . . . . . . . . . . . . 130
S39 An example showing PocketXMol could generate high-quality conformations for a 2D molecular graph (ChEMBL ID: CHEMBL1307825). . . . . 131
S40 Extended results for peptide docking on the PepBDB set. . . . . . . . . . . 132
S41 Peptide docking performance (on the PepBDB set) of small-molecule docking tools. . . . . . . . . . . . . . . . . . . . . . . . . . . . . . . . . . . 133
S42 Docking performance on the BioLiP peptide docking test set for AlphFold-Multimer, AlphaFold 3, and PocketXMol. . . . . . . . . . . . . . . . . . . 134
S43 Peptide docking on PepBDB set for different PocketXMol and different input pocket structures. . . . . . . . . . . . . . . . . . . . . . . . . . . . . 135
S44 Peptide docking on the Q-BioLiP set for different PocketXMol and different input pocket structures. . . . . . . . . . . . . . . . . . . . . . . . . . 135
S45 Relationship between DockQ of peptide docking (on the PepBDB set) and the peptide length in cases of different PocketXMol and input pocket structures. . . . . . . . . . . . . . . . . . . . . . . . . . . . . . . . . . . . 136

S46 **Relationship between DockQ of peptide docking (on the PepBDB set) and the ratio of secondary structures helix and sheet in cases of different PocketXMol and input pocket structures.** . . . 137
S47 **Relationship between DockQ of peptide docking (on the Q-BioLiP set) and the peptide properties in cases of different PocketXMol and input pocket structures.** . . . 138
S48 **Performance of substrate activity prediction by model confidence scores.** 139
S49 **Relationship between the enzyme-substrate activity prediction metric and ratio of active samples of the enzyme proteins.** . . . 140
S50 **Performance on enzyme activity prediction using logistic regression models and PocketXMol representation vectors.** . . . 141
S51 **Comparison of predicted enzyme–substrate structure characteristics between PocketXMol and AlphaFold 3.** . . . 142

### Supplementary Notes

**Table S1.** Main notations.

| | |
|---|---|
| **Molecules and protein pockets** | |
| $N$ | Number of heavy atoms of a molecule (or sub-molecule) |
| $\mathcal{M}$ | A molecule, $\mathcal{M} = \{\mathbf{A}, \mathbf{X}, \mathbf{B}\}$ |
| $K_{\text{atom}}$ | Number of atom types |
| $K_{\text{bond}}$ | Number of bond types |
| $\mathbf{a}_i \in \{0,1\}^{K_{\text{atom}}}$ | One-hot encoding for atom types of atom $i$ |
| $\mathbf{A} = [\mathbf{a}_i] \in \{0,1\}^{N \times K_{\text{atom}}}$ | Atom types for a certain molecule |
| $\mathbf{x}_i \in \mathbb{R}^3$ | 3D coordinate for atom $i$ |
| $\mathbf{X} = [\mathbf{x}_i] \in \mathbb{R}^{N \times 3}$ | Atom coordinates for a certain molecule |
| $\mathbf{b}_{ij} \in \{0,1\}^{K_{\text{bond}}}$ | One-hot encoding of bond type between atom $i$ and $j$ |
| $\mathbf{B} = [\mathbf{b}_{ij}] \in \{0,1\}^{N \times N \times K_{\text{bond}}}$ | Bond types for a certain molecule |
| $\mathcal{K}$ | Protein pocket |
| $\mathbb{A}$ | A set of atom indices |
| $\mathbb{B}$ | A set of bond indices (pairs of atom indices) |
| $\mathbb{M}$ | A union set of atom indices and bond indices: $\mathbb{A} \cup \mathbb{B}$ |
| $\mathcal{S} = [\mathbf{S}^{\text{atom}}, \mathbf{S}^{\text{coor}}, \mathbf{S}^{\text{bond}}]$ | predicted confidence scores |
| **Task prompt and noise** | |
| $\mathcal{P}$ | Task prompt, $\mathcal{P} = \{\mathbf{P}^{\text{atom}}, \mathbf{P}^{\text{coor}}, \mathbf{P}^{\text{bond}}, \mathbf{P}^{\text{dist}}, \mathbf{P}^{\text{pep}}\}$ |
| $\mathbf{P}^{\text{atom}} \in \{0,1\}^N$ | Indicators for whether the atoms types are fixed |
| $\mathbf{P}^{\text{coor}} \in \{0,1\}^N$ | Indicators for whether the atoms coordinates are fixed |
| $\mathbf{P}^{\text{pep}} \in \{0,1\}^N$ | Indicators for whether the atoms belong to peptides |
| $\mathbf{P}^{\text{bond}} \in \{0,1\}^{N \times N}$ | Indicators for whether the type of bonds are fixed |
| $\mathbf{P}^{\text{dist}} \in \{0,1\}^{N \times N}$ | Indicators for whether the atom distances are fixed |
| $T$ | Total steps of a generation process |
| $\beta$ | Noise scale (a scalar or a mapping function) |
| $\boldsymbol{\beta}$ | A sequence of noise scales $[\beta^1, \beta^2, \ldots, \beta^T]$ |
| $\xi$ | Molecular noise |
| $\sigma$ | Standard deviation of a Gaussian distribution |
| $\epsilon$ | Standard deviation of a Gaussian distribution in SO(3) |
| $\kappa$ | Concentration parameter of a circular normal distribution |
| $\mathbf{R}; \mathbf{t}$ | Rotation matrix; translation vector |
| $\Delta\omega$ | Change of a torsion angle |
| **Function** | |
| $F_\Theta$ | The denoiser with learnable parameters as $\Theta$ |
| $\Phi$ | The general molecular perturbation process |
| $\Psi$ | Molecular modification following prior knowledge |
| **Misc.** | |
| $[\![n]\!]$ | Abbreviation of $[1, 2, \ldots, n]$ |
| $\tilde{\cdot}$ | Noisy variable |
| $\hat{\cdot}$ | Variable related to model predictions |
| $\lVert \cdot \rVert$ | L2-norm of a vector or size of a set |

### 1 Model architecture

Generally, the denoiser $F_\Theta$ is composed of three major modules, the encoder to learn from the inputs, the decoders to predict the denoised molecular variables, and the projector to construct the final molecule from the variables. We utilized geometric graph neural networks in the encoders and decoders, where graphs were constructed by treating the atoms as nodes and certain atom pairs as edges. The details are described as follows.

#### 1.1 Encoder architectures

The encoder module contains two sub-modules: a pocket encoder and a molecule encoder. The protein pockets always serve as conditions in our generative model because we do not generate or modify them. Therefore, we first constructed a module named PocketEncoder to encode and process protein pockets. For molecules to be generated, we then utilized a module, named MoleculeEncoder to capture the interactions within the molecules as well as between the molecules and the atoms of the protein pockets.

**Algorithm 1 Encoder:** learn the representations of atoms and atom pairs for the molecule

**Input:** a noisy molecule $\tilde{\mathcal{M}} = \{\mathbf{A}, \mathbf{X}, \mathbf{B}\}$, pocket $\mathcal{K} = \{\mathbf{A}^{\text{poc}}, \mathbf{X}^{\text{poc}}\}$, task prompt $\mathcal{P}$.

**Output:** molecular node (atom) features $\mathbf{V}$, updated molecular coordinates $\mathbf{X}^{\text{new}}$, molecular edge (atom pair) features $\mathbf{E}$.

$\mathbf{V}^{\text{poc}} = \text{PocketEncoder}(\mathbf{A}^{\text{poc}}, \mathbf{X}^{\text{poc}})$

$\mathbf{V}, \mathbf{X}^{\text{new}}, \mathbf{E} = \text{MoleculeEncoder}(\mathcal{M}, \{\mathbf{V}^{\text{poc}}, \mathbf{X}^{\text{poc}}\}, \mathcal{P})$

##### *1.1.1 PocketEncoder architecture*

The PocketEncoder builds a $k$-nearest neighbor graph $\mathcal{G}_{\text{poc}}$ with $k = 32$ for all the pocket atoms and uses graph neural networks to learn the pocket node (atom) features $\mathbf{V}^{\text{poc}}$, as defined in Algorithm 2 (Figure S1). Here, Linear is the linear function, and MLP is the multilayer perceptron including a sequence of a linear layer, layer normalization, ReLU activation, and another linear layer. RBF in Line 5 is the radial basis function using 32 Gaussian kernels with kernel means ranging from 0 to 10. The output dimension of the Linear function in Line 1 is 32, and all other hidden dimensions in the algorithm are 128.

**Algorithm 2 PocketEncoder:** learn atom representations for a protein pocket

**Input:** pocket atom features $\mathbf{A} = [\mathbf{a}_i]$, pocket atom coordinates $\mathbf{X} = [\mathbf{x}_i]$.

**Output:** pocket node (atom) features $\mathbf{V} \in \mathbb{R}^{N_{\text{poc}} \times 128}$.

pocket atom graph $\mathcal{G}_{\text{poc}} \leftarrow k$-nearest neighbor graph ($k = 32$) of $\{\mathbf{A}, \mathbf{X}\}$

$\mathbf{v}_i = \text{Linear}(\mathbf{a}_i)$ ▷ $\forall i \in$ nodes of $\mathcal{G}_{\text{poc}}$. The same below.

$d_{ij} = \|\mathbf{x}_i - \mathbf{x}_j\|$ ▷ $\forall ij \in$ edges of $\mathcal{G}_{\text{poc}}$. The same below.

**for all** block $\in [1, 2, 3, 4]$ **do**

  $\tilde{\mathbf{v}}_i \leftarrow \text{MLP}(\mathbf{v}_i)$

  $\mathbf{e}_{ij} \leftarrow \text{MLP}\left(\text{RBF}(d_{ij})\right)$

  $\mathbf{m}_{ij} \leftarrow \text{Linear}(\mathbf{e}_{ij} + \tilde{\mathbf{v}}_i + \tilde{\mathbf{v}}_j)$

  $\tilde{\mathbf{v}}_i \leftarrow \text{Linear}(\tilde{\mathbf{v}}_i) + \sum_j \mathbf{m}_{ij}$

  $\mathbf{v}_i \leftarrow \mathbf{v}_i + \text{LayerNorm}\left(\text{MLP}(\tilde{\mathbf{v}}_i)\right)$

**end for**

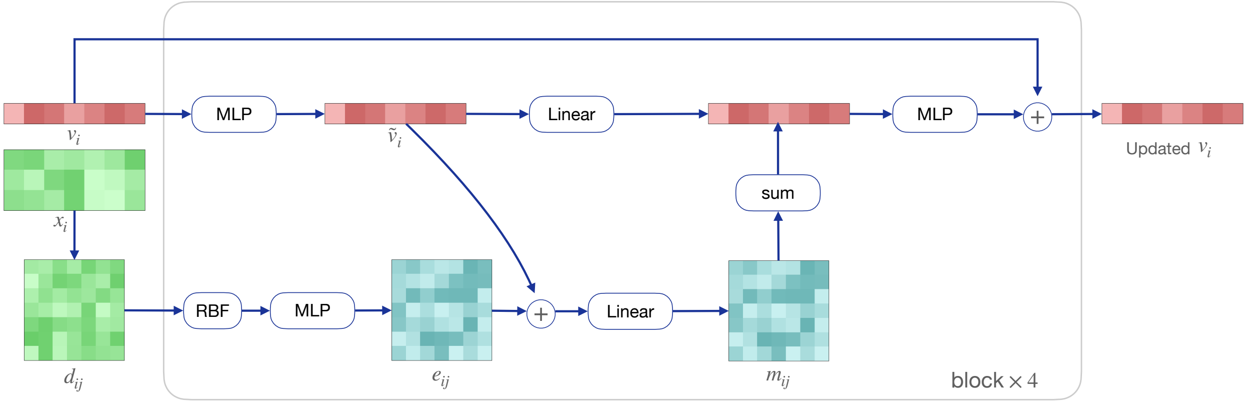

**Figure S1.** Architecture of PocketEncoder (Algorithm 2).

##### *1.1.2 MoleculeEncoder architecture*

The MoleculeEncoder is the central module of the denoiser to encode the atoms on either small molecules or peptides conditioned on the pocket atom representations derived from the PocketEncoder. It takes as input the molecule, the updated pocket features, and the task prompt (see Supplementary Notes 2 for details) and constructs a complete graph $\mathcal{G}_{\text{mol}}$ using the molecular atoms and $k$-NN dynamic bipartite graphs $\mathcal{G}_{\text{mol-poc}}$ between the molecule atoms and the pocket atoms ($k = 32$ in our implementation) to learn the node features, node coordinates and the edge features of the molecule $\mathcal{M}$, as defined in Algorithm 3.

**Algorithm 3 MoleculeEncoder:** update the node and edge feature and node coordinate of the molecule

**Input:** molecule $\mathcal{M} = \{\mathbf{A}, \mathbf{X}, \mathbf{B}\}$, pocket $\{\mathbf{V}^{\text{poc}}, \mathbf{X}^{\text{poc}}\}$, task prompt $\mathcal{P} = \{\mathbf{P}^{\text{atom}}, \mathbf{P}^{\text{coor}}, \mathbf{P}^{\text{bond}}, \mathbf{P}^{\text{dist}}, \mathbf{P}^{\text{pep}}\}$.

**Output:** molecular node (atom) features $\mathbf{V} \in \mathbb{R}^{N \times 320}$, updated node coordinates $\mathbf{X} \in \mathbb{R}^{N \times 3}$, molecular edge (atom pair) features $\mathbf{E} \in \mathbb{R}^{N \times N \times 96}$.

1: ▷ Input features
2: $\mathbf{V}^{\text{extra}} = \text{concat}(\mathbf{P}^{\text{atom}}, \mathbf{P}^{\text{coor}})$ ▷ $\mathbf{V}^{\text{extra}} \in \{0,1\}^{N \times 2}$
3: $\mathbf{E}^{\text{extra}} = \text{concat}(\mathbf{P}^{\text{bond}}, \mathbf{P}^{\text{dist}})$ ▷ $\mathbf{E}^{\text{extra}} \in \{0,1\}^{N \times N \times 2}$
4: $\mathbf{V} = \text{concat}(\text{Linear}(\mathbf{A}), \mathbf{P}^{\text{atom}}, \mathbf{P}^{\text{coor}}, \mathbf{P}^{\text{pep}})$ ▷ $\mathbf{V} \in \mathbb{R}^{N \times 320}$
5: $\mathbf{E} = \text{concat}(\text{Linear}(\mathbf{B}), \mathbf{P}^{\text{bond}}, \mathbf{P}^{\text{dist}})$ ▷ $\mathbf{E} \in \mathbb{R}^{N \times N \times 96}$
6: $\mathcal{G}_{\text{mol}} \leftarrow$ complete graph of molecule $\mathcal{M}$
7: **for all** block in $[1, 2, \ldots, 6]$ **do**
8: ▷ Update distance features
9: $\mathbf{E}^{\text{dist}}, \vec{\mathbf{R}}, \mathbf{D} = \text{DistanceBlock}(\mathcal{G}_{\text{mol}}, \mathbf{X})$ ▷ feature dimension of $\mathbf{E}^{\text{dist}}$ is 32
10: $\mathbf{E} \leftarrow \text{Linear}\left(\text{concat}(\mathbf{E}, \mathbf{E}^{\text{dist}})\right)$ ▷ feature dimension of $\mathbf{E}$ is 96
11: ▷ Update molecule-pocket edge features
12: $\mathcal{G}_{\text{mol-poc}} \leftarrow$ $k$-nearest neighbor bipartite graph ($k = 32$) from molecule $\mathbf{X}$ to pocket $\mathbf{X}^{\text{poc}}$
13: $\mathbf{E}^{\text{mol-poc}}, \vec{\mathbf{R}}^{\text{mol-poc}}, \mathbf{D}^{\text{mol-poc}} = \text{DistanceBlock}(\mathcal{G}_{\text{mol}}, \{\mathbf{X}, \mathbf{X}^{\text{poc}}\})$ ▷ feature dimension is 32
14: $\mathbf{E}^{\text{mol-poc}} \leftarrow \text{Linear}(\mathbf{E}^{\text{mol-poc}})$ ▷ feature dimension of $\mathbf{E}^{\text{mol-poc}}$ is 128
15: ▷ Update node features, edge features and coordinates
16: $\mathbf{V} \leftarrow \text{NodeBlock}(\mathbf{V}, \mathbf{E}, \mathcal{G}_{\text{mol}}, \mathbf{V}^{\text{poc}}, \mathbf{E}^{\text{mol-poc}}, \mathcal{G}_{\text{mol-poc}}, \mathbf{V}^{\text{extra}})$
17: $\mathbf{E} \leftarrow \text{EdgeBlock}(\mathbf{E}, \mathbf{V}, \mathbf{E}^{\text{extra}})$
18: $\mathbf{X} \leftarrow \mathbf{X} + \text{PosBlock}(\mathbf{V}, \mathbf{E}, \vec{\mathbf{R}}, \mathbf{D}, \mathbf{V}^{\text{extra}}, \mathbf{E}^{\text{extra}})$
19: $\mathbf{X} \leftarrow \mathbf{X} + \text{PosBlockFromPocket}(\mathbf{V}, \mathbf{E}^{\text{mol-poc}}, \vec{\mathbf{R}}^{\text{mol-poc}}, \mathbf{D}^{\text{mol-poc}}, \mathbf{V}^{\text{poc}})$
20: **end for**

The DistanceBlock function is designed to calculate the features, relative vectors, and distances of edges in a graph, as defined in Algorithm 4. The RBF of the DistanceBlock used in Line 9 and Line 13 of Algorithm 3 are within the intervals $[0, 15]$ and $[0, 20]$, respectively.

**Algorithm 4 DistanceBlock:** calculate distance features of edges

**Input:** a graph $\mathcal{G}$, node coordinates $\mathbf{X}$.

**Output:** RBF features $\mathbf{E} = [\mathbf{e}_{ij}]$ of edge distance, edge relative vectors $\vec{\mathbf{R}} = [\vec{\mathbf{r}}_{ij}]$, edge distances $\mathbf{D} = [d_{ij}]$.

```
for all ij ∈ edges of 𝒢 do
    r⃗_ij = x_i − x_j
    d_ij = ‖r⃗_ij‖
    e_ij = RBF(d_ij)
end for
```

**for all** $ij \in$ edges of $\mathcal{G}$ **do**
- $\vec{\mathbf{r}}_{ij} = \mathbf{x}_i - \mathbf{x}_j$
- $d_{ij} = \|\vec{\mathbf{r}}_{ij}\|$
- $\mathbf{e}_{ij} = \mathrm{RBF}(d_{ij})$

**end for**

The NodeBlock updates the molecular node features $\mathbf{V}$ using messages from other nodes in both $\mathcal{G}_{\text{mol}}$ and $\mathcal{G}_{\text{mol-poc}}$, as defined in Algorithm 5 (Figure S2). The hidden and output dimensions of all the Linear functions and MLP functions are 320.

**Algorithm 5 NodeBlock**: update molecular node features

**Input:** molecular node features $\mathbf{V}$, molecular edge features $\mathbf{E}$, molecular graph $\mathcal{G}_{\text{mol}}$, pocket node features $\mathbf{V}^{\text{poc}}$, edge feature of molecule-pocket graph, molecule-pocket graph $\mathcal{G}_{\text{mol-poc}}$, extra molecular node features $\mathbf{V}^{\text{extra}}$.

**Output:** updated molecular node features $\mathbf{V}$.

▷ Message in $\mathcal{G}_{\text{mol}}$

**for all** $i \in$ nodes , $ij \in$ edges of $\mathcal{G}_{\text{mol}}$ **do**
- $\tilde{\mathbf{v}}_i = \mathrm{MLP}(\mathbf{v}_i)$
- $\mathbf{m}_{ij} = \mathrm{Linear}(\mathrm{MLP}(\mathbf{e}_{ij}) + \tilde{\mathbf{v}}_i + \tilde{\mathbf{v}}_j)$
- $\mathbf{g}_{ij} = \mathrm{MLP}\left(\mathrm{concat}(\mathbf{e}_{ij}, \mathbf{v}_i, \mathbf{v}_j, \mathbf{v}_i^{\text{extra}}, \mathbf{v}_j^{\text{extra}})\right)$

**end for**

▷ Message in $\mathcal{G}_{\text{mol-poc}}$

**for all** $k \in$ pocket nodes, $ik \in$ edges of $\mathcal{G}_{\text{mol-poc}}$ **do**
- $\tilde{\mathbf{v}}_k^{\text{poc}} = \mathrm{MLP}(\mathbf{v}_k^{\text{poc}})$
- $\mathbf{m}_{ik}^{\text{mol-poc}} = \mathrm{Linear}\left(\mathrm{MLP}(\mathbf{e}_{ik}^{\text{mol-poc}}) \odot \tilde{\mathbf{v}}_k^{\text{poc}})\right)$
- $\mathbf{g}_{ik}^{\text{mol-poc}} = \mathrm{MLP}\left(\mathrm{concat}(\mathbf{e}_{ik}^{\text{mol-poc}}, \mathbf{v}_i, \mathbf{v}_k^{\text{poc}}, \mathbf{v}_i^{\text{extra}})\right)$

**end for**

▷ Aggregate all messages

**for all** $i \in$ nodes of $\mathcal{G}_{\text{mol}}$ **do**
- $\tilde{\mathbf{v}}_i \leftarrow \mathrm{Linear}(\tilde{\mathbf{v}}_i) + \sum_{ij \in \mathcal{G}_{\text{mol}}} \mathbf{m}_{ij} \odot \mathrm{sigmoid}(\mathbf{g}_{ij}) + \sum_{ik \in \mathcal{G}_{\text{mol-poc}}} \mathbf{m}_{ik}^{\text{mol-poc}} \odot \mathrm{sigmoid}(\mathbf{g}_{ik}^{\text{mol-poc}})$
- $\mathbf{v}_i \leftarrow \mathbf{v}_i + \mathrm{LayerNorm}\left(\mathrm{MLP}(\tilde{\mathbf{v}}_i)\right)$

**end for**

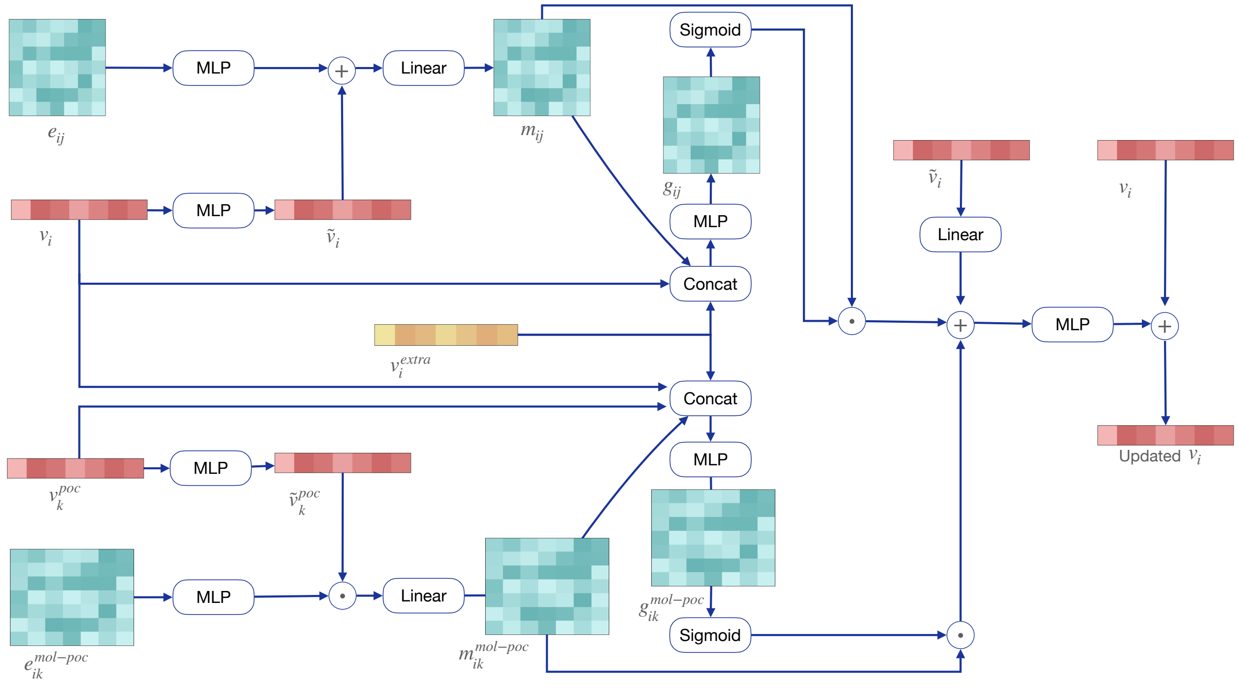

**Figure S2.** Architecture of NodeBlock (Algorithm 5).

The EdgeBlock updates the molecular edge features $\mathbf{E}$ using messages from other edges connecting to the same ends in $\mathcal{G}_{\text{mol}}$, as defined in Algorithm 6 (Figure S3). The feature dimension of the input $\mathbf{E}$ is 96. The output dimensions of the Linear functions in Line 3 and Line 8 are 192. The hidden dimensions of the MLPs in Line 3 and Line 8 are 192. The hidden dimensions of the MLPs in Line 4 and Line 9 are 32. The output dimensions of the MLPs in Line 3, 4, 8, and 9 are 96. The hidden and output dimensions in Line 13 and Line 14 are 96.

**Algorithm 6 EdgeBlock**: update molecular edge features

**Input:** molecular edge features $\mathbf{E}$, molecular node features $\mathbf{V}$, extra molecular node features $\mathbf{E}^{\text{extra}}$.

**Output:** updated molecular edge features. $\mathbf{E}$.

**for all** $ij \in$ edges of $\mathbf{E}$ **do**
- ▷ Message from left ends
- $\mathbf{e}_{ij}^{\text{left}} = \text{MLP}\left(\text{Linear}(\mathbf{e}_{ij}) + \text{Linear}(\mathbf{v}_i)\right)$
- $\mathbf{g}_{ij}^{\text{left}} = \text{MLP}\left(\text{concat}(\mathbf{e}_{ij}, \mathbf{v}_i, \mathbf{v}_i^{\text{extra}})\right)$
- $\mathbf{e}_{ij}^{\text{left}} \leftarrow \mathbf{e}_{ij}^{\text{left}} \odot \text{sigmoid}(\mathbf{g}_{ij}^{\text{left}})$
- $\mathbf{m}_i^{\text{left}} = \sum_l \mathbf{e}_{li}^{\text{left}}$
- ▷ Message from right ends
- $\mathbf{e}_{ij}^{\text{right}} = \text{MLP}\left(\text{Linear}(\mathbf{e}_{ij}) + \text{Linear}(\mathbf{v}_j)\right)$
- $\mathbf{g}_{ij}^{\text{right}} = \text{MLP}\left(\text{concat}(\mathbf{e}_{ij}, \mathbf{v}_j, \mathbf{v}_j^{\text{extra}})\right)$
- $\mathbf{e}_{ij}^{\text{right}} \leftarrow \mathbf{e}_{ij}^{\text{right}} \odot \text{sigmoid}(\mathbf{g}_{ij}^{\text{right}})$
- $\mathbf{m}_j^{\text{right}} = \sum_l \mathbf{e}_{jl}^{\text{right}}$
- ▷ Aggregate all messages
- $\tilde{\mathbf{e}}_{ij} = \text{Linear}(\mathbf{e}_{ij}) + \text{Linear}(\mathbf{v}_i) + \text{Linear}(\mathbf{v}_j) + \text{Linear}(\mathbf{m}_i^{\text{left}}) + \text{Linear}(\mathbf{m}_j^{\text{right}})$
- $\mathbf{e}_{ij} \leftarrow \mathbf{e}_{ij} + \text{LayerNorm}(\text{MLP}(\tilde{\mathbf{e}}_{ij}))$

**end for**

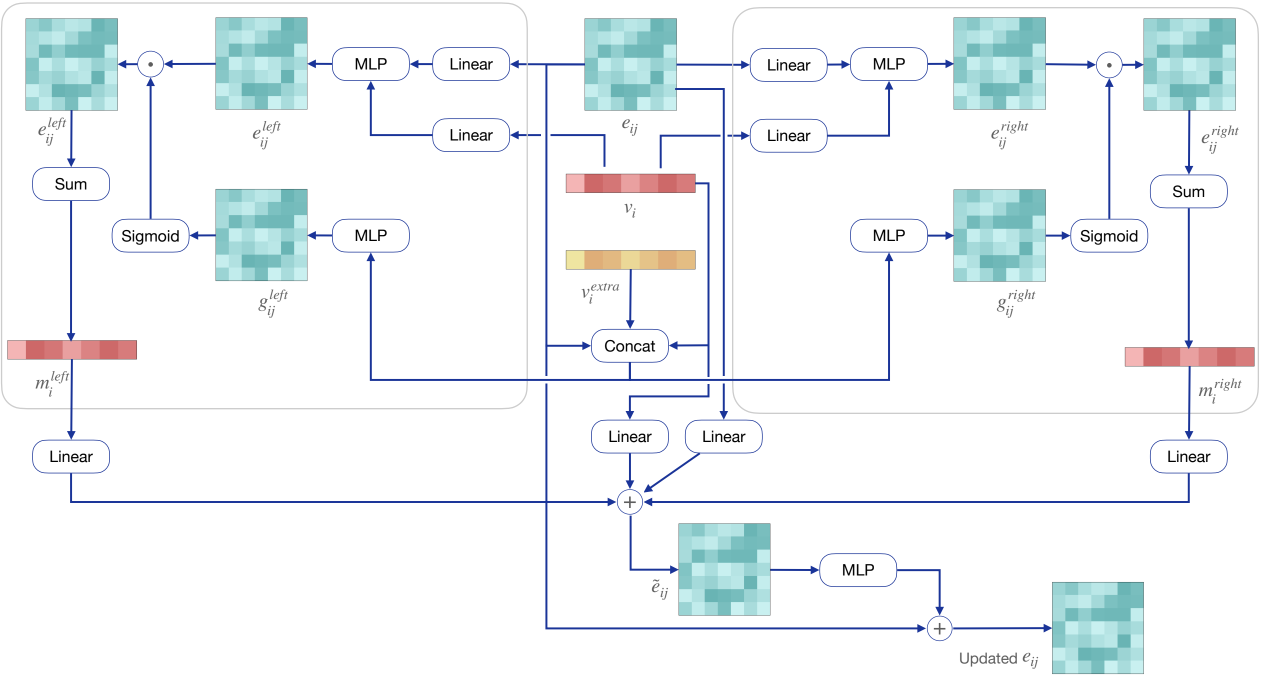

**Figure S3. Architecture of EdgeBlock (Algorithm 6).**

The PosBlock predicts the coordinate changes $\Delta\mathbf{x}$ using relative vectors among atoms in $\mathbf{G}_{\text{mol}}$, as defined in Algorithm 7 (Figure S4). The variable $d_0$ in Line 12 is a hyper-parameter as a scaling for edge weights and set as $d_0 = 5$ in the implementation. The hidden and output dimensions of the MLPs in Line 1 are 96 and 320, respectively. The output dimensions of the Linear functions and the hidden dimensions of the MLPs in Line 6 are 320. The hidden dimension of the MLP in Line 7 is 32. The hidden dimension of the MLP in Line 13 is 96. The output dimensions of the MLPs in Line 6, 7, and 13 are 1.

---

**Algorithm 7 PosBlock:** update molecular node coordinates in $\mathcal{G}_{\text{mol}}$

---

**Input:** molecular node features $\mathbf{V}$, molecular edge features $\mathbf{E}$, molecular edge vectors $\vec{\mathbf{R}}$, molecular edge distances $\mathbf{D}$, extra molecular node features $\mathbf{V}^{\text{extra}}$, extra molecular edge features $\mathbf{E}^{\text{extra}}$.

**Output:** changes of molecular node coordinates $\Delta\mathbf{X}$.

1: **for all** $ij \in$ edges of $\mathbf{E}$ **do**
2:     ▷ Integrate two ends into edges
3:     $\tilde{\mathbf{e}}_{ij} = \text{concat}\left(\text{MLP}(\mathbf{v}_i), \text{MLP}(\mathbf{v}_j)\right)$
4:     $\tilde{\mathbf{e}}_{ij}^{\text{extra}} = \text{concat}(\mathbf{v}_i^{\text{extra}}, \mathbf{e}_{ij}^{\text{extra}})$
5:     ▷ Predict edge weights
6:     $w_{ij} = \text{MLP}\left(\text{Linear}(\mathbf{e}_{ij}), \text{Linear}(\tilde{\mathbf{e}}_{ij})\right)$
7:     $g_{ij} = \text{MLP}\left(\text{concat}(\mathbf{e}_{ij}, \tilde{\mathbf{e}}_{ij}, \tilde{\mathbf{e}}_{ij}^{\text{extra}})\right)$
8:     $w_{ij} = w_{ij} \odot \text{sigmoid}(g_{ij})$
9: **end for**
10: **for all** $i \in$ nodes of $\mathbf{V}$ **do**
11:     ▷ Predict delta positions
12:     $\Delta\mathbf{x}_i = \sum_j \left(\frac{w_{ij}}{d_{ij}/d_0+1} \cdot \frac{\vec{\mathbf{r}}_{ij}}{d_{ij}}\right)$     ▷ $d_0 = 5$ is a scaling hyper-parameter
13:     $\tilde{g}_i = \text{MLP}\left(\text{concat}(\mathbf{v}_i, \mathbf{v}_i^{\text{extra}}, \|\Delta\mathbf{x}_i\|)\right)$
14:     $\Delta\mathbf{x}_i \leftarrow \text{sigmoid}(\tilde{g}_i) \cdot \Delta\mathbf{x}_i$
15: **end for**

---

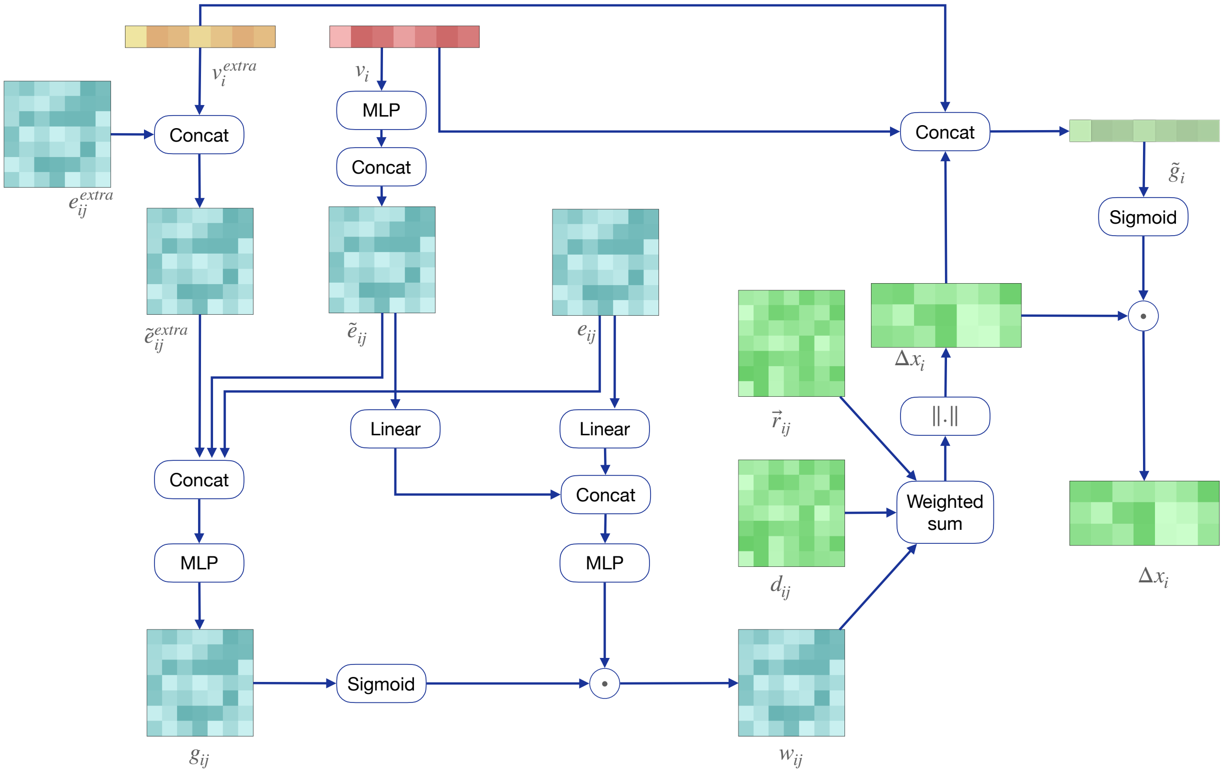

**Figure S4.** **Architecture of PosBlock (Algorithm 7).**

The PosBlockFromPocket works similarly to the PosBlock but uses the relative vectors of the edges in $\mathcal{G}_{\text{mol-poc}}$, as defined in Algorithm 8 (Figure S5). Here, the value of $d_0$ and the hidden dimensions and output dimensions of the MLPs and Linear functions are the same as the counterparts in the PosBlock.

**Algorithm 8 PosBlockFromPocket:** update molecular node coordinates in $\mathcal{G}_{\text{mol-poc}}$

**Input:** molecular node features $\mathbf{V}$, molecule-pocket edge features $\mathbf{E}^{\text{mol-poc}}$, molecule-pocket edge vectors $\vec{\mathbf{R}}^{\text{mol-poc}}$, molecule-pocket edge distances $\mathbf{D}^{\text{mol-poc}}$, pocket node features $\mathbf{V}^{\text{poc}}$.

**Output:** changes of molecular node coordinates $\Delta\mathbf{X}$.

**for all** $ik \in$ edges of $\mathbf{E}^{\text{mol-poc}}$ **do**
    ▷ Integrate two ends into edges
    $\tilde{\mathbf{e}}_{ik} = \text{concat}\left(\text{MLP}(\mathbf{v}_i), \text{MLP}(\mathbf{v}_k^{\text{poc}})\right)$
    $\tilde{\mathbf{e}}_{ik}^{\text{extra}} = \mathbf{v}_i^{\text{extra}}$
    ▷ Predict edge weights)
    $w_{ik} = \text{MLP}\left(\text{Linear}(\mathbf{e}_{ik}^{\text{mol-poc}}), \text{Linear}(\tilde{\mathbf{e}}_{ik})\right)$
    $g_{ik} = \text{MLP}\left(\text{concat}(\mathbf{e}_{ij}^{\text{mol-poc}}, \tilde{\mathbf{e}}_{ik}, \tilde{\mathbf{e}}_{ik}^{\text{extra}})\right)$
    $w_{ik} = w_{ik} \odot \text{sigmoid}(g_{ik})$
**end for**
**for all** $i \in$ nodes of $\mathbf{V}$ **do**
    ▷ Predict delta positions
    $\Delta\mathbf{x}_i = \sum_k \left(\frac{w_{ik}}{d_{ik}^{\text{mol-poc}}/d_0+1} \cdot \frac{\vec{\mathbf{r}}_{ik}^{\text{mol-poc}}}{d_{ik}^{\text{mol-poc}}}\right)$
    $\tilde{g}_i = \text{MLP}\left(\text{concat}(\mathbf{v}_i, \mathbf{v}_i^{\text{extra}}, \|\Delta\mathbf{x}_i\|)\right)$
    $\Delta\mathbf{x}_i \leftarrow \text{sigmoid}(\tilde{g}_i) \cdot \Delta\mathbf{x}_i$
**end for**

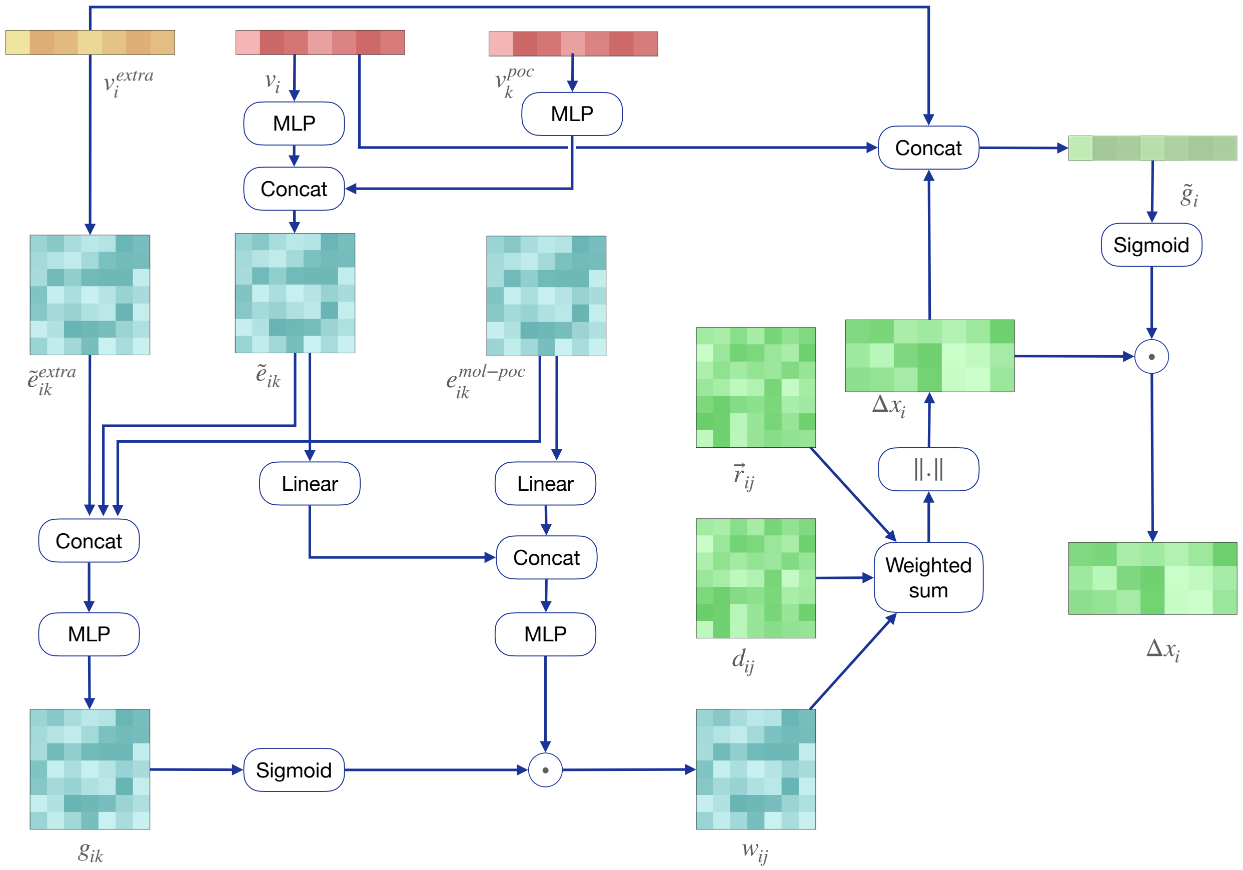

**Figure S5. Architecture of PosBlockFromPocket (Algorithm 8).**

364 ## 1.2 Decoder architectures

Here we introduce the three decoders AtomDecoder, CoordinateDecoder, and BondDecoder. The AtomDecoder takes the updated node features $\mathbf{V}$ as inputs and predicts the probabilities of atom types and model confidences, as defined in Algorithm 9. The hidden dimensions of the MLPs are 320 and 160, respectively. The output dimensions of the MLPs are equal to the number of atom types (i.e., 12, including the mask type) and 1, respectively.

---

**Algorithm 9 AtomDecoder:** predict atom types

**Input:** molecular node features $\mathbf{V}$.

**Output:** predicted probabilities of atom types $\mathbf{A}^{\text{raw}}$, confidence scores $\mathbf{S}^{\text{atom}}$ for atom types.

1: **for all** $i \in$ nodes of $\mathbf{V}$ **do**

2: $\quad \mathbf{a}_i^{\text{raw}} = \text{softmax}\left(\text{MLP}(\mathbf{v}_i)\right)$

3: $\quad s_i^{\text{atom}} = \text{sigmoid}\left(\text{MLP}(\mathbf{v}_i)\right)$

4: **end for**

---

The CoordinateDecoder takes the molecular node features $\mathbf{V}$ and coordinates $\mathbf{X}$ as inputs to predict the coordinates and the confidence scores, as defined in Algorithm 10. The hidden and output dimensions of the MLP are 160 and 1, respectively.

**Algorithm 10 CoordinateDecoder:** predict atom coordinates

**Input:** molecular node features $\mathbf{V}$, molecular node coordinates $\mathbf{X}$.

**Output:** predicted atom coordiantes $\mathbf{X}^{\text{raw}}$, confidence scores $\mathbf{S}^{\text{coor}}$ for atom coordinates.

```
for all i ∈ nodes of V do
    x_i^raw = x_i
    s_i^coor = sigmoid (MLP(v_i))
end for
```

The BondDecoder takes the molecular edge features $\mathbf{E}$ as inputs to predict the probabilities of bond types and confidence scores, as defined in Algorithm 11. The hidden dimensions of the MLPs are 96 and 48, respectively. The output dimensions of the MLPs are equal to the number of bond types (i.e., 6, including the mask type) and 1, respectively.

**Algorithm 11 BondDecoder:** predict bond types

**Input:** molecular edge features $\mathbf{E}$.

**Output:** predicted probabilities of bond types $\mathbf{E}^{\text{raw}}$, confidence scores $\mathbf{S}^{\text{bond}}$ for bond types.

```
for all ij ∈ edges of E do
    b_ij^raw = softmax (MLP(e_ij + e_ji))
    s_ij^bond = sigmoid (MLP(e_ij + e_ji))
end for
```

The learnable parameters of the denoiser are all in the encoders and the decoders. The outputs of the denoiser, named un-adapted denoised molecules, are used to calculate the loss function. We use $F_\Theta^{\text{raw}}$ to represent the combination of the encoders and the decoders:

$$\hat{\mathcal{M}}^{\text{raw}}, \hat{\mathcal{S}} = F_\Theta^{\text{raw}}(\mathcal{M}, \mathcal{P}, \xi, \beta) \tag{S1}$$

where $\hat{\mathcal{M}}^{\text{raw}} = \{\mathbf{A}^{\text{raw}}, \mathbf{X}^{\text{raw}}, \mathbf{B}^{\text{raw}}\}$.

#### 1.3 M-Projector

The M-Projector consists of two functions: 1) derive the atom and bond types from the predicted probabilities by choosing the types with the maximum probabilities and drop the atoms with the mask types; 2) adapt the predicted variables to align with the task prompt, i.e., the output variables with fixed prompt indicators must be consistent with the inputs. Although the neural networks are trained to keep these variables unchanged, they do not always follow the task prompt during the generation process. Therefore, an adaptation module is required to correct the output variables of the decoders to produce the final molecules.

Specifically, the atom types $\hat{a}_i$ and bond types $\hat{b}_{ij}$ are first derived as follows,

$$\begin{aligned} \hat{a}_i &= \arg\max \mathbf{a}_i^{\text{raw}} \\ \hat{b}_{ij} &= \arg\max \mathbf{b}_{ij}^{\text{raw}} \end{aligned} \tag{S2}$$

Then the atom and bond types were adapted by choosing from the types $a_i$ or $b_{ij}$ of the input molecule to the denoiser if the corresponding prompt indicators were fixed ($P_i^{\text{atom}} = 1$ or $P_{ij}^{\text{bond}} = 1$), which is defined as the follows,

$$\begin{aligned} \hat{a}_i &\leftarrow a_i P_i^{\text{atom}} + \hat{a}_i(1 - P_i^{\text{atom}}) \\ \hat{b}_{ij} &\leftarrow b_{ij} P_{ij}^{\text{bond}} + \hat{b}_{ij}(1 - P_{ij}^{\text{bond}}) \end{aligned} \tag{S3}$$

For the adaption of the atom coordinates, the prompts for both the coordinates $\mathbf{P}^{\text{coor}}$ and the pairwise distances $\mathbf{P}^{\text{dist}}$ should be considered. For atoms with $P_i^{\text{coor}} = 1$, their coordinates should be fixed and thus are directly adapted as the corresponding ones $\mathbf{x}_i$ of the input molecule to the denoiser. For atoms with unfixed coordinates and unfixed distances with other atoms, i.e. $P_i^{\text{coor}} = 0$ and $\sum_j P_{ij}^{\text{dist}} = 0$, their predicted coordinates $\hat{\mathbf{x}}_i$ do not need to be adapted. Otherwise, things are more difficult and the coordinates are adapted as $\hat{\mathbf{x}}_i'$ using a more complicated algorithm explained below.

$$\hat{\mathbf{x}}_i \leftarrow \begin{cases} \mathbf{x}_i & \text{if } P_i^{\text{coor}} = 1 \\ \hat{\mathbf{x}}_i & \text{if } P_i^{\text{coor}} = 0 \text{ and } \sum_j P_{ij}^{\text{dist}} = 0 \\ \hat{\mathbf{x}}_i' & \text{if } P_i^{\text{coor}} = 0 \text{ and } \sum_j P_{ij}^{\text{dist}} > 0 \end{cases} \tag{S4}$$

Now we introduce how to derive $\hat{\mathbf{x}}_i'$ for atoms with $P_i^{\text{coor}} = 0, \sum_j P_{ij}^{\text{dist}} > 0$. In this case, the atoms with fixed distances should be considered as an entire entity for adaptation. To guarantee the fixed atom distances, the adapted coordinates should be obtained through the rigid transformations and rotations of rotatable bonds from the input noisy molecule. The basic idea is to compare the predicted coordinates and the input coordinates of the denoiser to obtain the torsion angle changes of the rotatable bonds and the translation vector and rotation matrix of rigid transformation. Then the rotation of rotatable bonds and rigid transformation is directly applied to the input coordinates to produce the final adapted coordinates, as defined in Algorithm 12.

**Algorithm 12 AdaptCoordinateFixedDist:** adapt the 3D coordinates with fixed distances

**Input:** input molecule $\mathcal{M} = \{\mathbf{A}, \mathbf{X}, \mathbf{B}\}$, predicted coordinates $\mathbf{X}^{\text{raw}}$.

**Output:** adapted coordinates $\hat{\mathbf{X}}$.

```
1: 𝔹^chem, 𝔹^rot ← bond index set of chemical bonds and rotatable bonds of 𝓜
2: ▷ Calculate torsion angle changes of rotatable bonds
3: sin Δω, cos Δω = CalcTorsionAngleChanges(𝔹^chem, 𝔹^rot, X, X^raw)
4: ▷ Apply rotation to rotatable bonds
5: X̂ = RotBonds(A, 𝔹^chem, 𝔹^rot, X, {sin Δω, cos Δω})
6: ▷ Calculate translation vector and rotation matrix of rigid transformation
7: t, R = AlignStructure(X̂, X^raw)                      ▷ t ∈ ℝ^3, R ∈ ℝ^{3×3}
8: ▷ Apply rigid transformation
9: for all x̂_i ∈ X̂ do
10:     x̂_i ← R x̂_i + t
11: end for
```

The CalcTorsionAngleChanges (Algorithm 13) compares the torsion angles of rotatable bonds between the two coordinate sets to derive the changes of the torsion angles, where the torsion angles are calculated as an average of all dihedral angles formed by the rotatable bonds.

**Algorithm 13 CalcTorsionAngleChanges:** calculate torsion angle changes between two sets of molecular coordinates

**Input:** chemical bond set $\mathbb{B}^{\text{chem}}$, rotatable bond set $\mathbb{B}^{\text{rot}}$, predicted and input coordinates $\hat{\mathbf{X}}, \mathbf{X}$.

**Output:** torsion angle changes of rotatable bonds $\{\sin \Delta\boldsymbol{\omega}, \cos \Delta\boldsymbol{\omega}\}$.

```
for all ij ∈ B^rot do
    ▷ Calculate all related dihedral angle changes
    for all ki ∈ B^chem \ {ji}, jm ∈ B^chem \ {ji} do
        sin ψ_kijm, cos ψ_kijm = CalcDihedralAngle(x_k, x_i, x_j, x_m)
        sin ψ̂_kijm, cos ψ̂_kijm = CalcDihedralAngle(x̂_k, x̂_i, x̂_j, x̂_m)
        cos Δψ_kijm = cos(ψ̂_kijm − ψ_kijm) = cos ψ̂_kijm cos ψ_kijm + sin ψ̂_kijm sin ψ_kijm
        sin Δψ_kijm = sin(ψ̂_kijm − ψ_kijm) = sin ψ̂_kijm cos ψ_kijm − cos ψ̂_kijm sin ψ_kijm
    end for
    ▷ Average dihedral angle changes as torsion angle changes
    sin Δω_ij = mean_kl sin Δψ_kijm
    cos Δω_ij = mean_kl cos Δψ_kijm
    n_ij = sqrt(sin² Δω_ij + cos² Δω_ij)
    sin Δω_ij ← sin Δω_ij / n_ij, cos Δω_ij ← cos Δω_ij / n_ij        ▷ normalize
end for
sin Δω = {sin Δω_ij | ij ∈ B^rot}
cos Δω = {cos Δω_ij | ij ∈ B^rot}
```

The CalcDihedralAngle (Algorithm 14) calculates the dihedral angle formed by the four input atoms.

**Algorithm 14 CalcDihedralAngle:** calculate a dihedral angle

**Input:** the atom coordinates $[\mathbf{x}_0, \mathbf{x}_1, \mathbf{x}_2, \mathbf{x}_3]$

**Output:** sine and cosine of dihedral angle $\psi$ formed by the atoms $\{\sin \psi, \cos \psi\}$

$\vec{\mathbf{r}}_0 = \mathbf{x}_0 - \mathbf{x}_1$

$\vec{\mathbf{r}}_1 = (\mathbf{x}_1 - \mathbf{x}_2)/\|\mathbf{x}_1 - \mathbf{x}_2\|$

$\vec{\mathbf{r}}_2 = \mathbf{x}_3 - \mathbf{x}_2$

$\vec{\mathbf{r}}_0 \leftarrow \vec{\mathbf{r}}_0 - (\vec{\mathbf{r}}_0 \cdot \vec{\mathbf{r}}_1)\vec{\mathbf{r}}_1$

$\vec{\mathbf{r}}_2 \leftarrow \vec{\mathbf{r}}_2 - (\vec{\mathbf{r}}_2 \cdot \vec{\mathbf{r}}_1)\vec{\mathbf{r}}_1$

$\vec{\mathbf{r}}_0 \leftarrow \vec{\mathbf{r}}_0/\|\vec{\mathbf{r}}_0\|$ ▷ Normalize

$\vec{\mathbf{r}}_2 \leftarrow \vec{\mathbf{r}}_2/\|\vec{\mathbf{r}}_2\|$ ▷ Normalize

$\cos \psi = \vec{\mathbf{r}}_0 \cdot \vec{\mathbf{r}}_2$

$\sin \psi = \vec{\mathbf{r}}_0 \times \vec{\mathbf{r}} \cdot \vec{\mathbf{r}}_1$

The RotBonds (Algorithm 15) applies the rotations to all torsion angles of rotatable bonds. In

Line 7, the notation $\|\text{path}(\cdot,\cdot)\|$ denotes the length of the shortest path between two nodes on a graph.

**Algorithm 15 RotBonds:** rotate the bonds

**Input:** chemical bond set $\mathbb{B}^{\text{chem}}$, rotatable bond set $\mathbb{B}^{\text{rot}}$, atom coordinates $\mathbf{X}$, torsion angles changes $\{\sin\Delta\boldsymbol{\omega}, \cos\Delta\boldsymbol{\omega}\}$.

**Output:** updated coordinates $\tilde{\mathbf{X}}$.

1: $N \leftarrow$ the number of atoms of $\mathbf{X}$
2: $\mathbb{A} = \{i | i \in [\![N]\!]\}$
3: $\tilde{\mathbf{X}} \leftarrow \mathbf{X}$
4: **for all** $ij \in \mathbb{B}^{\text{rot}}$ **do**
5:     $\vec{\mathbf{r}}_{\text{axis}} = (\tilde{\mathbf{x}}_i - \tilde{\mathbf{x}}_j)/\|\tilde{\mathbf{x}}_i - \tilde{\mathbf{x}}_j\|$ ▷ rotation axis
6:     **for all** $k \in \mathbb{A} \setminus \{i, j\}$ **do**
7:         **if** $\|\text{path}(k,i)\| > \|\text{path}(k,j)\|$ on the graph composed of $\mathbb{B}^{\text{chem}}$ **then**
8:             ▷ atom $k$ is at the atom $j$ side of the bond $ij$
9:             $\vec{\mathbf{r}}_{\text{vec}} = \tilde{\mathbf{x}}_k - \tilde{\mathbf{x}}_i$
10:            $\vec{\mathbf{r}}_{\text{radius}} = \vec{\mathbf{r}}_{\text{vec}} - (\vec{\mathbf{r}}_{\text{vec}} \cdot \vec{\mathbf{r}}_{\text{axis}})\vec{\mathbf{r}}_{\text{axis}}$
11:            $\vec{\mathbf{r}}_{\text{center}} = \tilde{\mathbf{x}}_k - \vec{\mathbf{r}}_{\text{radius}}$
12:            $\tilde{\mathbf{x}}_k \leftarrow \text{RotAxisAngle}(\vec{\mathbf{r}}_{\text{radius}}, \vec{\mathbf{r}}_{\text{axis}}, \{\sin\Delta\omega_{ij}, \cos\Delta\omega_{ij}\}) + \vec{\mathbf{r}}_{\text{center}}$
13:        **end if**
14:    **end for**
15: **end for**

The RotAxisAngle uses the Rodrigues' rotation formula to rotate a vector $\vec{\mathbf{r}}_{\text{radius}}$ along the axis $\vec{\mathbf{r}}_{\text{axis}}$ by the angle $\Delta\omega$:

$$\vec{\mathbf{r}}_{\text{radius}} \leftarrow \vec{\mathbf{r}}_{\text{radius}} \cos\Delta\omega + (\vec{\mathbf{r}}_{\text{axis}} \times \vec{\mathbf{r}}_{\text{radius}}) \sin\Delta\omega + \vec{\mathbf{r}}_{\text{axis}}(\vec{\mathbf{r}}_{\text{axis}} \cdot \vec{\mathbf{r}}_{\text{radius}})(1 - \cos\Delta\omega) \tag{S5}$$

The AlignStructure (Algorithm 16) uses the Kabsch algorithm to calculate the translation vector and rotation matrix that can superimpose a set of coordinates to another.

**Algorithm 16 AlignStructure:** get the rigid transformation to align two sets of coordinates

**Input:** the two sets of coordinates $\mathbf{X} \in \mathbb{R}^{N\times 3}, \mathbf{Y} \in \mathbb{R}^{N\times 3}$.

**Output:** the translation vector $\mathbf{t}$ and rotation matrix $\mathbf{R}$ of a rigid transformation from $\mathbf{X}$ to $\mathbf{Y}$.

$\triangleright$ Centering
$\bar{\mathbf{x}} = 1/N \sum_i \mathbf{x}_i$
$\bar{\mathbf{y}} = 1/N \sum_i \mathbf{y}_i$
$\mathbf{X} \leftarrow \mathbf{X} - \mathbf{1}_N^\top \bar{\mathbf{x}}$
$\mathbf{Y} \leftarrow \mathbf{Y} - \mathbf{1}_N^\top \bar{\mathbf{y}}$
$\triangleright$ Covariance matrix and SVD
$\mathbf{U}, \mathbf{\Sigma}, \mathbf{V}^\top = \mathrm{SVD}(\mathbf{X}\mathbf{Y}^\top)$
$d = \mathrm{sign}(\det(\mathbf{U}\mathbf{V}^\top))$
$\triangleright$ Produce rotation and translation
$\mathbf{R} = \mathbf{V} \begin{pmatrix} 1 & 0 & 0 \\ 0 & 1 & 0 \\ 0 & 0 & d \end{pmatrix} \mathbf{U}^\top$
$\mathbf{t} = \bar{\mathbf{y}} - \mathbf{R}\bar{\mathbf{x}}$

### 2 Task prompt

#### 2.1 Definition of the prompt components

For a molecule with $N$ atoms, the complete task prompt for the denoiser includes the three groups of main prompt indicators $\{\mathbf{P}^{\text{atom}}, \mathbf{P}^{\text{coor}}, \mathbf{P}^{\text{bond}}\}$ and two groups of auxiliary prompt indicators $\{\mathbf{P}^{\text{dist}}, \mathbf{P}^{\text{pep}}\}$. The main prompt is enough for expressing different tasks but the auxiliary prompt enables more precise control of specific settings of tasks. All the prompt indicators are vectors or matrices of binary variables. Formally, the prompt $\mathbf{P}^{\text{atom}} \in \{0,1\}^N$ represents whether the individual atom types of the input molecules should be fixed:

$$\forall i \in [\![N]\!],\ P_i^{\text{atom}} = \begin{cases} 1, & \text{atom type of atom } i \text{ is fixed} \\ 0, & \text{atom type of atom } i \text{ is unfixed} \end{cases} \tag{S6}$$

The prompt $\mathbf{P}^{\text{coor}} \in \{0,1\}^N$ represents whether the individual atom coordinates of the input molecules should be fixed:

$$\forall i \in [\![N]\!],\ P_i^{\text{corr}} = \begin{cases} 1, & \text{coordinate of atom } i \text{ is fixed} \\ 0, & \text{coordinate of atom } i \text{ is unfixed} \end{cases} \tag{S7}$$

The prompt $\mathbf{P}^{\text{bond}} \in \{0,1\}^{N\times N}$ represents whether the individual bond types of the input molecules should be fixed and is a symmetric matrix:

$$\forall i,j \in [\![N]\!],\ P_{ij}^{\text{bond}} = \begin{cases} 1, & \text{bond type of bond } ij \text{ is fixed} \\ 0, & \text{bond type of bond } ij \text{ is unfixed} \end{cases} \tag{S8}$$

The prompt $\mathbf{P}^{\text{dist}} \in \{0,1\}^{N\times N}$ represents whether the pairwise atom distances of the input molecules should be fixed and is a symmetric matrix:

$$\forall i,j \in [\![N]\!],\ P_{ij}^{\text{dist}} = \begin{cases} 1, & \text{distance of atom pair } ij \text{ is fixed} \\ 0, & \text{distance of atom pair } ij \text{ is unfixed} \end{cases} \tag{S9}$$

This auxiliary prompt is useful when we only want the atom coordinates to be denoised through translation, rotation, and rotation of rotatable bonds.

The prompt $\mathbf{P}^{\mathrm{pep}} \in \{0,1\}^N$ represents whether the atom type should compose standard amino acids when generating atom types. Simply, it is set as an all-one vector when generating peptide side chains, otherwise, it is an all-zero vector.

$$\mathbf{P}^{\mathrm{pep}} = \begin{cases} \mathbf{1}_N, & \text{generating atoms of standard amino acids} \\ \mathbf{0}_N, & \text{otherwise} \end{cases} \tag{S10}$$

This auxiliary prompt is useful when we want the denoiser to generate atoms composing standard amino acids.

We now introduce how the prompt is specified for each generative task for a molecule with $N$ atoms. In the equations, we abbreviate $\forall i \in [\![N]\!]$ and $\forall i,j \in [\![N]\!]$ as $\forall i$ and $\forall ij$, respectively.

#### 2.2 Task prompt for canonical tasks

Here, we show how we specify the task prompt for canonical tasks. Users can also utilize the task prompt to express more complicated and specific task settings by themselves.

##### 2.2.1 Molecular docking

Docking tasks for molecules, including small molecules, linear peptides, cyclic peptides, and those containing non-standard amino acids, share the same prompt definitions. There are typically two strategies for the molecular docking task (small molecular docking and peptide docking): 1) individual atom coordinates are allowed to be freely moved in the 3D space, and 2) only the rigid transformation (translation and rotation) and the rotation of rotatable bonds are allowed to be applied to the molecule. The two strategies use different noise distributions for atom coordinates, which are the Gaussian noise and flexible noise (see Supplementary Notes 3.4.1), respectively.

The task prompt for docking using Gaussian noise is defined as:

$$\begin{aligned} \mathbf{P}^{\mathrm{atom}} &= \mathbf{1}_N \\ \mathbf{P}^{\mathrm{coor}} &= \mathbf{0}_N \\ \mathbf{P}^{\mathrm{bond}} &= \mathbf{1}_{N\times N} \\ \mathbf{P}^{\mathrm{dist}} &= \mathbf{0}_{N\times N} \\ \mathbf{P}^{\mathrm{pep}} &= \mathbf{0}_N \end{aligned} \tag{S11}$$

The task prompt for docking using flexible noise is the same as that for docking using Gaussian noise except for the indicators $\mathbf{P}^{\mathrm{dist}}$. Unlike Gaussian noise where the atom distances are changed during the perturbation, the flexible noise will not change the distances of certain atom pairs. All pairwise distances are not changed by merely applying the random translation and rotation to a group of atoms. By applying rotations on rotatable bonds, atom pairs that are always on the same side of rotatable bonds also maintain their distances during the perturbation. Therefore, by setting the indicators of pairwise distances properly, the denoiser can predict the atom coordinates with constraints of some fixed pairwise distances, performing just like applying rigid transformation and rotation on rotatable bonds on the atom coordinates. The indicators of distances $\mathbf{P}^{\mathrm{dist}}$ is derived using Algorithm 17.

**Algorithm 17 GetDistPromptFlexNoise:** get the distance prompt indicators using flexible noise

**Input:** molecule $\mathcal{M} = \{\mathbf{A}, \mathbf{X}, \mathbf{B}\}$.

**Output:** prompt indicators for atom distance $\mathbf{P}^{\text{dist}}$.

$N \leftarrow$ the number of atoms of $\mathcal{M}$
$\mathbf{P}^{\text{dist}} = \mathbf{1}_{N \times N}$ ▷ initialize as all fixed atom distances
$\mathbb{A} = \{i | i \in [\![N]\!]\}$ ▷ atom index set
$\mathbb{B}^{\text{chem}}, \mathbb{B}^{\text{rot}} \leftarrow$ bond index sets of chemical bonds and rotatable bonds of $\mathcal{M}$
**for all** $ij \in \mathbb{B}^{\text{rot}}$ **do**
    **for all** $k, m \in \mathbb{A} \setminus \{i, j\}$ **do**
        **if** no path from $k$ to $m$ exists on the graph of $\mathbb{B}^{\text{chem}} \setminus \{ij\}$ **then**
            $P^{\text{dist}}_{km} \leftarrow 0$ ▷ rotation of bond $ij$ can change distance between atoms $k$ and $m$
        **end if**
    **end for**
**end for**

##### 2.2.2 Molecular conformation generation

The prompt for molecular conformation generation is the same as that for molecular docking (Eq. S11 and Algorithm 17).

##### 2.2.3 Structure-based drug design

In structure-based drug design (SBDD), all variables of the molecules need to be generated and thus the complete prompt is

$$\begin{aligned} \mathbf{P}^{\text{atom}} &= \mathbf{0}_N \\ \mathbf{P}^{\text{coor}} &= \mathbf{0}_N \\ \mathbf{P}^{\text{bond}} &= \mathbf{0}_{N \times N} \\ \mathbf{P}^{\text{dist}} &= \mathbf{0}_{N \times N} \\ \mathbf{P}^{\text{pep}} &= \mathbf{0}_N \end{aligned} \tag{S12}$$

##### 2.2.4 3D molecule generation

The prompt is the same as that for SBDD (Eq. S12).

##### 2.2.5 Molecular optimization

For molecular optimization, the prompt depends on which part of the molecule is allowed to be modified, and thus the related variables are set as unfixed. Typically, if all variables are allowed to be modified, which is the case in our experiment, the prompt for molecular optimization is the same as that of the SBDD task (Eq. S12).

##### 2.2.6 Fragment linking

In fragment linking, the linker is designed to connect the fragments to form a complete molecule, and thus the prompt related to the fragment and the linker is different. For a molecule with $N$ atoms, assume the atom set is divided into the fragment and the linker, whose atom index sets are

$\mathbb{A}_{\text{frag}}$ and $\mathbb{A}_{\text{link}}$, respectively. Typically, the complete prompt is

$$\begin{aligned}
&\forall i, P_i^{\text{atom}} = \begin{cases} 1, & i \in \mathbb{A}_{\text{frag}} \\ 0, & i \in \mathbb{A}_{\text{link}} \end{cases} \\
&\forall i, P_i^{\text{coor}} = \begin{cases} 1, & i \in \mathbb{A}_{\text{frag}} \\ 0, & i \in \mathbb{A}_{\text{link}} \end{cases} \\
&\forall ij, P_{ij}^{\text{bond}} = \begin{cases} 1, & i \in \mathbb{A}_{\text{frag}} \text{ and } j \in \mathbb{A}_{\text{frag}} \\ 0, & \text{otherwise} \end{cases} \\
&\forall ij, P_{ij}^{\text{dist}} = \begin{cases} 1, & i \in \mathbb{A}_{\text{frag}} \text{ and } j \in \mathbb{A}_{\text{frag}} \\ 0, & \text{otherwise} \end{cases} \\
&\mathbf{P}^{\text{pep}} = \mathbf{0}_N
\end{aligned} \tag{S13}$$

The flexibility of the task prompt facilitates the specification of more complicated settings in fragment linking. When we want to specify which fragment atoms can possibly connect to the linker, we can achieve it by setting the bond types between the non-connecting atoms of fragments and the linker atoms as fixed. Assume the non-connecting atoms of the fragments are $\mathbb{A}_{\text{non-connect}} \in \mathbb{A}_{\text{frag}}$, then the prompt of bond is modified as

$$\forall ij,\ P_{ij}^{\text{bond}} = \begin{cases} 1, & i \in \mathbb{A}_{\text{frag}} \text{ and } j \in \mathbb{A}_{\text{frag}} \\ 1, & i \in \mathbb{A}_{\text{non-connect}} \text{ or } j \in \mathbb{A}_{\text{non-connect}} \\ 0, & \text{otherwise} \end{cases} \tag{S14}$$

Furthermore, in some cases, the fragment poses are not fixed and need to be determined together with the generation of the linker. Thus, we can also set the coordinates related to the fragments as unfixed and allow the denoiser to adapt these coordinates:

$$\begin{aligned}
\mathbf{P}^{\text{coor}} &= \mathbf{0}_N \\
\mathbf{P}^{\text{dist}} &= \mathbf{0}_{N \times N}
\end{aligned} \tag{S15}$$

In the experiments of fragment linking, we assumed the fragment poses were fixed and used Eq. S13 and Eq. S14 to derive the prompt.

###### 2.2.7 PROTAC design

The prompt is the same as that of fragment linking. In the experiment of PROTAC design, we assumed the connecting atoms were known and considered the cases that the fragment poses were known or not (Eq. S13 - S15).

###### 2.2.8 Fragment growing

Fragment growing designs a complete molecule containing the given fragments. We can divide the molecule into two parts: the fragment part and the growing part. The prompt is similar to that of the fragment linking, except that the growing part corresponds to the linker in fragment linking. In our experiment of fragment growing, the fragment poses were assumed to be completely unknown. Specifically, assume the atom index sets of the fragment part and growing part are $\mathbb{A}_{\text{frag}}$ and $\mathbb{A}_{\text{grow}}$, respectively, the prompt is

$$\begin{aligned}
\forall i, P_i^{\text{atom}} &= \begin{cases} 1, & i \in \mathbb{A}_{\text{frag}} \\ 0, & i \in \mathbb{A}_{\text{grow}} \end{cases} \\
\mathbf{P}^{\text{coor}} &= \mathbf{0}_N \\
\forall ij, P_{ij}^{\text{bond}} &= \begin{cases} 1, & i \in \mathbb{A}_{\text{frag}} \text{ and } j \in \mathbb{A}_{\text{frag}} \\ 0, & \text{otherwise} \end{cases} \\
\mathbf{P}^{\text{dist}} &= \mathbf{0}_{N\times N} \\
\mathbf{P}^{\text{pep}} &= \mathbf{0}_N
\end{aligned} \tag{S16}$$

##### 2.2.9 Other fragment-based drug designs

Other fragment-based drug designs, such as scaffold hopping and R-group replacement, can also be generally regarded as partial molecule designs. Specifically, the molecule to be generated can be divided into two parts, such that the atoms and bonds in the first part are known and all variables in the second part are to be designed. More generally, the molecules can be divided into multiple parts and distinct prompt indicators are defined for different parts to define more precise generation specifications.

##### 2.2.10 De novo peptide design

This task can be regarded as a special case of fragment growing, where the side-chain atoms are generated based on the backbone fragment. The backbone fragment poses are unknown, and the possible connecting atoms of the backbone are the alpha carbon atoms or the nitrogen atoms. Additionally, there is no guarantee to generate side-chain atoms of standard amino acid types merely based on the backbone fragment. After all, the denoiser can generate a small molecule that contains the backbone fragment. Therefore, we use a group of auxiliary prompt indicators $\mathbf{P}^{\text{pep}}$ to inform the model to generate new atoms composing standard amino acids.

Assume the set of backbone atom indices is $\mathbb{A}_{\text{bb}} = \mathbb{A}_{\text{bb-connect}} \cup \mathbb{A}_{\text{bb-noncon}}$, where $\mathbb{A}_{\text{bb-connect}}$ is the set of alpha carbon atoms and nitrogen atoms of the backbone, and $\mathbb{A}_{\text{bb-noncon}}$ is the set of other backbone atoms (i.e., the carbon atoms and oxygen atoms). Assume the set of side-chain atom indices is $\mathbb{A}_{\text{sc}}$, the prompt is defined as:

$$\begin{aligned}
\forall i, P_i^{\text{atom}} &= \begin{cases} 1, & i \in \mathbb{A}_{\text{bb}} \\ 0, & i \in \mathbb{A}_{\text{sc}} \end{cases} \\
\mathbf{P}^{\text{coor}} &= \mathbf{0}_N \\
\forall ij, P_{ij}^{\text{bond}} &= \begin{cases} 1, & i \in \mathbb{A}_{\text{bb}} \text{ and } j \in \mathbb{A}_{\text{bb}} \\ 1, & i \in \mathbb{A}_{\text{bb-noncon}} \text{ or } j \in \mathbb{A}_{\text{bb-noncon}} \\ 0, & \text{otherwise} \end{cases} \\
\mathbf{P}^{\text{dist}} &= \mathbf{0}_{N\times N} \\
\mathbf{P}^{\text{pep}} &= \mathbf{1}_N
\end{aligned} \tag{S17}$$

The prompts for linear peptides, cyclic peptides, and those containing non-standard amino acids are the same.

##### 2.2.11 Peptide inverse folding

Peptide inverse folding is to design the side-chain atoms based on the fixed backbone structures. Thus, the prompt is similar to that of *de novo* peptide design except for the items related to the backbone's atom coordinates:

$$
\begin{aligned}
\forall i, P_i^{\text{atom}} &= \begin{cases} 1, & i \in \mathbb{A}_{\text{bb}} \\ 0, & i \in \mathbb{A}_{\text{sc}} \end{cases} \\
\forall i, P_i^{\text{coor}} &= \begin{cases} 1, & i \in \mathbb{A}_{\text{bb}} \\ 0, & i \in \mathbb{A}_{\text{sc}} \end{cases} \\
\forall ij, P_{ij}^{\text{bond}} &= \begin{cases} 1, & i \in \mathbb{A}_{\text{bb}} \text{ and } j \in \mathbb{A}_{\text{bb}} \\ 1, & i \in \mathbb{A}_{\text{bb-noncon}} \text{ or } j \in \mathbb{A}_{\text{bb-noncon}} \\ 0, & \text{otherwise} \end{cases} \\
\forall ij, P_{ij}^{\text{dist}} &= \begin{cases} 1, & i \in \mathbb{A}_{\text{bb}} \text{ and } j \in \mathbb{A}_{\text{bb}} \\ 0, & \text{otherwise} \end{cases} \\
\mathbf{P}^{\text{pep}} &= \mathbf{1}_N
\end{aligned}
\tag{S18}
$$

###### 2.2.12 Other peptide design tasks

The prompt can also express other specific peptide design tasks. For example, if we only want to design partial peptides based on the protein-peptide complex, we can set the corresponding prompt indicators related to the atoms to be modified as unfixed.

##### 2.3 More task prompts

Here, we show more task prompts of non-canonical, more customized, or complicated tasks.

**Partially docking using Gaussian noise.** This task assumes some parts of the molecules are fixed during docking, which can be useful when we know the coordinates of some parts of the molecules and only dock the rest. In this case, we can divide all atom indices into two sets: the fixed set $\mathbb{A}_{\text{fixed}}$ and the unfixed set $\mathbb{A}_{\text{unfixed}}$. The complete prompt is:

$$
\begin{aligned}
\mathbf{P}^{\text{atom}} &= \mathbf{1}_N \\
\forall i, P_i^{\text{coor}} &= \begin{cases} 1, & i \in \mathbb{A}_{\text{fixed}} \\ 0, & i \in \mathbb{A}_{\text{unfixed}} \end{cases} \\
\mathbf{P}^{\text{bond}} &= \mathbf{1}_{N \times N} \\
\forall ij, P_{ij}^{\text{dist}} &= \begin{cases} 1, & i \in \mathbb{A}_{\text{fixed}} \text{ and } j \in \mathbb{A}_{\text{fixed}} \\ 0, & \text{otherwise} \end{cases} \\
\mathbf{P}^{\text{pep}} &= \mathbf{0}_N
\end{aligned}
\tag{S19}
$$

**Rigid docking.** If we want the molecule to be completely rigid during docking, we can set all atom distances as fixed based on the prompt of the canonical docking. The prompt is:

$$
\begin{aligned}
\mathbf{P}^{\text{atom}} &= \mathbf{1}_N \\
\mathbf{P}^{\text{coor}} &= \mathbf{0}_N \\
\mathbf{P}^{\text{bond}} &= \mathbf{1}_{N \times N} \\
\mathbf{P}^{\text{dist}} &= \mathbf{1}_{N \times N} \\
\mathbf{P}^{\text{pep}} &= \mathbf{0}_N
\end{aligned}
\tag{S20}
$$

**Rotation of fragments** This task is to rotate some molecular fragments, such as some R-groups of small molecules or some side chains of peptides. The task can be viewed as a special case of canonical docking with flexible noise, where only predefined rotatable bonds are permitted to rotate. We divide the molecule into two parts: the fixed part and the unfixed part, denoting their atoms set as $\mathbb{A}_{\text{fixed}}$ and $\mathbb{A}_{\text{unfixed}}$, respectively. Then we have:

$$
\begin{aligned}
\mathbf{P}^{\text{atom}} &= \mathbf{1}_N \\
\forall i, P_i^{\text{coor}} &= \begin{cases} 1, & i \in \mathbb{A}_{\text{fixed}} \\ 0, & i \in \mathbb{A}_{\text{unfixed}} \end{cases} \\
\mathbf{P}^{\text{bond}} &= \mathbf{1}_{N \times N}
\end{aligned}
\tag{S21}
$$

The prompt indicators $\mathbf{P}^{\text{pep}}$ are equal to $\mathbf{0}_N$ for small molecule and $\mathbf{1}_N$ for peptide. The prompt indicators $\mathbf{P}^{\text{dist}}$ are defined in a similar way as Algorithm 17 but with specified rotatable bonds, as defined in Algorithm 18. Here, the specified rotatable bonds are the chemical bonds connecting the fixed atoms $\mathbb{A}_{\text{fixed}}$ and unfixed atoms $\mathbb{A}_{\text{unfixed}}$ of the molecule.

---

**Algorithm 18 GetDistPromptSpecifyRot:** get the distance prompt indicators with specified rotatable bonds

---

**Input:** molecule $\mathcal{M} = \{\mathbf{A}, \mathbf{X}, \mathbf{B}\}$, specified rotatable bonds $\mathbb{B}^{\text{rot}}$.

**Output:** prompt indicators for atom distance $\mathbf{P}^{\text{dist}}$.

$N \leftarrow$ the number of atoms of $\mathcal{M}$
$\mathbf{P}^{\text{dist}} = \mathbf{1}_{N \times N}$ ▷ initialize as all fixed atom distances
$\mathbb{A} = \{i | i \in [\![N]\!]\}$ ▷ atom index set
$\mathbb{B}^{\text{chem}} \leftarrow$ bond index sets of chemical bonds of $\mathcal{M}$
**for all** $ij \in \mathbb{B}^{\text{rot}}$ **do**
    **for all** $k, m \in \mathbb{A} \setminus \{i, j\}$ **do**
        **if** no path from $k$ to $m$ exists on the graph of $\mathbb{B}^{\text{chem}} \setminus \{ij\}$ **then**
            $P_{km}^{\text{dist}} \leftarrow 0$ ▷ rotation of bond $ij$ can change distance between atoms $k$ and $m$
        **end if**
    **end for**
**end for**

---

**In-place local fragment editing.** This task involves redesigning specific parts of a ligand molecule based on a known protein–ligand complex. Given an existing binding structure, the goal is to modify specific molecular fragments while keeping the remaining parts unchanged and preserving their original binding pose. We can divide the molecule into two parts: the fixed part and the unfixed part that is to be redesigned, denoted as $\mathbb{A}_{\text{fixed}}$ and $\mathbb{A}_{\text{unfixed}}$, respectively. The complete prompt is:

$$
\begin{aligned}
&\forall i, P_i^{\text{atom}} = \begin{cases} 1, & i \in \mathbb{A}_{\text{fixed}} \\ 0, & i \in \mathbb{A}_{\text{unfixed}} \end{cases} \\
&\forall i, P_i^{\text{coor}} = \begin{cases} 1, & i \in \mathbb{A}_{\text{fixed}} \\ 0, & i \in \mathbb{A}_{\text{unfixed}} \end{cases} \\
&\forall ij, P_{ij}^{\text{bond}} = \begin{cases} 1, & i \in \mathbb{A}_{\text{fixed}} \text{ and } j \in \mathbb{A}_{\text{fixed}} \\ 0, & \text{otherwise} \end{cases} \\
&\forall ij, P_{ij}^{\text{dist}} = \begin{cases} 1, & i \in \mathbb{A}_{\text{fixed}} \text{ and } j \in \mathbb{A}_{\text{fixed}} \\ 0, & \text{otherwise} \end{cases} \\
&\mathbf{P}^{\text{pep}} = \mathbf{0}_N
\end{aligned}
\tag{S22}
$$

We can also specify the atoms in the fixed part that can connect to the new fragments. Assume the connecting and non-connecting atoms of the fixed part are $\mathbb{A}_{\text{connect}}$ and $\mathbb{A}_{\text{non-connect}}$, respectively, and they have $\mathbb{A}_{\text{connect}} \cup \mathbb{A}_{\text{non-connect}} = \mathbb{A}_{\text{fixed}}$, then we only need to modify $\mathbb{P}^{\text{bond}}$ as:

$$
\forall ij, P_{ij}^{\text{bond}} = \begin{cases} 1, & i \in \mathbb{A}_{\text{fixed}} \text{ and } j \in \mathbb{A}_{\text{fixed}} \\ 1, & i \in \mathbb{A}_{\text{non-connect}} \text{ or } j \in \mathbb{A}_{\text{non-connect}} \\ 0, & \text{otherwise} \end{cases}
\tag{S23}
$$

**Pose-agnostic local fragment editing** This task focuses on redesigning specific regions of a ligand molecule without access to the protein–ligand complex structure. A straightforward approach would be to first dock the ligand into the binding pocket and then perform in-place local fragment editing described above. However, PocketXMol is capable of performing both steps simultaneously in a unified generative process. Similarly, we divide the molecule into two parts: the fixed part, whose molecular graph (atom types and bond types) remains unchanged, and the unfixed region, which is subject to redesign. These are denoted as 𝔸fixed and 𝔸unfixed, respectively. The complete prompt is:

$$
\begin{aligned}
&\forall i, P_i^{\text{atom}} = \begin{cases} 1, & i \in \mathbb{A}_{\text{fixed}} \\ 0, & i \in \mathbb{A}_{\text{unfixed}} \end{cases} \\
&\mathbf{P}^{\text{coor}} = \mathbf{0}_N \\
&\forall ij, P_{ij}^{\text{bond}} = \begin{cases} 1, & i \in \mathbb{A}_{\text{fixed}} \text{ and } j \in \mathbb{A}_{\text{fixed}} \\ 0, & \text{otherwise} \end{cases} \\
&\mathbf{P}^{\text{dist}} = \mathbf{0}_{N\times N} \\
&\mathbf{P}^{\text{pep}} = \mathbf{0}_N
\end{aligned}
\tag{S24}
$$

**General local fragment editing** This task represents a more general form of the two aforementioned cases of local fragment editing. In this setting, we assume that some atoms are fixed in both the graph and 3D pose, some are fixed in the graph but have unfixed positions, and the remaining atoms are subject to redesign. We divide the molecule into three parts: the completed fixed part, the graph-fixed part, and the redesigned part. Their atom indices are denoted as $\mathbb{A}_{\text{fixed}}$, $\mathbb{A}_{\text{dock}}$, and

$\mathbb{A}_{\text{design}}$, respectively. The complete prompt is

$$\begin{aligned}
\forall i, P_i^{\text{atom}} &= \begin{cases} 1, & i \in \mathbb{A}_{\text{fixed}} \\ 1, & i \in \mathbb{A}_{\text{dock}} \\ 0, & i \in \mathbb{A}_{\text{design}} \end{cases} \\
\forall i, P_i^{\text{coor}} &= \begin{cases} 1, & i \in \mathbb{A}_{\text{fixed}} \\ 0, & i \in \mathbb{A}_{\text{dock}} \\ 0, & i \in \mathbb{A}_{\text{design}} \end{cases} \\
\forall ij, P_{ij}^{\text{bond}} &= \begin{cases} 1, & i \notin \mathbb{A}_{\text{design}} \text{ and } j \notin \mathbb{A}_{\text{design}} \\ 0, & \text{otherwise} \end{cases} \\
\forall ij, P_{ij}^{\text{dist}} &= \begin{cases} 1, & i \in \mathbb{A}_{\text{fixed}} \text{ and } j \in \mathbb{A}_{\text{fixed}} \\ 0, & \text{otherwise} \end{cases} \\
\mathbf{P}^{\text{pep}} &= \mathbf{0}_N
\end{aligned} \tag{S25}$$

By setting $\mathbb{A}_{\text{dock}} = \emptyset$ and $\mathbb{A}_{\text{design}} = \mathbb{A}_{\text{unfixed}}$, we can derive the task prompt of in-place local fragment editing (Eq. S22). By setting $\mathbb{A}_{\text{fixed}} = \emptyset$, $\mathbb{A}_{\text{design}} = \mathbb{A}_{\text{unfixed}}$, we can derive the task prompt of pose-agnostic local fragment editing (Eq. S24). More aggressively, by setting $\mathbb{A}_{\text{fixed}} = \mathbb{A}_{\text{dock}} = \emptyset$, the task prompt becomes the same as that of the canonical molecule design or SBDD tasks. By setting $\mathbb{A}_{\text{fixed}} = \mathbb{A}_{\text{design}} = \emptyset$, the task prompt becomes the same as that of the canonical docking tasks.

**Peptide design with specified residues.** This task involves designing peptides that contain specified residues and may include various specific constraints. Their task prompts can be modified from that of the canonical peptide design (Eq S17), and we will also follow the notations in Eq S17. Below, we elaborate on these settings and how their prompt are modified.

(1) Known residue types and indices. The designed peptides must contain residues with specified amino acid types at designated residue indices. For example, one may require the peptide to have a cysteine as the 4th residue. Another example is generating peptides with a partially fixed sequence such as XXVLIMHDQXX, where the "X" positions are to be generated. In this case, we divide all side chains $\mathbb{A}_{\text{sc}}$ into two parts: the side chains of specified residues and the side chains of generated residues, denoted as $\mathbb{A}_{\text{sc-known}}$ and $\mathbb{A}_{\text{sc-unknown}}$. This setting can be interpreted as incorporating the side chains of specified residues, $\mathbb{A}_{\text{sc-known}}$, into the backbone set $\mathbb{A}_{\text{bb}}$ used in canonical peptide design (Eq. S17). The complete prompt:

$$\begin{aligned}
\forall i, P_i^{\text{atom}} &= \begin{cases} 1, & i \in \mathbb{A}_{\text{bb}} \cup \mathbb{A}_{\text{sc-known}} \\ 0, & i \in \mathbb{A}_{\text{sc-unknown}} \end{cases} \\
\mathbf{P}^{\text{coor}} &= \mathbf{0}_N \\
\forall ij, P_{ij}^{\text{bond}} &= \begin{cases} 1, & i \in \mathbb{A}_{\text{bb}} \cup \mathbb{A}_{\text{sc-known}} \text{ and } j \in \mathbb{A}_{\text{bb}} \cup \mathbb{A}_{\text{sc-known}} \\ 1, & i \in \mathbb{A}_{\text{bb-noncon}} \text{ or } j \in \mathbb{A}_{\text{bb-noncon}} \\ 0, & \text{otherwise} \end{cases} \\
\mathbf{P}^{\text{dist}} &= \mathbf{0}_{N \times N} \\
\mathbf{P}^{\text{pep}} &= \mathbf{1}_N
\end{aligned} \tag{S26}$$

(2) Known residue types, indices, and coordinates. The setting additionally knows the coordinates of the specified residues based on the setting of the known residue types and indices. We assume the backbones and side chains of the specified residues are $\mathbb{A}_{\text{bb-known}}$ and $\mathbb{A}_{\text{sc-known}}$, and the

backbones and side chains of generated residues are $\mathbb{A}_{\text{bb-unknown}}$ and $\mathbb{A}_{\text{sc-unknown}}$. We further assume $\mathbb{A}_{\text{bb-noncon}}$ are the connecting atoms of $\mathbb{A}_{\text{bb-unknown}}$. We denote $\mathbb{A}_{\text{bb}}$ as $\mathbb{A}_{\text{bb-known}} \cup \mathbb{A}_{\text{bb-unknown}}$. The prompt is defined as:

$$
\begin{aligned}
\forall i, P_i^{\text{atom}} &= \begin{cases} 1, & i \in \mathbb{A}_{\text{bb}} \cup \mathbb{A}_{\text{sc-known}} \\ 0, & i \in \mathbb{A}_{\text{sc-unknown}} \end{cases} \\
\forall i, P_i^{\text{coor}} &= \begin{cases} 1, & i \in \mathbb{A}_{\text{bb-known}} \cup \mathbb{A}_{\text{sc-known}} \\ 0, & i \in \mathbb{A}_{\text{bb-unknown}} \cup \mathbb{A}_{\text{sc-unknown}} \end{cases} \\
\forall ij, P_{ij}^{\text{bond}} &= \begin{cases} 1, & i \in \mathbb{A}_{\text{bb}} \cup \mathbb{A}_{\text{sc-known}} \text{ and } j \in \mathbb{A}_{\text{bb}} \cup \mathbb{A}_{\text{sc-known}} \\ 1, & i \in \mathbb{A}_{\text{bb-noncon}} \text{ or } j \in \mathbb{A}_{\text{bb-noncon}} \\ 0, & \text{otherwise} \end{cases} \\
\forall ij, P_{ij}^{\text{dist}} &= \begin{cases} 1, & i \in \mathbb{A}_{\text{bb-known}} \cup \mathbb{A}_{\text{sc-known}} \text{ and } j \in \mathbb{A}_{\text{bb-known}} \cup \mathbb{A}_{\text{sc-known}} \\ 0, & \text{otherwise} \end{cases} \\
\mathbf{P}^{\text{pep}} &= \mathbf{1}_N
\end{aligned} \tag{S27}
$$

The difference of the task prompt from the previous setting (Eq. S26) is additionally setting the atom coordinates and distances of the known residues (both backbones and side chains) as fixed.

(3) Known residue type. The designed peptides should contain a certain residue type, but the residue index is not specified. We divide the atoms of the peptide into the backbone $\mathbb{A}_{\text{bb}}$ and side chains $\mathbb{A}_{\text{sc}}$. We further divide the side chains $\mathbb{A}_{\text{sc}}$ into side chains of the known residue type and unknown residues, denoted as $\mathbb{A}_{\text{sc-known}}$ and $\mathbb{A}_{\text{sc-unknown}}$, respectively. We denote the non-connecting atoms of $\mathbb{A}_{\text{sc-known}}$ as $\mathbb{A}_{\text{sc-known-noncon}}$ and the non-connecting atoms of $\mathbb{A}_{\text{bb}}$ as $\mathbb{A}_{\text{bb-noncon}}$. Then the task prompt is:

$$
\begin{aligned}
\forall i, P_i^{\text{atom}} &= \begin{cases} 1, & i \in \mathbb{A}_{\text{bb}} \cup \mathbb{A}_{\text{sc-known}} \\ 0, & i \in \mathbb{A}_{\text{sc-unknown}} \end{cases} \\
\mathbf{P}^{\text{coor}} &= \mathbf{0}_N \\
\forall ij, P_{ij}^{\text{bond}} &= \begin{cases} 1, & i \in \mathbb{A}_{\text{bb}} \text{ and } j \in \mathbb{A}_{\text{bb}} \\ 1, & i \in \mathbb{A}_{\text{sc-known}} \text{ and } j \in \mathbb{A}_{\text{sc-known}} \\ 1, & i \in \mathbb{A}_{\text{bb-noncon}} \text{ or } j \in \mathbb{A}_{\text{bb-noncon}} \\ 1, & i \in \mathbb{A}_{\text{sc-knwon-noncon}} \text{ or } j \in \mathbb{A}_{\text{sc-knwon-noncon}} \\ 0, & \text{otherwise} \end{cases} \\
\mathbf{P}^{\text{dist}} &= \mathbf{0}_{N \times N} \\
\mathbf{P}^{\text{pep}} &= \mathbf{1}_N
\end{aligned} \tag{S28}
$$

Actually, compared to the setting of known residue types and indices (Eq. S26), this setting further sets the bond types between the connecting atoms of the known side chain and the connecting atoms of the backbones as unfixed, indicating that the attachment points of the side chains of the known residue to the backbone are not constrained.

(4) Known residue type and coordinates. The setting constrains specific a residue type with known coordinates of its side-chain atoms, but does not specify its residue index. Using the same

notations as Eq. S28, the prompt of this task is:

$$
\begin{aligned}
\forall i, P_i^{\text{atom}} &= \begin{cases} 1, & i \in \mathbb{A}_{\text{bb}} \cup \mathbb{A}_{\text{sc-known}} \\ 0, & i \in \mathbb{A}_{\text{sc-unknown}} \end{cases} \\
\forall i, P_i^{\text{coor}} &= \begin{cases} 1, & i \in \mathbb{A}_{\text{sc-known}} \\ 0, & i \in \mathbb{A}_{\text{bb}} \cup \mathbb{A}_{\text{sc-unknown}} \end{cases} \\
\forall ij, P_{ij}^{\text{bond}} &= \begin{cases} 1, & i \in \mathbb{A}_{\text{bb}} \text{ and } j \in \mathbb{A}_{\text{bb}} \\ 1, & i \in \mathbb{A}_{\text{sc-known}} \text{ and } j \in \mathbb{A}_{\text{sc-known}} \\ 1, & i \in \mathbb{A}_{\text{bb-noncon}} \text{ or } j \in \mathbb{A}_{\text{bb-noncon}} \\ 1, & i \in \mathbb{A}_{\text{sc-knwon-noncon}} \text{ or } j \in \mathbb{A}_{\text{sc-knwon-noncon}} \\ 0, & \text{otherwise} \end{cases} \\
\forall ij, P_{ij}^{\text{dist}} &= \begin{cases} 1, & i \in \mathbb{A}_{\text{sc-known}} \text{ and } j \in \mathbb{A}_{\text{sc-known}} \\ 0, & \text{otherwise} \end{cases} \\
\mathbf{P}^{\text{pep}} &= \mathbf{1}_N
\end{aligned}
\tag{S29}
$$

Compared to the setting of known residue type (Eq. S28), this setting further sets the atom coordinates and distances of the side chains of the known residue type as fixed.

### 3 Task noise and perturbation

#### 3.1 Basic noise distributions

The task noise for any task comprises the following four types of basic noise components. Assume the noise scale is $\beta \in [0, 1]$ used to control how much noise is introduced to the variables. The noise components are defined as:

1. **Gaussian distribution.** Gaussian noise is used to perturb individual atom coordinates or randomly translate a set of coordinates in 3D space. For atom coordinate $\mathbf{x}_i$, the noisy coordinate $\tilde{\mathbf{x}}_i$ is sampled using the variance-exploding noise form[1]:

   $$p(\tilde{\mathbf{x}}_i|\mathbf{x}, \sigma, \beta) = \mathcal{N}(\mathbf{x}, \beta\sigma^2\mathbf{I}) \tag{S30}$$

   where $\sigma$ and $\mathbf{I} \in \mathbb{R}^{3\times 3}$ are the standard deviation and the identity matrix, respectively. When the noise is used for random translation of a group of $N'$ atoms, we add Gaussian noise to the center of the atoms $\mathbf{x}^{\text{center}} = 1/N' \sum_i \mathbf{x}_i$ using the variance-preserving noise form[1]

   $$p(\tilde{\mathbf{x}}^{\text{center}}|\sigma, \beta) = \mathcal{N}(\sqrt{1-\beta}\mathbf{x}^{\text{center}}, \beta\sigma^2\mathbf{I}) \tag{S31}$$

   The translation is derived as $\mathbf{t} = \tilde{\mathbf{x}}^{\text{center}} - \mathbf{x}^{\text{center}}$, which is applied to the coordinates of all corresponding atoms $i$ as $\tilde{\mathbf{x}}_i = \mathbf{x}_i + \mathbf{t}$.

2. **Categorical distribution.** The discrete variables including atom types and the bond types are perturbed using the categorical distributions. Let $\mathbf{a}_i$ and $\mathbf{b}_{ij}$ be the one-hot encoding vectors of atom type and bond type, respectively, the noisy types $\tilde{\mathbf{a}}_i$ and $\tilde{\mathbf{b}}_{ij}$ are sampled from:

   $$
   \begin{aligned}
   p(\tilde{\mathbf{a}}_i|\mathbf{a}_i, \bar{\mathbf{a}}, \beta) &= \text{Cat}\left((1-\beta)\mathbf{a}_i + \beta\bar{\mathbf{a}}\right) \\
   p(\tilde{\mathbf{b}}_{ij}|\mathbf{b}_{ij}, \bar{\mathbf{b}}, \beta) &= \text{Cat}\left((1-\beta)\mathbf{b}_{ij} + \beta\bar{\mathbf{b}}\right)
   \end{aligned}
   \tag{S32}
   $$

   where $\bar{\mathbf{a}}$ and $\bar{\mathbf{b}}$ are the prior distributions for atom and bond types, respectively.

3. **Isotropic Gaussian distribution in SO(3).** The random rotation can be applied to a set of atom coordinates by multiplying the coordinates with a rotation matrix $\mathbf{R}$ that is sampled from the Isotropic Gaussian distribution in SO(3)[2,3]:

$$p(\mathbf{R}|\epsilon, \beta) = \mathcal{IG}_{SO(3)}(0, \beta\epsilon^2) \tag{S33}$$

where $\epsilon$ is the standard deviation. Then the noisy coordinates of a set of $N'$ atoms $\mathbf{X} \in \mathbb{R}^{N' \times 3}$ are produced as $\tilde{\mathbf{X}}^\top = \mathbf{R}\mathbf{X}^\top$.

4. **Circular normal distribution.** The circular normal distribution, also known as the von Mises distribution, is a close approximation to the circular analog of the normal distribution[4] and is used to perturb the torsion angles of rotatable bonds as an indirect perturbation to atom coordinates. A random rotation $\Delta\omega$ is sampled from the distribution $\mathcal{N}_{\text{circ}}$ with zero mean as:

$$p(\Delta\omega|\kappa, \beta) = \mathcal{N}_{\text{circ}}(0, \beta/\kappa), \tag{S34}$$

where $\kappa$ is the concentration parameter (analogous to the reciprocal of the variance). The probability density function of the circular distribution with concentration parameter $\kappa$ is

$$p(\Delta\omega|\kappa) = \frac{\exp(\kappa\cos(\Delta\omega))}{2\pi I_0(\kappa)} \tag{S35}$$

where $I_0(\kappa)$ is the modified Bessel function of the first kind of order 0 to scale the probability density. The noisy coordinates are obtained by rotating the corresponding bonds by $\Delta\omega$ angles.

Having defined the basic types of noise distributions, we now define the task-specific molecular noise $\xi$ which specifies the noise distributions for individual molecular variables. If the different atoms or bonds of the entire molecule use consistent distributions, the molecular noise is defined as a set of noise parameters:

$$\xi = \{\bar{\mathbf{a}}, \bar{\mathbf{b}}, \sigma^{\text{Gauss}}, \sigma^{\text{trans}}, \epsilon, \kappa\} \tag{S36}$$

where $\bar{\mathbf{a}}, \bar{\mathbf{b}}, \sigma^{\text{Gauss}}, \sigma^{\text{trans}}, \epsilon$ and $\kappa$ are the parameters for the categorical distribution of atom types, categorical distribution of bond types, the Gaussian distribution of individual coordinates, the Gaussian distribution of translation vector, the $\mathcal{IG}_{\text{SO3}}$ distributions of a rotation matrix, and the circular normal distribution of torsion angles, respectively. Note that Eq. S36 is merely a general form and does not indicate that all these distributions should be defined. For example, in the docking tasks, there is no need to define $\bar{\mathbf{a}}_k$ and $\bar{\mathbf{b}}_k$ because the atom and bond types do not need to be generated.

In more general cases, different atoms or bonds use different noise distribution types or distribution parameters. We thus decompose the molecule $\mathcal{M}$ into $k$ parts $\{\mathbb{M}_k | k \in [\![n_{\text{part}}]\!]\}$. Each part $\mathbb{M}_k$ is a union of the atom index set $\mathbb{A}_k$ and bond (atom pair) index set $\mathbb{B}_k$ of the part, i.e., $\mathbb{M}_k = \mathbb{A}_k \cup \mathbb{B}_k$ Then the molecular noise is defined as a mapping $\xi(\mathbb{M}_k)$ from individual parts of the molecule to a set of noise distribution parameters applied for the part. A general form of $\xi(\mathbb{M}_k)$ is:

$$\xi(\mathbb{M}_k) = \{\bar{\mathbf{a}}_k, \bar{\mathbf{b}}_k, \sigma_k^{\text{Gauss}}, \sigma_k^{\text{trans}}, \epsilon_k, \kappa_k\},\ k \in [\![n_{\text{parts}}]\!] \tag{S37}$$

with the domain of $\xi$ as $\{\mathbb{M}_k | k \in [\![n_{\text{part}}]\!]\}$.

#### 3.2 Basic perturbation functions

Based on the distributions above, we define four basic perturbation functions for individual molecular variables. Typically, a perturbation function takes the corresponding molecular variable, the parameters of the distributions, and the noise scale $\beta \in [0, 1]$ as input and adds noise to the variables. There are four basic perturbation functions $\Phi^{\text{atom}}, \Phi^{\text{bond}}, \Phi^{\text{Gauss}}$ and $\Phi^{\text{flex}}$, which correspond to the perturbation of atom types, the perturbation of bond types, the perturbation of atom coordinates with individual Gaussian noise, and the perturbation of a group of atom coordinates using flexible noise (random rigid transformation and rotation of rotatable bonds), respectively.

The perturbations for atom type and bond types are defined as $\Phi^{\text{atom}}(\mathbf{a}_i, \bar{\mathbf{a}}, \beta)$ and $\Phi^{\text{bond}}(\mathbf{b}_{ij}, \bar{\mathbf{b}}, \beta)$, respectively, which involve directly sampling from the corresponding categorical distributions:

$$\begin{aligned} \Phi^{\text{atom}}(\mathbf{a}_i, \bar{\mathbf{a}}, \beta) &: \tilde{\mathbf{a}}_i \sim \text{Cat}\left((1-\beta)\mathbf{a}_i + \beta\bar{\mathbf{a}}\right) \\ \Phi^{\text{bond}}(\mathbf{b}_{ij}, \bar{\mathbf{b}}, \beta) &: \tilde{\mathbf{b}}_{ij} \sim \text{Cat}\left((1-\beta)\mathbf{b}_{ij} + \beta\bar{\mathbf{b}}\right) \end{aligned} \tag{S38}$$

where the $\bar{\mathbf{a}}$ and $\bar{\mathbf{b}}$ are the prior distributions. These perturbation functions are used in the generation of atom types and bond types. The perturbation on atom coordinates with Gaussian noise is defined as:

$$\Phi^{\text{Gauss}}(\mathbf{x}_i, \sigma, \beta) : \tilde{\mathbf{x}} \sim \mathcal{N}(\mathbf{x}, \beta\sigma^2\mathbf{I}) \tag{S39}$$

The perturbation function is used for the generation of atom coordinates when designing molecules or the generation of coordinates without the restrictions of atom distances.

Another type of perturbation on coordinates, named flexible noise, applies random rigid transformation on a set of atoms (can be the whole molecule or molecular fragment) and rotations on the rotatable bonds. Generally, assume the atom types and the bond types of a group of $N$ atoms are $\mathbf{A}$ and $\mathbf{B}$, respectively. The fragment atom coordinates before and after perturbation are $\tilde{\mathbf{X}} \in \mathbb{R}^{N\times 3}$ and $\mathbf{X} \in \mathbb{R}^{N\times 3}$, respectively. The perturbation function is denoted as:

$$\tilde{\mathbf{X}} = \Phi^{\text{flex}}\left(\mathbf{X}, \{\mathbf{A}, \mathbf{B}\}, \sigma, \epsilon, \kappa, \beta\right) \tag{S40}$$

where $\sigma$, $\epsilon$, and $\kappa$ are the parameters of the distributions. The perturbation function is defined in Algorithm 19, which applies a random rigid transformation to coordinates and subsequently applies random rotations of rotatable bonds.

**Algorithm 19 PertCoordinateFlex:** $\Phi^{\text{flex}}$, perturb coordinates with flexible noise

**Input:** atom coordinates $\mathbf{X}$, atom types and bond types $\{\mathbf{A}, \mathbf{B}\}$, standard deviation $\sigma$ of Gaussian distribution for translation vectors, standard deviation $\epsilon$ of $\mathcal{IG}_{SO(3)}$ for rotation matrices, the concentration parameter $\kappa$ of $\mathcal{N}_{\text{circ}}$ for torsion angles, noise scale $\beta$.

**Output:** perturbed fragment coordinates $\tilde{\mathbf{X}}$.

$N \leftarrow$ number of atoms of $\mathbf{X}$
$\tilde{\mathbf{X}} \leftarrow \mathbf{X}$
$\mathbf{x}^{\text{center}} = 1/N \sum_i \mathbf{x}_i$
$\tilde{\mathbf{x}}^{\text{center}} \sim \mathcal{N}(\sqrt{1-\beta}\mathbf{x}^{\text{center}}, \beta\sigma^2\mathbf{I})$
$\mathbf{t} = \tilde{\mathbf{x}}^{\text{center}} - \mathbf{x}^{\text{center}}$ ▷ Sample a translation vector
$\mathbf{R} = \mathcal{IG}_{SO(3)}(0, \beta\epsilon^2)$ ▷ Sample a rotation matrix
**for all** $\tilde{\mathbf{x}}_i \in \tilde{\mathbf{X}}$ **do**
    $\tilde{\mathbf{x}}_i \leftarrow \mathbf{R}\tilde{\mathbf{x}}_i + \mathbf{t}$ ▷ Apply rigid transformation
**end for**
$\mathbb{B}^{\text{chem}}, \mathbb{B}^{\text{rot}} \leftarrow$ bond index sets of chemical bonds and rotatable bonds of the molecule $\{\mathbf{A}, \mathbf{B}\}$
**for all** $ij \in \mathbb{B}^{\text{rot}}$ **do**
    $\Delta\omega_{ij} = \mathcal{N}_{\text{circ}}(0, \beta/\kappa)$ ▷ Sample torsion angle changes for all rotatable bonds
**end for**
$\sin\Delta\boldsymbol{\omega} = \{\sin\Delta\omega_{ij} | ij \in \mathbb{B}^{\text{rot}}\}$
$\cos\Delta\boldsymbol{\omega} = \{\cos\Delta\omega_{ij} | ij \in \mathbb{B}^{\text{rot}}\}$
$\tilde{\mathbf{X}} \leftarrow$ OrthogonalRotBonds($\mathbb{B}^{\text{chem}}, \mathbb{B}^{\text{rot}}, \tilde{\mathbf{X}}, \{\sin\Delta\boldsymbol{\omega}, \cos\Delta\boldsymbol{\omega}\}$)

**Algorithm 20 OrthogonalRotBonds:** rotate rotatable bonds orthogonally to rigid transformation.

**Input:** chemical bond set $\mathbb{B}^{\text{chem}}$, rotatable bond set $\mathbb{B}^{\text{rot}}$, atom coordinates $\mathbf{X}$, torsion angles changes $\{\sin\Delta\boldsymbol{\omega}, \cos\Delta\boldsymbol{\omega}\}$.

**Output:** updated coordinates $\mathbf{X}'$ by rotating rotatable bonds orthogonally to rigid transformation.

$\mathbf{X}' =$ RotBonds($\mathbb{B}^{\text{chem}}, \mathbb{B}^{\text{rot}}, \mathbf{X}, \{\sin\Delta\boldsymbol{\omega}, \cos\Delta\boldsymbol{\omega}\}$) ▷ rotate bonds in arbitrary order
$\mathbf{R}, \mathbf{t} =$ AlignStructure($\mathbf{X}', \mathbf{X}$) ▷ orthogonal to rigid transformation
**for all** $\mathbf{x}'_i \in \mathbf{X}'$ **do**
    $\mathbf{x}'_i \leftarrow \mathbf{R}\mathbf{x}'_i + \mathbf{t}$
**end for**

##### 3.3 General perturbation step

Having defined the molecular noise distributions and perturbation functions, we can derive the perturbation process on the whole molecule based on the task prompt and the parameter of the noise distributions.

Given the molecule $\mathcal{M}$, the molecular noise $\xi$, the task prompt $\mathcal{P}$ and the noise scale $\beta$, the general perturbation step $\Phi\big(\mathcal{M}, \mathcal{P}, \xi, \beta\big)$ (Eq. 1 in the main text) is defined in Algorithm 21, which applies the perturbation functions defined in Eq. S38-S40 to the molecular variables according to the task prompt and molecular noise.

**Algorithm 21 GeneralPert:** general perturbation step

**Input:** the molecule $\mathcal{M} = \{\mathbf{A}, \mathbf{X}, \mathbf{B}\}$, task prompt $\mathcal{P}$, molecular noise $\xi$, noise scale $\beta$.

**Output:** noisy molecule $\tilde{M} = \{\tilde{\mathbf{A}}, \tilde{\mathbf{X}}, \tilde{\mathbf{B}}\}$.

$\mathbf{P}^{\text{atom}}, \mathbf{P}^{\text{coor}}, \mathbf{P}^{\text{bond}}, \mathbf{P}^{\text{dist}}, \mathbf{P}^{\text{pep}} \leftarrow \mathcal{P}$
molecular decomposition $\big\{\mathbb{M}_k \big| k \in [\![n_{\text{part}}]\!]\big\} \leftarrow$ the domain of $\xi$
▷ Initialize
$\tilde{\mathbf{A}}, \tilde{\mathbf{X}}, \tilde{\mathbf{B}} \leftarrow \mathbf{A}, \mathbf{X}, \mathbf{B}$
**for all** $k \in [\![n_{\text{parts}}]\!]$ **do**
    $\mathbb{A}_k, \mathbb{B}_k \leftarrow \mathbb{M}_k$ ▷ atom index set and bond index set of the part
    $\{\bar{\mathbf{a}}_k, \bar{\mathbf{b}}_k, \sigma_k^{\text{Gauss}}, \sigma_k^{\text{trans}}, \epsilon_k, \kappa_k\} \leftarrow \xi(\mathbb{M}_k)$ ▷ noise distributions for this part
    $\beta_k \leftarrow \beta(\mathbb{M}_k)$ ▷ noise scale can also be part-specific
    **for all** $i \in \mathbb{A}_k$ **do** ▷ perturb atom types
        **if** $P_i^{\text{atom}} = 0$ **then**
            $\tilde{\mathbf{a}}_i = \Phi^{\text{atom}}(\mathbf{a}_i, \bar{\mathbf{a}}_k, \beta_k)$
        **end if**
    **end for**
    **for all** $ij \in \mathbb{B}_k$ **do** ▷ perturb bond types
        **if** $P_{ij}^{\text{bond}} = 0$ **then**
            $\tilde{\mathbf{b}}_{ij} = \Phi^{\text{bond}}(\mathbf{b}_{ij}, \bar{\mathbf{b}}_k, \beta_k)$
        **end if**
    **end for**
    **if** $\exists i \in \mathbb{A}_k, P_i^{\text{coor}} = 0$ **then** ▷ perturb atom coordinates
        **if** $\forall ij \in \mathbb{B}_k, P_{ij}^{\text{dist}} = 0$ **then** ▷ perturb using Gaussian noise
            **for all** $i \in \mathbb{A}_k$ **do**
                **if** $P_i^{\text{coor}} = 0$ **then**
                    $\tilde{\mathbf{x}}_i = \Phi^{\text{Gauss}}(\mathbf{x}_i, \sigma_k^{\text{Gauss}}, \beta_k)$
                **end if**
            **end for**
        **else** ▷ perturb using flexible noise
            $\tilde{\mathbf{X}}_{\mathbb{A}_k} = \Phi^{\text{flex}}\left(\mathbf{X}_{\mathbb{A}_k}, \{\mathbf{A}_{\mathbb{A}_k}, \mathbf{B}_{\mathbb{B}_k}\}, \sigma_k^{\text{trans}}, \epsilon_k, \kappa_k, \beta_k\right)$
        **end if**
    **end if**
**end for**

##### 3.4 Specifications for individual tasks

Here, we introduce the perturbation step for each generative task, which is derived based on the general perturbation process (Algorithm 21), with the specification of the molecular noise. For a molecule $\mathcal{M}$ with $N$ atoms, we assume the atom types, the atom coordinates, and the bond types of the molecule are denoted as $\mathbf{A}, \mathbf{X}$ and $\mathbf{B}$, respectively. The atom set and the bond set of the molecule $\mathbb{A}$ and $\mathbb{B}$, respectively. The molecular noise and the perturbation steps for individual tasks are defined in the following.

###### *3.4.1 Molecular docking*

Docking tasks for molecules, including small molecules, linear peptides, cyclic peptides, and peptides containing non-standard amino acids, utilize the same noise definitions. Molecular docking can use either Gaussian noise or flexible noise. For the Gaussian noise, we use one parameter $\sigma$ of the Gaussian distribution for the whole molecule:

$$\xi^{\text{Gaussian}} = \{\sigma\} \tag{S41}$$

where we set $\sigma = \sqrt{N}$ in our implementation and $N$ is the number of atoms. The perturbation step is defined in Algorithm 22.

**Algorithm 22 PertDockSmallMolGauss:** perturbation for molecular docking using Gaussian noise.

**Input:** molecule $\mathcal{M} = \{\mathbf{A}, \mathbf{X}, \mathbf{B}\}$, molecular noise $\xi$, noise scale $\beta$.

**Output:** noisy molecule $\tilde{\mathcal{M}} = \{\tilde{\mathbf{A}}, \tilde{\mathbf{X}}, \tilde{\mathbf{B}}\}$.

1: $\{\sigma\} \leftarrow \xi$

2: $\tilde{\mathbf{A}}, \tilde{\mathbf{X}}, \tilde{\mathbf{B}} \leftarrow \mathbf{A}, \mathbf{X}, \mathbf{B}$

3: **for all** $i \in \{1, 2, \ldots N\}$ **do**

4: $\quad \tilde{\mathbf{x}}_i = \Phi^{\text{Gauss}}(\mathbf{x}_i, \sigma, \beta)$

5: **end for**

For molecular docking using flexible noise, we use one set of parameters $\{\sigma, \epsilon, \kappa\}$ for the entire molecule:

$$\xi^{\text{flex}} = \{\sigma, \epsilon, \kappa\} \tag{S42}$$

where $\sigma$, $\epsilon$, and $\kappa$ are the parameters for the Gaussian distribution of the translation vector, $\mathcal{IG}_{SO(3)}$ of the rotation matrix, and the circular normal distribution of torsion angles and are set as 1, 0.0002, and 0.2, respectively in our implementation. The perturbation step is defined in Algorithm 23.

**Algorithm 23 PertDockSmallMolFlex:** perturbation for molecular docking using flexible noise.

**Input:** molecule $\mathcal{M} = \{\mathbf{A}, \mathbf{X}, \mathbf{B}\}$, molecular noise $\xi$, noise scale $\beta$.

**Output:** noisy molecule $\tilde{\mathcal{M}} = \{\tilde{\mathbf{A}}, \tilde{\mathbf{X}}, \tilde{\mathbf{B}}\}$.

1: $\{\sigma, \epsilon, \kappa\} \leftarrow \xi$

2: $\tilde{\mathbf{A}}, \tilde{\mathbf{B}} \leftarrow \mathbf{A}, \mathbf{B}$

3: $\tilde{\mathbf{X}} = \Phi^{\text{flex}}(\mathbf{X}, \{\mathbf{A}, \mathbf{B}\}, \sigma, \epsilon, \kappa, \beta)$

##### 3.4.2 Molecular conformation generation

The molecular noise using Gaussian noise is defined as:

$$\xi = \{\sigma\} \tag{S43}$$

where $\sigma$ is the standard deviation of Gaussian noise for atom coordinates and is set as 1 in the implementation. The molecular noise using flexible noise is defined as:

$$\xi = \{\kappa\} \tag{S44}$$

where $\kappa$ is the parameter of the circular normal distribution of torsion angles and is set as 0.3. Unlike the flexible noise of docking, there is no noise for translation or rotation for conformation generation because the translation or rotation does not change the molecular conformations.

The perturbation processes for molecular conformation generation are the same as those for molecular docking (Algorithm 22 and 23). When using the perturbation algorithm with flexible noise (Algorithm 23), we can define fake parameters $\sigma = 0$ and $\epsilon = 0$ for translation and rotation noise.

##### 3.4.3 Structure-based drug design

The perturbation process for structure-based drug design regards the molecule as a complete part and defines the molecular noise as:

$$\xi = \{\bar{\mathbf{a}}, \bar{\mathbf{b}}, \sigma\} \tag{S45}$$

where $\bar{\mathbf{a}}$ and $\bar{\mathbf{b}}$ are the prior distribution of the categorical distributions for atom types and bond types, respectively, and $\sigma$ is the standard deviation of Gaussian distribution for atom coordinates. $\sigma$ is set as 1. $\bar{\mathbf{a}}$ is set as a normalized weight vector and the weight vector is defined as 3, 2, 2, 2, 1, 1, 1, 0.3, 0.3, 0.3, 0.3 and 13.2 for C, N, O, F, P, S, Cl, B, Br, I, Se, and the mask type, respectively. Similarly, $\bar{\mathbf{b}}$ uses the weight vector as 1, 1, 1, 1, 1, and 5 for the none-bond, single bond, double bond, triple bond, aromatic bond, and the mask type, respectively. The perturbation step is defined in Algorithm 24.

**Algorithm 24 PertSBDD:** perturbation for structure-based drug design

**Input:** molecule $\mathcal{M} = \{\mathbf{A}, \mathbf{X}, \mathbf{B}\}$, molecular noise $\xi$, noise scale $\beta$.

**Output:** noisy molecule $\tilde{\mathcal{M}} = \{\tilde{\mathbf{A}}, \tilde{\mathbf{X}}, \tilde{\mathbf{B}}\}$.

```
1: {ā, b̄, σ} ← ξ
2: for all i ∈ ⟦N⟧ do
3:     ã_i = Φ^atom(a_i, ā, β)
4:     x̃_i = Φ^Gauss(x_i, σ, β)
5: end for
6: for all i, j ∈ ⟦N⟧ do
7:     b̃_ij = Φ^bond(b_ij, b̄, β)
8: end for
```

##### 3.4.4 3D molecule generation

Both the molecular noise and the perturbation process for 3D molecule generation are the same as those for the structure-based drug design (Eq. S45 and Algorithm 24).

##### 3.4.5 Molecular optimization

In our setting, all variables of the molecules are allowed to be modified. Therefore, the molecular noise and the perturbation process have no difference from those for the structure-based drug design or 3D molecule generation (Eq. S45 and Algorithm 24).

##### 3.4.6 Fragment linking

For fragment linking, we divide the molecule into two parts: the fragment part $\mathbb{M}_{\text{frag}}$ and the linker part $\mathbb{M}_{\text{link}}$ and use different noise parameters for them. The fragment part includes the fragment atom set $\mathbb{A}_{\text{frag}}$ and the inner fragment bond set $\mathbb{B}_{\text{frag}}$. The linker part includes the linker atom set $\mathbb{A}_{\text{frag}}$, the inner linker bond set $\mathbb{B}_{\text{link}}$, and the bond set of atoms between the fragment and linker $\mathbb{B}_{\text{frag-link}}$. The molecular noise is defined as a mapping:

$$\xi(\mathbb{M}_k) = \begin{cases} \{\sigma_{\text{frag}}\}, & \mathbb{M}_k = \mathbb{A}_{\text{frag}} \cup \mathbb{B}_{\text{frag}} \\ \{\bar{\mathbf{a}}, \bar{\mathbf{b}}, \sigma_{\text{link}}\}, & \mathbb{M}_k = \mathbb{A}_{\text{link}} \cup \mathbb{B}_{\text{link}} \cup \mathbb{B}_{\text{frag-link}} \end{cases} \tag{S46}$$

The parameters for the fragment part and linker part are $\{\sigma_{\text{frag}}\}$ and $\{\bar{\mathbf{a}}, \bar{\mathbf{b}}, \sigma_{\text{frag}}\}$, respectively. $\sigma_{\text{frag}}$ is the standard deviation of Gaussian distribution for the coordinates of fragment atoms and is set as 1. $\bar{\mathbf{a}}, \bar{\mathbf{b}}, \sigma_{\text{frag}}$ are the parameters of the noise distributions for linker part. Their meaning and values are the same as those in the SBDD task (Eq. S45). The perturbation process is defined in Algorithm 25.

**Algorithm 25 PertFragLinking:** perturbation for fragment linking

**Input:** molecule $\mathcal{M} = \{\mathbf{A}, \mathbf{X}, \mathbf{B}\}$ , task prompt $\mathcal{P} = \{\mathbf{P}^{\text{atom}}, \mathbf{P}^{\text{coor}}, \mathbf{P}^{\text{bond}}, \mathbf{P}^{\text{dist}}, \mathbf{P}^{\text{pep}}\}$, molecular noise $\xi$, noise scale $\beta$.

**Output:** noisy molecule $\tilde{\mathcal{M}} = \{\tilde{\mathbf{A}}, \tilde{\mathbf{X}}, \tilde{\mathbf{B}}\}$.

```
Ã, X̃, B̃ ← A, X, B
M_frag, M_link ← the domain of ξ
{σ_frag} = ξ(M_frag)
{ā, b̄, σ_link} = ξ(M_link)
for all i ∈ {1, 2, ..., N} do
    if i ∈ M_frag then
        if P_i^coor = 0 then
            x̃_i = Φ^Gauss(x_i, σ_frag, β)
        end if
    else
        ã_i = Φ^atom(a_i, ā, β)
        x̃_i = Φ^Gauss(x_i, σ_link, β)
    end if
end for
for all i, j ∈ {1, 2, ..., N} do
    if ij ∈ B_link and P_ij^bond = 0 then
        b̃_ij = Φ^bond(b_ij, b̄, β)
    end if
end for
```

##### 3.4.7 PROTAC design

The molecular noise and the perturbation for PROTAC design are the same as those for fragment linking (Eq. S46 and Algorithm 25).

##### 3.4.8 Fragment growing

The molecular noise and the perturbation for fragment growing are similar to that for fragment linking (Eq. S46 and Algorithm 25), except that in fragment growing the molecule is divided into the fragment part and growing part. The growing part in fragment growing is treated in the same way as the linker part in fragment linking.

##### 3.4.9 De novo peptide design

For *de novo* peptide design, the molecule is divided into the backbone part $\mathbb{M}_{\text{bb}}$ and the side-chain part $\mathbb{M}_{\text{sc}}$. We define the backbone part as the atom set $\mathbb{A}_{\text{bb}}$, the inner backbone bond set $\mathbb{B}_{\text{bb}}$ and the bond set $\mathbb{B}_{\text{bbcon-sc}}$ of atoms between the backbone connecting atoms (alpha carbon and nitrogen atoms) and side-chain atoms. We define the side-chain part as the atom set $\mathbb{A}_{\text{sc}}$, the inner side-chain bond set $\mathbb{B}_{\text{sc}}$, and the bond set $\mathbb{B}_{\text{bbnoc-sc}}$ of atoms between the backbone non-connecting atoms (carbon and oxygen atoms) and side-chain atoms. The molecular noise is defined as a mapping:

$$\xi(\mathbb{M}_k) = \begin{cases} \{\sigma_{\text{bb}}\}, & \mathbb{M}_k = \mathbb{A}_{\text{bb}} \cup \mathbb{B}_{\text{bb}} \cup \mathbb{B}_{\text{bbcon-sc}} \\ \{\bar{\mathbf{a}}, \bar{\mathbf{b}}, \sigma_{\text{sc}}\}, & \mathbb{M}_k = \mathbb{A}_{\text{sc}} \cup \mathbb{B}_{\text{sc}} \cup \mathbb{B}_{\text{bbnoc-sc}} \end{cases} \tag{S47}$$

where $\sigma_{\text{bb}}$ and $\sigma_{\text{sc}}$ are the standard deviations of Gaussian distributions for backbone atoms and side-chain atoms, respectively, and both are set as three. $\bar{\mathbf{a}}$ is the prior distribution of the categorical distribution for atom types and set as a normalized vector with weights 75 for C, 10 for N, 13 for O, 0.3 for S atoms, 98 for the mask type, and 0 for all other element types. $\bar{\mathbf{b}}$ is the prior distribution of the categorical distribution for atom types and set as a normalized vector with 1, 1, 1, 1, 1, and 5 for none-type, single bond, double bond, triple bond, aromatic bond, and the mask type, respectively. The perturbation process is defined in Algorithm 26.

**Algorithm 26 PertPepDesign:** perturbation for *de novo* peptide design

**Input:** molecule $\mathcal{M} = \{\mathbf{A}, \mathbf{X}, \mathbf{B}\}$ , molecular noise $\xi$, noise scale $\beta$.

**Output:** noisy molecule $\tilde{\mathcal{M}} = \{\tilde{\mathbf{A}}, \tilde{\mathbf{X}}, \tilde{\mathbf{B}}\}$.

```
$\tilde{\mathbf{A}}, \tilde{\mathbf{X}}, \tilde{\mathbf{B}} \leftarrow \mathbf{A}, \mathbf{X}, \mathbf{B}$
$\mathbb{M}_{\text{bb}}, \mathbb{M}_{\text{sc}} \leftarrow$ the domain of $\xi$
$\{\sigma_{\text{bb}}\} = \xi(\mathbb{M}_{\text{bb}})$
$\{\bar{\mathbf{a}}, \bar{\mathbf{b}}, \sigma_{\text{sc}}\} = \xi(\mathbb{M}_{\text{sc}})$
for all $i \in [\![N]\!]$ do
    if $i \in \mathbb{M}_{\text{bb}}$ then
        $\tilde{\mathbf{x}}_i = \Phi^{\text{Gauss}}(\mathbf{x}_i, \sigma_{\text{bb}}, \beta)$
    else
        $\tilde{\mathbf{a}}_i = \Phi^{\text{atom}}(\mathbf{a}_i, \bar{\mathbf{a}}, \beta)$
        $\tilde{\mathbf{x}}_i = \Phi^{\text{Gauss}}(\mathbf{x}_i, \sigma_{\text{sc}}, \beta)$
    end if
end for
for all $i, j \in [\![N]\!]$ do
    if $ij \in \mathbb{M}_{\text{sc}}$ then
        $\tilde{\mathbf{b}}_{ij} = \Phi^{\text{bond}}(\mathbf{b}_{ij}, \bar{\mathbf{b}}, \beta)$
    end if
end for
```

Linear peptides, cyclic peptides, and peptides containing non-standard amino acids share the same noise definitions.

###### *3.4.10 Peptide inverse folding*

The molecular noise and the perturbation process for peptide inverse folding are similar to those for de novo peptide design. The difference is that no noise is added for the backbone atom coordinates. Similarly, we divided the molecule into the backbone part and the side-chain part. The backbone part is the atom set $\mathbb{A}_{\text{bb}}$, the inner backbone bond set $\mathbb{B}_{\text{bb}}$ and the bond set $\mathbb{B}_{\text{bbcon-sc}}$ of atoms between the backbone connecting atoms (alpha carbon and nitrogen atoms) and side-chain atoms. The side-chain part is the atom set $\mathbb{A}_{\text{sc}}$, the inner side-chain bond set $\mathbb{B}_{\text{sc}}$, and the bond set $\mathbb{B}_{\text{bbnoc-sc}}$ of atoms between the backbone non-connecting atoms (carbon and oxygen atoms) and side-chain atoms. The molecular noise is defined as a mapping function as the following,

$$\xi(\mathbb{M}_k) = \begin{cases} \varnothing, & \mathbb{M}_k = \mathbb{A}_{\text{bb}} \cup \mathbb{B}_{\text{bb}} \cup \mathbb{B}_{\text{bbcon-sc}} \\ \{\bar{\mathbf{a}}, \bar{\mathbf{b}}, \sigma_{\text{sc}}\}, & \mathbb{M}_k = \mathbb{A}_{\text{sc}} \cup \mathbb{B}_{\text{sc}} \cup \mathbb{B}_{\text{bbnoc-sc}} \end{cases} \tag{S48}$$

where the parameters have the same meaning and values as those of the *de novo* peptide design. The perturbation function is defined in Algorithm 27.

**Algorithm 27 PertPepInvFolding:** perturbation for peptide inverse folding

**Input:** molecule $\mathcal{M} = \{\mathbf{A}, \mathbf{X}, \mathbf{B}\}$, molecular noise $\xi$, noise scale $\beta$.

**Output:** noisy molecule $\tilde{\mathcal{M}} = \{\tilde{\mathbf{A}}, \tilde{\mathbf{X}}, \tilde{\mathbf{B}}\}$.

```
1: Ã, X̃, B̃ ← A, X, B
2: M_bb, M_sc ← the domain of ξ
3: {ā, b̄, σ_sc} = ξ(M_sc)
4: for all i ∈ ⟦N⟧ do
5:     if i ∈ M_sc then
6:         ã_i = Φ^atom(a_i, ā, β)
7:         x̃_i = Φ^Gauss(x_i, σ_sc, β)
8:     end if
9: end for
10: for all i, j ∈ ⟦N⟧ do
11:     if ij ∈ M_sc then
12:         b̃_ij = Φ^bond(b_ij, b̄, β)
13:     end if
14: end for
```

### 4 Generation process

#### 4.1 General generation process

For a generative task, we need to define the task prompt $\mathcal{P}$ and provide the pocket $\mathcal{K}$ and the initial molecule $\mathcal{M}_{\text{init}}$. The pocket $\mathcal{K}$ can be set as $\varnothing$ to indicate that the generation is not conditioned on any pocket. The initial molecule $\mathcal{M}_{\text{init}}$ can be provided by users or sampled from molecular noise distributions. Then the model repeatedly adds noise and removes noise for the molecule, with a predefined sequence of noise scale $\boldsymbol{\beta} = [\beta^1, \beta^2, \ldots, \beta^T]$. A modification function $\Psi$ can be invoked to modify the molecule to align with prior knowledge before adding noise in each step. The general generation process is defined as Algorithm 28.

**Algorithm 28 GenerationProcess:** general generation process of PocketXMol

**Input:** initial molecule $\mathcal{M}_{\text{init}}$, pocket $\mathcal{K}$, task prompt $\mathcal{P}$, molecular noise $\xi$, total step $T$, sequence of noise scale $\boldsymbol{\beta} = [\beta^1, \beta^2, \ldots, \beta^T]$, (optional) prior-knowledge-guided modification function $\Psi$.

**Output:** generated molecule $\mathcal{M}_{\text{final}}$, last confidence scores $\mathcal{S}_{\text{final}}$, trajectory confidence scores $\mathcal{S}_{\text{traj}}$.

$\mathcal{M}^0 = \mathcal{M}_{\text{init}}$
**for all** $t \in [\![T]\!]$ **do**
    **if** $\Psi$ is provided **then**
        $\tilde{\mathcal{M}}^t = \Psi(\mathcal{M}^{t-1}, \mathcal{K}, \mathcal{P}, t)$    ▷ modify based on prior knowledge
    **else**
        $\tilde{\mathcal{M}}^t = \mathcal{M}^t$
    **end if**
    $\tilde{\mathcal{M}}^t \leftarrow \Phi(\tilde{\mathcal{M}}^t, \mathcal{P}, \xi, \beta^t)$    ▷ add noise to molecule
    $\mathcal{M}^t, \mathcal{S}^t = F_\Theta(\tilde{\mathcal{M}}^t, \mathcal{P}, \mathcal{K})$    ▷ remove noise using denoiser
**end for**
$\mathcal{M}_{\text{final}} = \mathcal{M}^T$
$\mathcal{S}_{\text{final}} = \mathcal{S}^T$
$\mathcal{S}_{\text{traj}} = [\mathcal{S}^1, \ldots, \mathcal{S}^T]$

#### 4.2 Schedule of noise scale

The schedule of the noise scale $\boldsymbol{\beta} = [\beta^1, \beta^2, \ldots, \beta^T]$ can be any pre-defined sequences of values between zero and ones. In our implementation, we mainly adopted two forms of schedules. The first one is a decaying schedule based on the sigmoid function, proposed in previous work[5]. Given the total step $T$, the parameters of the range $(\beta_{\text{min}}, \beta_{\text{max}})$, and a hyper-parameter $k$ controlling the slope of the curve, the value of noise scale $\beta^t$ for $t \in [\![T]\!]$ is defined as:

$$\begin{aligned}
\tau &= 1 - \frac{t-1}{T} \\
\alpha_{\text{max}} &= 1 - \beta_{\text{min}} \\
\alpha_{\text{min}} &= 1 - \beta_{\text{max}} \\
w &= \frac{\alpha_{\text{min}} - \alpha_{\text{max}}}{\text{sigmoid}(-k) - \text{sigmoid}(k)} \\
u &= 0.5 \times (\alpha_{\text{max}} + \alpha_{\text{min}} - w) \\
\beta^t &= 1 - w \times \text{sigmoid}\left(-k(2\tau/T - 1)\right) + u
\end{aligned} \tag{S49}$$

Note that $\tau$ decreases from 1 to $1/T$ as $t$ increases from 1 to $T$. In our implementation (source code), the variable $\tau$ instead of $t$ was directly used and we maintained $1 - \beta^t$ instead of $\beta^t$. We defined $\Upsilon(\beta_{\text{max}}, \beta_{\text{min}}, k, T)$ as the sequences of noise scale $\boldsymbol{\beta} = [\beta^1, \beta^2, \ldots, \beta^T]$ derived from Eq. S49. We further defined $\Upsilon'$ to be the same as $\Upsilon$ except that the first value was set as one ($\beta^1 = 1$). Note that $\beta^1 = 1$ will remove all information about the molecule in the perturbation step, which is used to sample the initial molecule directly from the noise.

The second one is a uniform form, which defines the noise scale as a linear decaying function of the step $t$. Specifically, given the total step $T$, the range parameters $(\beta_{\text{min}}, \beta_{\text{max}})$, the noise scale $\beta^t$

is defined as:

$$\begin{aligned}\tau &= 1 - \frac{t-1}{T} \\ \alpha_{\max} &= 1 - \beta_{\min} \\ \alpha_{\min} &= 1 - \beta_{\max} \\ \beta^t &= 1 - (1-\tau)(\alpha_{\max} - \alpha_{\min}) - \alpha_{\min}\end{aligned} \tag{S50}$$

This scheduler function is denoted as $\Gamma(\beta_{\max}, \beta_{\min}, T)$. Similarly, we defined $\Gamma'$ to be the same as $\Gamma$ except that the first value $\beta^1$ was set as one.

#### 4.3 Generation settings for individual tasks

##### *4.3.1 Small molecule docking*

For protein-small molecule docking, both the molecule pose and the score representing the docking confidence should be produced. For protein-small molecule docking, either Gaussian noise or flexible noise can be used in the generation process. We can choose one of them and apply the generation process defined in Algorithm 29. The preparation of the initial molecule for docking using Gaussian noise is in Algorithm 30.

---

**Algorithm 29 SimpleGenDockSmallMol:** simple generation process for protein-small molecule docking

---

**Input:** molecular atoms and bonds $\{\mathbf{A}, \mathbf{B}\}$, protein pocket $\mathcal{K}$, selection $\varkappa$ of Gaussian noise or flexible noise.

**Output:** generated molecule $\mathcal{M}$, last confidence scores $\mathcal{S}$, trajectory confidence scores $\mathcal{S}_{\text{traj}}$.

▷ Set parameters and initialization
**if** $\varkappa =$ `Gaussian` **then**
  $\mathcal{P} \leftarrow$ define task prompt using Eq. S11
  $\xi \leftarrow$ define molecular noise using Eq. S41
  $N \leftarrow$ the number of the atoms of the molecule $\{\mathbf{A}, \mathbf{B}\}$
  $\mathbf{X} =$ PrepareInitCoordinates$(N, \mathcal{K})$
**else if** $\varkappa =$ `flexible` **then**
  $\mathcal{P} \leftarrow$ define task prompt using Eq. S11 and Algorithm 17
  $\xi \leftarrow$ define molecular noise using Eq. S42
  $\mathbf{X} \leftarrow$ generate a conformation using RDKit
  **if** $\mathcal{K} \neq \varnothing$ **then**
    $\mathbf{X} \leftarrow$ put it at the center of the pocket $\mathcal{K}$
  **end if**
**end if**
$\mathcal{M}_{\text{init}} = \{\mathbf{A}, \mathbf{X}, \mathbf{B}\}$
$T = 100$
$\boldsymbol{\beta} = \Upsilon'(\beta_{\max} = 1 - 10^{-5}, \beta_{\min} = 10^{-5}, k = 3, T)$
$\mathcal{M}, \mathcal{S}, \mathcal{S}_{\text{traj}} =$ GenerationProcess$(\mathcal{M}_{\text{init}}, \mathcal{K}, \mathcal{P}, \xi, T, \boldsymbol{\beta})$

---

**Algorithm 30 PrepareInitCoordinates:** prepare the initial coordinates for docking

**Input:** the number of atoms of the molecule $N$, protein pocket $\mathcal{K}$.

**Output:** initial coordinates $\mathbf{X}$.

**if** $\mathcal{K} = \varnothing$ **then**

  $\bar{\mathbf{x}} \leftarrow \mathbf{0}_3$

**else**

  $\bar{\mathbf{x}} \leftarrow$ average of atom coordinates of the pocket $\mathcal{K}$

**end if**

$\mathbf{X} = [\bar{\mathbf{x}}; \bar{\mathbf{x}}; \ldots; \bar{\mathbf{x}}] \in \mathbb{R}^{N \times 3}$

For generative docking, it is common to generate multiple poses and select one pose using some ranking scores[6,7]. Here, we can rely on the confidence scores produced by our model for ranking. Additionally, following AlphaFold 3, we also consider the penalties for violating stereochemistry and clashes. The stereochemistry penalty was calculated by comparing the stereochemistry between the generated molecules and the true one and adding the penalty if they were not identical. The clash penalty was added if the generated molecules were too close to the pockets, which were measured by comparing the pairwise atom distances and the der Waals radiuses. The complete generative docking process is in Algorithm 31 and the self ranking scores are calculated in Algorithm 32.

**Algorithm 31 GenDockSmallMol:** generation process for protein-small-molecule docking

**Input:** molecular atoms and bonds $\{\mathbf{A}, \mathbf{B}\}$, protein pocket $\mathcal{K}$, selection $\varkappa$ of Gaussian noise or flexible noise.

**Output:** generated molecule $\mathcal{M}$, ranking score $s$.

$\mathcal{M}, \mathcal{S}, \mathcal{S}_{\text{traj}} =$ SimpleGenDockSmallMol$(\{\mathbf{A}, \mathbf{B}\}, \mathcal{K}, \varkappa)$

$s =$ GenSelfRankingScore$(\mathcal{M}, \mathcal{K}, \mathcal{S}_{\text{traj}}, \varkappa)$

**Algorithm 32 GenSelfRankingScore:** get self ranking scores

**Input:** Generated molecule $\mathcal{M} = \{\mathbf{A}, \mathbf{X}, \mathbf{B}\}$, protein pocket $\mathcal{K}$, trajectory confidence scores $\mathcal{S}_{\text{traj}}$, selection $\varkappa$ of Gaussian noise or flexible noise.

**Output:** self-ranking score $s$.

$[\mathbf{S}_1^{\text{coor}}, \ldots, \mathbf{S}_T^{\text{coor}}] \leftarrow$ coordinate trajectory confidence scores from $\mathcal{S}_{\text{traj}}$
$\forall t \in [\![T]\!], \bar{s}_t = \text{mean}(\mathbf{S}_t^{\text{coor}})$ ▷ Average over all atoms
**if** $\varkappa =$ `Gaussian` **then**
    $s =$ average over $[\bar{s}_{\lfloor T/2 \rfloor}, \ldots, \bar{s}_T]$ ▷ Last half trajectory
**else if** $\varkappa =$ `flexible` **then**
    $s = \bar{s}_T$ ▷ Last one only
**end if**
**if** $\mathcal{M}$ violates sterechemisty **then**
    $s \leftarrow s - 1$
**end if**
**if** $\mathcal{M}$ has clashes with $\mathcal{K}$ **then**
    $s \leftarrow s - 1$
**end if**

In our experiments, we observed that docking using Gaussian noise has a better generation ability than flexible noise and thus used it as the default docking option. In practice, given a protein pocket, we generated $n_{\text{gen}}$ molecule poses ($n_{\text{gen}} = 100$ in our evaluation), sorted them according to the ranking scores, and provided $n_{\text{sel}}$ molecule poses ($n_{\text{sel}} = 1$ in our evaluation) as the docking results, as defined in Algorithm 33.

**Algorithm 33 GenSelDockSmallMol:** generation and selection for protein-small molecule docking

**Input:** molecular atoms and bonds $\{\mathbf{A}, \mathbf{B}\}$, protein pocket $\mathcal{K}$, number of generated poses $n_{\text{gen}}$, number of selected poses $n_{\text{sel}}$.

**Output:** a set $\mathbf{M}$ of $n_{\text{sel}}$ generated molecule poses.

**for all** $i \in [n_{\text{gen}}]$ **do**
    $\mathcal{M}_i, s_i =$ GenDockSmallMol$(\{\mathbf{A}, \mathbf{B}\}, \mathcal{K})$
**end for**
sorted indices $\boldsymbol{\eta} \leftarrow$ sort $[s_1, s_2, \ldots, s_{n_{\text{gen}}}]$ in descending order
$\mathbf{M} = \{\mathcal{M}_{\eta_i} | i \in [\![n_{\text{sel}}]\!]\}$

We also explored more ranking strategies. We considered training a separate pose ranker to score the protein-small molecule structures. This ranker was obtained through further fine-tuning the PocketXMol by only using the confidence losses (see Supplementary Notes 5.2). The tuned ranking scores are calculated in Algorithm 34.

**Algorithm 34 GenTunedRankingScore:** get tuned ranking scores

**Input:** Generated molecule $\mathcal{M} = \{\mathbf{A}, \mathbf{X}, \mathbf{B}\}$, protein pocket $\mathcal{K}$, tuned ranker $F_\Theta^{\text{ranker}}$.

**Output:** tuned ranking score $s$.

$\mathcal{S} = F_\Theta^{\text{ranker}}(\mathcal{M}, \mathcal{K})$
$\mathbf{S}^{\text{atom}}, \mathbf{S}^{\text{coor}}, \mathbf{S}^{\text{bond}} = \mathcal{S}$
$s = \text{mean}(\mathbf{S}^{\text{coor}})$
**if** $\mathcal{M}$ violates sterechemisty **then**
    $s \leftarrow s - 1$
**end if**
**if** $\mathcal{M}$ has clashes with $\mathcal{K}$ **then**
    $s \leftarrow s - 1$
**end if**

We also found that the denoiser can directly serve as a pose ranker given the task prompt of docking with flexible noise. Therefore, for a molecular pose to be scored, we can use the molecule as the initial molecule to re-dock using flexible noise and prompt. But we infer only one step and do not add substantial noise to the molecule. Then the output confidence scores can be used for ranking. This is defined in Algorithm 35. The ranking strategy had a slightly weaker performance than the previous two ranking strategies for docking. However, the advantage of this ranking strategy is that any protein-small molecule/peptide complex (even those obtained through other tasks) can be fed back to the model to obtain a confidence score without the need to fine-tune a new ranker. For example, we can design novel molecules using PocketXMol and then use this strategy to score the designed molecules for selection.

**Algorithm 35 GenFlexRankingScore:** get confidence scores using flexible noise

**Input:** Generated molecule $\mathcal{M} = \{\mathbf{A}, \mathbf{X}, \mathbf{B}\}$, protein pocket $\mathcal{K}$.

**Output:** confidence score $s$.

$\mathcal{P} \leftarrow$ define task prompt Eq. S11 and Algorithm 17 (flexible noise)
$\xi \leftarrow$ define molecular noise using Eq. S42 (flexible noise)
$T = 1$
$\boldsymbol{\beta} = [0]$ ▷ No noise is added
$\mathcal{M}', \mathcal{S}_{\text{flex}} = \text{GenerationProcess}(\mathcal{M}, \mathcal{K}, \mathcal{P}, \xi, T, \boldsymbol{\beta})$
$\mathbf{S}^{\text{atom}}, \mathbf{S}^{\text{coor}}, \mathbf{S}^{\text{bond}} \leftarrow \mathcal{S}$
$s = \text{mean}(\mathbf{S}^{\text{coor}})$ ▷ average of the confidence scores of atom coordinates

Now we introduce how to incorporate different types of prior knowledge into the docking process.

**Molecular center.** The prior knowledge is that we know the approximate position of the center of the docked molecule, which can be incorporated into the docking process by simply translating the denoised molecule if its center is not located at the prior position. In our implementation, we assume the center position is within a sphere with a radius of 2 Å and the center at the true molecular pose's center.

**Algorithm 36 ModifyCenter:** modify the molecule to align the reference center position

**Input:** denoised molecule $\mathcal{M}$, reference center $\mathbf{x}^{\text{ref}}$.

**Output:** modified molecule $\mathcal{M}^{\text{mod}}$.

$\mathbf{A}, \mathbf{X}, \mathbf{B} \leftarrow \mathcal{M}$

$\mathbf{x}^{\text{center}} = \text{mean}(\mathbf{X})$

$\Delta\mathbf{x} = \mathbf{x}^{\text{ref}} - \mathbf{x}^{\text{center}}$

$d = \|\Delta\mathbf{x}\|$

**if** $d > 2$ **then**

  $\forall i, \mathbf{x}_i^{\text{mod}} = \mathbf{x}_i + \Delta\mathbf{x}$

  $\mathcal{M}^{\text{mod}} = \{\mathbf{A}, \mathbf{X}^{\text{mod}}, \mathbf{B}\}$

**else**

  $\mathcal{M}^{\text{mod}} = \mathcal{M}$

**end if**

**Bond lengths.** Knowing the bond lengths of the molecule indicates some atom distances are known (Algorithm 17). Therefore, we can incorporate this information by modifying some atom distances of the denoised molecule. In our implementation, we provide a reference molecule with a valid conformation and use gradient descent to modify the atom distances of the denoised molecule, as defined in Algorithm 37.

**Algorithm 37 ModifyBondLength:** modify the molecule to align the bond lengths

**Input:** denoised molecule $\mathcal{M}$, reference molecule $\mathcal{M}^{\text{ref}}$, prompt indicators for atom distances $\mathbf{P}^{\text{coor}}$.

**Output:** modified molecule $\mathcal{M}^{\text{mod}}$.

$\mathbf{A}, \mathbf{X}, \mathbf{B} \leftarrow \mathcal{M}$
$N \leftarrow$ number of atoms of $\mathcal{M}$
$\mathbf{P}^{\text{dist}} = \text{GetDistPromptFlexNoise}(\mathcal{M})$
$[d_{ij}^{\text{ref}}] \leftarrow$ pairwise distances of $\mathcal{M}^{\text{ref}}$
**for all** $\tau \in [\![10]\!]$ **do**
    $[d_{ij}] \leftarrow$ pairwise distances of $\mathcal{M}$
    **for all** $i \in [\![N]\!]$ **do**
        **if** $P_i^{\text{coor}} = 0$ and $\sum_j P_{ij}^{\text{dist}} > 0$ **then**
            $\Delta \mathbf{x}_i = \sum_j [2P_{ij}^{\text{dist}}(d_{ij} - d_{ij}^{\text{ref}})/d_{ij}^{\text{ref}}(\mathbf{x}_i - \mathbf{x}_j)] / \sum_j P_{ij}^{\text{dist}}$
            $\mathbf{x}_i \leftarrow \mathbf{x}_i - 0.05\Delta \mathbf{x}_i$
        **end if**
    **end for**
**end for**
$\mathcal{M}^{\text{mod}} = \{\mathbf{A}, \mathbf{X}, \mathbf{B}\}$

**Anchor atom coordinates.** Similar to the molecular center, if the approximate coordinates of some atoms are known, the denoised molecule can be translated so that the pose aligns with the prior knowledge. In our implementation, we defined the anchor atom as the atom of the molecule that is closest to the protein atoms in the dataset and assumed this atom should be located at a sphere with radius 2 Å, as defined in Algorithm 38.

**Algorithm 38 ModifyAtomCoordinate:** modify the molecule to align the reference atom coordinate

**Input:** denoised molecule $\mathcal{M}$, reference atom index $k$, reference coordinates $\mathbf{x}^{\text{ref}}$.

**Output:** modified molecule $\mathcal{M}^{\text{mod}}$.

$\mathbf{A}, \mathbf{X}, \mathbf{B} \leftarrow \mathcal{M}$
$\Delta \mathbf{x} = \mathbf{x}^{\text{ref}} - \mathbf{x}_k$
$d = \|\Delta \mathbf{x}\|$
**if** $d > 2$ **then**
    $\forall i, \mathbf{x}_i^{\text{mod}} = \mathbf{x}_i + \Delta \mathbf{x}$
    $\mathcal{M}^{\text{mod}} = \{\mathbf{A}, \mathbf{X}^{\text{mod}}, \mathbf{B}\}$
**else**
    $\mathcal{M}^{\text{mod}} = \mathcal{M}$
**end if**

Furthermore, if the atom coordinate is exactly known, i.e., the atom coordinate is fixed, we can directly incorporate this prior knowledge into the task prompt. Assume the anchor atom index is $k$, we only need to modify the prompt of coordinates as:

$$\forall i, P_i^{\text{coor}} = \begin{cases} 1, & i = k \\ 0, & \text{otherwise} \end{cases} \tag{S51}$$

###### 4.3.2 Peptide docking

The generation process for protein-peptide docking is exactly the same as that for protein-small molecule docking (Algorithm 31).

In practice, for protein-peptide docking, the peptide was usually provided using `.pdb` files. We only need to extract the atom types and the bond types from the file using RDKit and discard the information about residue types. Then treat them the same as small molecules during the generation process. Additionally, linear peptides, cyclic peptides, and peptides containing non-standard amino acids share the same generation process.

Now we introduce how to incorporate prior knowledge into the docking process. In our experiments, we showed peptide docking with exactly known coordinates of different atoms, all of which can be implemented by modifying the prompt indicators of coordinates. Assume $\mathbb{A}_{\text{fixed}}$ be the index set of atoms with exactly known coordinates, the prompt indicators of coordinates are defined as:

$$\forall i, P_i^{\text{coor}} = \begin{cases} 1, & i \in \mathbb{A}_{\text{fixed}} \\ 0, & \text{otherwise} \end{cases} \tag{S52}$$

Docking with fixed anchor atoms defines $\mathbb{A}_{\text{fixed}}$ as the atom of the peptide that is closest to the protein atoms. Docking with fixed first residues defines $\mathbb{A}_{\text{fixed}}$ as the atoms of the first residues. Docking with terminal residues defines $\mathbb{A}_{\text{fixed}}$ as the atoms of the first and the last residues. Docking with fixed backbone atoms defines $\mathbb{A}_{\text{fixed}}$ as all the backbone atoms.

###### 4.3.3 Molecular conformation generation

For molecular conformation generation, the generation process is similar to that for molecular docking. The difference is that this task does not require the pocket as input. Besides, confidence is not required in this task and we only use Gaussian noise as the default noise during generation. The generation process is defined in Algorithm 39.

**Algorithm 39 GenConf:** generation process for molecular conformation generation

**Input:** molecular atoms and bonds $\{\mathbf{A}, \mathbf{B}\}$.

**Output:** generated molecule $\mathcal{M}_{\text{gen}}$.

1: $\mathcal{P} \leftarrow$ derive task prompt using Eq. S11
2: $\Omega \leftarrow$ define molecular noise using Eq. S43
3: $\mathcal{K} = \varnothing$ ▷ empty pocket
4: $N \leftarrow$ The number of the atoms of the molecule
5: $\mathbf{X} = \text{PrepareInitCoordinates}(N, \mathcal{K})$
6: $\mathcal{M}_{\text{init}} = \{\mathbf{A}, \mathbf{X}, \mathbf{B}\}$
7: $T = 100$
8: $\boldsymbol{\beta} = \Upsilon'(\beta_{\max} = 1 - 10^{-5}, \beta_{\min} = 10^{-5}, k = 3, T)$
9: $\mathcal{M}_{\text{gen}}, \mathcal{S}_{\text{gen}}, \mathcal{S}_{\text{traj}} = \text{GenerationProcess}(\mathcal{M}_{\text{init}}, \mathcal{K}, \mathcal{P}, \Omega, T, \boldsymbol{\beta})$

###### 4.3.4 Structure-based drug design

For structure-based drug design (SBDD), PocketXMol can adopt a generation process similar to the previous diffusion-based SBDD models[8,9], as defined in Algorithm 40, which takes the pocket and the number of atoms of the generated molecules as inputs. In our experiments, the number of atoms of the molecules can be determined according to the sizes of the pockets (Algorithm 42), which is based on the observation that the numbers of atoms of the molecules relate to the sizes of the pockets in the dataset. In practice, the number of atoms of the generated molecules can also be sampled from other distributions. For example, it can be sampled from a distribution related to the reference molecules (Algorithm 43).

**Algorithm 40 SimpleGenSBDD:** simple generation process for structure-based drug design

**Input:** number of molecular (heavy) atoms $N$, protein pocket $\mathcal{K}$.

**Output:** generated molecule $\mathcal{M}_{\text{gen}}$, confidence scores $\mathcal{S}_{\text{gen}}$.

1: $\mathcal{M}_{\text{init}} = \text{PrepareInitMolecule}(N, \mathcal{K})$
2: $\mathcal{P} \leftarrow$ derive task prompt for SBDD using Eq. S12
3: $\xi \leftarrow$ define molecular noise using Eq. S45
4: $T = 100$
5: $\boldsymbol{\beta} = \Upsilon'(\beta_{\max} = 1 - 10^{-5}, \beta_{\min} = 10^{-5}, k = 3, T)$
6: $\mathcal{M}_{\text{gen}}, \mathcal{S}_{\text{gen}}, \mathcal{S}_{\text{traj}} = \text{GenerationProcess}(\mathcal{M}_{\text{init}}, \mathcal{K}, \mathcal{P}, \xi, \boldsymbol{\beta}, T)$

**Algorithm 41 PrepareInitMolecule:** prepare initial molecule for molecule design

**Input:** number of molecular atoms $N$, protein pocket $\mathcal{K}$.

**Output:** initialize molecule $\mathcal{M} = \{\mathbf{A}, \mathbf{X}, \mathbf{B}\}$.

1: $\mathbf{X} = \text{PrepareInitCoordinates}(N, \mathcal{K})$
2: $\mathbf{A} = \mathbf{0}_{N \times K_{\text{atom}}}$
3: $\mathbf{B} = \mathbf{0}_{N \times N \times K_{\text{bond}}}$
4: $\mathcal{M} = \{\mathbf{A}, \mathbf{X}, \mathbf{B}\}$

**Algorithm 42 GetMolSizeFromPocket:** sample number of molecular atoms based on pocket size

**Input:** protein pocket $\mathcal{K}$.

**Output:** the number $N$ of atoms of the molecule for the pocket.

$N_{\text{pocket}} \leftarrow$ the number of atoms of the pocket $\mathcal{K}$
$\mu = 0.0592 N_{\text{pocket}} + 0.1478$
$\sigma = 6.3037$
$N_{\text{min}} = 5$
**while** true **do**
    $N \sim \mathcal{N}(\mu, \sigma^2)$
    $N \leftarrow$ round of $N$
    **if** $N \geq N_{\text{min}}$ **then**
        break
    **end if**
**end while**

**Algorithm 43 GetMolSizeFromRef:** sample number of molecular atoms based on reference size

**Input:** the number $N_{\text{ref}}$ of atoms of the reference molecule.

**Output:** the number $N$ of atoms of molecule to be generated.

$\sigma = 0.02 N_{\text{ref}} + 1$
$N_{\text{min}} = 5$
**while** true **do**
    $N \sim \mathcal{N}(N_{\text{ref}}, \sigma^2)$
    $N \leftarrow$ round of $N$
    **if** $N \geq N_{\text{min}}$ **then**
        break
    **end if**
**end while**

Compared with diffusion-based models, our denoiser additionally predicts the confidence scores for each individual variable of the generated molecules, which can be utilized to further re-generate particular parts of the molecules with low confidence scores for better performance. We designed a novel generation algorithm for SBDD (Algorithm 44), which iteratively refines molecule generation by applying higher noise scales to molecular parts with low confidence scores for multiple rounds. Initially, molecules are generated using standard noise distribution (Eq. S45) and noise scale. Subsequently, atoms and bonds with low confidence scores are identified as parts requiring re-generation. These parts are regenerated using standard noise scales ranging from 1 to 0, while the rest of the molecules are with much lower noise scales in this round. These regeneration processes aim to enhance low-confidence parts while minimizing the modification to the rest of the parts.

The re-generated part is defined in Algorithm 45, which first determines atoms with low confidence scores of the predicted atom types, atom coordinates, and chemical bond types and then defines the re-generated part as atoms and bonds close to these low-confidence atoms. The molecular noise and the noise scale in the re-generation process are similar to those in the standard generation process. The only difference is that the noise scale for the non-re-generated parts is much lower and its schedule is defined as a linear decay from a value less than 1 to 0 (Algorithm 47).

**Algorithm 44 GenSBDD:** generation process with refining using confidence scores for SBDD

**Input:** the number of molecular atoms $N$, protein pocket $\mathcal{K}$.

**Output:** generated molecule $\mathcal{M}_{\text{gen}}$, confidence scores $\mathcal{S}_{\text{gen}}$.

```
𝒫 ← derive task prompt for SBDD using Eq. S12
T = 100
ℳ⁰, 𝒮⁰ = SimpleGenSBDD(N, 𝒦)
▷ Re-generate
n_regen = 9
for all τ ∈ ⟦n_regen⟧ do
    ℳ^τ_init, ξ, β = PrepareRegen(ℳ^{τ−1}, 𝒮^{τ−1}, 𝒦, T)
    if ℳ^{τ−1} = ℳ^τ_init then                              ▷ no need for re-generation
        ℳ^τ = ℳ^{τ−1}, 𝒮^τ = 𝒮^{τ−1}
        break
    end if
    ℳ^τ, 𝒮^τ, 𝒮^τ_traj = GenerationProcess(ℳ^τ_init, 𝒦, 𝒫, ξ, β, T)
end for
ℳ_gen = ℳ^τ, 𝒮_gen = 𝒮^τ
```

**Algorithm 45 PrepareRegen:** prepare for re-generation round of SBDD

**Input:** generated molecule $\mathcal{M}$, confidence scores $\mathcal{S}$, total step $T$.

**Output:** initial molecule $\mathcal{M}_{\text{init}}$ for re-generation, molecular noise $\xi$, noise scale $\boldsymbol{\beta}$.

$\triangleright$ Get atoms with low confidence scores
$\mathbb{B}^{\text{chem}} \leftarrow$ chemical bond set of $\mathcal{M}_{\text{init}}$
$[s_i^{\text{atom}}], [s_i^{\text{coor}}], [s_{ij}^{\text{bond}}] = \mathcal{S}$
$\forall i, s_i^{\text{atom\_bonds}} \leftarrow$ average items of $\{s_{ij}^{\text{bond}} | ij \in \mathbb{B}^{\text{chem}}\}$
$\check{s}^{\text{atom}} = 0.98,\ \check{s}^{\text{coor}} = 0.91,\ \check{s}^{\text{atom\_bond}} = 0.98$ $\triangleright$ set thresholds for low confidence scores
$\mathbb{A}_{\text{low}} \leftarrow \{i | s_i^{\text{atom}} \le \check{s}^{\text{atom}} \text{ or } s_i^{\text{coor}} \le \check{s}^{\text{coor}} \text{ or } s_i^{\text{atom\_bond}} \le \check{s}^{\text{bond}}\}$
$\triangleright$ Get re-generated parts
$[\mathbf{x}_i] \leftarrow$ atom coordinates of $\mathcal{M}$
$\check{d} = 3$ $\triangleright$ set distance threshold
$\mathbb{A}_{\text{regen}} = \{i | \exists j \in \mathbb{A}_{\text{low}}, \|\mathbf{x}_i - \mathbf{x}_j\| \le \check{d}\}$
$\mathbb{B}_{\text{regen}} = \{ij | i \in \mathbb{A}_{\text{regen}} \text{ or } j \in \mathbb{A}_{\text{regen}}\}$
$\triangleright$ Prepare initial molecule
$\mathcal{M}_{\text{init}} = \text{PrepareRegenInitMolecule}(\mathcal{M}, \mathbb{A}_{\text{regen}}, \mathbb{B}_{\text{regen}})$
$\triangleright$ Define noise scale
$\xi, \beta = \text{DefineRegenNoise}(\mathbb{A}_{\text{regen}}, \mathbb{B}_{\text{regen}}, \mathcal{M}_{\text{init}}, T)$

**Algorithm 46 PrepareRegenInitMolecule:** prepare initial molecule for re-generation

**Input:** molecule $\mathcal{M}$, atom index set of re-generated part $\mathbb{A}_{\text{regen}}$, bond index set of re-generated part $\mathbb{B}_{\text{regen}}$, protein pocket $\mathcal{K}$.

**Output:** initialize molecule for re-generation $\mathcal{M}^{\text{init}} = \{\mathbf{A}^{\text{init}}, \mathbf{X}^{\text{init}}, \mathbf{B}^{\text{init}}\}$.

$\mathbf{A}, \mathbf{X}, \mathbf{B} \leftarrow \mathcal{M}$
$\mathbf{A}_{\text{regen}} = \mathbf{A}, \mathbf{X}_{\text{regen}} = \mathbf{X}, \mathbf{B}_{\text{regen}} = \mathbf{B}$
$N' = \|\mathbf{A}_{\text{regen}}\|$
$\mathbf{X}^{\text{init}}_{\mathbb{A}_{\text{regen}}} = \text{PrepareInitCoordinates}(N', \mathcal{K})$
$\mathbf{A}^{\text{init}}_{\mathbb{A}_{\text{regen}}} = \mathbf{0}_{N' \times K_{\text{atom}}}$
$\mathbf{B}^{\text{init}}_{\mathbb{B}_{\text{regen}}} = \mathbf{0}_{N' \times N' \times K_{\text{bond}}}$
$\mathcal{M}^{\text{init}} = \{\mathbf{A}^{\text{init}}, \mathbf{X}^{\text{init}}, \mathbf{B}^{\text{init}}\}$

**Algorithm 47 DefineRegenNoise:** define molecular noise and noise scale for re-generation

**Input:** molecule $\mathcal{M}$, atom index set of re-generated part $\mathbb{A}_{\text{regen}}$, bond index set of re-generated part $\mathbb{B}_{\text{regen}}$, total step $T$.

**Output:** molecular noise $\xi$, schedule of noise scale $[\beta^1, \beta^2, \dots, \beta^T]$.

$N \leftarrow$ number of atoms of $\mathcal{M}$

$\mathbb{M}_{\text{regen}} = \mathbb{A}_{\text{regen}} \cup \mathbb{B}_{\text{regen}}$

$\mathbb{M}_{\text{others}} = \{i | i \in [\![N]\!]\} \cup \{ij | i, j \in [\![N]\!]\} \setminus \mathbb{M}_{\text{regen}}$

$\xi_{\text{part}} \leftarrow$ define molecular part noise using Eq. S45

$\xi_{\text{regen}} = \xi_{\text{frag}}$, $\xi_{\text{others}} = \xi_{\text{frag}}$

$\xi$ is defined as a mapping: $\xi(\mathbb{M}_k) = \begin{cases} \xi_{\text{others}}, & \mathbb{M}_k = \mathbb{M}_{\text{others}} \\ \xi_{\text{regen}}, & \mathbb{M}_k = \mathbb{M}_{\text{regen}} \end{cases}$

$[{\beta^t}_{\text{others}}] = \Gamma(\beta_{\max} = 0.4, \beta_{\min} = 0, T)$

$[{\beta^t}_{\text{regen}}] = \Upsilon'(\beta_{\max} = 1 - 10^{-5}, \beta_{\min} = 10^{-5}, k = 3, T)$

$\forall t \in [\![T]\!], \beta^t$ is defined as a mapping: $\beta^t(\mathbb{M}_k) = \begin{cases} \beta^t_{\text{others}}, & \mathbb{M}_k = \mathbb{M}_{\text{others}} \\ \beta^t_{\text{regen}}, & \mathbb{M}_k = \mathbb{M}_{\text{regen}} \end{cases}$

Utilizing the confidence scores, we can also employ an auto-regressive-like approach to generate molecules (Algorithm 48). The concept involves generating molecular atoms iteratively, conditioning each generation process on previously generated atoms.

**Algorithm 48 GenSBDDAR:** generation process using confidence scores for SBDD in an auto-regressive-like manner

**Input:** number of molecular atoms $N$, protein pocket $\mathcal{K}$.

**Output:** generated molecule $\mathcal{M}_{\text{gen}}$, confidence scores $\mathcal{S}_{\text{gen}}$.

$\mathcal{P} \leftarrow$ derive task prompt for SBDD using Eq. S12

$T = 30$

$\mathcal{M}^0 = \text{PrepareInitMolecule}(N, \mathcal{K})$

$\mathcal{S}^0 = \varnothing$, $\mathbb{A}^0_{\text{prev}} = \varnothing$

**for all** $\tau \in [\![N]\!]$ **do**

    $\mathcal{M}^\tau_{\text{init}}, \xi^\tau, \boldsymbol{\beta}^\tau, \mathbb{A}^\tau_{\text{prev}} = \text{PrepareAR}(\mathcal{M}^{\tau-1}, \mathcal{S}^{\tau-1}, \mathcal{K}, \mathbb{A}^{\tau-1}_{\text{prev}}, T)$

    **if** $\|\mathbb{A}^\tau_{\text{prev}}\| =$ number of atoms of $\mathcal{M}^\tau_{\text{init}}$ **then**

        $\mathcal{M}^\tau \leftarrow \mathcal{M}^{\tau-1}, \mathcal{S}^\tau \leftarrow \mathcal{S}^{\tau-1}$

        break

    **else**

        $\mathcal{M}^\tau, \mathcal{S}^\tau, \mathcal{S}^\tau_{\text{traj}} = \text{GenerationProcess}(\mathcal{M}^\tau_{\text{init}}, \mathcal{K}, \mathcal{P}, \xi^\tau, \boldsymbol{\beta}^\tau, T)$

    **end if**

**end for**

$\mathcal{M}_{\text{gen}} = \mathcal{M}^\tau, \mathcal{S}_{\text{gen}} = \mathcal{S}^\tau$

**Algorithm 49 PrepareAR:** prepare for auto-regressive-like generation

**Input:** previously generated molecule $\mathcal{M}$, confidence scores $\mathcal{S}$, protein pocket $\mathcal{K}$, atom set $\mathbb{A}_{\text{prev}}$ of previously generated atoms of $\mathcal{M}$, total step $T$.

**Output:** initial molecule $\mathcal{M}_{\text{init}}$ for the next generation round, molecular noise $\xi$, noise scale $\boldsymbol{\beta} = [\beta^1, \beta^2, \ldots, \beta^T]$, atom set $\mathbb{A}'_{\text{prev}}$ of already generated atoms of $\mathcal{M}_{\text{init}}$.

▷ Get generated atoms in this round
$N \leftarrow$ number of atoms of $\mathcal{M}$
$\mathbb{A}_{\text{togen}} = \{i | i \in [\![N]\!]\} \setminus \mathbb{A}_{\text{prev}}$
$N_{\text{add}} = \min(6, \|\mathbb{A}_{\text{togen}}\|)$ ▷ Each round selects at most 6 atoms as newly generated ones
**if** $\mathcal{S} = \varnothing$ **then** ▷ the initial step
    $\mathbb{A}_{\text{add}} = \varnothing$
**else**
    $[s_i^{\text{coor}}] \leftarrow$ confidence scores of atom coordinates from $\mathcal{S}$
    $\forall i \in \mathbb{A}_{\text{togen}}$, $p_i = (s_i^{\text{coor}} - \min \mathbf{s}^{\text{coor}})/(\max \mathbf{s}^{\text{coor}} - \min \mathbf{s}^{\text{coor}} + 0.0001) + 0.001$
    $\mathbb{A}_{\text{add}} \leftarrow$ sample $N_{\text{add}}$ atoms from $\mathbb{A}_{\text{togen}}$ with normalized $p_i$ as probabilities
**end if**
$\mathbb{A}'_{\text{prev}} = \mathbb{A}_{\text{prev}} \cup \mathbb{A}_{\text{add}}$
$\mathbb{A}'_{\text{togen}} = \{i | i \in [\![N]\!]\} \setminus \mathbb{A}'_{\text{prev}}$
$\mathbb{B}'_{\text{togen}} = \{ij | i \in \mathbb{A}'_{\text{togen}}$ or $j \in \mathbb{A}'_{\text{togen}}\}$
▷ Prepare initial molecule
$\mathcal{M}_{\text{init}} =$ PrepareRegenInitMolecule$(\mathcal{M}, \mathbb{A}'_{\text{togen}}, \mathbb{B}'_{\text{togen}})$
▷ Define noise scale
$\xi, \boldsymbol{\beta} =$ DefineRegenNoise$(\mathbb{A}'_{\text{togen}}, \mathbb{B}'_{\text{togen}}, \mathcal{M}_{\text{init}}, T)$

##### 4.3.5 3D molecule generation

The generation process for 3D molecule generation is the same as that for SBDD (Algorithm 44 and Algorithm 48). Since no protein pocket is needed for 3D molecule generation, the pocket is set as $\varnothing$ as input for the generation algorithms.

Now we introduce how to incorporate the prior knowledge of shape into the generation process. The basic idea is to modify the atom coordinates of the denoised molecule so that its atom pairwise distances are close to the pairwise distances of a reference point cloud with a specific shape, as defined in Algorithm 50. In our implementation, the values of distance thresholds $d_{\text{th}}$ were set as 2 or 3.

**Algorithm 50 ModifyShape:** modify the molecule to align with shape

**Input:** molecule $\mathcal{M}$, reference point cloud with specified shape $\mathbf{X}^{\text{ref}}$, generation step $t$, distance threshold $d_{\text{th}}$.

**Output:** modified molecule $\mathcal{M}^{\text{mod}}$.

$t_{\text{mod}} = 90$
**if** $t \leq t_{\text{mod}}$ **then**
  $\mathbf{A}, \mathbf{X}, \mathbf{B} = \mathcal{M}$
  $N \leftarrow$ number of atoms of $\mathcal{M}$
  $\forall i, j \in [\![N]\!], [d_{ij}] = \|\mathbf{x}_i - \mathbf{x}_j^{\text{ref}}\|$
  $\boldsymbol{\eta} \leftarrow$ permutation of $[\![N]\!]$ by solving the assignment problem to minimize $\sum_i d_{i,\eta_i}$
  $\mathbf{X}^{\text{ref}} \leftarrow \mathbf{X}^{\text{ref}}_{\boldsymbol{\eta}}$
  **for all** $i \in [\![N]\!]$ **do**
    $\Delta \mathbf{x}_i = \mathbf{x}_i^{\text{ref}} - \mathbf{x}_i$
    **if** $\|\Delta \mathbf{x}_i\| > d_{\text{th}}$ **then**
      $\mathbf{x}_i \leftarrow \mathbf{x}_i + (t_{\text{mod}} - (t-1))/t_{\text{mod}} \Delta \mathbf{x}_i$
    **end if**
  **end for**
  $\mathcal{M}^{\text{mod}} = \{\mathbf{A}, \mathbf{X}, \mathbf{B}\}$
**else**
  $\mathcal{M}^{\text{mod}} = \mathcal{M}$
**end if**

###### *4.3.6 Fragment linking*

For fragment linking, the molecule is composed of the fragment part and the linker part. The fragment part is provided by the users and the linker part is initialized in a similar way as SBDD. We initialized the molecule composed of the fragment part and the linker part, where the fragment part was provided by the users and the linker part was sampled from the noise distributions

**Algorithm 51 GenFragLinking:** generation process for fragment linking

**Input:** the number $N$ of atoms of the generated molecule, the fragment $\{\mathbf{A}_{\text{frag}}, \mathbf{X}_{\text{frag}}, \mathbf{B}_{\text{frag}}\}$ protein pocket $\mathcal{K}$, task prompt $\mathcal{P}$.

**Output:** generated molecule $\mathcal{M}_{\text{gen}}$, confidence scores $\mathcal{S}_{\text{gen}}$.

$\mathcal{M}_{\text{init}} = \text{PrepareInitMoleculeFrag}(N, \{\mathbf{A}_{\text{frag}}, \mathbf{X}_{\text{frag}}, \mathbf{B}_{\text{frag}}\}, \mathcal{K}, \mathcal{P})$

$\mathcal{P} \leftarrow$ derive task prompt for fragment lining using Eq. S13 - S15

$\xi \leftarrow$ define molecular noise using Eq. S46

$\{\mathbb{M}_{\text{frag}}, \mathbb{M}_{\text{link}}\} \leftarrow$ the domain of $\xi$

$T = 100$

$[{\beta^t}_{\text{frag}}] = \Gamma(\beta_{\max} = 0.3, \beta_{\min} = 0, T)$

$[{\beta^t}_{\text{link}}] = \Upsilon'(\beta_{\max} = 1 - 10^{-5}, \beta_{\min} = 10^{-5}, k = 3, T)$

$\forall t \in [\![T]\!], \beta^t$ is defined as a mapping: $\beta^t(\mathbb{M}_k) = \begin{cases} \beta^t_{\text{frag}}, & \mathbb{M}_k = \mathbb{M}_{\text{frag}} \\ \beta^t_{\text{link}}, & \mathbb{M}_k = \mathbb{M}_{\text{link}} \end{cases}$

$\boldsymbol{\beta} = [\beta^1, \beta^2, \ldots, \beta^T]$

$\mathcal{M}_{\text{gen}}, \mathcal{S}_{\text{gen}}, \mathcal{S}_{\text{traj}} = \text{GenerationProcess}(\mathcal{M}_{\text{init}}, \mathcal{K}, \mathcal{P}, \xi, \boldsymbol{\beta}, T)$

**Algorithm 52 PrepareInitMoleculeFrag:** prepare initial molecule with fragment

**Input:** number of atoms $N$ of the molecule, fragment $\{\mathbf{A}_{\text{frag}}, \mathbf{X}_{\text{frag}}, \mathbf{B}_{\text{frag}}\}$, protein pocket $\mathcal{K}$, task prompt $\mathcal{P}$.

**Output:** initialize molecule $\mathcal{M} = \{\mathbf{A}, \mathbf{X}, \mathbf{B}\}$.

$N_{\text{frag}} \leftarrow$ the number of atoms of the fragment $\mathbf{A}_{\text{frag}}$<br>
$N_{\text{link}} = N - N_{\text{frag}}$<br>
**if** $\mathbf{X}_{\text{frag}} = \varnothing$ **then**<br>
    $\mathbf{X} =$ PrepareInitCoordinates$(N, \mathcal{K})$<br>
**else**<br>
    $\mathbf{X}_{\text{link}} =$ PrepareInitCoordinates$(N_{\text{link}}, \mathcal{K})$<br>
    $\mathbf{X} = \text{concat}(\mathbf{X}_{\text{frag}}, \mathbf{X}_{\text{link}})$<br>
**end if**<br>
$\mathbf{A}_{\text{link}} = \mathbf{0}_{N_{\text{link}} \times K_{\text{atom}}}$<br>
$\mathbf{A} = \text{concat}(\mathbf{A}_{\text{frag}}, \mathbf{A}_{\text{link}})$<br>
$\mathbf{B} = \mathbf{0}_{N \times N \times K_{\text{bond}}}$<br>
$\mathbf{P}^{\text{bond}} \leftarrow$ prompt indicators for bond types from $\mathcal{P}$<br>
**for all** $i, j \in [\![N]\!]$ **do**<br>
    **if** $i \leq N_{\text{frag}}, j \leq N_{\text{frag}}$ **then**<br>
        $\mathbf{b}_{ij} = \mathbf{b}_{ij}^{\text{frag}}$<br>
    **else if** $(i \leq N_{\text{frag}}, j > N_{\text{frag}})$ or $(i > N_{\text{frag}}, j \leq N_{\text{frag}})$ **then** ▷ between fragment and linker<br>
        **if** $P_{ij}^{\text{bond}} = 1$ **then**<br>
            $\mathbf{b}_{ij} \leftarrow$ one-hot encoding of none-bond type<br>
        **end if**<br>
    **end if**<br>
**end for**<br>
$\mathcal{M} = \{\mathbf{A}, \mathbf{X}, \mathbf{B}\}$

##### 4.3.7 PROTAC design

The generation algorithm for PROTAC design is the same as that for fragment linking. In our setting for this task, we assume the protein pockets are not provided. Therefore, the protein pocket is set as $\varnothing$ for the input for the generation algorithm.

##### 4.3.8 Fragment growing

The generation algorithm for fragment growing is similar to that for fragment linking by corresponding the growing part in the fragment growing task to the linker part in the fragment linking task. In our setting of this task, we assumed the fragment poses are completely unknown. Therefore, the coordinates of fragment atoms also require initialization and the noise scale of the fragment part for this task is different from that for fragment linking (Algorithm 53).

**Algorithm 53 GenFragGrowing:** generation process for fragment growing

**Input:** the number $N$ of atoms of the generated molecule, the fragment $\{\mathbf{A}_{\text{frag}}, \mathbf{B}_{\text{frag}}\}$ protein pocket $\mathcal{K}$, task prompt $\mathcal{P}$.

**Output:** generated molecule $\mathcal{M}_{\text{gen}}$, confidence scores $\mathcal{S}_{\text{gen}}$.

$\mathbf{X}_{\text{frag}} = \varnothing$
$\mathcal{M}_{\text{init}} = \text{PrepareInitMoleculeFrag}(N, \{\mathbf{A}_{\text{frag}}, \mathbf{X}_{\text{frag}}, \mathbf{B}_{\text{frag}}\}, \mathcal{K}, \mathcal{P})$
$\mathcal{P} \leftarrow$ derive task prompt for fragment growing using Eq. S16
$\xi \leftarrow$ define molecular noise using Eq. S46
$T = 100$
$\boldsymbol{\beta} = \Upsilon'(\beta_{\max} = 1 - 10^{-5}, \beta_{\min} = 10^{-5}, k = 3, T)$ ▷ all parts use the same noise scale
$\mathcal{M}_{\text{gen}}, \mathcal{S}_{\text{gen}} = \text{GenerationProcess}(\mathcal{M}_{\text{init}}, \mathcal{K}, \mathcal{P}, \xi, \boldsymbol{\beta}, T)$

##### 4.3.9 Molecule optimization

The generation process is similar to the SBDD task. The difference is that the noise scale starts from a value less than 1. In our implementation, we optimized the LogP values of molecules using Algorithm 54.

**Algorithm 54 GenOptLogP:** generation process for optimizing the LogP of molecule

**Input:** reference molecule $\mathcal{M}^{\text{ref}}$, protein pocket $\mathcal{K}$.

**Output:** generated molecule $\mathcal{M}_{\text{gen}}$.

```
∀i ∈ ⟦10⟧, M_i^seed = M^ref
for all round ∈ [1, 2, 3] do
    ▷ Generate molecules
    M = ∅
    for all i ∈ ⟦10⟧ do
        for all j ∈ ⟦10⟧ do
            M_ij, S_ij = GenSimilarMol(M_i^seed, K)
            M ← M ∪ {M_ij}
        end for
    end for
    ▷ Select seeds of next round
    ∀M ∈ M, calculate the LogP values
    [M_1^seed, ..., M_10^seed] ← molecules with top ten best LogP in M (minimum errors to 1.8)
end for
M_gen ← M_1^seed
```

**Algorithm 55 GenSimilarMol:** generation process for generating a similar molecule

**Input:** reference molecule $\mathcal{M}$, protein pocket $\mathcal{K}$.

**Output:** generated molecule $\mathcal{M}_{\text{gen}}$, confidence scores $\mathcal{S}_{\text{gen}}$.

1: $\mathcal{P} \leftarrow$ derive task prompt for SBDD using Eq. S12
2: $\xi \leftarrow$ define molecular noise using Eq. S45
3: $[\beta^1, \beta^2, \ldots, \beta^{100}] = \Upsilon'(\beta_{\max} = 1 - 10^{-5}, \beta_{\min} = 10^{-5}, k = 3, T = 100)$
4: $T = 30$
5: $\boldsymbol{\beta} = [\beta^{71}, \beta^{72}, \ldots, \beta^{100}]$ ▷ only the last several noise scales
6: $\mathcal{M}_{\text{gen}}, \mathcal{S}_{\text{gen}}, \mathcal{S}_{\text{traj}} = \text{GenerationProcess}(\mathcal{M}, \mathcal{K}, \mathcal{P}, \xi, \boldsymbol{\beta}, T)$

###### 4.3.10 De novo peptide design

*De novo* peptide design can be regarded as a special case of fragment growing in our model, i.e., generating the side-chain atoms based on the backbone fragment. However, there is a problem in practice. Given the peptide length $L$, the number of atoms of the backbone fragment is $4L$ but the number of atoms of the side chain is unknown because different amino acid types have different numbers of side-chain atoms. We devised a solution wherein we sample a slightly larger number of atoms, allowing for a margin of flexibility. After the generation process, any atoms decoded as the mask type are discarded. Through this approach, our model autonomously adjusts the quantity of side-chain atoms by predicting atom types. As a result, different types of amino acids could be efficiently generated. The generation algorithm is described in Algorithm 56. Given the peptide length $L$, the number of atoms is sampled from a Gaussian distribution whose parameters are fitted from the dataset (Algorithm 58). Then the backbone fragment atoms and bonds are derived using Algorithm 57. Subsequent steps are similar to those of fragment growing. Finally, the chemical toolbox Open Babel[10] is utilized to identify the amino acid types from the atoms and bonds of the generated molecules. Specifically, the `OBConversion` module of the Open Babel read the generated molecules (peptides) in the `.sdf` files and save them in `.pdb` files. In this way, standard residues will be annotated with their amino acid types in the `.pdb` files.

**Algorithm 56 GenPepDesign:** generation process for *de novo* peptide design

**Input:** peptide length $L$, protein pocket $\mathcal{K}$.

**Output:** generated molecule $\mathcal{M}_{\text{gen}}$, peptide sequence, confidence scores $\mathcal{S}_{\text{gen}}$.

1: $N = \text{GetNumAtomsPep}(L)$
2: $\{\mathbf{A}_{\text{bb}}, \mathbf{B}_{\text{bb}}\} = \text{GetBackboneFrag}(L)$
3: $\mathbf{X}_{\text{bb}} = \varnothing$
4: $\mathcal{M}_{\text{init}} = \text{PrepareInitMoleculeFrag}(N, \{\mathbf{A}_{\text{bb}}, \mathbf{X}_{\text{bb}}, \mathbf{B}_{\text{bb}}\}, \mathcal{K})$
5: $\mathcal{P} \leftarrow$ derive task prompt for *de novo* peptide design using Eq. S17
6: $\xi \leftarrow$ define molecular noise using Eq. S47
7: $T = 100$
8: $\boldsymbol{\beta} = \Upsilon'(\beta_{\max} = 1 - 10^{-5}, \beta_{\min} = 10^{-5}, k = 3, T)$
9: $\mathcal{M}_{\text{gen}}, \mathcal{S}_{\text{gen}}, \mathcal{S}_{\text{traj}} = \text{GenerationProcess}(\mathcal{M}_{\text{init}}, \mathcal{K}, \mathcal{P}, \xi, \boldsymbol{\beta}, T)$
10: peptide sequence $\leftarrow$ Open Babel identifies the amino acid types of $\mathcal{M}_{\text{gen}}$

**Algorithm 57 GetBackboneFrag:** get the backbone fragment for linear peptide generation

**Input:** peptide length $L$.

**Output:** atom types and bond types $\{\mathbf{A}_{\text{bb}}, \mathbf{B}_{\text{bb}}\}$ of the backbone fragment of the peptide.

```
N_bb = 4L
A_bb = [a_i] ← 0_{N_bb × K_atom}
B_bb = [b_ij] ∈ {0,1}^{N_bb × N_bb × K_atom} ← initialize all b_ij as one-hot encoding of none-bond
for all i ∈ ⟦N_bb⟧ do
    if mod (i, 4) = 1 then                                        ▷ nitrogen atom
        a_i ← one-hot encoding of nitrogen
        b_{i,i+1} ← one-hot encoding of single bond               ▷ N-C_α
        if i ≥ 4 then
            b_{i,i-2} ← one-hot encoding of single bond           ▷ peptide bond
        end if
    else if mod (i, 4) = 2 then                                   ▷ alpha carbon atom
        a_i ← one-hot encoding of carbon
        b_{i,i-1} ← one-hot encoding of single bond               ▷ C_α-N
        b_{i,i+1} ← one-hot encoding of single bond               ▷ C_α−C
    else if mod (i, 4) = 3 then                                   ▷ carbon atom
        a_i ← one-hot encoding of carbon
        b_{i,i-1} ← one-hot encoding of single bond               ▷ C−C_α
        b_{i,i+1} ← one-hot encoding of double bond               ▷ C=O
        if i < 4(L − 1) then
            b_{i,i+2} ← one-hot encoding of single bond           ▷ peptide bond
        end if
    else                                                          ▷ oxygen atom
        a_i ← one-hot encoding of oxygen
        b_{i,i-1} ← one-hot encoding of double bond               ▷ O=C
    end if
end for
```

**Algorithm 58 GetNumAtomsPep:** sample the number of atoms for peptide generation

**Input:** peptide length $L$

**Output:** the number $N$ of atoms for peptide generation

```
μ = 8
σ = 0.3817L + 1.8727
N ∼ 𝒩(μ, σ²)
N ← round of N
```

The generation process for cyclic peptide design is similar to that of linear peptides. The only difference lies in the backbone fragments provided: linear backbones are used for linear peptides, while cyclic backbones are used for cyclic peptides. The cyclic backbone can be obtained by just connecting the nitrogen atom of the first residue and the carbon atom of the last residue with a single bond, as shown in Algorithm 59.

**Algorithm 59 GetBackboneFragCyclic:** get the backbone fragment for cyclic peptide generation

**Input:** peptide length $L$.

**Output:** atom types and bond types $\{\mathbf{A}_{\text{bb}}, \mathbf{B}_{\text{bb}}\}$ of the backbone fragment of the cyclic peptide.

1: $\mathbf{A}_{\text{bb}}, \mathbf{B}_{\text{bb}} = \text{GetBackboneFrag}(L)$

2: $\mathbf{b}_{1,4L-1} \leftarrow$ one-hot encoding of single bond ▷ head-to-tail single bond for cycle

The generation process for peptides containing non-standard amino acids is the same as that for linear peptides. Their distinction arises during post-generation processing, where peptides are selected based on whether their side chains can be mapped to standard amino acids.

###### 4.3.11 Peptide inverse folding

The generation process for peptide inverse folding is similar to that for *de novo* peptide design. The only difference is that the atom coordinates of the backbones are known in peptide inverse folding.

**Algorithm 60 GenPepInvFolding:** generation process for peptide inverse folding

**Input:** peptide length $L$, the backbone fragment $\{\mathbf{A}_{\text{bb}}, \mathbf{X}_{\text{bb}}, \mathbf{B}_{\text{bb}}\}$, protein pocket $\mathcal{K}$.

**Output:** generated molecule $\mathcal{M}_{\text{gen}}$, peptide sequence, confidence scores $\mathcal{S}_{\text{gen}}$.

1: $N = \text{GetNumAtomsPep}(L)$

2: $\mathcal{M}_{\text{init}} = \text{PrepareInitMoleculeFrag}(N, \{\mathbf{A}_{\text{bb}}, \mathbf{X}_{\text{bb}}, \mathbf{B}_{\text{bb}}\}, \mathcal{K})$

3: $\mathcal{P} \leftarrow$ derive task prompt for peptide inverse folding using Eq. S18

4: $\xi \leftarrow$ define molecular noise using Eq. S47

5: $T = 100$

6: $\boldsymbol{\beta} = \Upsilon'(\beta_{\text{max}} = 1 - 10^{-5}, \beta_{\text{min}} = 10^{-5}, k = 3, T)$

7: $\mathcal{M}_{\text{gen}}, \mathcal{S}_{\text{gen}}, \mathcal{S}_{\text{traj}} = \text{GenerationProcess}(\mathcal{M}_{\text{init}}, \mathcal{K}, \mathcal{P}, \xi, \boldsymbol{\beta}, T)$

8: peptide sequence $\leftarrow$ Open Babel identifies the amino acid types of $\mathcal{M}_{\text{gen}}$

##### 4.4 Generation with variant pocket structures

The generation process remains unchanged when using variant pocket structures compared to using experimentally determined pocket structures. When multiple candidate pocket structures are available for a protein, for instance, when AlphaFold provides a set of predicted conformations for one protein, we retain all these structural candidates rather than selecting a single structure. For each generation attempt, one pocket structure is randomly sampled from the set of candidates.

##### 4.5 Implementation availability

We have released the source code of the model at `https://github.com/pengxingang/PocketXMol`. To facilitate quick adoption and provide practical demonstrations, we also offer interactive Colab notebooks at https://github.com/pengxingang/PocketXMol/tree/master/notebooks. These notebooks streamline the use of PocketXMol for sampling on user-provided data. Users can simply follow the step-by-step instructions to upload their data and specify generation requirements. The notebooks then automatically install necessary dependencies, generate the configuration file, execute the sampling process, and display the results.

### 5 Model training

The denoiser was optimized to recover the molecules from the noisy ones, with randomly sampled molecules, tasks, and noise scales at each training step (Algorithm 61 and Figure S6). At each training step, we randomly sampled a training task from the training task set and constructed the corresponding task prompt and the molecular noise (Supplementary Notes 5.1). Then we randomly sampled a data point from the training dataset (Supplementary Notes 6.2), including the molecule and possibly the binding protein pocket. Next, we sampled a noise scale from the interval $(0, 1)$ and then perturbed the molecule. Subsequently, the noisy molecule was fed into the denoiser to predict the denoised molecule, which was used to calculate the loss function (Supplementary Notes 5.2) for parameter optimization. The training tasks include the typical generative tasks introduced in the previous sections, as well as more general formulated tasks. The setup of the training task enables the model to generalize to new generative tasks it has never considered in the training phase. Therefore, the training task often employs more complicated perturbations than that of the generative tasks used for evaluation.

**Algorithm 61 ModelTraining:** train the denoiser neural networks

```
while not converge do
    training task ← sample a task from all training tasks (Supplementary Notes 5.1)
    {M, K} ← sample a data point from training dataset for the task
    P, ξ ← define the task prompt and noise distributions for the molecule
    β ← sample a noise scale from (0, 1)
    M̃ = Φ(M, P, ξ, β)                                  ▷ add noise
    M̂, Ŝ = F_Θ^raw(M̃, P, K)                            ▷ denoise (un-adapted outputs, Eq. S1)
    L = L(M, M̂, Ŝ, P)
    Θ ← optimize parameter Θ to minimize L
end while
```

#### 5.1 Training tasks

We defined seven training tasks including 3D molecule generation, molecular conformation generation, structure-based drug design (SBDD), molecular docking, mask-fill, fragment-based drug design (FBDD), and peptide design. These training tasks can cover all generative tasks in practice. The first four training tasks (3D molecule generation, molecular conformation generation, SBDD, and docking) adopt the same task prompt, molecular noise, and perturbation procedure with their corresponding generative tasks, and the noise scale is uniformly sampled from $(0, 1)$ during training.

For molecular docking and 3D molecular conformation tasks, whether using the Gaussian noise or the flexible noise is also randomly sampled during training, with the probabilities of 0.999 for

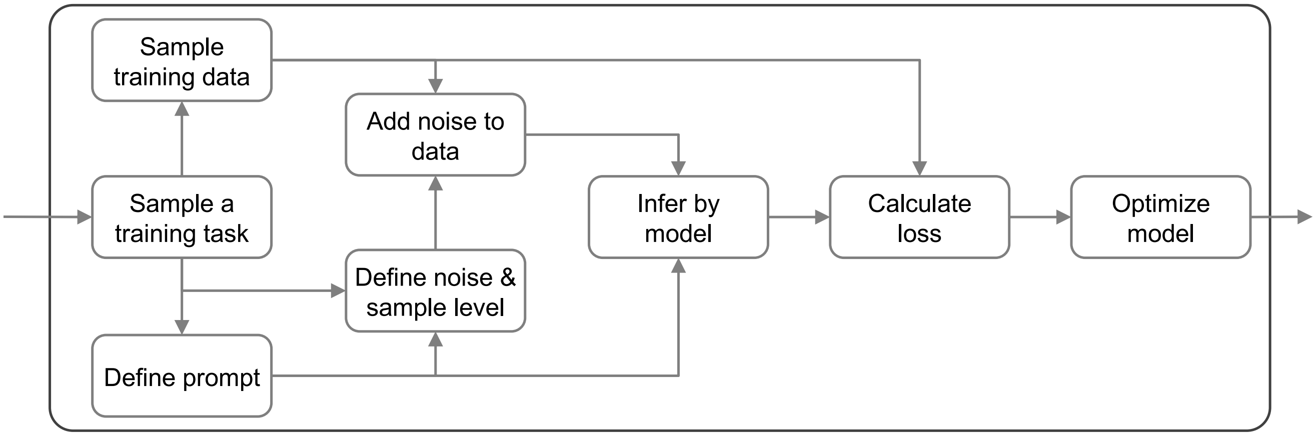

**Figure S6. A schematic diagram demonstrating one training step of the training loop.**

Gaussian noise and 0.001 for flexible noise. Additionally, in the training task of 3D conformation generation, we have addressed an issue in the denoised molecular generation. Due to the presence of molecular symmetry, the optimal strategy for restoring several perturbed atoms may not necessarily involve returning an atom to its original position. For example, to rapidly recover a benzene ring in 3D space, the model should learn to restore atoms to positions symmetrical to their counterparts, rather than to their original locations. Thus, we traversed all possible atom permutation schemes that did not alter the molecular graph topology and selected the one that minimized the distance between the corresponding molecules before and after perturbation.

The mask-fill task is a general version of tasks including fragment linking and PROTAC design. To achieve this task, we divide the molecule into two parts $\mathbb{M}_1$ and $\mathbb{M}_2$. $\mathbb{M}_1$ includes a set of atoms and all the inner bonds, and $\mathbb{M}_2$ includes all the remaining atoms and bonds. The task prompt is defined similarly to the task of fragment linking (Eqs. S13-S15), where the fragment part and the linker part correspond to $\mathbb{M}_1$ and $\mathbb{M}_2$, respectively. The molecular noise $\xi$ is defined as:

$$\xi(\mathbb{M}_k) = \begin{cases} \{\bar{\mathbf{a}}_1, \bar{\mathbf{b}}_1, \sigma_1^{\text{coor}}, \sigma_1^{\text{trans}}, \epsilon, \kappa\}, & \mathbb{M}_k = \mathbb{M}_1 \\ \{\bar{\mathbf{a}}_2, \bar{\mathbf{b}}_2, \sigma_2\}, & \mathbb{M}_k = \mathbb{M}_2 \end{cases} \tag{S53}$$

where $\{\bar{\mathbf{a}}_1, \bar{\mathbf{b}}_1, \sigma_1^{\text{coor}}\}$ and $\{\bar{\mathbf{a}}_2, \bar{\mathbf{b}}_2, \sigma_2\}$ share the same meaning and values as the molecular noise of SBDD (Eq. S45), and $\{\sigma_1^{\text{trans}}, \epsilon, \kappa\}$ share the same meaning and values as the molecular noise of docking using flexible noise (Eq. S42). The noise scale is a mapping function defined as the follows,

$$\beta(\mathbb{M}_k) = \begin{cases} 0.3\bar{\beta}, & \mathbb{M}_k = \mathbb{M}_1 \\ \bar{\beta}, & \mathbb{M}_k = \mathbb{M}_2 \end{cases} \tag{S54}$$

where $\bar{\beta}$ is uniformly sampled from $(0, 1)$.

In the training phase, we employed different strategies at each stage of the mask-fill task to ensure that the model could generalize better according to the diverse needs of users. Whether each strategy was adopted was independently decided at each training step. These four strategies are described as follows,

1. **Decomposition strategy.** To improve the generalizability, we need a proper strategy to determine how to decompose the molecule. Our strategy is to first decompose the molecule into

multiple basic units, and subsequently separate these units into two parts $\mathbb{M}_1$ and $\mathbb{M}_2$. At this step, we developed three options: BRICS decomposition, MMPA decomposition, and atomic-level decomposition. The BRICS decomposition decomposed the molecule into molecular fragments using the BRICS decomposition algorithm[11], and the MMPA decomposition used an algorithm based on matched molecular pair analysis (MMPA)[12]. Both approaches are implemented using RDKit[13]. The atomic-level decomposition uses the atom as the basic decomposed unit. During training, these three decomposition methodologies were sampled with probabilities of 0.2, 0.3, and 0.5, respectively.

2. **Assembly strategy.** The assembly strategy introduces how we defined $\mathbb{M}_1$ and $\mathbb{M}_2$ based on the basic units from the decomposition. It includes three options: tree assembly, inverse tree assembly, and random assembly. For the tree assembly, we randomly selected a node as the root of the tree, then constructed a spanning tree starting from the root node using a breadth-first search for the graph. Next, we traversed the spanning tree in preorder, randomly cut the path of all the visited nodes in the middle, and chose the atoms of the part closer to the root node as $\mathbb{M}_1$, and the remaining atoms as $\mathbb{M}_2$. For the inverse tree assembly, we took the part close to the root as $\mathbb{M}_2$ and the other one as $\mathbb{M}_1$. The random assembly does not build any tree structure but randomly selects a set of nodes (fragments or atoms) as $\mathbb{M}_1$ and the remaining part as $\mathbb{M}_2$. In the training phase, these three assembly strategies were sampled with probabilities of 0.35, 0.3, and 0.35, respectively.

3. **Part-one perturbation strategy.** We define five different noise types to perturb $\mathbb{M}_1$, including 1) Gaussian noise for atom coordinates; 2) noise used in the task of SBDD; 3) coordinate noise using flexible noise without injected to the rotation of rotatable bonds; 4) coordinate noise with flexible noise; 5) no noise introduced to $\mathbb{M}_1$. The transformation between different types of noise can be achieved by removing the corresponding noise parameters of $\mathbb{M}_1$ in Eq. S53. During training, these different strategies were sampled with probabilities of 0.4, 0.4, 0.198, 0.001, and 0.001, respectively.

4. **Anchor strategy.** This strategy controls whether the atoms in $\mathbb{M}_1$ that connect with $\mathbb{M}_2$, which are defined as anchor atoms, are known or not, which includes three options: 1) knowing all connecting atoms; 2) knowing connecting atoms of part of the fragments, and 3) knowing none of the connecting atoms. If the connecting atoms are provided beforehand, the bond types between the corresponding non-connecting atoms and all atoms in $\mathbb{M}_2$ are set as none-bond and the corresponding prompt indicators for bond types are set as fixed. During training, the options are sampled with probabilities of 0.15, 0.2, and 0.65, respectively.

The FBDD training task is an extension of the mask-fill task. The FBDD task also divides the molecule into two parts $\mathbb{M}_1$ and $\mathbb{M}_2$. The differences between these two tasks are their noise scales and noise distributions introduced to $\mathbb{M}_1$. Here, the noise scale of $\mathbb{M}_1$ is equal to that of $\mathbb{M}_2$, i.e., uniformly sampled from $(0, 1)$. The noise of $\mathbb{M}_1$ does not perturb the atom types or bond types, i.e., $\xi(\mathbb{M}_1) = \{\sigma_1^{\text{coor}}, \sigma_1^{\text{trans}}, \epsilon, \kappa\}$. The FBDD task has the same four strategies as the mask-fill task but with different probability distributions. The probabilities for the BRICS decomposition, MMPA decomposition, and atomic-level decomposition are 0.6, 0.4, and 0, respectively. For the assembly strategies, the probabilities for tree, inverse tree, and random are 0.15, 0.35, and 0.5, respectively. For the perturbation strategy, the probabilities for the three options discussed above are 0, 0.998,

0, 0.001, and 0.001, respectively. For the anchor strategy, the probabilities for the three options are 0.25, 0.25, and 0.5, respectively.

For this training task, the molecular noise is the same as the generative task of *de novo* peptide design (Eq. S47), and the noise scale is uniformly sampled from $(0, 1)$. This training task has three sub-tasks. The first two tasks correspond to the generative tasks of *de novo* peptide design (Eq. S17) and peptide inverse folding (Eq. S18). The third sub-task is side-chain packing in which atom and bond types of side-chains are fixed and atom coordinates are to be predicted. These three sub-tasks were sampled with probabilities of 0.7, 0.2, and 0.1, respectively. Moreover, dummy atoms annotated as the mask type were introduced to the molecules before perturbation. For a peptide with $N_{\text{sc}}$ atoms, the number of dummy atoms had half probability of being zero, and other half to be sampled from the uniform distribution with range 0 to $0.1N_{\text{sc}}$. Then we sampled an equal number of side-chain atoms as center atoms and sampled vectors with lengths following Gaussian distribution (mean as 1.4 Å and standard deviation as 0.2Å) as the relative between the center atoms to the dummy atoms. The coordinates of the dummy atoms are determined by adding the relative vectors to the coordinates of the center atoms. Their atom types are assigned as the mask type and their bond types with other atoms are assigned as none-bond. Then these dummy atoms are added to the side-chain part and treated in the same way as other side-chain atoms.

During training, the training tasks were sampled with probabilities of 0.14 for 3D molecule generation, 0.12 for molecular conformation generation, 0.205 for SBDD, 0.25 for docking, 0.11 for mask-fill, 0.045 for FBDD, and 0.13 for peptide design.

#### 5.2 Training loss

The loss for a denoised molecule was calculated using Algorithm 62. It requires the un-adapted denoised molecule $\hat{\mathcal{M}}^{\text{raw}} = \{\mathbf{A}^{\text{raw}}, \mathbf{X}^{\text{raw}}, \mathbf{B}^{\text{raw}}\}$ (i.e., not processed by the M-Projector, Eq. S1) to calculate the noise. The items in $\mathbf{A}^{\text{raw}}$ and $\mathbf{B}^{\text{raw}}$ are the predicted probabilities of the atom and bond types instead of the one-hot encoding vectors. The loss function consists of six terms, including atom types, atom coordinates, and bond types, distances between two atoms, dihedral angles, and the loss for predicted confidence scores.

---

**Algorithm 62 CalcLoss:** calculate the loss for a denoised molecule

---

**Input:** un-adapted denoised molecule $\hat{\mathcal{M}}^{\text{raw}}$, predicted confidence scores $\mathcal{S}$, true (clean) molecule $\mathcal{M}$, task prompt $\mathcal{P}$.

**Output:** loss $L$ for the molecule.

$\hat{\mathbf{A}}, \hat{\mathbf{X}}, \hat{\mathbf{B}} \leftarrow \hat{\mathcal{M}}^{\text{raw}}$ ▷ omit the superscript raw for simplicity

$\mathbf{A}, \mathbf{X}, \mathbf{B} \leftarrow \mathcal{M}$

$N \leftarrow$ the number of atoms of $\mathcal{M}$

$L^{\text{atom}} = \text{CalcAtomLoss}(\hat{\mathbf{A}}, \mathbf{A}, \mathbf{P}^{\text{atom}})$

$L^{\text{coor}} = \text{CalcCoordinateLoss}(\hat{\mathbf{X}}, \mathbf{X}, \mathbf{P}^{\text{coor}})$

$L^{\text{bond}} = \text{CalcBondLoss}(\hat{\mathbf{B}}, \mathbf{B}, \mathbf{P}^{\text{bond}})$

$L^{\text{dist}} = \text{CalcDistanceLoss}(\hat{\mathbf{X}}, \mathbf{X}, \mathbf{P}^{\text{dist}})$

$L^{\text{dih}} = \text{CalcDihedralLoss}(\hat{\mathbf{X}}, \mathbf{X}, \mathbf{P}^{\text{dist}}, \mathcal{M})$

$L^{\text{cfd}} = \text{CalcConfidenceLoss}(\hat{\mathcal{M}}^{\text{raw}}, \hat{\mathcal{S}}, \mathcal{M}, \mathcal{P})$

$L = 1.5L^{\text{atom}} + 2.5L^{\text{coor}} + 1.5L^{\text{bond}} + 0.0005L^{\text{dist}} + 0.0005L^{\text{dih}} + L^{\text{cfd}}$

---

**Algorithm 63 CalcAtomLoss:** calculate the loss for predicted atom types

**Input:** predicted probabilities of atom types $\hat{\mathbf{A}}$, true atom types $\mathbf{A}$, prompt indicators for atom types $\mathbf{P}^{\text{atom}}$.

**Output:** loss for predicted atom types $L^{\text{atom}}$.

1: $N \leftarrow$ the number of atoms of $\mathbf{A}$
2: **for all** $i \in [\![N]\!]$ **do**
3: $\quad w_i^{\text{atom}} = (1 - P_i^{\text{atom}}) + 1/15 P_i^{\text{atom}}$
4: $\quad l_i^{\text{atom}} = w_i^{\text{atom}} \times \text{CrossEntropyLoss}(\tilde{\mathbf{a}}_i, \mathbf{a}_i)$
5: **end for**
6: $L^{\text{atom}} = 1/N \sum_i l_i^{\text{atom}}$

**Algorithm 64 CalcCoordinateLoss:** calculate the loss for predicted atom coordinates

**Input:** predicted atom coordinates $\hat{\mathbf{X}}$, true atom coordinates $\mathbf{X}$, prompt indicators for atom coordinate $\mathbf{P}^{\text{coor}}$.

**Output:** loss for predicted atom coordinates $L^{\text{coor}}$.

1: $N \leftarrow$ the number of atoms of $\mathbf{X}$
2: **for all** $i \in [\![N]\!]$ **do**
3: $\quad w_i^{\text{coor}} = (1 - P_i^{\text{coor}}) + 1/25 P_i^{\text{coor}}$
4: $\quad l_i^{\text{coor}} = w_i^{\text{coor}} \times \|\hat{\mathbf{x}}_i - \mathbf{x}_i\|^2$
5: **end for**
6: $L^{\text{coor}} = 1/N \sum_i l_i^{\text{coor}}$

**Algorithm 65 CalcBondLoss:** calculate the loss for predicted bond types

**Input:** predicted probabilities of bond types $\hat{\mathbf{B}}$, true atom types $\mathbf{B}$, prompt indicators for bond types $\mathbf{P}^{\text{bond}}$.

**Output:** loss for predicted bond types $L^{\text{bond}}$.

1: **for all** $i, j \in [\![N]\!]$ **do**
2: $\quad w_{ij}^{\text{bond}} = (1 - P_{ij}^{\text{bond}}) + 1/15 P_{ij}^{\text{bond}}$
3: $\quad l_{ij}^{\text{bond}} = w_{ij}^{\text{bond}} \times \text{CrossEntropyLoss}(\tilde{\mathbf{b}}_{ij}, \mathbf{b}_{ij})$
4: **end for**
5: $L^{\text{bond}} = 1/N^2 \sum_{ij} l_{ij}^{\text{bond}}$

**Algorithm 66 CalcDistanceLoss:** calculate the loss for predicted atom distances

**Input:** predicted atom coordinates $\hat{\mathbf{X}}$, true atom coordinates $\mathbf{X}$, prompt indicators for atom coordinate $\mathbf{P}^{\text{dist}}$.

**Output:** loss for predicted atom distances $L^{\text{dist}}$.

**for all** $i, j \in [\![N]\!]$ **do**

$\quad w_{ij}^{\text{dist}} = \begin{cases} 1, & \exists k, P_{ik}^{\text{dist}} = 1 \text{ or } P_{kj}^{\text{dist}} = 1 \\ 0, & \text{otherwise} \end{cases}$

$\quad \hat{d}_{ij} = \|\hat{\mathbf{x}}_i - \hat{\mathbf{x}}_j\|$

$\quad d_{ij} = \|\mathbf{x}_i - \mathbf{x}_j\|$

$\quad l_{ij}^{\text{dist}} = \|\hat{d}_{ij} - d_{ij}\|^2$

**end for**

$L^{\text{dist}} = \left(\sum_{ij} l_{ij}^{\text{dist}}\right) / \left(\sum_{ij} w_{ij}^{\text{dist}}\right)$

**Algorithm 67 CalcDihedralLoss:** calculate the loss for predicted dihedral angles

**Input:** predicted atom coordinates $\hat{\mathbf{X}}$, true atom coordinates $\mathbf{X}$, prompt indicators for atom coordinate $\mathbf{P}^{\text{dist}}$, true molecule $\mathcal{M}$.

**Output:** loss for predicted dihedral angles $L^{\text{dih}}$.

$\Omega = \text{GetFlexDihedralAngleIndices}(\mathcal{M}, \mathbf{P}^{\text{dist}})$

**for all** $kijm \in \Omega$ **do**

$\quad \sin\hat{\psi}_{kijm}, \cos\hat{\psi}_{kijm} = \text{CalcDihedralAngle}(\hat{\mathbf{x}}_k, \hat{\mathbf{x}}_i, \hat{\mathbf{x}}_j, \hat{\mathbf{x}}_l)$

$\quad \sin\psi_{kijm}, \cos\psi_{kijm} = \text{CalcDihedralAngle}(\mathbf{x}_k, \mathbf{x}_i, \mathbf{x}_j, \mathbf{x}_l)$

$\quad \cos(\hat{\psi}_{kijm} - \psi_{kijm}) = \cos\hat{\psi}_{kijm}\cos\psi_{kijm} + \sin\hat{\psi}_{kijm}\sin\psi_{kijm}$

$\quad l_{kijm}^{\text{dih}} = 1 - \cos(\hat{\psi}_{kijm} - \psi_{kijm})$

**end for**

$L^{\text{dih}} = 1/\|\Omega\| \sum_{kijm} l_{kijm}^{\text{dih}}$

**Algorithm 68 GetFlexDihedralAngleIndices**: get the indices of the active dihedral angles in the flexible noise

**Input:** molecule $\mathcal{M}$, prompt indicators for distances $\mathbf{P}^{\text{dist}}$.

**Output:** index set $\Omega$ of active dihedral angles.

$\{\mathbb{A}_u | u \in [\![n_{\text{groups}}]\!]\} \leftarrow$ flexible groups from $\mathbf{P}^{\text{dist}}$

$\mathbf{A}, \mathbf{X}, \mathbf{B} \leftarrow \mathcal{M}$

$\Omega = \varnothing$ ▷ index

**for all** $u \in [\![n_{\text{groups}}]\!]$ **do**

  $\mathbb{B}_u = \{ij | i \in \mathbb{A}_u \text{ and } j \in \mathbb{A}_u\}$

  $\mathbb{B}_u^{\text{chem}}, \mathbb{B}_u^{\text{rot}} \leftarrow$ bond set of chemical bonds and rotatable bonds of $\{\mathbf{A}_{\mathbb{A}_u}, \mathbf{B}_{\mathbb{B}_u}\}$

  **for all** $ij \in \mathbb{B}_u^{\text{rot}}$ **do**

    **for all** $ki \in \mathbb{B}_u^{\text{chem}} \setminus \{ji\}, jl \in \mathbb{B}_u^{\text{chem}} \setminus \{ji\}$ **do**

      $\Omega \leftarrow \Omega \cup \{kijm\}$

    **end for**

  **end for**

**end for**

**Algorithm 69 CalcConfidenceLoss:** calculate the loss for predicted confidence scores

**Input:** un-adapted denoised molecule $\hat{\mathcal{M}}^{\text{raw}}$, predicted confidence scores $\hat{\mathcal{S}}$, true molecule $\mathcal{M}$, task prompt $\mathcal{P}$.

**Output:** loss $L^{\text{cfd}}$ for predicted confidence scores.

$\hat{\mathbf{A}}, \hat{\mathbf{X}}, \hat{\mathbf{B}} \leftarrow \hat{\mathcal{M}}^{\text{raw}}$ ▷ Omit superscript
$\mathbf{A}, \mathbf{X}, \mathbf{B} \leftarrow \mathcal{M}$
$N \leftarrow$ the number of atoms of $\mathcal{M}$
▷ Make labels
$\hat{\mathbf{A}}, \hat{\mathbf{B}} \leftarrow$ get one-hot encoding of the types with max probabilities from $\hat{\mathbf{A}}, \hat{\mathbf{B}}$
**for all** $i \in [\![N]\!]$ **do**
    $s_i^{\text{atom}} = \mathbb{I}(\hat{\mathbf{a}}_i = \mathbf{a}_i)$ ▷ $\mathbb{I}$ is the indicator function
    $s_i^{\text{coor}} = \exp\left(\|\hat{\mathbf{x}}_i - \mathbf{x}_i\| \ln 0.2\right)$
    **for all** $j \in [\![N]\!]$ **do**
        $s_{ij}^{\text{bond}} = \mathbb{I}(\hat{\mathbf{b}}_{ij} = \mathbf{b}_{ij})$
    **end for**
**end for**
▷ Calculate loss
$\hat{\mathbf{S}}^{\text{atom}}, \hat{\mathbf{S}}^{\text{coor}}, \hat{\mathbf{S}}^{\text{bond}} \leftarrow \hat{\mathcal{S}}$
**for all** $i \in [\![N]\!]$ **do**
    $l_i^{\text{atom}} = -[s_i^{\text{atom}} \log \hat{s}_i^{\text{atom}} + (1 - s_i^{\text{atom}}) \log(1 - \hat{s}_i^{\text{atom}})]$ ▷ binary cross entropy
    $l_i^{\text{coor}} = \|\hat{s}_i^{\text{atom}} - s_i^{\text{atom}}\|^2$ ▷ squared error
    **for all** $j \in [\![N]\!]$ **do**
        $l_{ij}^{\text{bond}} = -[s_{ij}^{\text{bond}} \log \hat{s}_{ij}^{\text{bond}} + (1 - s_{ij}^{\text{bond}}) \log(1 - \hat{s}_{ij}^{\text{bond}})]$ ▷ binary cross entropy
    **end for**
**end for**
$L^{\text{cfd}} = 1/N \sum_i l_i^{\text{atom}} + 20/N \sum_i l_i^{\text{coor}} + 1/N^2 \sum_{ij} l_{ij}^{\text{bond}}$

#### 5.3 Training implementation

The entire project was implemented using PyTorch 2.0.1 and PyTorch-Lightning 2.0.4. We trained the model with a batch size (per GPU) of 40 on eight 80G A100 GPUs for 180,000 steps (around 54 hours). The optimization was performed using AdamW[14] with weight decay $\lambda = 0.001$, $\beta_1 = 0.99$ and $\beta_2 = 0.999$. The learning rate warmed up linearly from 0 to 0.001 at the first 1000 steps. For designing molecules for wet lab validation, we incorporated more data (including more test set redundancy) for training the model with the same hyperparameters.

We did not extensively explore the hyperparameter space due to the large number of hyperparameters involved. During model development, we found that the model was not highly sensitive to the choice of these hyperparameters. For example, learning rates ranging from 1e-4 to 1e-3 allowed the model to converge with similar training and validation losses. We selected reasonable hyperparameter values and accepted them as long as the training and validation losses showed reasonable reductions.

##### 5.4 Training of PocketXMol-PF

PocketXMol-PF (PocketXMol with pocket-flexible adaptation) was developed by fine-tuning the original PocketXMol on perturbed pocket structures to enhance its robustness to pocket variability. The perturbed pockets were obtained by repacking the side chains of experimentally determined pocket structures using Rosetta, resulting in five alternative conformations per pocket. The reason that we did not use other pocket variants for fine-tuning was that repacking perturbations were easier and more efficient to be obtained, and subsequent experiments showed that using only these perturbations was sufficient for generalizing to different structure variants. During fine-tuning, a pocket structure was sampled from the perturbed set with a probability of 90%, and from the original true pocket with a probability of 10%. If the perturbed structure was chosen, one of the five repacked variants was selected at random. The fine-tuning was conducted using only the docking tasks and associated training data. The model was trained for 20,000 steps with a learning rate of $1 \times 10^{-4}$, while all other hyperparameters were kept consistent with the original PocketXMol configuration.

#### 6 Evaluation settings and data

Before we describe the evaluation setting and the training and testing data, we want to emphasize that the training data that are redundant to ANY test data were removed. For example, although the test set of the SBDD task is a subset of the CrossDocked2020 dataset, any training data that share similar sequences with the test data, even those training data not from the CrossDocked2020 dataset, were removed. See Supplementary Notes 6.2 for more details.

##### 6.1 Benchmark settings for each task

Here we described the benchmark settings for each generative task used to evaluate the performance of different computational approaches. The basic principle is that we try to follow the benchmark settings used in previous work of corresponding tasks unless the previous settings are not suitable.

###### Small molecule docking

- **Data**: We used the test dataset proposed in PoseBusters[15] (Version 1), which contains 428 protein-small-molecule complexes.
- **Protocols**: We also followed the benchmark settings proposed by PoseBusters. Each model provided one docked pose for each pocket to measure the RMSD between the predicted molecular pose and the true one. The performance was measured by the RMSD between the predicted docking pose and the true one in the test set with aligned protein pockets. Note that energy minimization was not used for all models in our evaluation. The values of the docking RMSDs for the baseline models EquiBind, TankBind, DeepDock, DiffDOck, Gold, and Vina were directly adopted from the article of PoseBusters. The values of Uni-Mol Docking V2[16] and SurfDock, RFAA[17] and AlphaFold 3[6] were taken from their corresponding papers[6,16–18]. The PB-valid check was conducted using the PoseBusters checker (`https://posebusters.readthedocs.io/en/latest/`).

###### Molecular conformation generation

- **Data**: We used the test dataset from the work Uni-Mol[19], which contains 200 molecules with multiple conformations for each molecule.

- **Protocols**: Each method generated twice the number of 3D conformations provided by the benchmark dataset for each molecule. The metrics (coverage and matching) were calculated for each molecule, following the same procedure as Uni-Mol. The values of metrics of baselines, except the torsional diffusion series, were also adopted in the paper of Uni-Mol. We installed the torsional diffusion methods by ourselves and produced the values in the test set. The reason is that the original test set used by the torsional diffusion series was different from the test set used in our evaluation. One molecule in the test set contains the element Si that cannot be processed by our model and thus was ignored.

##### Linear peptide docking

- **Data**: We collected the test set proposed in the work of protein-peptide docking[20], which originally contained 122 protein-peptide complexes. We then removed those complexes that did not exist in the PepBDB database[21] because those might not be related to peptide-mediated protein interactions, which resulted in a test set with 79 protein-peptide complexes. For the Q-BioLiP peptide docking test set, it was constructed from protein-peptide pairs released later than 2022-01-01 in the Q-BioLiP[22] database. To ensure no data leakage, we used MMseqs2[23] to compute receptor sequence similarity between the Q-BioLiP database and PocketXMol's training set, and excluded any entries with over 30% sequence identity, resulting in 112 protein-peptide pairs.

- **Protocols**: Each model provided one or multiple docked peptide poses for each protein pocket, and we calculated the DockQ[24] between the predicted poses and the one solved by experimental procedure. For the baseline method FlexPepDock, we used its web server with default settings to dock the peptide. The server provided ten docking poses for each protein pocket. For each protein-peptide pair, the baseline model AlphaFold-Linker added 200 Glycines between the protein and the peptide to form a single chain as the input of AlphaFold 2.3.2[25,26]. It predicted five protein-peptide complexes and adopted the one with the largest pLDDT by AlphaFold as the final prediction. For the baseline AlphaFold-Multiple, we installed it and directly used AlphaFold-Multimer 2.3.2 to predict 25 protein-peptide complexes and the first one ranked by AlphaFold-Multimer was selected as the final prediction. For the baseline AlphaFold 3, we used the server (https://alphafoldserver.com) to predict the protein-peptide complexes and the first one ranked by AlphaFold 3 was selected as the final prediction. For PocketXMol, we generated 100 peptide poses per pocket, ranked them using self-ranking scores or tuned ranking scores, and selected the top-ranked peptide as the final prediction. In the main text, we compared the performance of these ranking methods and then used the tuned ranking as the default ranking strategy for other analyses.

##### Cyclic peptide docking

- **Data**: We began with 34 protein–cyclic peptide complexes collected from a previous study[27]. Peptides longer than 30 amino acids or those whose binding proteins shared identical sequences with any protein in the training set were subsequently excluded. This filtering process yielded a final test set of 26 protein–cyclic peptide complexes. Next, to mitigate the potential dominance of these similar targets in the evaluation, we used MMseqs2 to assess sequence similarity among the test receptors and clustered them into nine distinct groups.

- **Protocols**: Each model provided one docked peptide pose for each protein pocket, and we calculated the DockQ[24] between the predicted poses and the one solved by experimental procedure. For the results of AlphaFold 2 and HighFold, we run the implementation of the HighFold paper[27] with default parameters. For the results of AfCycDesign, we run the implementation of the AfCycDesign paper[28] with default parameters.

##### Structure-based drug design

- **Data**: We used a standard test set widely adopted in previous SBDD works[8,29–32], which contained 100 protein-small-molecule complexes and was a subset of the CrossDocked2020 dataset[33].
- **Protocols**: Each model generated 100 molecules for each protein pocket. The generated molecules were evaluated using metrics proposed by previous work related to molecular generation[5,8]. For the baseline IPDiff and AliDiff, we installed them and generated molecules by ourselves. For other baselines, we used the generated molecules provided by previous works[8,32] for evaluation.

The metric definitions are as follows:

- QED (Quantitative Estimation of Drug-likeness). This metric quantifies the drug-likeness of a molecule based on the distributions of multiple molecular properties ranging from 0 to 1[34]. QED was calculated using the RDKit[13] implementation.
- SA (Synthetic Accessibility). It estimates the ease of synthesizing a molecule, and was calculated based on fragment contributions and complexity penalties[35]. It provides practical guidance for selecting molecules for synthesis and wet-lab validation. We employed the implementation from previous work[30], which normalized the SA score between 0 and 1 and higher values indicate easier synthesis.
- LogP. LogP represents the octanol-water partition coefficient. It is a crucial drug property indicating a molecule's lipophilicity[36]. It was quantified using the Jensen-Shannon Divergence (JSD, derived from KL-divergence) between the distribution of LogP values (predicted by RDKit) of generated molecules and those in the test set. A lower JSD indicates greater similarity between the LogP distributions of generated and real molecules, helping detect potential biases of the lipophilicity of generated molecules.
- Lipinski. Lipinski's Rule of Five provides empirical guidelines for drug-likeness[37]. We used RDKit to count the number of Lipinski rules satisfied by each molecule. While Lipinski's Rule of Five was initially intended to be strictly satisfied in all aspects, subsequent studies have shown that many drugs do not fully comply with all five rules. As a result, we evaluated the distance between the distributions of our generated molecules and real compounds using JSD.
- Diversity. This metric quantifies the chemical diversity among the generated molecules. It is defined as one minus the average Tanimoto similarity (based on molecular topological fingerprints) across all pairs of generated molecules. A desirable generative model should produce a diverse set of molecules rather than repeatedly sampling similar structures from a limited chemical space.

- Bond counts. It measures the distribution of the number of bonds per molecule. It helps detect potential biases towards molecules with an unusually high or low number of bonds (e.g., a bias towards long aliphatic chains might skew this distribution). The score is the JSD between the bond count distributions of generated molecules and test set molecules.

- Ring counts. This metric assesses the distribution of the number of rings per molecule, identifying potential biases towards excessive or insufficient ring structures. It was estimated as the JSD between the ring count distributions of generated molecules and reference molecules in the corresponding protein pockets.

- Molecular weight. This metric examines the distribution of molecular weights to detect biases towards generating overly large or small molecules. The score was calculated as the JSD between the molecular weight distributions of generated molecules and test set molecules.

- Bond types. This metric evaluates the distribution of fundamental bond types (single, double, triple). It helps identify biases towards specific bond types, which can occur if, for example, a model generates atoms first and adds bonds post-hoc without considering atom-bond interdependencies. The score is the JSD between the distributions of bond type ratios (proportions of single, double, triple bonds) in generated molecules and test set molecules.

- Ring sizes. This metric measures the distribution of ring sizes. It detects biases towards specific ring sizes, such as the excessive generation of small (3- or 4-membered) or large (>10-membered) rings observed in some earlier models[8,30,31]. The score is the JSD between the distributions of ring size ratios in generated molecules and test set molecules.

- Bond lengths. This metric assesses the accuracy of bond lengths in the generated 3D molecular structures. It is calculated as the average JSD across distributions of bond lengths for common bond types. Bond types are defined by the bond order (single, double, triple, aromatic) and the elements of the two connected atoms. For each type, the JSD compares the length distribution in generated molecules versus the test set. The SMARTS (SMILES arbitrary target specification) of considered bond types include: c:c, [#6]-[#6], [#6]-[#7], [#6]-O, c:n, [#6]=O, [#6]-S, O=S, c:o, c:s, [#6]-F, n:n, [#6]-Cl, [#6]=[#6], [#7]-S, [#6]=[#7], [#7]-[#7], [#7]-O, [#6]=S, and [#7]=O.

- Bond angles. This metric evaluates the accuracy of bond angles within the 3D structures. It was calculated as the average JSD across distributions of angles formed by common types of bonded atom triplets. Triplet types are defined by the elements of the three atoms and the types of the two connecting bonds. For each type, the JSD compares the angle distribution in generated molecules versus the test set. The SMARTS of considered types include: c:c:c, [#6]-[#6]-[#6], [#6]-[#7]-[#6], [#7]-[#6]-[#6], c:c-[#6], [#6]-O-[#6], O=[#6]-[#6], [#7]-c:c, n:c:c, c:c-O, c:n:c, [#6]-[#6]-O, and O=[#6]-[#7].

- Dihedral angles (JSD). This metric measures the accuracy of dihedral angles in the generated 3D structures. It is calculated as the average JSD across distributions of dihedral angles formed by common types of bonded atom quadruplets. Quadruplet types are defined by the elements of the four atoms and the types of the three connecting bonds. For each type, the JSD compares the dihedral angle distribution in generated molecules versus the test set. The SMARTS of considered types include: c:c:c:c, [#6]-[#6]-[#6]-[#6], [#6]-[#7]-[#6]-[#6], [#6]-c:c:c, [#7]-[#6]-[#6]-[#6], [#7]-c:c:c, O-c:c:c, [#6]-[#7]-c:c, [#7]-[#6]-c:c, n:c:c:c, [#6]-[#7]-[#6]=O, [#6]-[#6]-c:c, c:c-[#7]-[#6], c:n:c:c, and [#6]-O-c:c.

- 3D validity. This score assesses the overall validity of the generated 3D structure and topology using the PoseBusters[15] tool. PoseBusters performs various checks, such as the connectivity, steric clashes, internal energy, and proximity to the protein receptor. The final score is the proportion of generated molecules that pass all PoseBusters checks.

##### 3D molecule generation

- **Data**: The test set of 3D molecule generation contains 500 small molecules randomly sampled from the GEOM-Drug dataset[38], and the function of the test set is to provide molecular sizes for generation.

- **Protocols**: Each model generated 1000 small molecules for evaluation. The t-SNE visualization of distributions of these molecules was achieved by calculating the RDKit fingerprints of the generated molecules and the real molecules in the GEOM-Drug dataset. Since some methods use RDKit as post-processing, the generated molecules were defined as 2D valid if they were complete and were defined as 3D valid if their 3D structures passed all the filters of the PoseBusters validity checker[15]. We downloaded and installed the baselines to generate molecules by ourselves. For shape-conditioned molecules, we used PoseBusters to assess their validity.

##### Fragment linking

- **Data**: We used the test set proposed by the previous work DiffLinker[39], which originally contained 566 complexes of protein and fragment-linker pairs. It was built from the Binding MOAD dataset[40]. However, the molecules provided by DiffLinker did not contain the bond information. We tried to align the molecules in their test set with the Binding MOAD database processed by ourselves to add the bond information. Finally, we constructed a test set with 416 complexes of protein and fragment-linker pairs.

- **Protocols**: For each fragment within the pocket, each model generated 100 linkers with the same size as the corresponding molecule in the dataset. All the baseline models, including DeLinker[41], 3DLinker[42], and DiffLinker[39] were invoked using their default settings based on their papers. The generated molecules were defined as 2D valid if they could be loaded by RDKit and were defined as 3D valid if their 3D structures passed all the filters of the PoseBusters validity checker[15]. For each fragment, if any of the 100 generated molecules had the same non-isomeric SMILES as the true molecule, it was annotated as a molecule that recovered 2D graphs. If any of the 100 generated molecules had an RMSD less than 0.2 Å with the reference molecular linker, it was annotated as a molecule that recovered both 2D and 3D structures. The similarity between the generated molecules and the reference molecular linker in the test set was also calculated using the similarity function proposed by previous work DeLinker[41].

##### PROTAC design

- **Data**: We used the test set utilized by the previous work LinkerNet[43], which contained 43 fragment-linker pairs and was a subset of the PROTAC-DB database[44].

- **Protocols**: For the setting where fragment poses were known, each model generated 100 linkers for each fragment. For the setting where fragment poses were unknown, we randomly perturbed the true fragment poses as the input to the model. The perturbation contained five levels, which represented how different the input fragment poses were from the true ones. We generated 30 molecules for each fragment and perturbation level. We installed and generated molecules following the papers of the baselines[39,43].

##### Fragment growing

- **Data**: We used the test set of the SBDD task, which originally contained 100 protein-small-molecule complexes. We discarded samples whose small molecules had fewer than 20 atoms. For the remaining data, we decomposed the molecules into multiple fragments using BRICS decomposition. Starting from the fragments that were closest to the protein pockets, we kept connecting the neighboring fragments until the number of atoms was larger than 19. We then removed data whose remaining atoms were fewer than five. In this way, we constructed a test set for fragment growing, which contained 53 complexes of proteins and molecular fragments.

- **Protocols**: For each fragment in the test set, we generated 100 molecules that could potentially bind the protein pocket, and ensured the generated molecule had the same number of atoms as the original molecules in the dataset. We installed the baseline DiffDec and generated molecules by ourselves.

##### Molecule optimization

- **Data**: We used the same test set as the SBDD task for this task.

- **Protocols**: For each pocket–small molecule pair, the small molecule was used as the initial denoising input with an initial noise scale of 0.3 and a total of 30 denoising steps. For each pair, PocketXMol sampled 100 new molecules, from which the top 10 closest to the target LogP value were selected as optimized candidates. These selected molecules then served as starting points for the subsequent optimization round. A total of three optimization rounds were performed.

##### Linear peptide design

- **Data**: We used the test set of the peptide docking task, which originally contained 79 protein-peptide complexes. We then removed the peptides larger than 15, peptides with numbers of atoms more than 150, or peptides with non-standard amino acid types or backbones. Finally, we obtained 35 protein-peptide complexes for benchmarking *de novo* peptide design. For the Q-BioLiP peptide design test set, we used the Q-BioLiP peptide docking test set, which originally contained 112 complexes. We then removed the peptides longer than 15 residues, shorter than 3 residues, or containing non-standard amino acids, which resulted in 57 protein-peptide complexes for benchmarking peptide design.

- **Protocols**: Each model generated 100 valid peptides for each pocket. We installed RFdiffusion[45], ProteinMPNN[46], and Rosetta[47] to generate peptides for the test set with their default settings. For the metric sequence recovery, we calculated the sequence identity of the generated peptides and the true ones and selected the highest recovery per pocket. For the metric backbone RMSD, we calculated the backbone alignment RMSDs between the generated peptides and the true ones and selected the lowest RMSD per pocket. The Rosetta binding energy was calculated by using the Interface Analyzer of the Rosetta tool[47]. We installed the MolProbity tool locally to assess the MolProbity metrics of the peptides.

##### Cyclic peptide design

- **Data**: Based on the cyclic peptide docking test set, we selected one representative from each cluster to construct the test set for cyclic peptide design.

- **Protocols**: The metrics are the same as the evaluation of linear peptide design. Each model generated 100 standard peptides for each test pocket. For the baseline AfDesign(cyclic), we used the provided Colab notebook from the work[48] and generated peptides for the test set with default parameters.

##### Peptide inverse folding

- **Data**: We used the same test set as the *de novo* peptide design.

- **Protocols**: Each model generated 100 peptides for each pair of proteins and the backbone structures. We installed ProteinMPNN[46] to generate peptides for the test set with the default setting.

#### 6.2 Training data and redundancy control

##### 6.2.1 Data filtering

We used three types of data from multiple databases or datasets: 1) protein-small-molecule complexes, including PDBBind[49], Binding MOAD[40], and the CrossDocked2020 dataset[33]; 2) protein-peptide complexes including PepBDB[21] and a synthetic dataset of protein-binding loop structures from AlphaFoldDB[50]; 3) small molecule or peptide structures, including the GEOM dataset[38] (GEOM-Drug and GEOM-QM9), CREMP dataset[51], and a large molecular data dataset proposed by Uni-Mol[19] which contained small molecules from ZINC[52], ChEMBL[53] and other purchasable molecular databases.

We filtered out data that violated any of the following criteria: 1) successful loading by RDKit[13]; 2) inclusion of only pre-defined atom types including C, N, O, F, P, S, Cl, B, Br, I, and Se; 3) being a complete molecule; 4) for small molecules, the count of heavy atoms should be within the range of 4 to 122; 5) for peptides, the length should be smaller than 15 and the count of heavy atoms should not exceed 150. After filtering for individual datasets, we eliminated duplicate entries from each dataset to compile a comprehensive dataset.

##### 6.2.2 Redundancy control

To prevent data leakage, we excluded training samples that shared significant similarity with any data point in the test sets. For tasks not involving protein pockets, we removed training samples that had identical non-isomeric SMILES to those in the test sets. For tasks involving protein pockets, we applied MMseqs2 clustering[23] to all protein sequences across training and test sets using a sequence identity threshold of 0.3. Any training sample whose protein sequence fell into the same cluster as a test sequence was removed. In cases where a data point contained multiple protein chains, we selected the chain with the greatest number of contacts with the ligand for clustering and comparison.

##### 6.2.3 Dataset splits

We randomly divided the collected dataset into training and validation sets and the sizes of validation sets are determined considering the sizes of the datasets. The numbers of data points in the training and validation sets are shown in the Table S2.

**Table S2. The number of samples in the training and validation sets from individual source datasets.**

| Source datasets | Training set | Validation set |
|---|---|---|
| PDBBind | 8,916 | 100 |
| Bind MOAD | 24,789 | 100 |
| CrossDocked2020 | 51,429 | 100 |
| AlphaFold DB | 36,319 | 100 |
| PepBDB | 3,422 | 70 |
| GEOM-Drug | 299,762 | 500 |
| GEOM-QM9 | 132,366 | 500 |
| Uni-Mol data | 11,515,474 | 500 |
| CREMP | 35,698 | 500 |

As shown in Algorithm 61, we need to sample a molecule at each training step. Due to the unbalanced sizes of each individual dataset in the assembled training dataset, we defined different probabilities for each individual dataset to be sampled during the training phase. The probabilities also emphasized different datasets for different training tasks. The un-normalized probabilities were described in Table S3.

**Table S3. Sampling ratio of training datasets for different training tasks.**

| Source datasets | 3D molecule generation | Molecule conformation generation | SBDD | Docking | Mask-fill | FBDD | Peptide design |
|---|---|---|---|---|---|---|---|
| PDBBind | N/A | N/A | 0.012 | 0.115 | 0.09 | 0.1 | N/A |
| Bind MOAD | N/A | N/A | 0.03 | 0.03 | 0.2 | 0.2 | N/A |
| CrossDocked2020 | N/A | N/A | 0.064 | 0.032 | 0.5 | 0.5 | N/A |
| AlphaFold DB | N/A | N/A | 0.01 | 0.06 | 0 | 0 | 37 |
| PepBDB | N/A | N/A | 0.003 | 0.03 | 0 | 0 | 7 |
| GEOM-Drug | 0.6 | 0.3 | N/A | N/A | 0.2 | 0.4 | N/A |
| GEOM-QM9 | 0.08 | 0.133 | N/A | N/A | 0.01 | 0.02 | N/A |
| Uni-Mol data | 11.5 | 0.3 | N/A | N/A | 2 | 4 | N/A |
| CREMP | 0 | 0.03 | N/A | N/A | 0 | 0 | N/A |

#### 6.3 Data processing

We used RDKit to process the small molecules and peptides. We removed the hydrogen atoms and collected the atom types, bond types, and conformations (atom coordinates). If multiple conformations existed, all conformations were reserved, from which one conformation was sampled at the training step. We then used RDKit to annotate the rotatable bonds and calculated the decompositions using the BRICS and MMPA algorithms. For peptides, we annotated the backbone atoms and side-chain atoms using BioPython[54]. We parsed the protein receptor files and defined the pocket as the amino acids whose distances with any atom in the molecule were less than 10Å. The pocket was represented as a set of atoms, whose features included their atom coordinates, atom types, amino acids they belong to, and whether they were in the backbone.

##### 6.4 Variant pocket structures

The variant pocket structures were derived from the receptor protein structures from four sources:

1. **AlphaFold-predicted structures.** We used AlphaFold 2[25] to predict the receptor structures. For multi-chain receptors, we employed AlphaFold-Multimer[55]. All 25 predicted structures from AlphaFold 2 or AlphaFold-Multimer for each receptor were retained. These predictions were backbone-aligned to the true structures to determine the pockets using our previous strategy, i.e., selecting residues within 10 Å of the ligand. The resulting pocket structures then replaced the true structures as input.

2. **Rosetta-repacked structures.** We applied the `fixbb` function in Rosetta[47] to repack the side chains of the native receptor structures. For each receptor, five repacked structures were randomly sampled. Pocket identification followed the same procedure as above.

3. **Retrieved structures from RCSB PDB.** We searched for experimentally determined receptor structures in the RCSB PDB[56] with identical amino acid sequences. The search procedure followed a template matching strategy similar to that used in AlphaFold's inference pipeline[25]. Candidate structures containing the same ligand CCD ID as the true ligand were excluded. For each remaining candidate, we used PyMol to align its backbone to that of the true structure and verified the consistency of residue types at pocket positions. Structures with mismatched residue types or PyMol-refined pocket backbone RMSD greater than 2 Å were discarded. The protein chains containing aligned pocket residues were extracted from the retrieved complex as the final retrieved receptors. Pockets were then defined as residues within 10 Å of the true ligand.

4. **Retrieved apo structures from RCSB PDB.** This followed the same procedure as above, with an additional filtering step to remove holo structures. After pocket alignment, any candidate structure containing atoms other than receptor or solvent within 2 Å of the ligand in the true structure was considered holo and excluded. Structures lacking such atoms were retained as apo variants.

#### 7 Application to enzyme-substrate interactions

##### 7.1 Enzyme-substrate datasets

We collected four enzyme-substrate datasets from related studies[57–61]. These datasets contained pairs of enzymes and small molecule substrates from four distinct enzyme families: halogenases, esterases, glycosyltransferases, and phosphatases (Table S18). Each pair within these datasets was assigned a binary label (active/inactive) based on the findings reported in the original publications, indicating whether the enzyme could catalyze the substrate. As the datasets provided only the enzyme sequences, we predicted the protein structures using AlphaFold-Multimer[55]. We identified the active site pockets by aligning these predicted structures to reference structures from the RCSB PDB known to contain the active site for each enzyme family (Table S18).

##### 7.2 Activity prediction

For each substrate and enzyme pocket pair, we utilized PocketXMol to generate 50 docking poses and selected the pose with the highest PocketXMol self-confidence score. Similarly, we also used the baseline AlphaFold 3 to predict the enzyme-substrate complex structures and used the best confidence score among its 25 predictions per pair. To have a quantitative evaluation, we calculated the Area Under the Receiver Operating Characteristic curve (AUROC) and the Area Under the Precision-Recall curve (AUPR) based on the confidence scores for individual proteins.

To adapt PocketXMol for enzyme activity prediction, we used it to generate structural representations of enzyme–substrate pairs and fine-tuned a simple logistic regression model on activity-labeled data. Specifically, PocketXMol was used to dock each pair, and the feature vectors from its final layer were extracted as representations, which were then input into the logistic regression model to predict enzyme activity. We evaluated performance using cross-validation across different data-splitting schemes, including splits for new substrates, new enzymes, new enzyme–substrate pairs, and new substrates and enzymes (both new). Note that only enzymes with at least five active and five inactive substrates (named as valid enzymes in Table S18) were considered here to ensure reliable calculation of AUROC and AUPR scores.

##### 7.3 Docking structure analyses

We identified catalytic residues by aligning each enzyme sequence to its corresponding reference sequences (reference PDB IDs listed in Table S18) whose catalytic resides were identified by referring to UniProt[62]. For halogenases, catalytic residues were defined as those aligned with the lysine in the AFKAA motif of the reference. For glycosyltransferases, they corresponded to the histidine in the GTHAA motif, and for phosphatases, to the aspartic acid in the FLDRD motif. Esterases were excluded from this analysis due to the inability to reliably identify their catalytic residues through sequence alignment.

For the substrates, we identified potential reactive atoms by referencing prior studies on the corresponding enzyme families[58,60,61]. In halogenases, reactive sites typically involve activated C–H bonds; thus, we considered aromatic ring carbon atoms bearing hydrogen atoms as candidate reactive atoms. In glycosyltransferases, nucleophilic groups such as hydroxyl, amino, and thiol moieties act as acceptors, so we matched substructures corresponding to –OH, $-NH_2$, and –SH as potential reactive groups. For phosphatases, where the phosphate group directly participates in the catalytic process, all phosphate moieties within the substrates were designated as reactive groups.

Based on the above criteria, we identified the catalytic residues and reactive groups for each positive enzyme–substrate pair in the three datasets. We then used PocketXMol and AlphaFold 3 to predict the corresponding binding structures and computed the distances between the catalytic residue side chains and various atoms of the substrates. Specifically, we evaluated: (1) the substrate distance, defined as the minimum distance from any substrate atom to the catalytic residues; (2) the reactive group distance, defined similarly but restricted to atoms within the reactive group; and (3) the substrate center distance, calculated as the average distance between all substrate atoms and the catalytic residues. We quantified the proportion of samples satisfying specific distance thresholds (5Å, 8Å, and 10Å) for both the full substrates and reactive groups. Additionally, we measured the fraction of cases in which the reactive group was positioned closer to the catalytic residues than the center of the substrate, considering all samples as well as the subset where the substrate distance was less than 5Å.

### 8 Wet-lab validated molecule design

#### 8.1 Caspase-9 binding small molecule design

##### 8.1.1 De novo generation of molecules

We used the caspase-9 structure (PDB ID: 2AR9) as the binding protein structure. To determine the binding pocket, we used another protein in the caspase family as a reference. Specifically, we used a ligand-binding caspase-8 complex structure (PDB ID: 3KJN). We downloaded their structure files from RCSB PDB[56] and aligned the structures of caspase-9 to caspase-8. Then pocket center of the caspase-9 structure is specified by the position of a medium atom (an oxygen atom of the amide group) of the ligand (CCD ID: B93). We defined the pocket as the amino acids of caspase-9 within 15 Å of the center positions. We then set molecular sizes sampled from Gaussian distributions with a mean of 25 and a standard deviation of 5. Then we utilized PocketXMol to generate novel small molecules for the pocket through the SBDD task.

We finally collected 270,281 small molecules. We first used RDKit[13] and followed previous SBDD works[31] to calculate the basic properties of the molecules, including QED (quantitative estimate of druglikeness) score, SA (synthesis accessibility) score, predicted LogP, the number of hydrogen acceptors, the number of hydrogen donors, the number of rotatable bonds, the number of satisfied criteria of Lipinski's rule of five, and whether belonging to PAINS (pan-assay interference compounds). We removed molecules with any of the following properties: (1) QED scores less than 0.4; (2) SA less than 0.7; (3) the number of satisfied criteria of Lipinski's rule of five less than three; (4) belonging to PAINS. This filtering resulted in 135,315 remaining molecules. Next, we analyzed the molecular graph and removed those with any of the following criteria: (1) containing three-membered rings or four-membered rings; (2) containing large rings with sizes $\geq 7$; (3) containing too many rings (over four rings); (4) the number of atoms are not between 10 to 100; (5) the number of atoms that belonged to more than one ring was over two (i.e., containing over one fused ring). This filtering resulted in 110,779 molecules.

Next, we used RDKit with the MMFF94 force field to optimize the molecules and used OpenMM[63] to optimize the generated complex structures of caspase-9 and molecules through energy minimization. We then utilized a protein-ligand binding affinity predictor[8] to predict the binding affinities between generated molecules and the caspase-9. We selected those with high binding affinities, i.e., with predicted dissociation constant $K_D < 10^{-7}$M, resulting in 2,153 molecules. We then predicted their binding affinities with the proteins caspase-1/3/8 whose pocket structures were similar to that of caspase-9 to evaluate their specificity. We finally selected molecules with predicted $K_D < 10^{-8}$M for caspase-9 and with $K_D > 10^{-6}$M for caspase-1/3/8, which resulted in 67 molecules. We then applied AutoDock Vina to calculate the Vina scores of these molecules. Taking the ease of synthesis and all the above metrics into consideration, we prioritized 16 molecules to experimentally synthesize and validate.

##### 8.1.2 Re-optimization of the molecule 84663

The second-round candidate molecules were derived based on the PocketXMol-predicted binding structure of molecule 84663 with caspase-9 (Figure 3b), employing two complementary design strategies. The first strategy utilized the molecular optimization capability of our model. Starting from 84663, a small perturbation was introduced via a series of small noise levels (0.1, 0.2, and 0.3), followed by iterative denoising. This process generated analogs that preserved the core scaffold of 84663 while potentially improving molecular properties or binding interactions. The second strategy involved straightforward structure-guided manual modification. By analyzing the predicted protein–ligand complex and referencing known caspase inhibitors from the literature[64], we applied structure–activity relationship (SAR) insights to design new derivatives, such as including alterations to hydrophobic ring systems and the incorporation of covalent warheads, to enhance binding affinity or reactivity. All candidate molecules from both strategies were subsequently redocked using PocketXMol, and evaluated based on confidence scores and redocking RMSD. Additional assessments included Vina docking scores, synthetic accessibility (SA) scores, detailed binding mode analyses, and retrosynthetic route evaluations. Based on these comprehensive criteria, a subset of optimized candidates was selected for synthesis and experimental validation in the second round.

#### 8.2 PD-L1 binding peptide design

##### *8.2.1 Peptide design with filtering*

We used three PD-L1 structures (PDB ID: 3BIK, 4ZQK, and 5IUS) for binding peptide design. We selected the three structures from the PDB based on the criterion that they were complex structures of PD-1 and PD-L1 without mutations on PD-L1 and with relatively low resolutions. The resolutions for 3BIK, 4ZQK, and 5IUS are 2.65Å, 2.45Å, and 2.89Å, respectively. This selection allowed for more reliable protein structures and peptide design..

We downloaded their complex structure files from RCSB PDB[56] and aligned their PD-L1 structures. We defined the centers of the protein pockets at the intermediate positions between the surfaces of PD-1 and PD-L1. We then removed atoms not belonging to PD-L1 from the structures. We defined the pockets as the amino acids of PD-L1 within 20 Å of the center positions. We set the generated peptide length as 11 to allow peptides with enough flexibility to bind the target. Algorithm 56 was adopted to generate peptides for the three pockets.

In the end, we collected 944,552 generated valid peptides. We then filtered them as follows. We first used the flexible noise to calculate the confidence score, named self-confidence score, for the generated peptides (Algorithm 32) and removed peptides with the lowest 50% self-confidence scores, resulting in 472,275 peptides. We then used the commands `AnalyseComplex` and `Stability` of FoldX[65] to calculate the FoldX binding energies, clashes of peptides, and stability energies of the complexes of remaining peptides and PD-L1. We removed peptides with the worst 30% FoldX binding energies and subsequently removed peptides with the worst 30% stability energies of individual pockets, resulting in 253,026 peptides. FoldX was used as a preliminary filter to reduce the number of generated peptides to a manageable scale before applying the more computationally intensive Rosetta calculations. While both tools serve similar purposes, their combined use enhanced filtering efficiency and provided a more comprehensive reference for selecting peptide candidates.

Next, we used the refinement mode, rescoring mode, and the minimization mode of the FlexPepDock Refinement protocol of Rosetta[47,66] to score the remaining complexes and removed those with the worst 30% refinement scores of individual pockets, resulting in 184,140 peptides. Subsequently, we removed those with minimization scores larger than 100 or rescoring scores larger than 1000, resulting in 176,225 peptides. We used the InterfaceAnalyzer protocol of Rosetta[47,67] to calculate the change in Rosetta energy, the solvent accessible area, and the number of residues at the surface, and removed peptides with the worst 30% change in Rosetta energy and subsequently those with the worst 10% solvent accessible area at the interface, resulting 116,657 peptides.

We then considered all the above metrics simultaneously. We defined two sets of criteria. The first set was: (1) refinement score: top 40%; (2) change of Rosetta energy: top 40%; (3)

minimization score: top 80%; (4) rescoring score: top 80%; (5) clash of peptide: top 40%; (6) FoldX binding energy: top 40%; (7) FoldX stability energy: top 40%; (8) solvent accessible area at the interface: top 60%; (9) number of residues at the interface: top 80%; and (10) self-confidence score: top 60%. The second set was: (1) refinement score: top 200; (2) change of Rosetta energy: top 200; (3) minimization score: top 100; (4) rescoring score: top 100; (5) FoldX binding energy: top 200; (6) FoldX stability energy: top 200; (7) solvent accessible area at the interface: top 100; and (8) self-confidence score: top 200. From the remaining peptides, we selected those satisfying at least seven out of ten criteria in the first set or any of the criteria in the second set, resulting in 39,548 peptides.

Next, we employed AlphaFold-Multimer[55] to predict the complexes of PD-L1 and the generated peptides and used the confidence scores of AlphaFold-Multimer, named AlphaFold confidence score, as a new metric. We only predicted one complex per generated peptide for efficiency. We calculated the RMSDs of the peptide structures generated by PocketXMol and predicted by AlphaFold-Multimer, named RMSD consistency. We defined two sets of criteria. The first set was (1) consistency RMSD less than 4.5Å and (2) AlphaFold confidence score larger than 0.5. The second was (1) consistency RMSD less than 2.5Å; (2) AlphaFold confidence score larger than 0.8; and (3) self-confidence score larger than -0.9. We selected 568 peptides satisfying the two criteria in the first set or any criterion in the second set.

For the remaining 568 peptides, we utilized AlphaFold-Multimer to predict their complex structures. This time we used the default setting, i.e., predicted 25 complexes per generated peptide, and chose the one with the best AlphaFold confidence scores per peptide. Similarly, we also defined two sets of criteria. The first set was (1) consistency RMSD less than 4.2Å and (2) AlphaFold confidence score larger than 0.5. The second was (1) consistency RMSD less than 2.5Å; (2) AlphaFold confidence score larger than 0.85; and (3) self-confidence score larger than -0.899. There were 172 peptides satisfying the two criteria in the first set and 231 peptides satisfying any criterion in the second set. A combination of them resulted in 382 peptides, which were used for further wet lab experiments.

###### 8.2.2 Peptide design without filtering

The process was similar to the above with minor modifications. Specifically, we only used the PDB structure 5IUS as the pocket structure. Following the same sampling procedure as above, we selected the first 382 standard peptides for wet-lab evaluation, without applying any filtering or selection criteria.

### Supplementary Tables

**Table S4. The purchasable molecular databases from the Uni-Mol dataset.** This table is based on the Uni-Mol paper[19].

| Database | Number of molecules | Link |
|---|---|---|
| Targetmol | 10,000 | https://www.targetmol.com/ |
| Chemdiv | 1,613,931 | https://www.chemdiv.com/ |
| Enamine | 2,734,581 | https://enamine.net/ |
| Chembridge | 1,557,942 | https://www.chembridge.com/ |
| Life Chemical | 509,975 | https://lifechemicals.com/ |
| Specs | 208,670 | https://www.specs.net/ |
| Vitas-M | 1,409,339 | https://vitasmlab.biz/ |
| InterBioScreen | 48,627 | https://www.ibscreen.com/ |
| Maybridge | 53,352 | https://www.thermofisher.in/ |
| Bionet-Key Organics | 259,244 | https://www.keyorganics.net/ |
| Asinex | 530,881 | https://www.asinex.com/ |
| UkrOrgSynthesis | 688,952 | https://uorsy.com/ |
| Eximed | 61,009 | https://eximedlab.com/ |
| HTS Biochemie Innovationen | 58,437 | https://www.hts-biochemie.de/ |
| Princeton BioMolecular | 1,532,542 | https://princetonbio.com/ |
| Otava | 270,835 | https://otavachemicals.com/ |
| Alinda Chemical | 202,332 | https://www.alinda.ru/ |
| Analyticon | 42,664 | https://www.analyticon-diagnostics.com/ |

**Table S5. Performance on structure-based drug design.** These numbers are the raw values of each metric. They were transformed to the normalized scores in Table S6 for the radar plot in the main article.

| | PocketX Mol | PocketX Mol (AR) | liGAN | 3DSBD D | Pocket2 Mol | Target Diff | Decomp Diff | IPDiff | AliDiff |
|---|---|---|---|---|---|---|---|---|---|
| QED (↑) | 0.52 | 0.51 | 0.39 | 0.51 | 0.57 | 0.48 | 0.45 | 0.50 | 0.50 |
| SA (↑) | 0.77 | 0.75 | 0.59 | 0.63 | 0.76 | 0.58 | 0.61 | 0.56 | 0.56 |
| LogP (JSD. ↓) | 0.16 | 0.15 | 0.17 | 0.20 | 0.25 | 0.15 | 0.24 | 0.40 | 0.42 |
| Lipinski (JSD. ↓) | 0.11 | 0.10 | 0.10 | 0.20 | 0.28 | 0.11 | 0.15 | 0.19 | 0.20 |
| Diversity (↑) | 0.74 | 0.74 | 0.66 | 0.70 | 0.74 | 0.72 | 0.67 | 0.74 | 0.73 |
| Bond counts (JSD. ↓) | 0.19 | 0.19 | 0.20 | 0.25 | 0.27 | 0.20 | 0.39 | 0.22 | 0.22 |
| Ring counts (JSD. ↓) | 0.17 | 0.17 | 0.19 | 0.22 | 0.23 | 0.18 | 0.26 | 0.25 | 0.25 |
| Molecular weight (JSD. ↓) | 0.20 | 0.20 | 0.20 | 0.27 | 0.30 | 0.23 | 0.37 | 0.22 | 0.22 |
| Bond types (JSD. ↓) | 0.01 | 0.01 | 0.02 | 0.03 | 0.06 | 0.01 | 0.03 | 0.03 | 0.03 |
| Ring sizes (JSD. ↓) | 0.13 | 0.13 | 0.43 | 0.32 | 0.15 | 0.25 | 0.24 | 0.31 | 0.31 |
| Bond lengths (JSD. ↓) | 0.43 | 0.43 | 0.60 | 0.50 | 0.49 | 0.41 | 0.35 | 0.48 | 0.49 |
| Bond angles (JSD. ↓) | 0.16 | 0.16 | 0.51 | 0.33 | 0.23 | 0.23 | 0.18 | 0.30 | 0.31 |
| Dihedral angles (JSD. ↓) | 0.28 | 0.29 | 0.38 | 0.47 | 0.34 | 0.32 | 0.31 | 0.36 | 0.36 |
| 3D validity (↑) | 0.85 | 0.83 | 0.03 | 0.59 | 0.72 | 0.51 | 0.49 | 0.22 | 0.25 |

**Table S6. Normalized scores on structure-based drug design.** These scores are the normalized version of values in Table S5. Each metric was normalized to a scale from 0 to 1, with 1 representing the best method and 0 indicating the worst. The average score was calculated as the non-weighted average over all metrics. These numbers are used for the radar plot of the SBDD task in the main article.

| | PocketX Mol | PocketX Mol(AR) | liGAN | 3D SBDD | Pocket2 Mol | Target Diff | Decomp Diff | IPDiff | AliDiff |
|---|---|---|---|---|---|---|---|---|---|
| Average score | 0.94 | 0.93 | 0.43 | 0.51 | 0.62 | 0.74 | 0.48 | 0.49 | 0.48 |
| QED | 0.70 | 0.63 | 0.00 | 0.65 | 1.00 | 0.49 | 0.35 | 0.61 | 0.62 |
| SA | 1.00 | 0.90 | 0.14 | 0.35 | 0.95 | 0.11 | 0.22 | 0.00 | 0.01 |
| LogP (JSD) | 0.98 | 1.00 | 0.93 | 0.82 | 0.65 | 1.00 | 0.69 | 0.09 | 0.00 |
| Lipinski (JSD) | 0.90 | 0.95 | 1.00 | 0.45 | 0.00 | 0.92 | 0.69 | 0.46 | 0.40 |
| Diversity | 0.96 | 1.00 | 0.00 | 0.49 | 1.00 | 0.71 | 0.16 | 0.94 | 0.92 |
| Bond counts (JSD) | 1.00 | 0.99 | 0.93 | 0.70 | 0.60 | 0.92 | 0.00 | 0.83 | 0.84 |
| Ring counts (JSD) | 0.94 | 1.00 | 0.77 | 0.46 | 0.35 | 0.89 | 0.00 | 0.16 | 0.09 |
| Molecular weight (JSD) | 0.99 | 1.00 | 0.98 | 0.58 | 0.44 | 0.82 | 0.00 | 0.84 | 0.87 |
| Bond types (JSD) | 1.00 | 0.94 | 0.87 | 0.69 | 0.00 | 0.95 | 0.66 | 0.67 | 0.66 |
| Ring sizes (JSD) | 1.00 | 1.00 | 0.00 | 0.37 | 0.93 | 0.61 | 0.61 | 0.42 | 0.40 |
| Bond lengths (JSD) | 0.70 | 0.68 | 0.00 | 0.41 | 0.47 | 0.76 | 1.00 | 0.49 | 0.44 |
| Bond angles (JSD) | 1.00 | 1.00 | 0.00 | 0.52 | 0.81 | 0.80 | 0.96 | 0.62 | 0.59 |
| Dihedral angles (JSD) | 1.00 | 0.95 | 0.46 | 0.00 | 0.66 | 0.79 | 0.82 | 0.56 | 0.57 |
| 3D validity | 1.00 | 0.98 | 0.00 | 0.68 | 0.84 | 0.58 | 0.56 | 0.23 | 0.27 |

**Table S7.** **Performance on the SBDD task of PocketXMol with different input pocket structures.**

| | Holo structure | AFM predicted | Retrieved from PDB | Retrieved from PDB (apo only) | Rosetta repacked |
|---|---|---|---|---|---|
| QED (↑) | 0.52 | 0.52 | 0.52 | 0.54 | 0.52 |
| SA (↑) | 0.77 | 0.77 | 0.77 | 0.78 | 0.78 |
| LogP (JSD. ↓) | 0.16 | 0.16 | 0.16 | 0.18 | 0.17 |
| Lipinski (JSD. ↓) | 0.11 | 0.14 | 0.13 | 0.17 | 0.15 |
| Diversity (↑) | 0.74 | 0.75 | 0.75 | 0.77 | 0.76 |
| Bond counts (JSD. ↓) | 0.19 | 0.19 | 0.18 | 0.20 | 0.18 |
| Ring counts (JSD. ↓) | 0.17 | 0.18 | 0.17 | 0.15 | 0.18 |
| Molecular weight (JSD. ↓) | 0.20 | 0.20 | 0.20 | 0.23 | 0.22 |
| Bond types (JSD. ↓) | 0.01 | 0.02 | 0.01 | 0.02 | 0.03 |
| Ring sizes (JSD. ↓) | 0.13 | 0.13 | 0.12 | 0.13 | 0.12 |
| Bond lengths (JSD. ↓) | 0.42 | 0.43 | 0.43 | 0.42 | 0.41 |
| Bond angles (JSD. ↓) | 0.16 | 0.17 | 0.16 | 0.17 | 0.17 |
| Dihedral angles (JSD. ↓) | 0.28 | 0.28 | 0.28 | 0.28 | 0.28 |
| 3D validity (↑) | 0.85 | 0.83 | 0.83 | 0.84 | 0.75 |

**Table S8.** **Validity checks of the molecules generated in the constraints of letter shapes.** The check was conducted by the PoseBusters tool.

| Letter | C | E | K | L | M | O | P | T | X |
|---|---|---|---|---|---|---|---|---|---|
| Successfully loaded | 100.0 | 100.0 | 100.0 | 100.0 | 100.0 | 100.0 | 100.0 | 100.0 | 100.0 |
| Successful sanitization | 100.0 | 100.0 | 100.0 | 100.0 | 100.0 | 100.0 | 100.0 | 100.0 | 100.0 |
| All atoms connected | 100.0 | 99.0 | 99.0 | 100.0 | 95.6 | 100.0 | 99.0 | 99.0 | 98.9 |
| Bond length within bounds | 99.8 | 100.0 | 100.0 | 100.0 | 99.8 | 100.0 | 99.0 | 100.0 | 98.9 |
| Bond angle within bounds | 99.8 | 97.9 | 98.0 | 99.8 | 96.6 | 98.5 | 99.0 | 100.0 | 93.4 |
| No internal steric clash | 98.3 | 99.0 | 100.0 | 98.1 | 98.3 | 55.1 | 93.8 | 100.0 | 98.9 |
| No deformed aromatic rings | 100.0 | 100.0 | 100.0 | 100.0 | 100.0 | 100.0 | 100.0 | 100.0 | 100.0 |
| No defomed double bonds | 100.0 | 99.0 | 98.0 | 99.8 | 99.6 | 99.6 | 100.0 | 100.0 | 98.9 |
| Energy not too high | 99.2 | 93.8 | 100.0 | 99.4 | 91.8 | 97.0 | 95.9 | 93.8 | 96.7 |
| Pass all tests | 97.3 | 90.7 | 94.9 | 97.1 | 82.9 | 52.5 | 87.6 | 92.8 | 89.0 |

**Table S9.** **Surface plasmon resonance (SPR) analysis of D12 and 84663 binding to caspase-9 and its mutants.**

| | 84663 | D12 |
|---|---|---|
| WT CASP9 | 2.11e-06 | 9.11e-07 |
| CASP9 C287A | 7.96e-05 | 3.39e-06 |
| CASP9 R180A | 4.44e-06 | 2.19e-06 |
| CASP9 Q285A | 8.30e-06 | 4.79e-06 |

**Table S10.** **Ratios satisfying MolProbity metrics for peptides in the peptide design test set and generated by RFdiffusion or PocketXMol.** The results of PocketXMol peptides with filtering of high confidence is also included where only those with self confidence larger than 0.8 were chosen.

| Metric | Criterion | Test set | RFdiffusion | PocketXMol | PocketXMol (high confidence) |
|---|---|---|---|---|---|
| Clashscore | =0 | 0.86 | 0.54 | 0.58 | 0.80 |
| CaBLAM disfavored ratios | =0 | 0.69 | 0.97 | 0.80 | 1.00 |
| CaBLAM outlier ratios | =0 | 0.86 | 0.98 | 0.91 | 1.00 |
| CA Geometry outlier ratios | =0 | 0.94 | 1.00 | 0.98 | 1.00 |
| C$\beta$ deviations > 0.25Å | =0 | 1.00 | 1.00 | 0.53 | 0.87 |
| cis prolines out of PRO | =0 | 1.00 | 1.00 | 1.00 | 1.00 |
| twisted prolines out of PRO | =0 | 1.00 | 0.99 | 0.95 | 1.00 |
| cis prolines out of nonPRO | =0 | 1.00 | 1.00 | 1.00 | 1.00 |
| twisted prolines out of PRO | =0 | 1.00 | 0.99 | 0.97 | 1.00 |
| Ramachandran outlier ratios | =0 | 0.97 | 0.97 | 0.85 | 1.00 |
| Ramachandran favored ratios | =100% | 0.69 | 0.90 | 0.61 | 0.73 |
| Rotamer outlier ratios | =0 | 0.63 | 1.00 | 0.19 | 0.27 |
| Bond length outlier ratios | <10% | 1.00 | 0.94 | 0.85 | 1.00 |
| Bond angle outlier ratios | <10% | 1.00 | 0.97 | 0.59 | 0.80 |

**Table S11.** **Ratios satisfying MolProbity metrics for the peptides generated by PocketXMol within different confidence intervals.**

| Metric | Criterion | PocketXMol confidence interval | | | | |
|---|---|---|---|---|---|---|
| | | <0.5 | (0.5, 0.6] | (0.6, 0.7] | (0.7, 0.8] | >0.8 |
| Clashscore | =0 | 0.52 | 0.47 | 0.61 | 0.70 | 0.80 |
| CaBLAM disfavored ratios | =0 | 0.62 | 0.61 | 0.93 | 0.99 | 1.00 |
| CaBLAM outlier ratios | =0 | 0.83 | 0.81 | 0.97 | 0.99 | 1.00 |
| CA Geometry outlier ratios | =0 | 0.97 | 0.94 | 1.00 | 0.99 | 1.00 |
| C$\beta$ deviations > 0.25Å | =0 | 0.40 | 0.35 | 0.63 | 0.69 | 0.87 |
| Cis prolines out of PRO | =0 | 1.00 | 1.00 | 1.00 | 1.00 | 1.00 |
| Twisted prolines out of PRO | =0 | 0.90 | 0.96 | 0.97 | 0.95 | 1.00 |
| Cis prolines out of nonPRO | =0 | 1.00 | 1.00 | 1.00 | 1.00 | 1.00 |
| Twisted prolines out of PRO | =0 | 0.95 | 0.94 | 0.98 | 0.99 | 1.00 |
| Ramachandran outlier ratios | =0 | 0.70 | 0.74 | 0.94 | 0.99 | 1.00 |
| Ramachandran favored ratios | =100% | 0.33 | 0.40 | 0.80 | 0.87 | 0.73 |
| Rotamer outlier ratios | =0 | 0.24 | 0.23 | 0.13 | 0.18 | 0.27 |
| Bond length outlier ratios | <10% | 0.81 | 0.73 | 0.90 | 0.93 | 1.00 |
| Bond angle outlier ratios | <10% | 0.66 | 0.64 | 0.55 | 0.49 | 0.80 |

**Table S12.** **Sequence recovery rates and the backbone structure RMSDs of cyclic peptides generated by AfDesign(cyclic) and PocketXMol.** Peptides with the highest sequence recovery rates among the 100 generated samples for each pocket were selected for comparison. The minimum backbone RMSDs among 100 generated peptides for each pocket were selected for comparison.

| PDB ID | Sequence recovery (%) | | Structure RMSD (Å) | |
|---|---|---|---|---|
| | AfDesign (cyclic) | PocketXMol | AfDesign (cyclic) | PocketXMol |
| 1sfi | 42.86 | 42.86 | 1.95 | 1.56 |
| 3av9 | 37.50 | 37.50 | 1.34 | 2.47 |
| 3p8f | 35.71 | 35.71 | 2.31 | 1.44 |
| 3wne | 33.33 | 33.33 | 1.66 | 2.15 |
| 3zgc | 57.14 | 71.43 | 2.23 | 2.24 |
| 4k1e | 35.71 | 28.57 | 0.86 | 1.34 |
| 6d3x | 35.71 | 35.71 | 1.70 | 1.74 |
| 6d3y | 28.57 | 28.57 | 2.30 | 1.14 |
| 6u22 | 35.71 | 28.57 | 1.95 | 1.41 |

**Table S13.** **The 9 peptides achieving $K_D$ ranges of $10^{-8}$M among 382 non-filtering peptides designed for PD-L1.**

| Sequence | kD(M) |
|---|---|
| MHNVDFEGLI | 4.50E-08 |
| VVVAMWGVYG | 4.76E-08 |
| LDELHAFVDS | 1.08E-08 |
| PAFYDDPLNI | 3.37E-08 |
| AFLAVPVDIF | 9.48E-08 |
| SVAEWVAILG | 1.26E-09 |
| FELGFLAGAD | 1.56E-08 |
| SAAEDFWARL | 8.56E-08 |
| PAHYDDFVFA | 2.05E-08 |

**Table S14.** **Performance on small molecules docking of repeated evaluations to demonstrate the robustness and consistency of PocketXMol.**

| | Average RMSD (Å) | | | Ratio of RMSD < 2Å | | | Ratio of RMSD < 2Å & PB-valid | | |
|---|---|---|---|---|---|---|---|---|---|
| | tuned ranking | self ranking | oracle ranking | tuned ranking | self ranking | oracle ranking | tuned ranking | self ranking | oracle ranking |
| Reported | 1.293 | 1.475 | 0.652 | 83.4% | 82.5% | 96.5% | 79.4% | 78.5% | 90.7% |
| Repeat 1 | 1.294 | 1.386 | 0.645 | 84.8% | 84.6% | 96.5% | 80.1% | 81.8% | 90.7% |
| Repeat 2 | 1.316 | 1.425 | 0.646 | 84.3% | 83.9% | 96.5% | 79.7% | 81.1% | 90.4% |
| Repeat 3 | 1.397 | 1.400 | 0.662 | 83.6% | 83.2% | 96.3% | 79.4% | 80.4% | 91.8% |
| Repeat 4 | 1.315 | 1.496 | 0.638 | 84.6% | 82.5% | 97.2% | 78.7% | 78.0% | 90.7% |
| Repeat 5 | 1.300 | 1.488 | 0.649 | 85.0% | 82.0% | 96.7% | 79.4% | 78.5% | 90.4% |
| Repeat 6 | 1.317 | 1.412 | 0.643 | 85.5% | 82.2% | 97.0% | 79.0% | 78.0% | 91.1% |
| Repeat 7 | 1.398 | 1.414 | 0.634 | 83.6% | 84.1% | 96.7% | 77.8% | 80.1% | 90.7% |
| Repeat 8 | 1.334 | 1.461 | 0.653 | 83.4% | 82.2% | 96.3% | 78.0% | 78.3% | 90.0% |
| Repeat 9 | 1.335 | 1.446 | 0.646 | 84.1% | 82.2% | 96.0% | 79.7% | 78.7% | 90.0% |
| Repeat 10 | 1.339 | 1.493 | 0.641 | 84.1% | 82.5% | 97.2% | 79.4% | 79.2% | 92.3% |
| Mean | 1.331 | 1.445 | 0.646 | 84.2% | 82.9% | 96.6% | 79.2% | 79.3% | 90.8% |
| Std. | 0.037 | 0.040 | 0.008 | 0.7% | 0.9% | 0.4% | 0.7% | 1.3% | 0.7% |
| Max. | 1.398 | 1.496 | 0.662 | 85.5% | 84.6% | 97.2% | 80.1% | 81.8% | 92.3% |
| Min. | 1.293 | 1.386 | 0.634 | 83.4% | 82.0% | 96.0% | 77.8% | 78.0% | 90.0% |
| Range | 0.105 | 0.109 | 0.027 | 2.1% | 2.6% | 1.2% | 2.3% | 3.7% | 2.3% |
| 95% CI (left) | 1.259 | 1.368 | 0.631 | 79.9% | 78.9% | 94.8% | 75.6% | 74.6% | 87.9% |
| 95% CI (right) | 1.402 | 1.523 | 0.661 | 88.2% | 88.0% | 98.2% | 83.9% | 85.4% | 93.4% |

**Table S15.** **Performance on 3D conformation generation for small molecules.**

| Methods | Coverage(↑,%) | | Matching(↓,Å) | |
|---|---|---|---|---|
| | Mean | Median | Mean | Median |
| RDKit | 60.91 | 65.70 | 1.2026 | 1.1252 |
| CVGAE | 0.00 | 0.00 | 3.0702 | 2.9937 |
| GraphDG | 8.27 | 0.00 | 1.9722 | 1.9845 |
| CGCF | 53.96 | 57.06 | 1.2487 | 1.2247 |
| ConfVAE | 53.14 | 53.98 | 1.2392 | 1.2447 |
| ConfGF | 62.15 | 70.93 | 1.1629 | 1.1596 |
| GeoMol | 67.16 | 71.71 | 1.0875 | 1.0586 |
| DGSM | 78.73 | 94.39 | 1.0154 | 0.9980 |
| GeoDiff | 88.45 | 97.09 | 0.8651 | 0.8598 |
| DMCG | 91.27 | 100.00 | 0.8287 | 0.7908 |
| Uni-Mol | 91.91 | 100.00 | 0.7863 | 0.7794 |
| Torsional diffusion* (TD) | 94.95 | 100.00 | 0.6742 | 0.6318 |
| TD w/ particle guidance (recall)* | 93.91 | 98.47 | 0.6365 | 0.6115 |
| TD w/ particle guidance (precision)* | 90.23 | 95.16 | 0.6846 | 0.6724 |
| PocketXMol | 86.21 | 94.15 | 0.8734 | 0.8596 |

* These methods might use training set that had overlap with the test set here.

**Table S16.** **Performance on peptide docking of repeated evaluations to demonstrate the robustness and consistency of PocketXMol.**

| | Average DockQ | | | Median DockQ | | |
|---|---|---|---|---|---|---|
| | self ranking | tuned ranking | orcale ranking | self ranking | tuned ranking | orcale ranking |
| Reported | 0.53 | 0.58 | 0.73 | 0.55 | 0.63 | 0.78 |
| Repeat 1 | 0.54 | 0.58 | 0.74 | 0.61 | 0.63 | 0.77 |
| Repeat 2 | 0.55 | 0.59 | 0.75 | 0.61 | 0.65 | 0.80 |
| Repeat 3 | 0.52 | 0.56 | 0.73 | 0.53 | 0.62 | 0.77 |
| Repeat 4 | 0.56 | 0.58 | 0.74 | 0.62 | 0.64 | 0.78 |
| Repeat 5 | 0.49 | 0.57 | 0.74 | 0.47 | 0.62 | 0.79 |
| Repeat 6 | 0.51 | 0.57 | 0.74 | 0.53 | 0.65 | 0.76 |
| Repeat 7 | 0.50 | 0.58 | 0.74 | 0.59 | 0.65 | 0.78 |
| Repeat 8 | 0.48 | 0.56 | 0.74 | 0.47 | 0.61 | 0.80 |
| Repeat 9 | 0.54 | 0.58 | 0.73 | 0.62 | 0.65 | 0.80 |
| Repeat 10 | 0.52 | 0.60 | 0.75 | 0.54 | 0.65 | 0.80 |
| Mean | 0.52 | 0.58 | 0.74 | 0.56 | 0.63 | 0.78 |
| Std. | 0.02 | 0.01 | 0.01 | 0.06 | 0.02 | 0.01 |
| Max. | 0.56 | 0.60 | 0.75 | 0.62 | 0.65 | 0.80 |
| Min. | 0.48 | 0.56 | 0.73 | 0.47 | 0.61 | 0.76 |
| Range | 0.08 | 0.05 | 0.02 | 0.16 | 0.04 | 0.04 |
| 95% CI (left) | 0.47 | 0.55 | 0.73 | 0.45 | 0.60 | 0.76 |
| 95% CI (right) | 0.57 | 0.60 | 0.75 | 0.67 | 0.66 | 0.81 |

**Table S17.** DockQ of cyclic peptide docking for all data in the test set.

| PDB ID | Cluster ID | Alpha Fold 2 | HighFold | AfCyc Design | PocketX Mol (self ranking) | PocketX Mol (tuned ranking) | PocketX Mol (oracle ranking) |
|---|---|---|---|---|---|---|---|
| 3wne | 0 | 0.93 | 0.93 | 0.50 | 0.59 | 0.86 | 0.95 |
| 3wng | 0 | 0.04 | 0.91 | 0.90 | 0.78 | 0.56 | 0.96 |
| 1sfi | 1 | 0.98 | 0.98 | 0.98 | 0.96 | 0.96 | 0.96 |
| 6bvh | 1 | 0.97 | 0.97 | 0.96 | 0.95 | 0.93 | 0.97 |
| 4k1e | 2 | 0.94 | 0.94 | 0.84 | 0.86 | 0.83 | 0.94 |
| 4kel | 2 | 0.94 | 0.94 | 0.83 | 0.87 | 0.86 | 0.88 |
| 6d3x | 3 | 0.80 | 0.80 | 0.80 | 0.96 | 0.94 | 0.97 |
| 6d3z | 3 | 0.90 | 0.91 | 0.90 | 0.93 | 0.93 | 0.97 |
| 6q1u | 3 | 0.78 | 0.78 | 0.53 | 0.96 | 0.96 | 0.96 |
| 3av9 | 4 | 0.92 | 0.92 | 0.91 | 0.76 | 0.92 | 0.93 |
| 3ava | 4 | 0.92 | 0.92 | 0.91 | 0.73 | 0.75 | 0.84 |
| 3avb | 4 | 0.91 | 0.91 | 0.93 | 0.71 | 0.52 | 0.76 |
| 3avf | 4 | 0.90 | 0.90 | 0.89 | 0.77 | 0.67 | 0.88 |
| 3avg | 4 | 0.91 | 0.92 | 0.91 | 0.76 | 0.39 | 0.91 |
| 3avh | 4 | 0.85 | 0.85 | 0.91 | 0.62 | 0.72 | 0.89 |
| 3avi | 4 | 0.82 | 0.82 | 0.85 | 0.45 | 0.66 | 0.73 |
| 3avj | 4 | 0.81 | 0.85 | 0.82 | 0.47 | 0.50 | 0.72 |
| 3avk | 4 | 0.86 | 0.86 | 0.87 | 0.48 | 0.47 | 0.73 |
| 3avm | 4 | 0.88 | 0.88 | 0.87 | 0.69 | 0.72 | 0.88 |
| 3avn | 4 | 0.90 | 0.89 | 0.90 | 0.74 | 0.72 | 0.85 |
| 6d3y | 5 | 0.80 | 0.80 | 0.80 | 0.97 | 0.96 | 0.97 |
| 6d40 | 5 | 0.80 | 0.80 | 0.82 | 0.94 | 0.94 | 0.99 |
| 3p8f | 6 | 0.94 | 0.94 | 0.95 | 0.96 | 0.88 | 0.96 |
| 3zgc | 7 | 0.59 | 0.59 | 0.97 | 0.43 | 0.95 | 1.00 |
| 6u22 | 8 | 0.97 | 0.97 | 0.55 | 0.96 | 0.97 | 0.97 |
| 6vxy | 8 | 0.95 | 0.95 | 0.55 | 0.94 | 0.93 | 0.97 |

**Table S18. Statistic of enzyme databases.** Here, the valid proteins are those enzymes with at least five positive or negative substrates in the datasets and were used to analyze the performance of PocketXMol+LR for activity prediction. Reference PDBs are the structures used for defining and aligning the enzyme active pockets.

| Enzyme family | Unique proteins | Unique substrates | Unique pairs | Valid proteins | Reference PDB |
|---|---|---|---|---|---|
| Esterases | 132 | 96 | 12,672 | 96 | 5A6V |
| Glycosyltransferases | 54 | 89 | 4,297 | 48 | 3HBF |
| Halogenases | 42 | 62 | 2,604 | 17 | 2AR8 |
| Phosphatases | 218 | 165 | 35,970 | 102 | 3L8E |

### Supplementary Figures

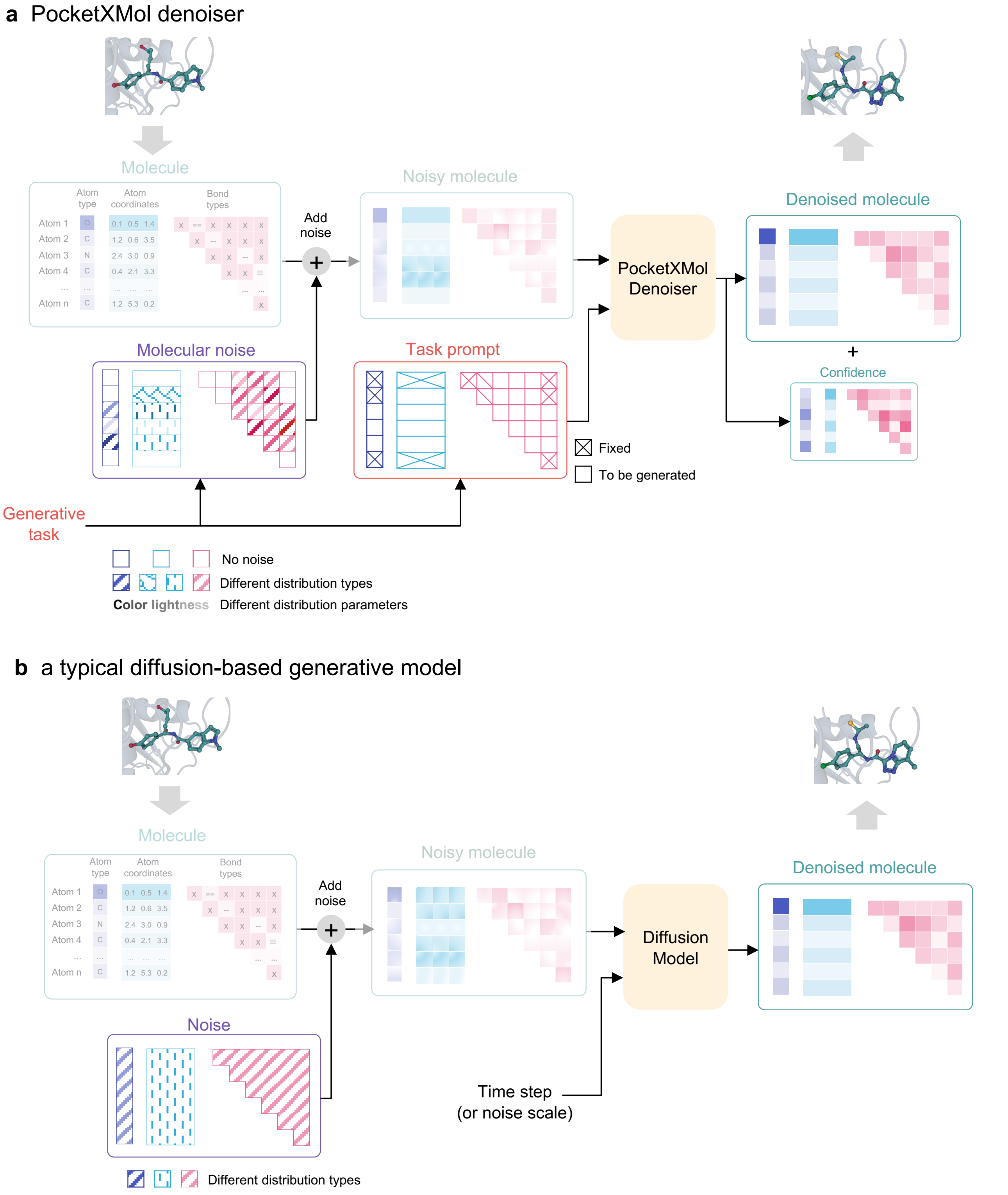

**Figure S7. Comparison of the denoisers between PocketXMol and the diffusion-based generative model. a**, Denoiser of PocketXMol. **b**, Denoiser of a typical diffusion-based generative model.

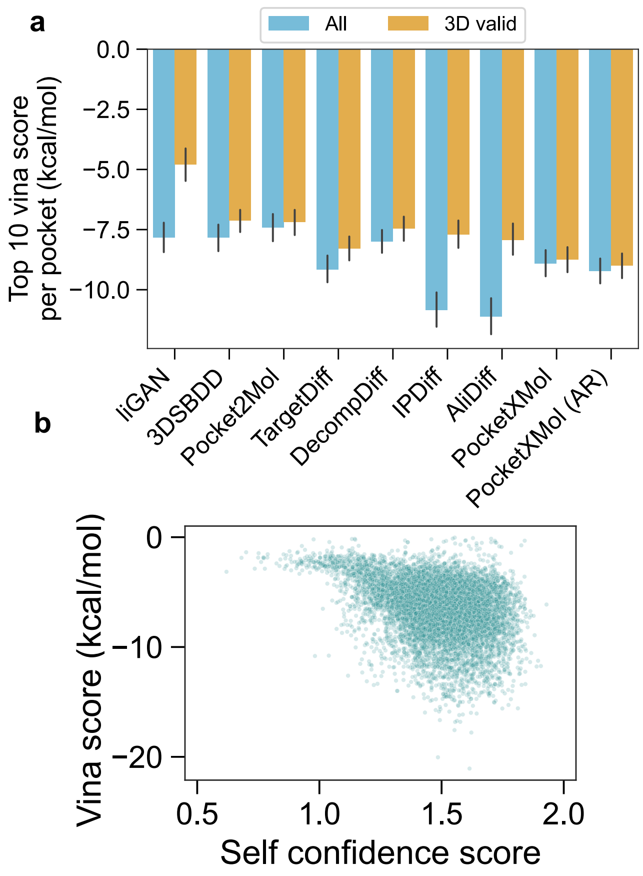

**Figure S8. Extended results of Vina scores for the SBDD task. a**, average over top 10 Vina scores per protein pocket by different methods and filter criteria. The filter criteria include no filtering (all) and passing PoseBusters 3D validity check (3D valid). **b**, Correlation between the Vina scores and the self confidence scores of molecules generated by PocketXMol.

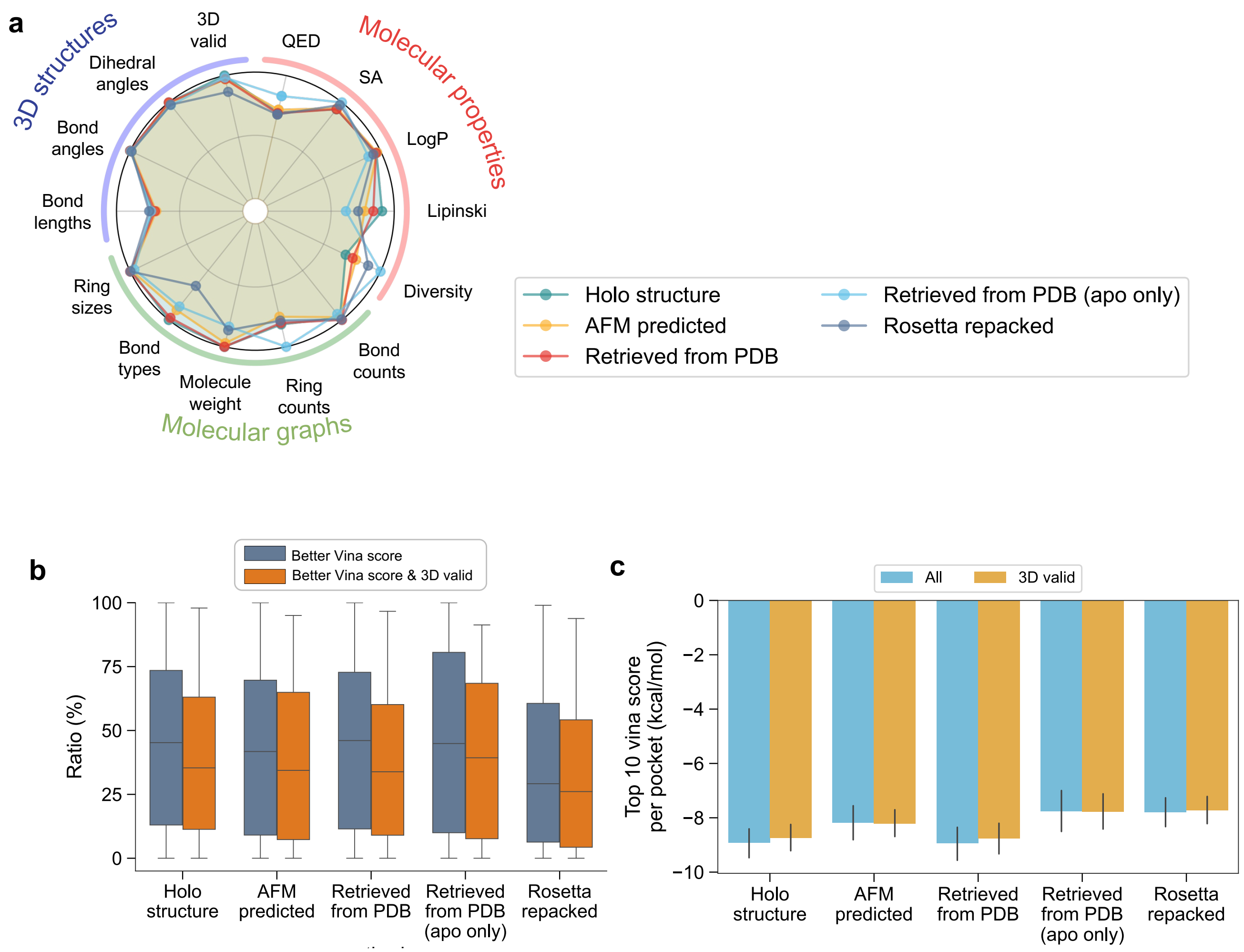

**Figure S9. Performance of PocketXMol on the SBDD task with different input pocket structures. a**, Basic metrics of PocketXMol with different input pocket structures. Each metric on the radar plot was normalized to a scale from 0 to 1. The normalizing scales are the same as the corresponding radar plot of the baselines in the main text. **b**, Ratios of the generated molecules exhibiting both better Vina scores and 3D validity for each pocket. Better Vina score is defined as those with Vina scores better than the reference molecule in the test set. The 3D validity was annotated by the PoseBusters validity checker. **c**, average Vina scores per protein pocket by PocketXMol with different input pocket structures and filter criteria. The filter criteria include no filtering (all) and passing PoseBusters 3D validity check (3D valid).

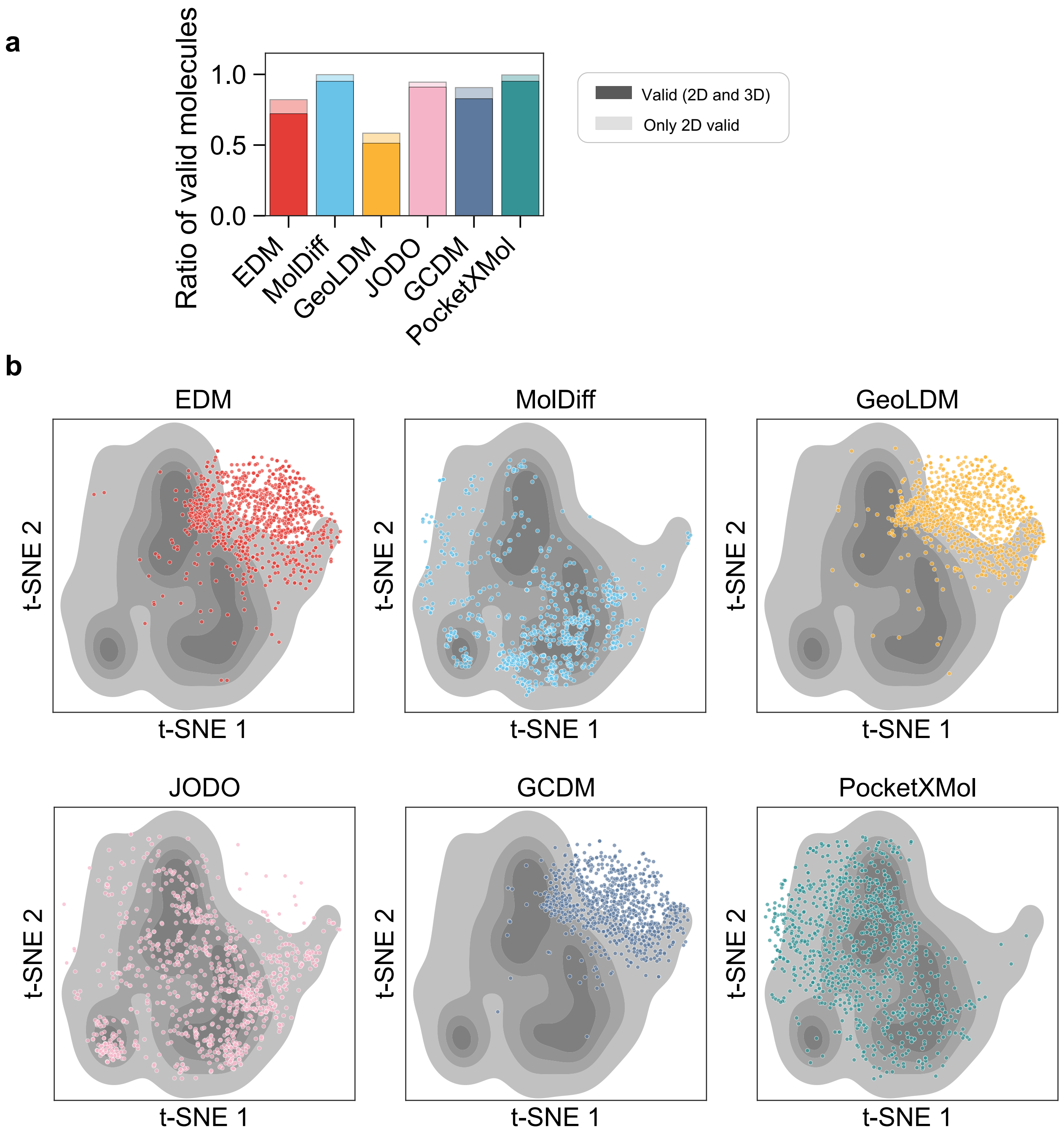

**Figure S10.** **Comparison of different models for pocket-free 3D small molecule generation.** **a**, Ratio of valid molecules for 3D molecule generation. Here, the 2D validity was defined as complete molecules by RDKit, and 3D validity was annotated by the PoseBusters. **b**, t-SNE visualization depicting the molecular fingerprints of the real molecules in the GEOM-Drug dataset and molecules generated by different 3D molecule generation methods. The grey density map of each subfigure represents the real molecules, and the colored scatters represent molecules generated by different methods.

**a**

| Models | Generate linker positions | Unknown connecting atoms | Pocket-aware | Multiple fragments | Unknown fragment positions |
|---|---|---|---|---|---|
| DeLinker | | | | | |
| 3DLinker | √ | √ | | | |
| DiffLinker | √ | √ | √ | √ | |
| LinkerNet | √ | √ | | | √ |
| PocketXMol | √ | √ | √ | √ | √ |

**b** **c**

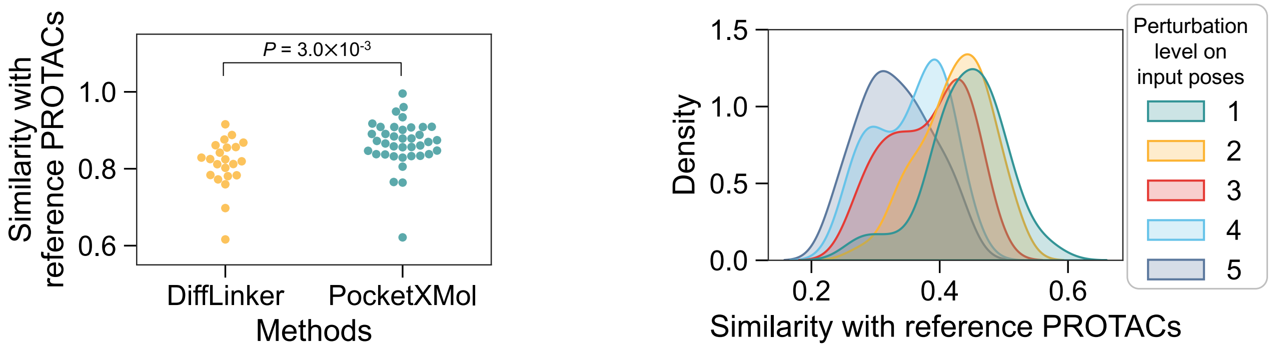

**Figure S11. Extended results for molecular linker design tasks (fragment linking and PROTAC design). a**, Comparison of the functionality of different fragment linking models. **b**, Similarities between the true PROTACs and those generated by DiffLinker and PocketXMol. $P$ value was calculated using one-sided paired $t$-test with $n = 18$. **c**, Similarities between the true PROTACs and those generated by PocketXMol for different perturbation levels.

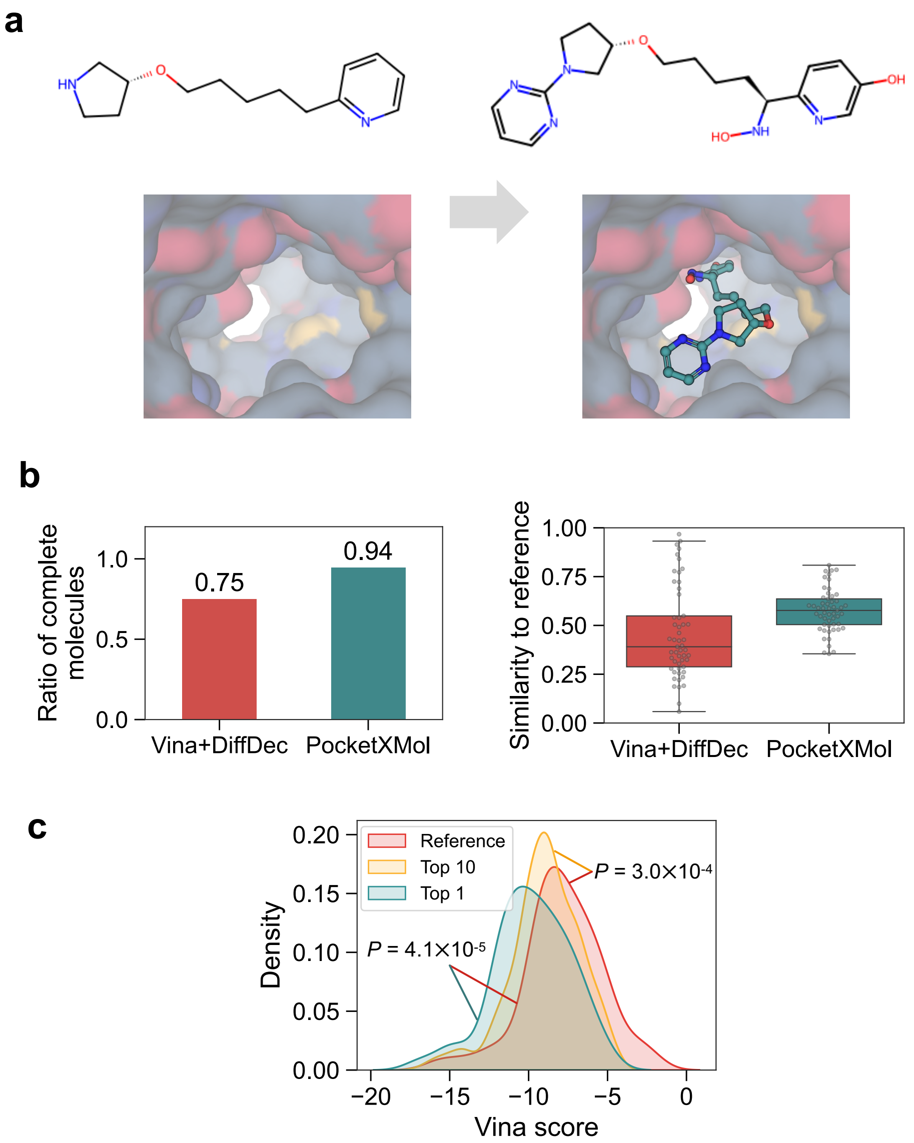

**Figure S12. Extended results for fragment growing. a**, An example showing a small molecule growing from an initial fragment based on the 3D pocket. The inputs are the fragment graph (without 3D poses) and the pocket structure and the outputs are the complex structure of the pocket and the generated molecules. **b**, Comparison between the baseline Vina+DiffDec and PocketXMol on the fragment growing task. The ratio of complete molecules among generated ones (left) and the similarity between the generated molecules and the reference molecule in the test set (right) are shown. **c**, Vina score distributions of the reference molecules and the molecules generated by PocketXMol in the fragment growing task. Reference molecules were the complete molecules used to derive the input fragment in the test set. The average Vina scores of the top 1 and the top 10 generated molecules for each input fragment were displayed. $P$ value was calculated using one-sided paired t-test with $n = 53$.

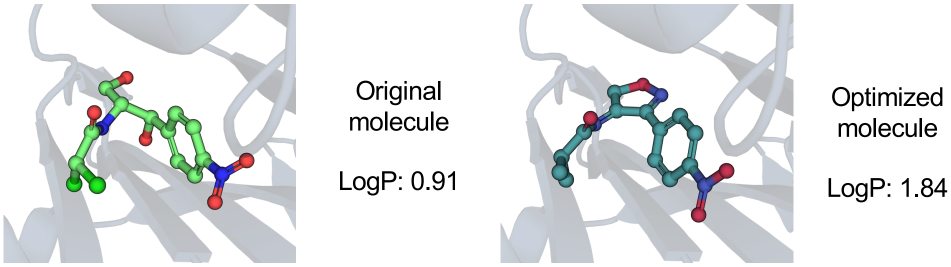

**Figure S13.** An example for molecule optimization.

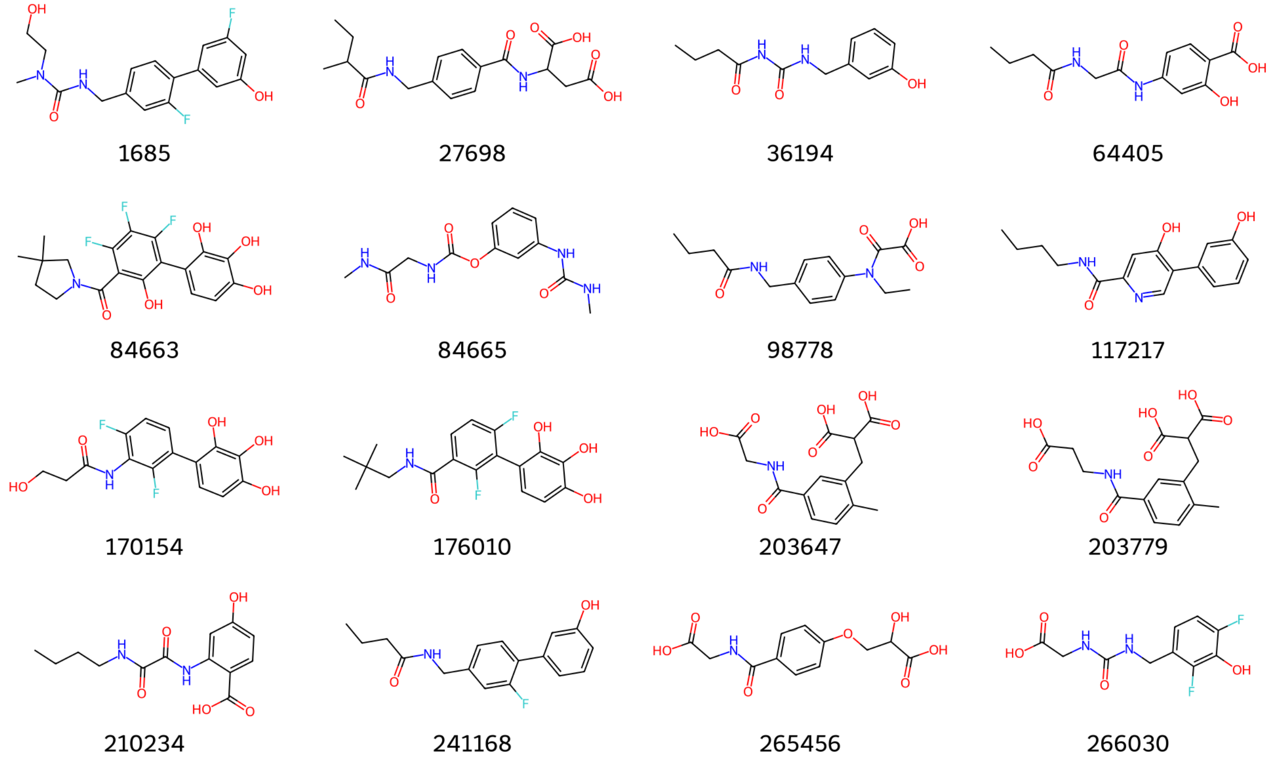

**Figure S14.** The 16 synthesized molecules designed by PocketXMol for caspase-9.

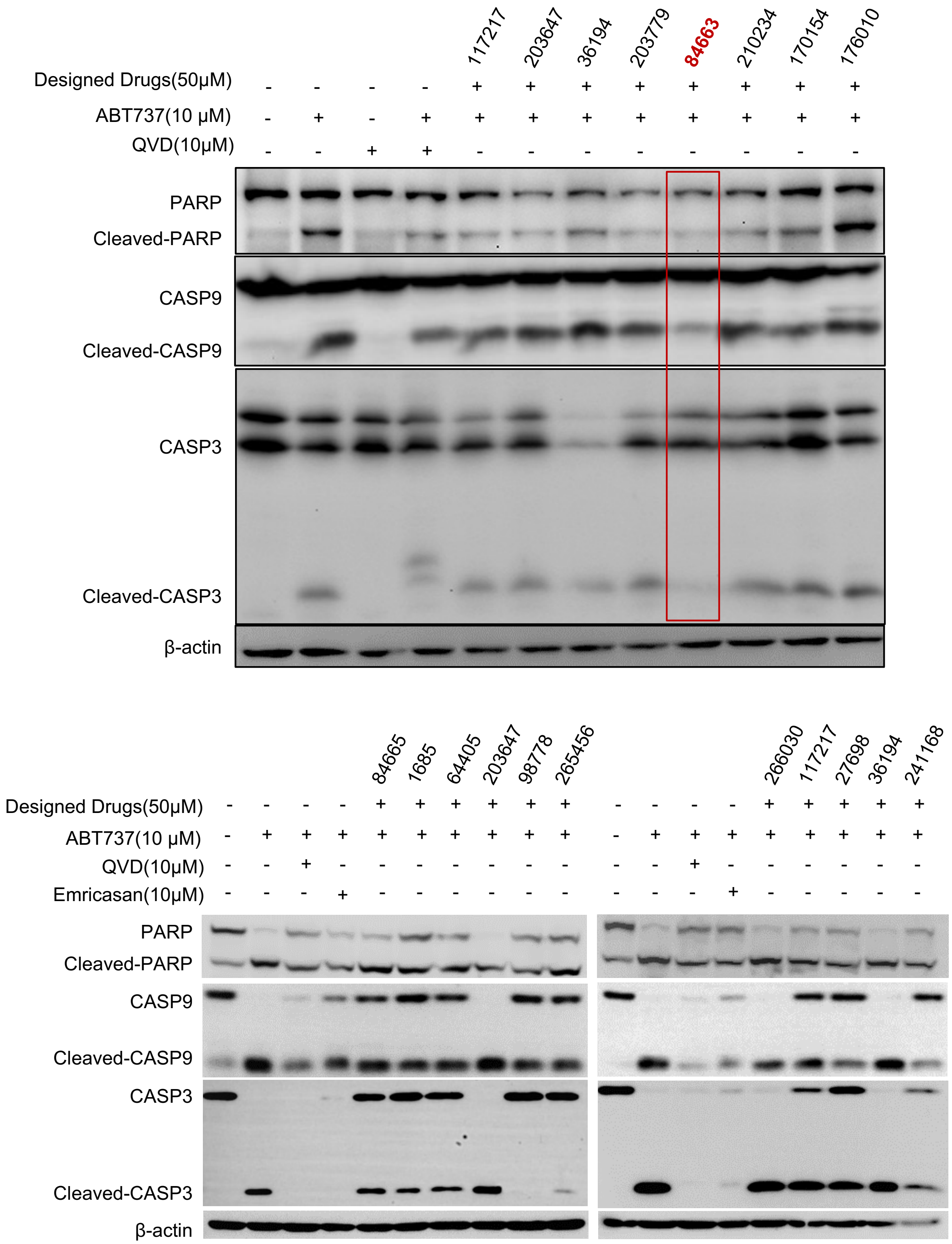

**Figure S15. Western blot analysis of the expression of PARP and caspase-9/3 with all the synthesized molecules.** MC38 cells were treated with QVD(10 µM), and other inhibitors (50 µM) combined with ABT-737 (10 µM) or not. Emricasan is a pan-caspases inhibitor and was used as additional reference inhibitors in some experiments.

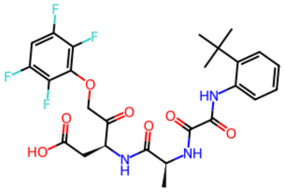

Similarity: 0.36
Emricasan

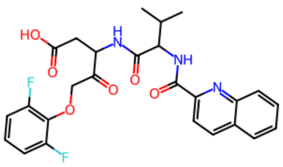

Similarity: 0.40
Q-VD-Oph

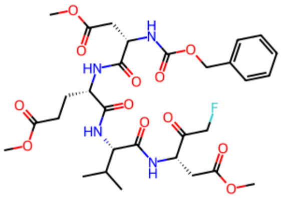

Similarity: 0.30
Z-DEVD-FMK

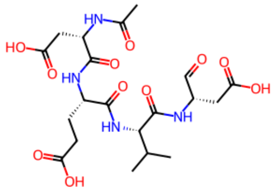

Similarity: 0.19
Ac-DEVD-CHO

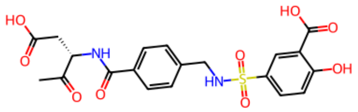

Similarity: 0.36
5-[4-(1-Carboxymethyl-2-Oxo-Propylcarbamoyl)-Benzylsulfamoyl]-2-Hydroxy-Benzoic Acid

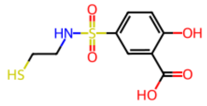

Similarity: 0.26
2-HYDROXY-5-(2-MERCAPTO-ETHYLSULFAMOYL)-BENZOIC ACID

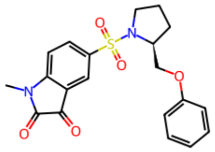

Similarity: 0.41
1-METHYL-5-(2-PHENOXYMETHYL-PYRROLIDINE-1-SULFONYL)-1H-INDOLE-2,3-DIONE

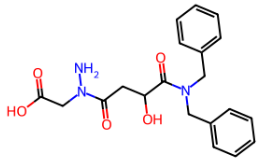

Similarity: 0.28
[N-(3-dibenzylcarbamoyl-oxiranecarbonyl)-hydrazino]-acetic acid

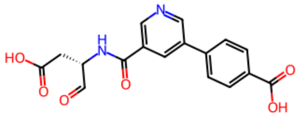

Similarity: 0.35
4-[5-(2-CARBOXY-1-FORMYL-ETHYLCARBAMOYL)-PYRIDIN-3-YL]-BENZOIC ACID

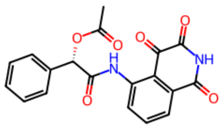

Similarity: 0.41
(1S)-2-oxo-1-phenyl-2-[(1,3,4-trioxo-1,2,3,4-tetrahydroisoquinolin-5-yl)amino]ethyl acetate

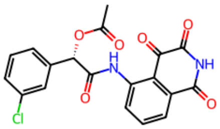

Similarity: 0.42
(1S)-1-(3-chlorophenyl)-2-oxo-2-[(1,3,4-trioxo-1,2,3,4-tetrahydroisoquinolin-5-yl)amino]ethyl acetate

**Figure S16. The common caspase inhibitors and their similarities with 84663.** The similarities were calculated from the molecular fingerprint using RDKit.

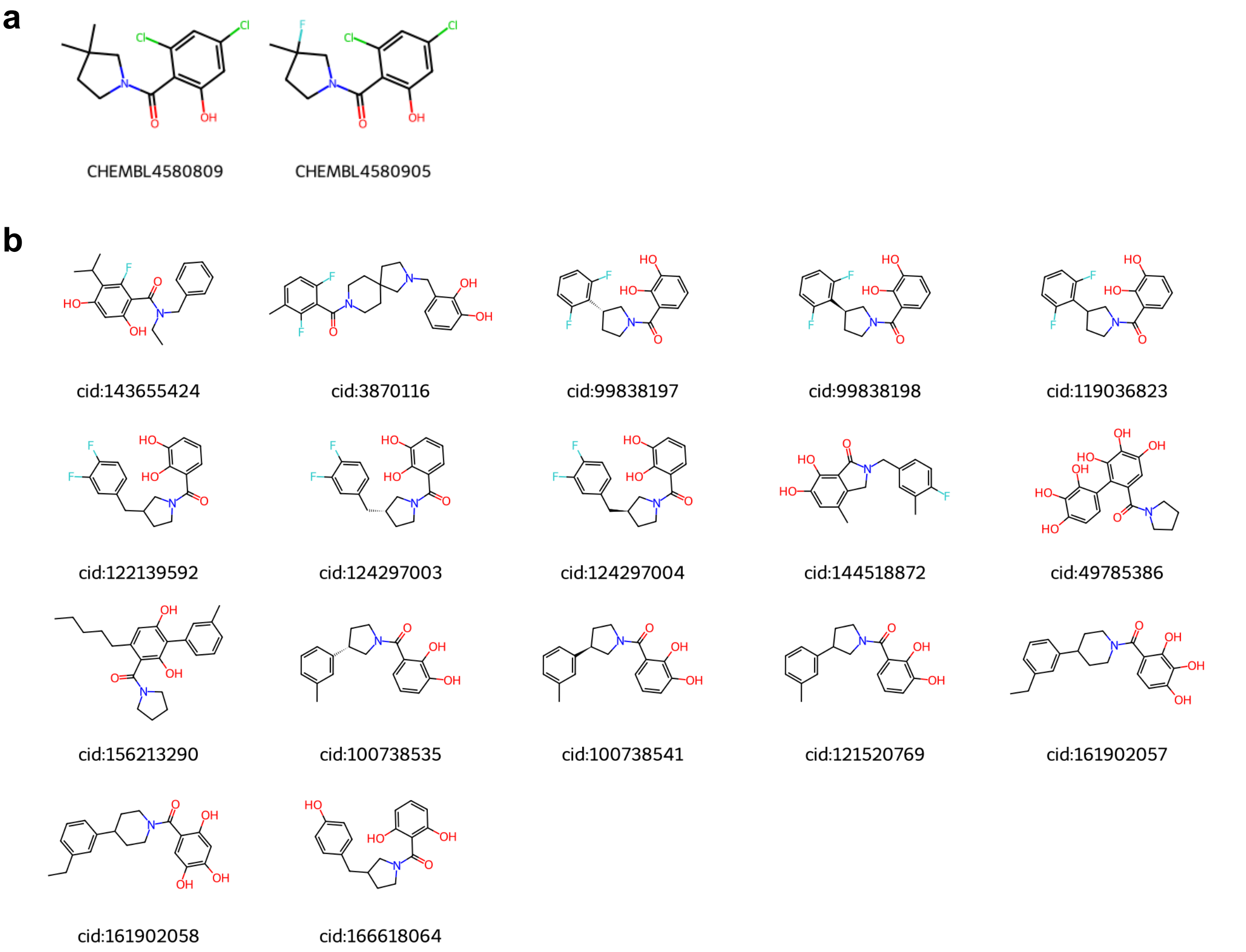

**Figure S17. The most similar molecules with 84663 found in different chemical databases. a**, The most similar molecules with 84663 in the ChEMBL database, with a similarity threshold of 40%. The similarities were calculated by the ChEMBL database. **b**, The most similar molecules with 84663 in the PubChem database, with a similarity threshold of 86%. The similarities were calculated by the PubChem database. Note that the similarity values calculated by different databases may not be comparable.

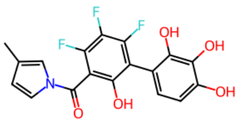

D9
+

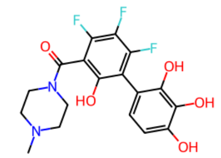

D11
-

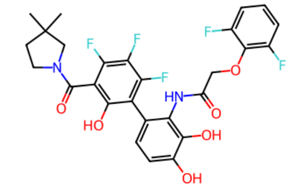

D12
+++

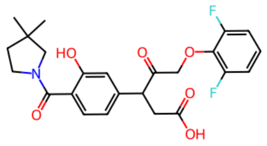

D16
-

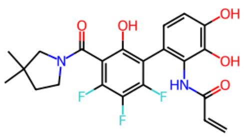

D18
++

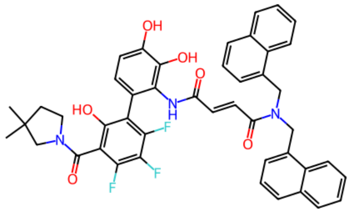

D19
++++

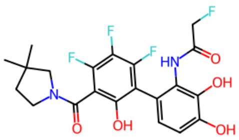

D20
+

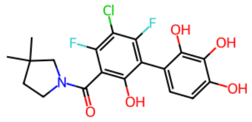

D1
+

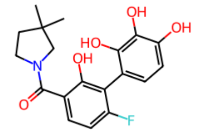

D3
+

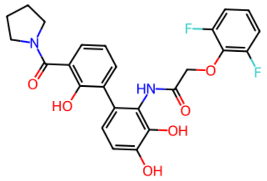

D13
++

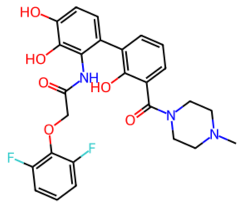

D14
-

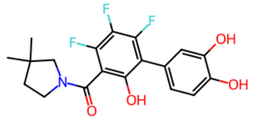

D4
+

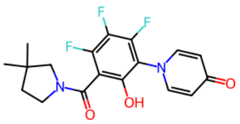

D5
-

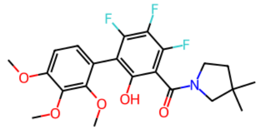

D7
-

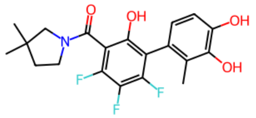

D6
+

**Figure S18.** The molecules that were manually designed and synthesized based on 84663 and their experimental results.

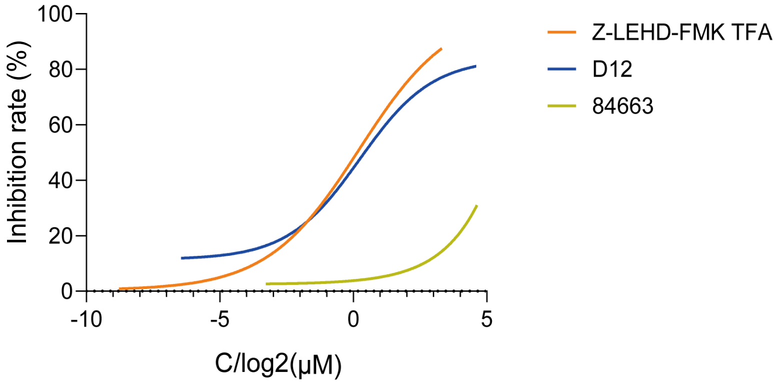

**Figure S19. Caspase-3 activity assay for D12 and 84663.** This was to determine the $EC_{50}$ values of D12, 84663, and a validated caspase-9 inhibitor Z-LEHD-FMK TFA for suppressing the ABT-737 induced caspase-9 mediated caspase-3 activation.

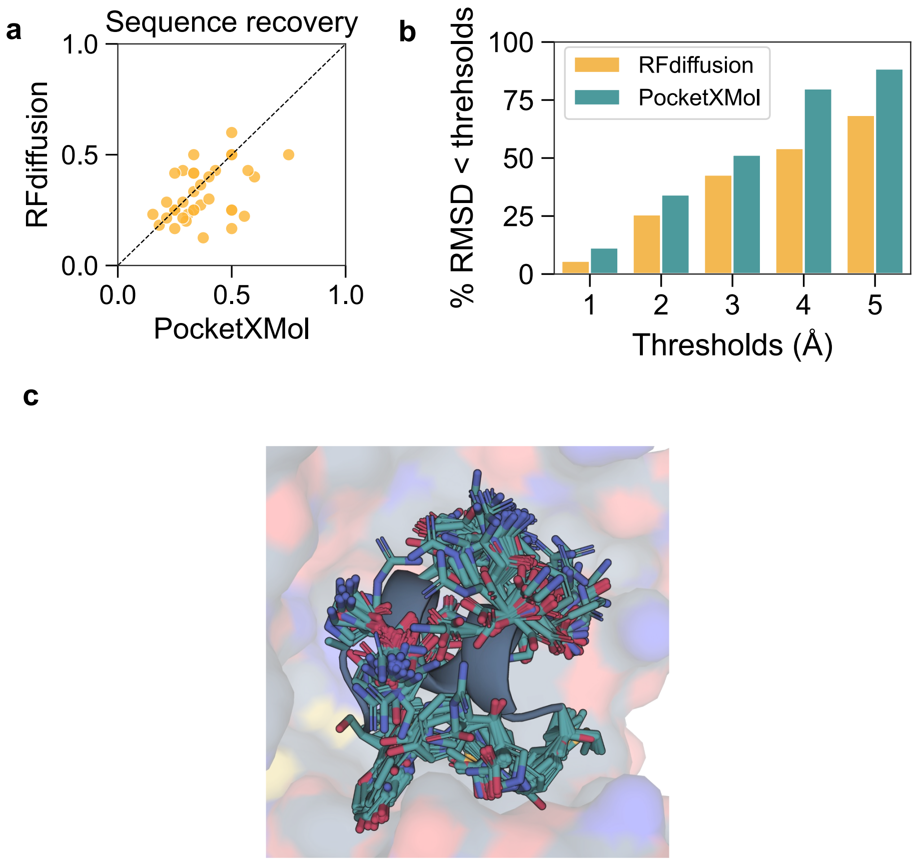

**Figure S20.** **Extended results for peptide design.** **a**, Comparison of the sequence recovery rates between the RFdiffusion pipeline and PocketXMol. For both methods, the peptides with the highest sequence recovery rates were selected for comparison among the 100 generated samples for each pocket. **b**, Ratios of peptides with backbone atom RMSD less than thresholds for RFdiffusion and PocketXMol. The minimum backbone RMSD among 100 generated peptides for each pocket was selected for comparison. **c**, Structures of 100 peptides generated by PocketXMol for a protein pocket (PDB ID: 6NWE) in the peptide inverse folding task.

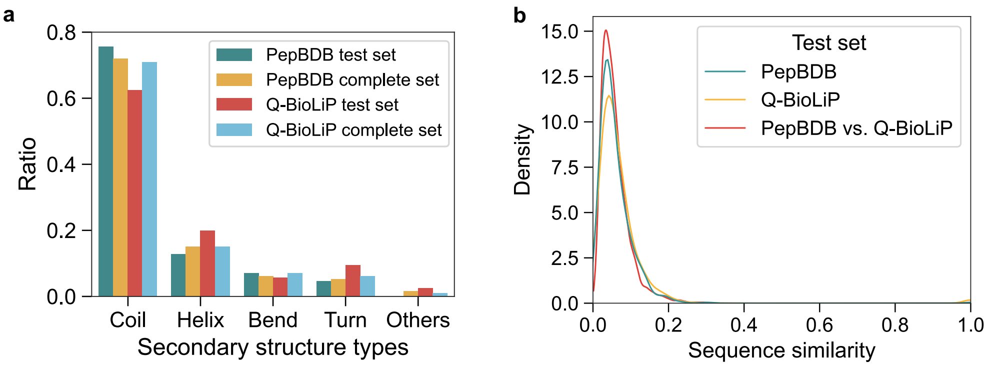

**Figure S21.** **Distributions of secondary structure types of the PepBDB peptide test set and the Q-BioLiP peptide test set.**

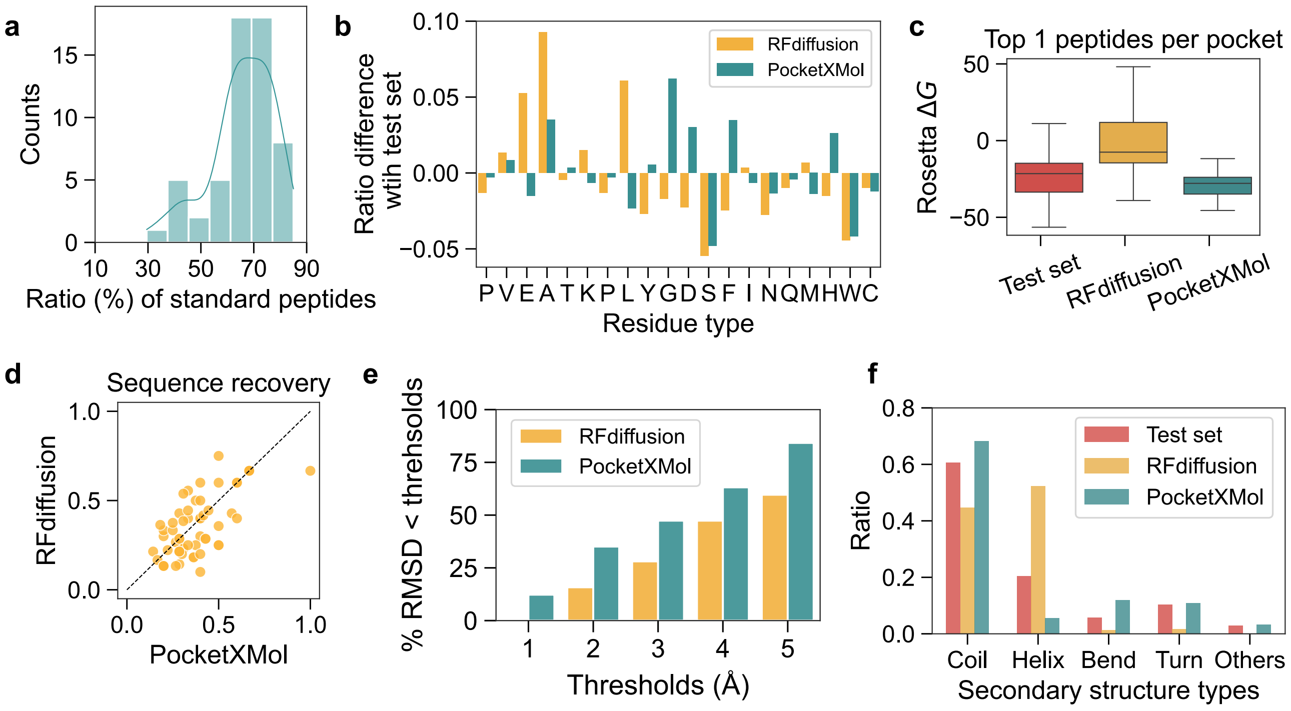

**Figure S22.** **The performance on peptide design on the Q-BioLiP test set.** **a**, Ratio distribution of standard peptides (i.e., peptides containing only standard amino acid types) generated by PocketXMol for each pocket in the test set. **b**, Ratio differences of amino acid types between the generated peptides and the test set. **c**, Distributions of the Rosetta binding energy for the test set, RFdiffusion, and PocketXMol. The binding energy was calculated using the Rosetta Interface Analyzer, and the peptides with the lowest Rosetta energies for each pocket were selected for comparison (one-sided paired $t$-test with $n = 57$). **d**, Comparison of the sequence recovery rates between the RFdiffusion pipeline and PocketXMol. For both methods, the peptides with the highest sequence recovery rates were selected for comparison among the 100 generated samples for each pocket. **e**, Ratios of peptides with backbone atom RMSD less than thresholds for RFdiffusion and PocketXMol. The minimum backbone RMSD among 100 generated peptides for each pocket was selected for comparison. **f**, Ratio of secondary structures in the peptides from the test set, RFdiffusion and PocketXMol.

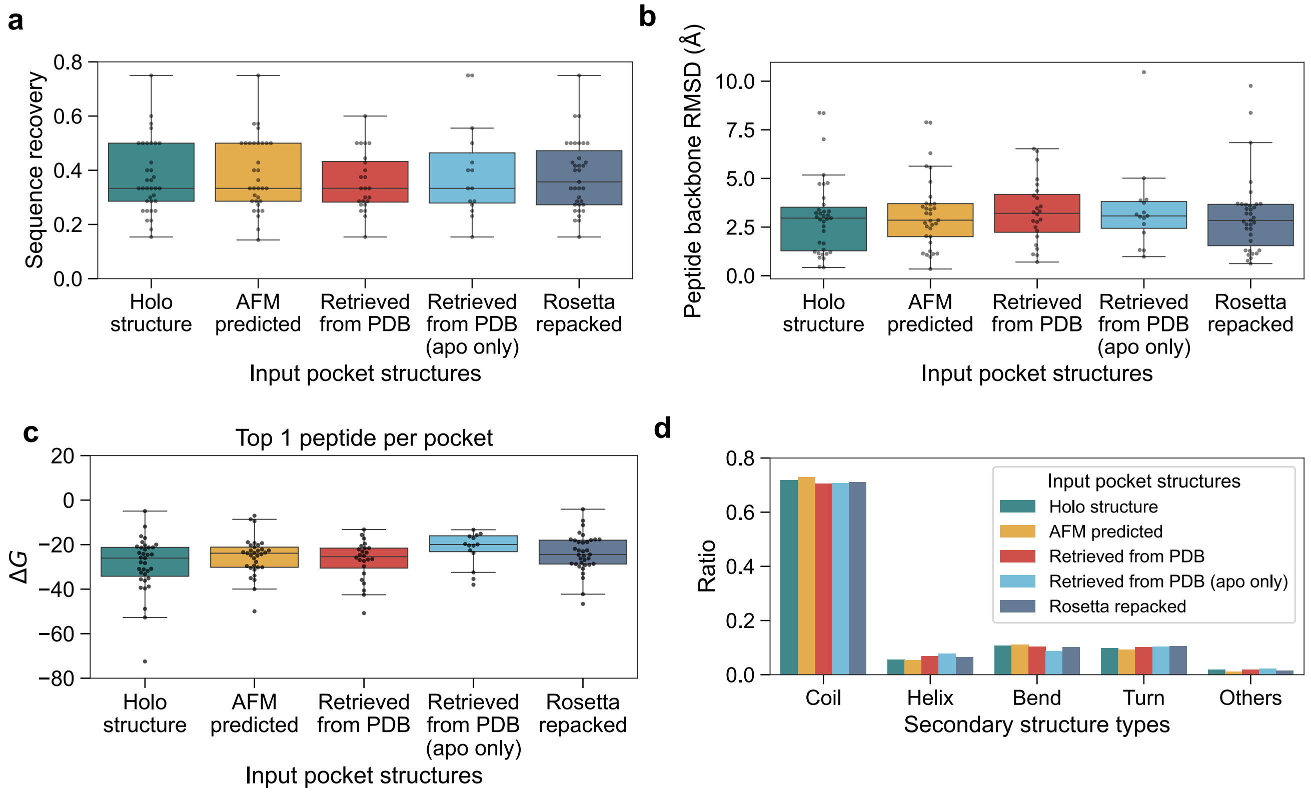

**Figure S23. Performance on *de novo* peptide design of PocketXMol with different input pocket structures (The true holo structures, AlphaFold-Multimer predicted protein structures, all structures of the same sequences retrieved from RCSB PDB, apo structures of the same sequences retrieved from RCSB PDB, and the true structures with side-chain repacked by the Rosetta).** **a**, Sequence recoveries for different input pocket structures. Peptides with the highest sequence recovery rates among the 100 generated samples for each pocket were selected for comparison. **b**, Peptide backbone RMSDs for different input pocket structures. The minimum backbone RMSDs among 100 generated peptides for each pocket were selected for comparison. **c**, Top 1 Rosetta binding energies per pocket for different input pocket structures. **d**, Distributions of secondary structure types for different input pocket structures.

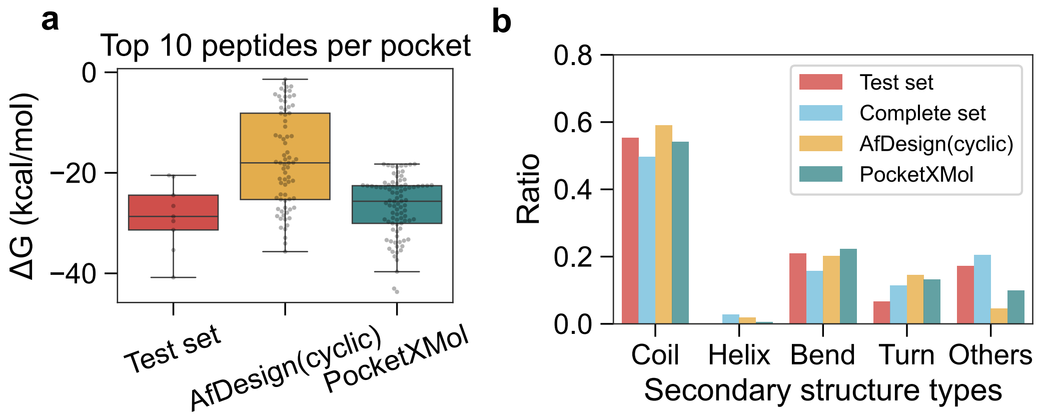

**Figure S24. The Rosetta binding energies and the distributions of secondary structure for the cyclic peptides in the datasets or generated by different models. a**, Top 10 Rosetta binding energies of the generated cyclic peptides per pocket and the Rosetta binding energies of those in the cyclic peptide test set. The $P$ value of one-sided $t$-test is $2.18 \times 10^{-10}$ with $n = 90$ for the comparison between PocketXMol and AfDesign(cyclic). **b**, Distributions of secondary structure types of cyclic peptides in the cyclic peptide test set, cyclic peptide complete set, and generated by AfDesign(cyclic) or PocketXMol.

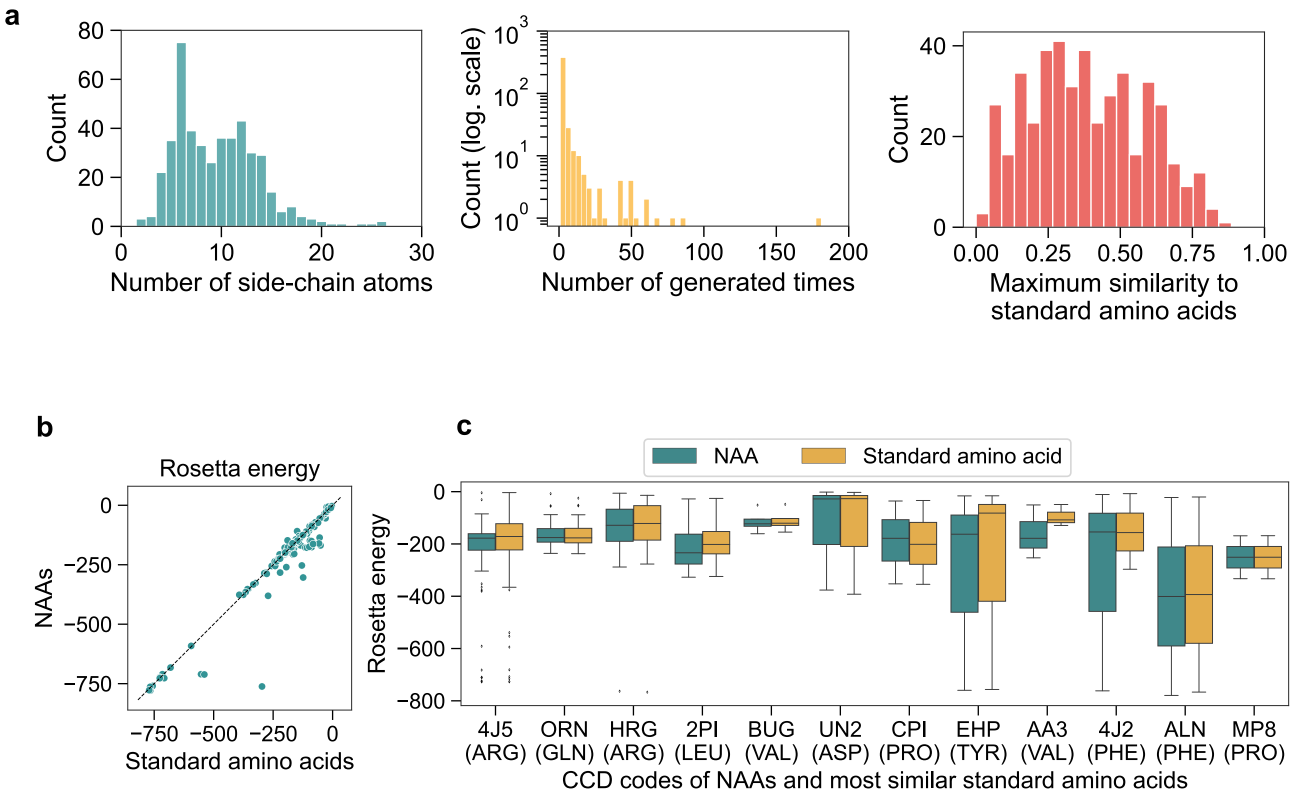

**Figure S25. Properties and Rosetta energy of NAAs.** (**a**) Distributions of the number of side-chain atoms, the number of generated times, and the maximum similarity to standard amino acids of the NAAs generated by PocketXMol. For peptides containing NAAs, we collected the side chains of all NAAs and calculated the counts of individual unique NAAs. We then calculated the Tanimoto similarity of the topological fingerprints of the side chains between these NAAS and 19 standard amino acids (Glycine was excluded because it did not contain heavy atoms in side chains) to get the maximum similarity to standard amino acids of NAAs. (**b**) Comparison of the Rosetta energy between the peptides with NAAs and peptides by mapping NAAs to their most similar standard amino acids. Here, only the NAAs that can be mapped to known CCD codes in the Rosetta residue databases were considered because they can be directly processed by Rosetta with consistent energy terms. For each peptide with NAAs, we made its standard version by replacing one NAA with its most similar amino acid and packing the side chain using Rosetta. To make a fair comparison, we also re-packed the peptides with NAAs before calculating the Rosetta energy. Finally, the interface analyzer of Rosetta was used to calculate the Rosetta energy of the complexes formed by these peptides. (**c**) Distributions of the Rosetta energy between the peptides with NAAs and peptides by mapping NAAs to their most similar standard amino acids of most frequent NAAs. The x-axis is the CCD codes of the NAAs and the most similar standard amino acids.

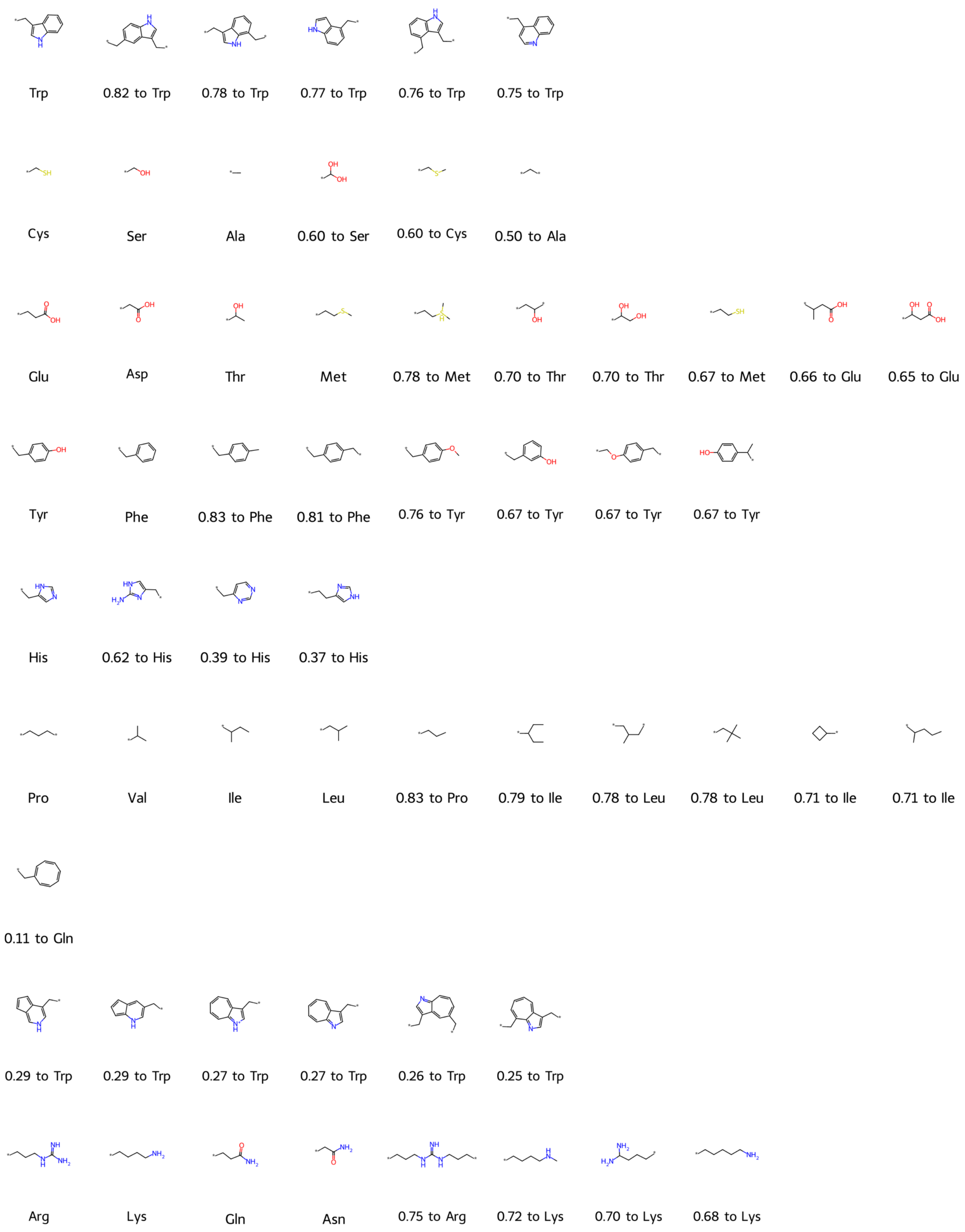

**Figure S26. Side chains of standard amino acids and some frequent NAAs generated by PocketXMol.** Each row represents a cluster group. Standard amino acids are annotated as their three-letter names while NAAs are annotated as the similarities with the most similar standard amino acids. At most ten amino acids per cluster were shown, and NAAs that appeared only once were excluded from the display. The similarity was calculated as the Tanimoto similarity of the topological fingerprints of the side-chain fragments.

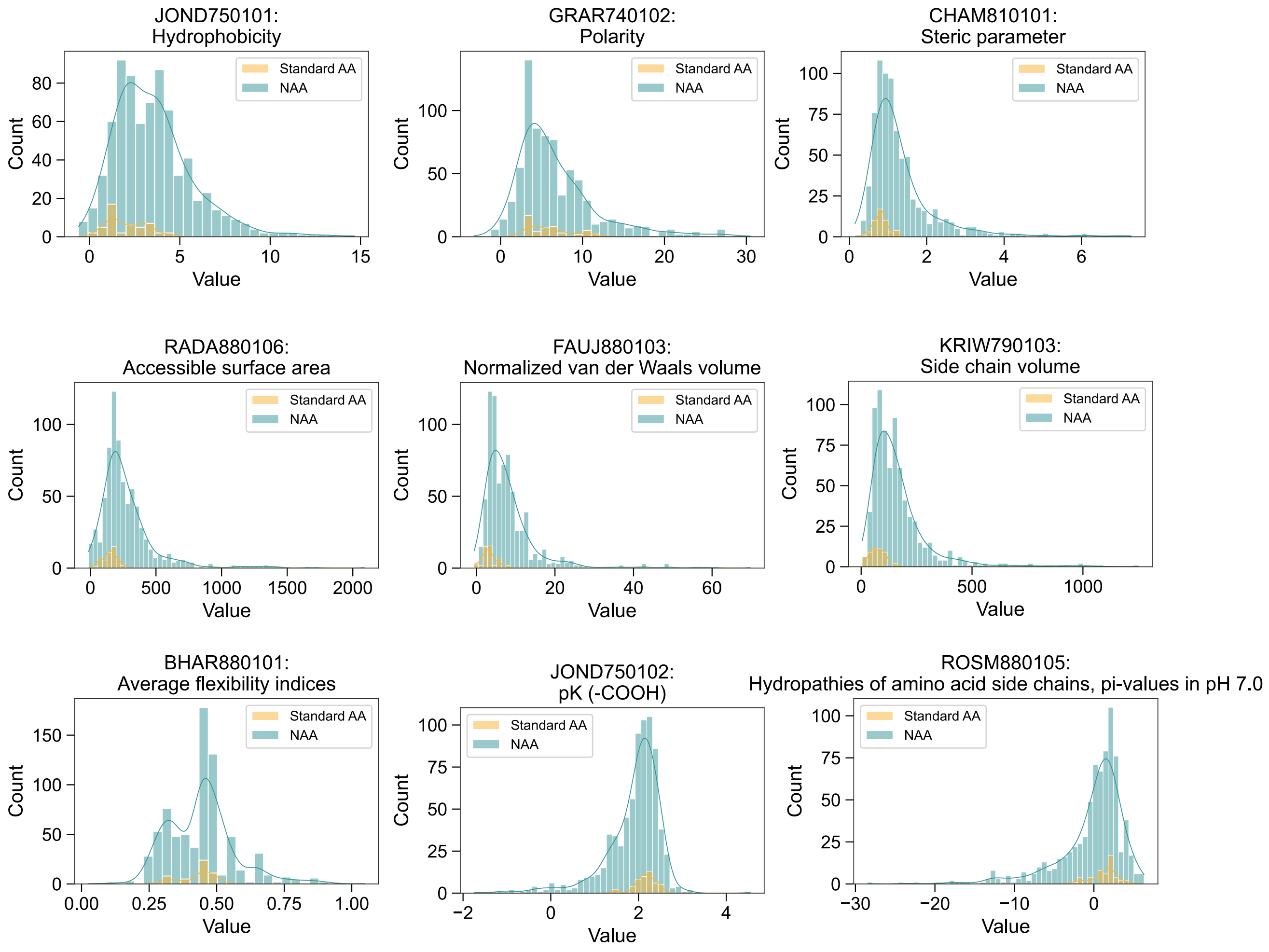

**Figure S27.** Some physical and chemical properties of the standard amino acids and NAAs calculated by AAindexNC.

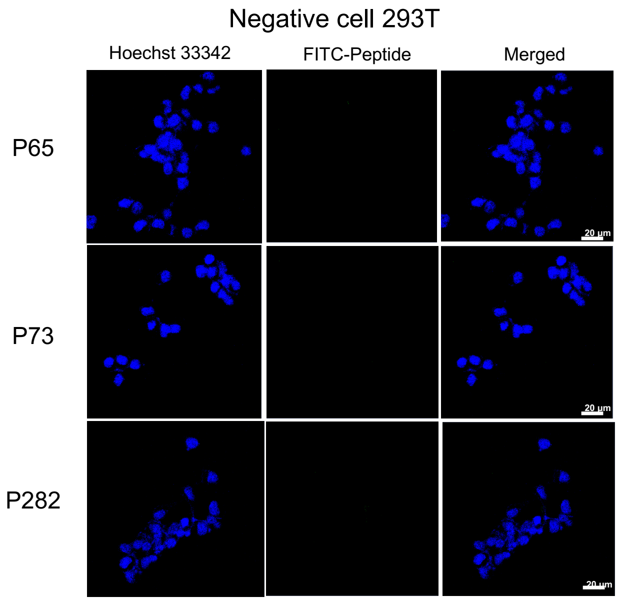

**Figure S28.** Confocal image of FITC labeled peptides binding toward negative 293T cells.

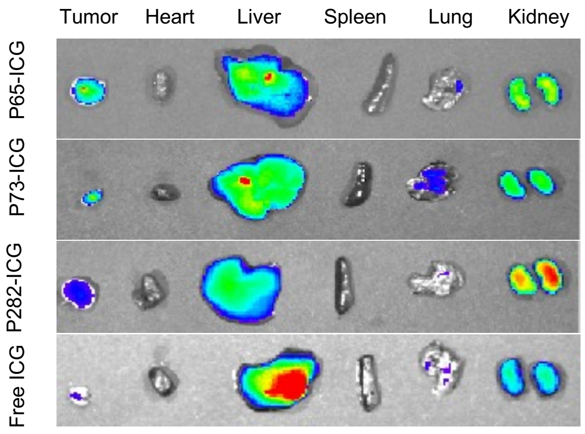

**Figure S29.** Ex vivo images of tumors and main organs (heart, liver, spleen, lung, kidney) that were taken 12 hours after post-injection.

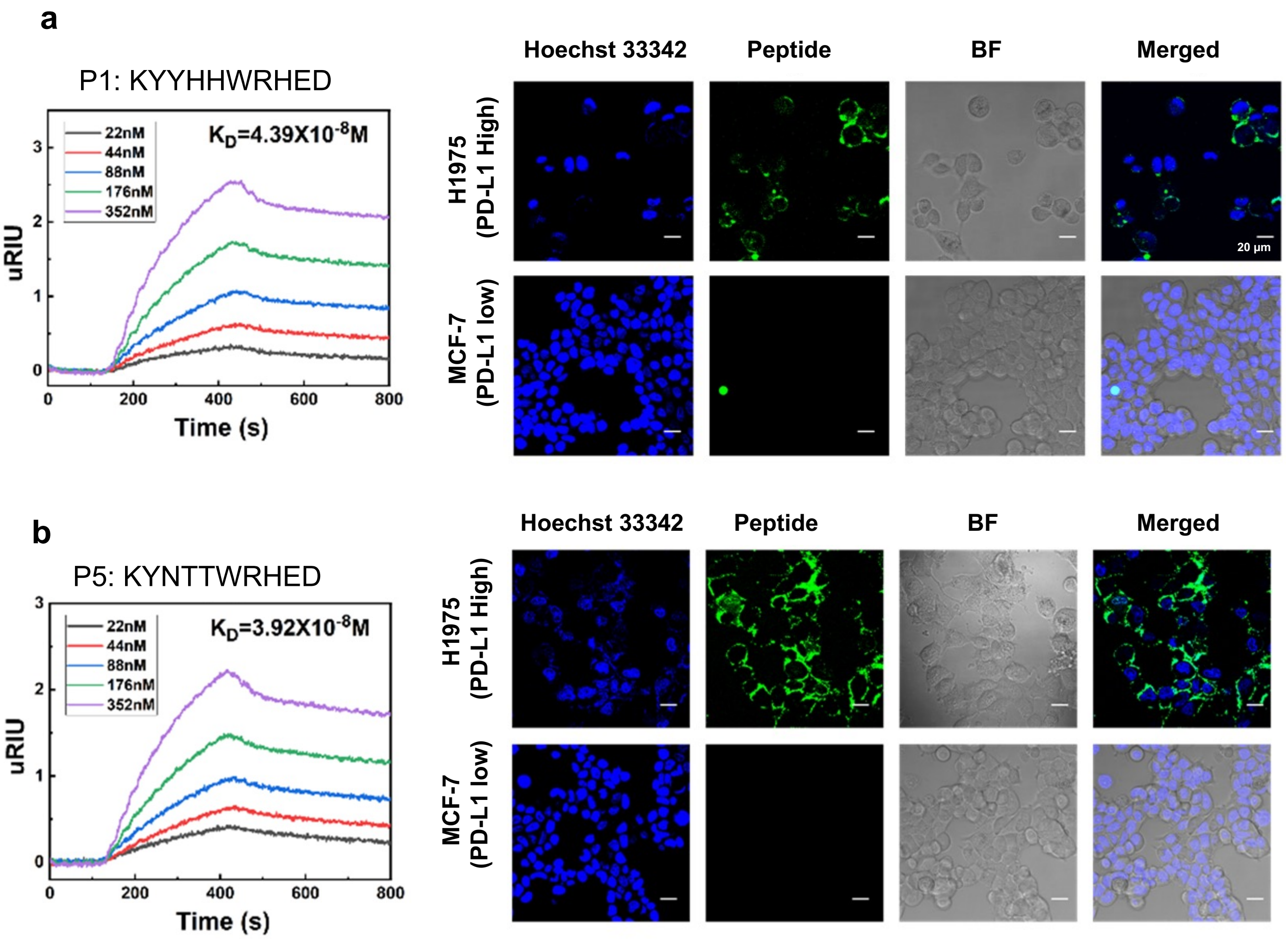

**Figure S30.** The SPRi binding curves and the confocal image of FITC-labeled peptides binding with cells for the two positive peptides obtained through library screening: **a**, P1 and **b**, P5. Scale bar is 20 μm.

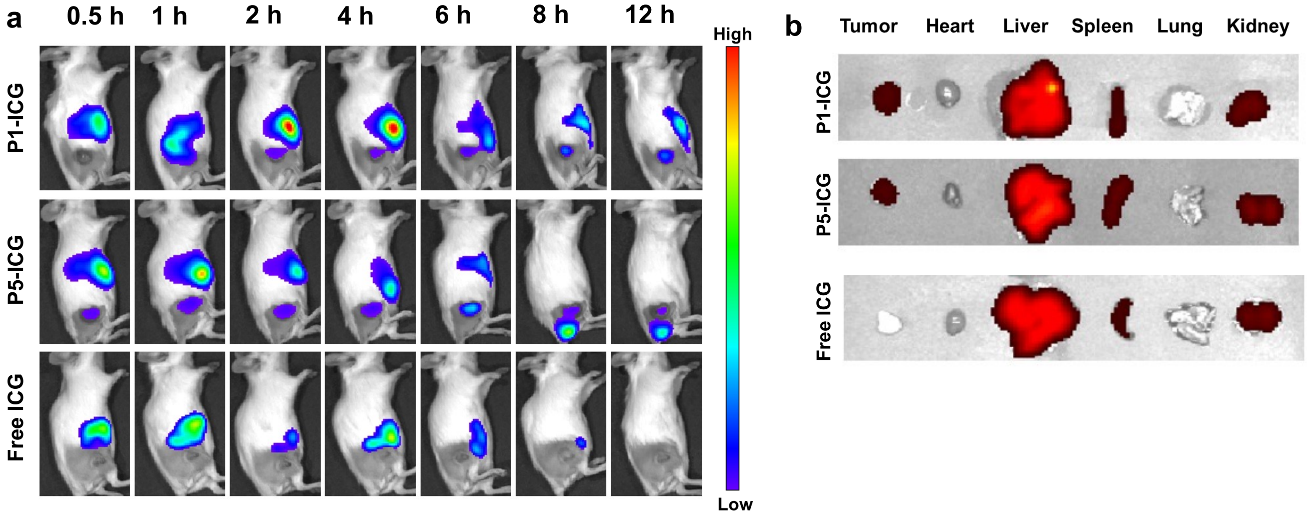

**Figure S31. The in vivo experiments for the two positive peptides (P1 and P5) obtained through library screening. a**, In vivo fluorescence imaging was conducted on H1975 xenograft model mice at various time points after intravenous injection of P1-ICG, P5-ICG, and free ICG, each administered at a dose of 50 µM. **b**, Ex vivo images of tumors and main organs (heart, liver, spleen, lung, kidney) that were taken 12 hours after post-injection.

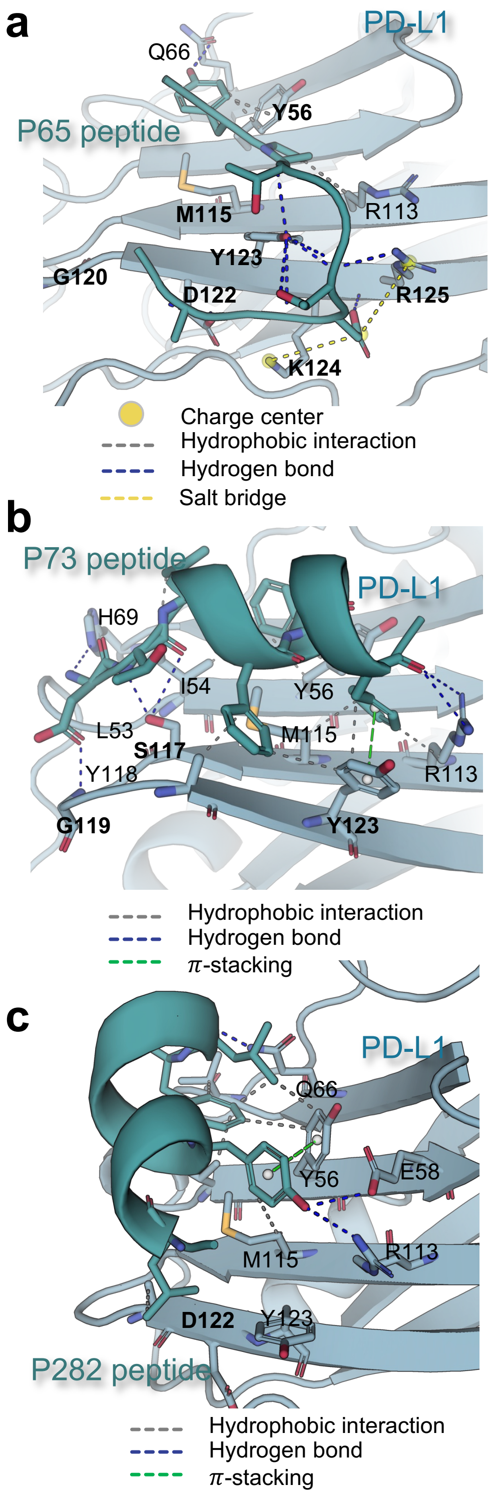

**d**

| Peptide | Receptor | KD (M) | KD ratio |
|---|---|---|---|
| P65 | WT | 2.61 ×10-8 | NA |
| | **Y56A** | 7.00 ×10-7 | 26.82 |
| | Q66A | 1.03 ×10-7 | 3.95 |
| | R113A | 2.10 ×10-7 | 8.05 |
| | **M115A** | 5.90 ×10-7 | 22.61 |
| | **G120A** | 2.83 ×10-6 | 108.43 |
| | **D122A** | 1.93 ×10-6 | 73.95 |
| | **Y123A** | 1.12 ×10-6 | 42.91 |
| | **K124A** | 8.18 ×10-7 | 31.34 |
| | **R125A** | 1.22 ×10-5 | 467.43 |

**e**

| Peptide | Receptor | KD | KD ratio |
|---|---|---|---|
| P73 | WT | 4.49 ×10-8 | NA |
| | L53A | 1.62 ×10-8 | 0.36 |
| | I54A | 6.05 ×10-8 | 1.35 |
| | Y56A | 3.44 ×10-7 | 7.66 |
| | H69A | 1.31 ×10-7 | 2.92 |
| | R113A | 4.88 ×10-8 | 1.09 |
| | M115A | 1.26 ×10-7 | 2.81 |
| | **S117A** | 7.06 ×10-7 | 15.72 |
| | Y118A | 1.60 ×10-7 | 3.56 |
| | **G119A** | 4.16 ×10-7 | 9.27 |
| | **Y123A** | 6.08 ×10-7 | 13.54 |

**f**

| Peptide | Receptor | KD | KD ratio |
|---|---|---|---|
| P282 | WT | 6.18 ×10-8 | NA |
| | Y56A | 4.90 ×10-8 | 0.79 |
| | E58A | 2.41 ×10-8 | 0.39 |
| | Q66A | 4.83 ×10-8 | 0.78 |
| | R113A | 7.54 ×10-8 | 1.22 |
| | M115A | 9.32 ×10-8 | 1.51 |
| | **D122A** | 8.18 ×10-7 | 13.24 |
| | Y123A | 1.50 ×10-7 | 2.43 |

**Figure S32. Binding structures generated by PocketXMol and measured $K_D$ changes for binding-residue mutants of three designed peptides (P65, P73, and P282). a-c**, Predicted binding structures with interaction profiles identified by the Protein–Ligand Interaction Profiler (PLIP)[68]. Binding residues are annotated. **d-f**, Experimental $K_D$ values for wild-type receptors and mutants with alanine substitutions at binding residues. $K_D$ ratios indicate the mutant $K_D$ relative to the corresponding wild-type. Residues with $K_D$ close to or exceeding a 10-fold decrease are shown in bold.

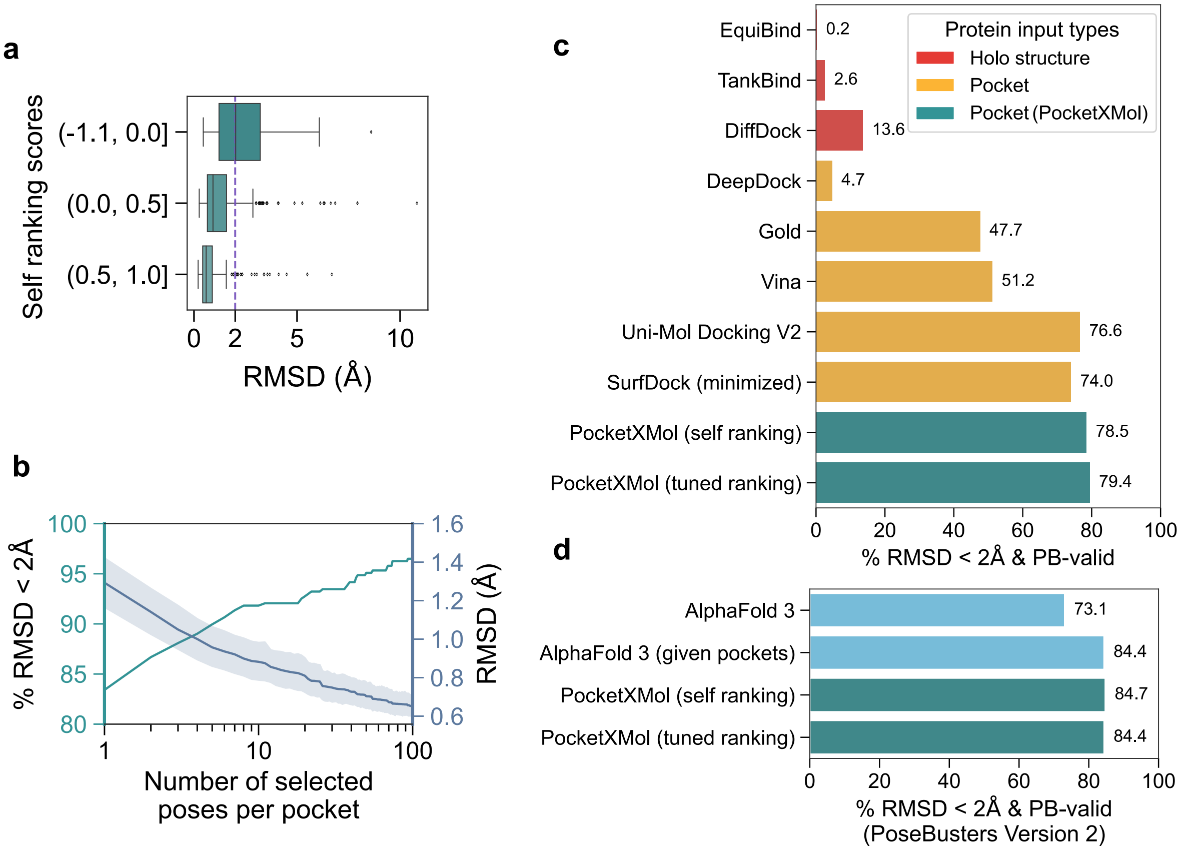

**Figure S33. Extended results for small molecule docking. a**, The relationship between PocketXMol's self-ranking scores and the RMSDs for the docked poses. **b**, Docking performance of selecting different numbers of poses from the 100 generated ones per pocket for evaluation. **c**, The ratios of docked poses that were both RMSD <2Å and PB-valid for different methods in the PoseBusters set (version 1, 428 samples). **d**, The ratios of docked poses that were both RMSD <2Å and PB-valid for PocketXMol and AlphaFold 3 in version 2 of the PoseBusters set (308 samples).

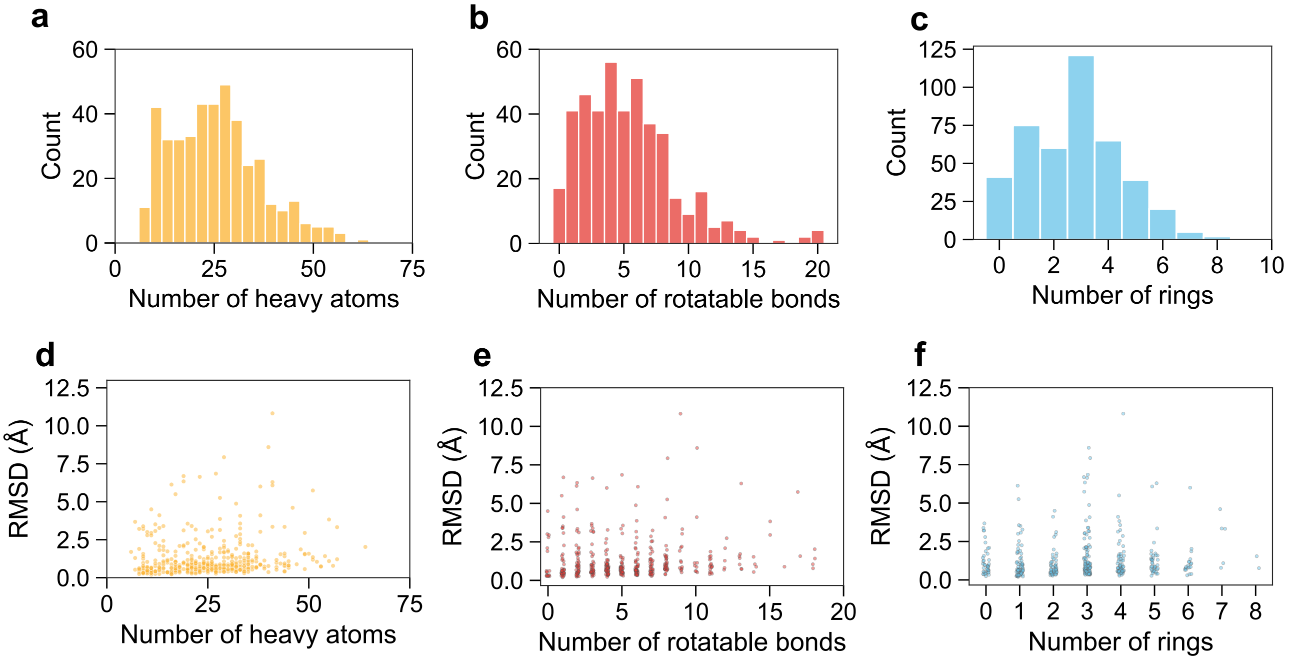

**Figure S34.** **Relationships between docking performance and different molecular properties.** Count distributions for different molecules for (**a**) heavy atoms, (**b**) rotatable bonds, and (**c**) rings in the test set. Docking RMSDs for different (**d**) heavy atoms, (**e**) rotatable bonds, and (**f**) rings of the molecules.

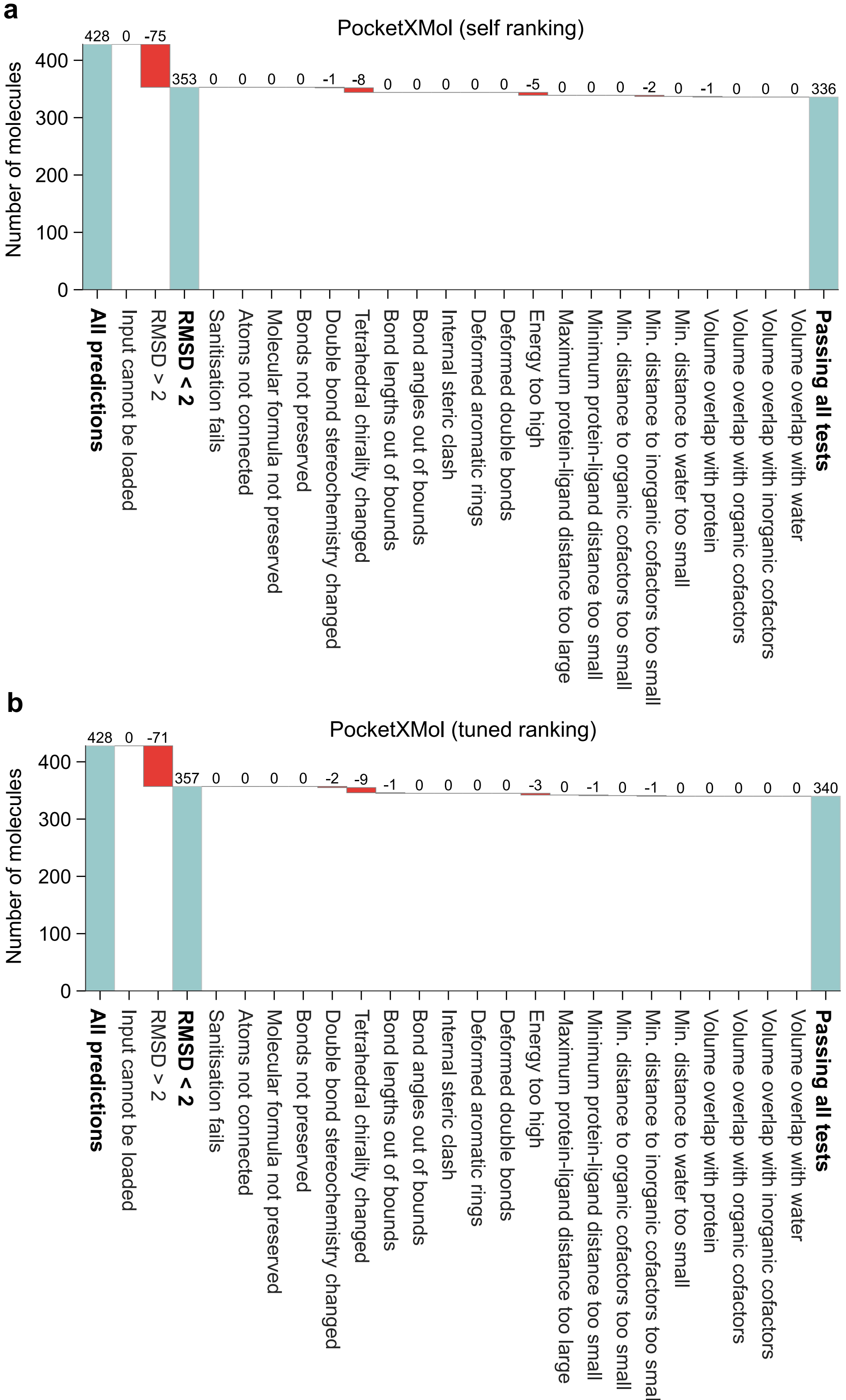

**Figure S35. The number of molecules that satisfy the 3D structural screening criteria proposed by Posebusters. a**, Generated by PocketXMol with self ranking scores. **b**, Generated by PocketXMol with tuned ranking scores.

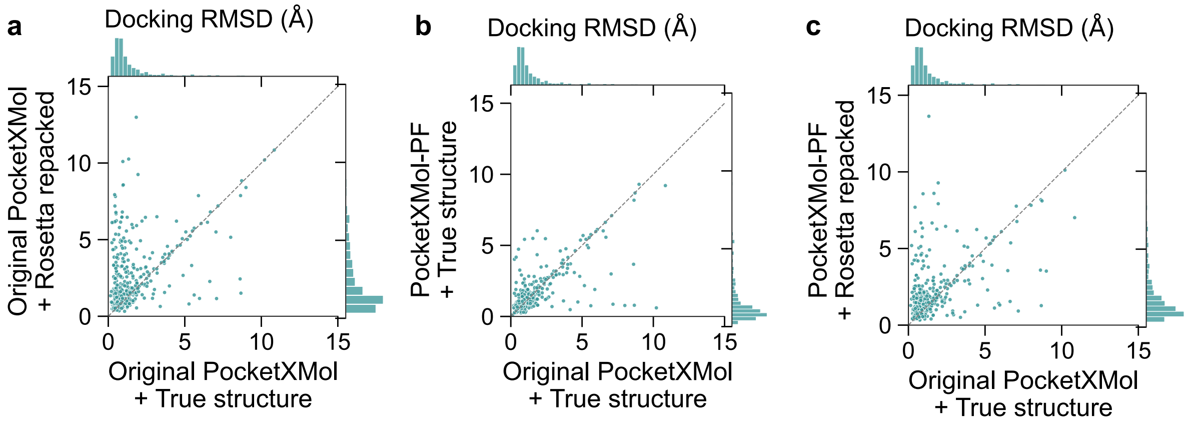

**Figure S36. Small molecular docking with different PocketXMol and input pocket structures.** **a**, Comparison of docking RMSDs of the original PocketXMol with true pocket structures and with true structures. **b**, Comparison of docking RMSDs between the original PocketXMol with true pocket structures and the finetuned PocketXMol with true structures. **c**, Comparison of docking RMSDs between the original PocketXMol with true pocket structures and the finetuned PocketXMol with Rosetta-repacked structures.

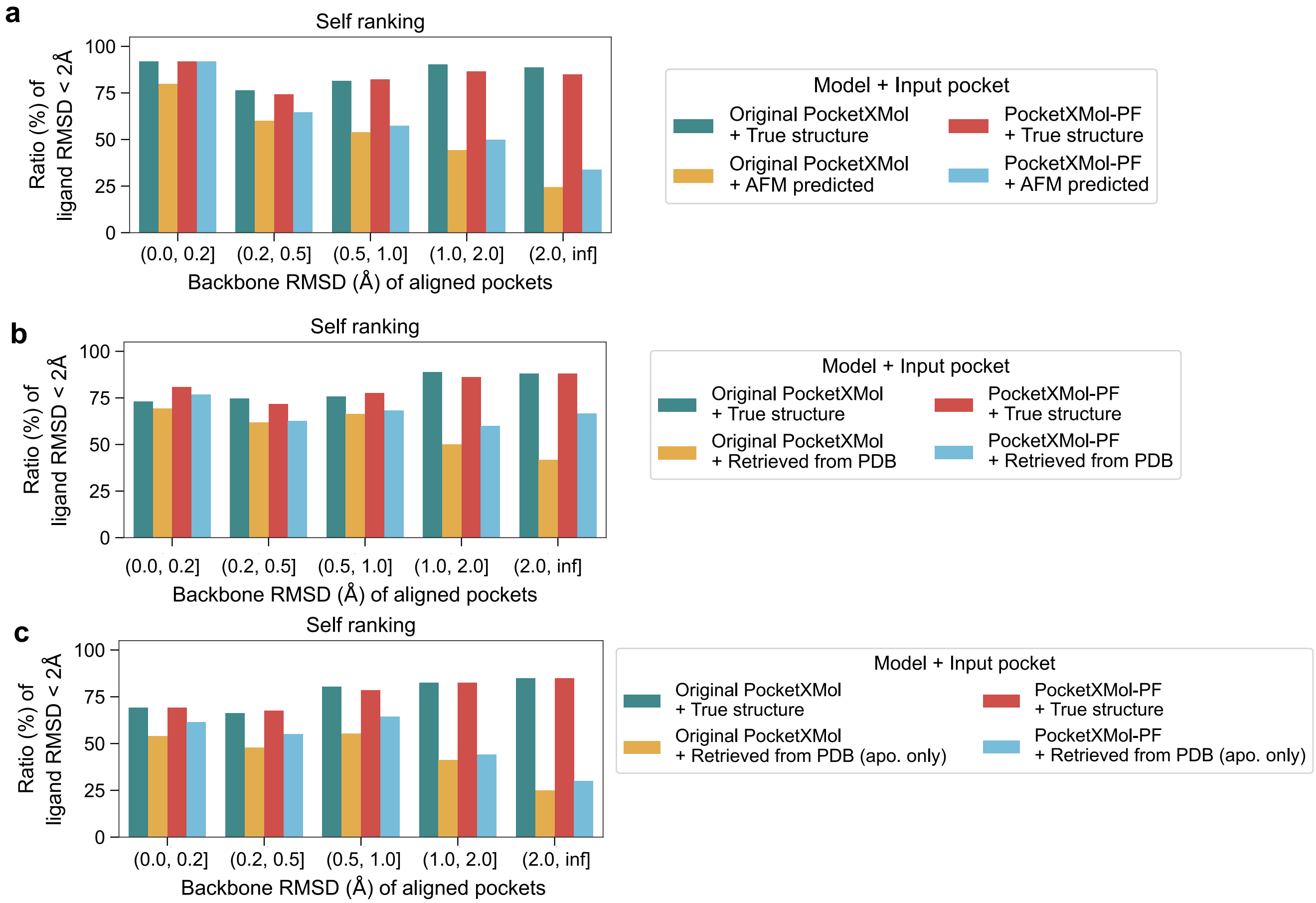

**Figure S37. Relationship between the performance of small molecular docking and the pocket RMSDs for PocketXMol (self ranking) with different input structures. a**, Comparison for input pocket structures predicted by AlphaFold-Multimer. **b**, Comparison for input pocket structures retrieved from RCSB PDB. **c**, Comparison for input pocket structures retrieved from apo structures in RCSB PDB.

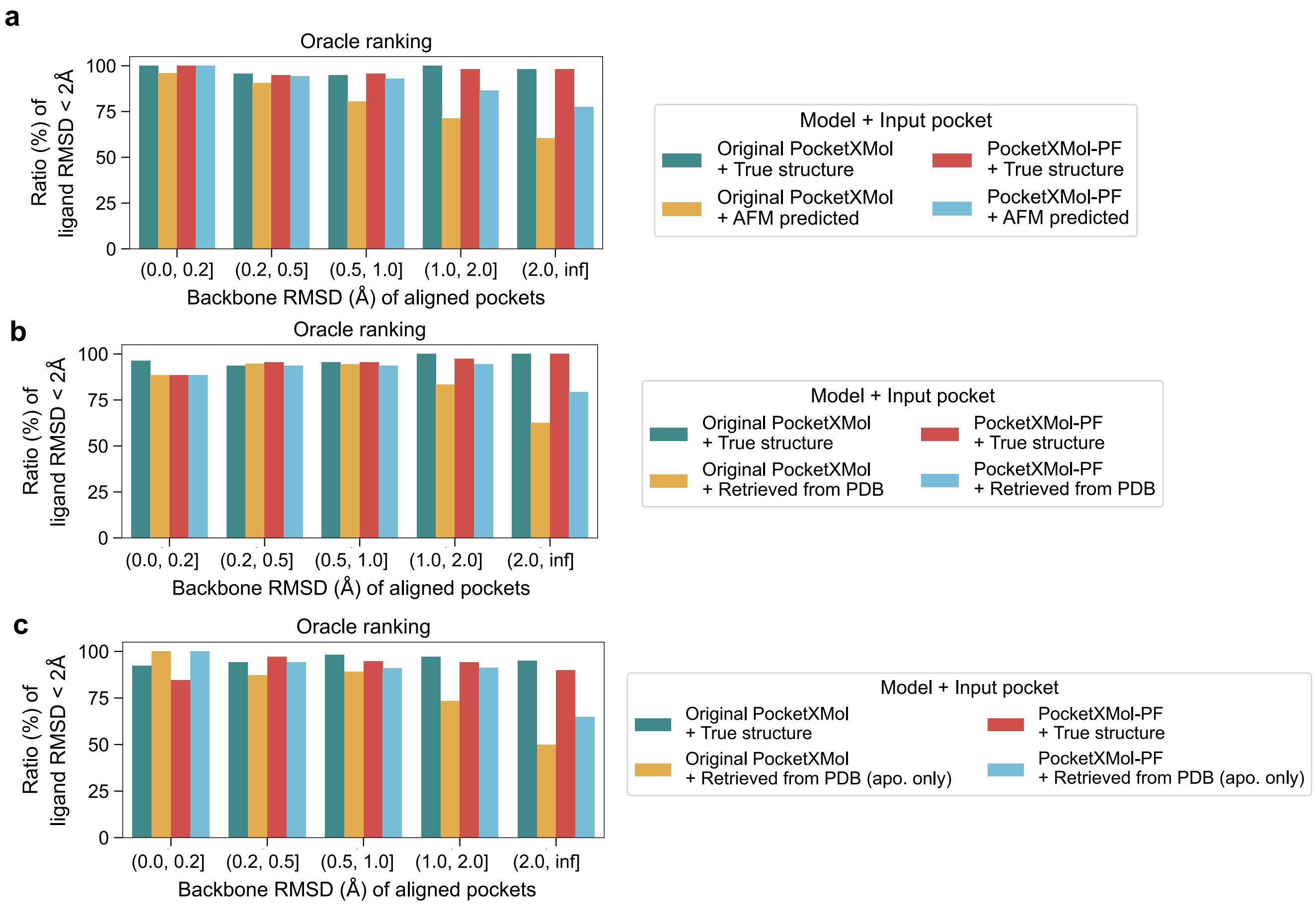

**Figure S38. Relationship between the performance of small molecular docking and the pocket RMSDs for PocketXMol (oracle ranking) with different input structures. a**, Comparison for input pocket structures predicted by AlphaFold-Multimer. **b**, Comparison for input pocket structures retrieved from RCSB PDB. **c**, Comparison for input pocket structures retrieved from apo structures in RCSB PDB.

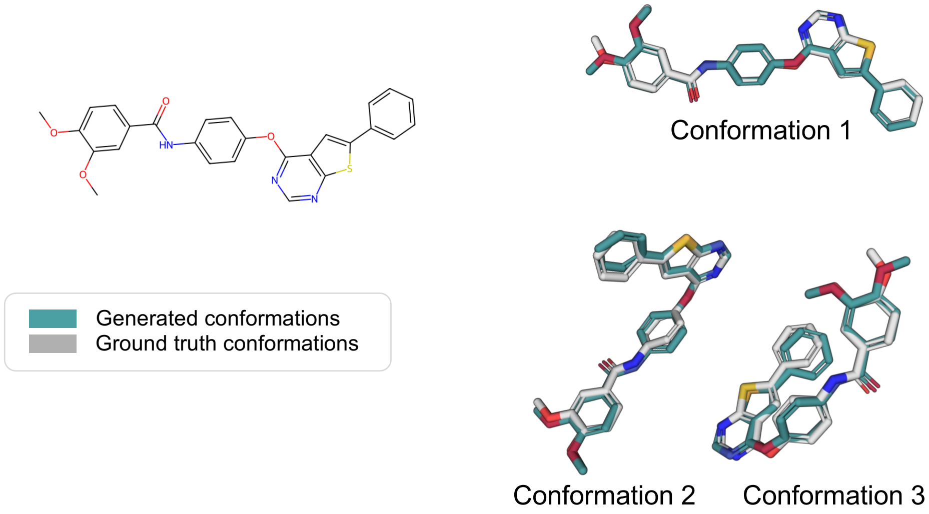

**Figure S39.** **An example showing PocketXMol could generate high-quality conformations for a 2D molecular graph (ChEMBL ID: CHEMBL1307825).** The three conformations here represent three low-energy conformations of this molecule in the dataset.

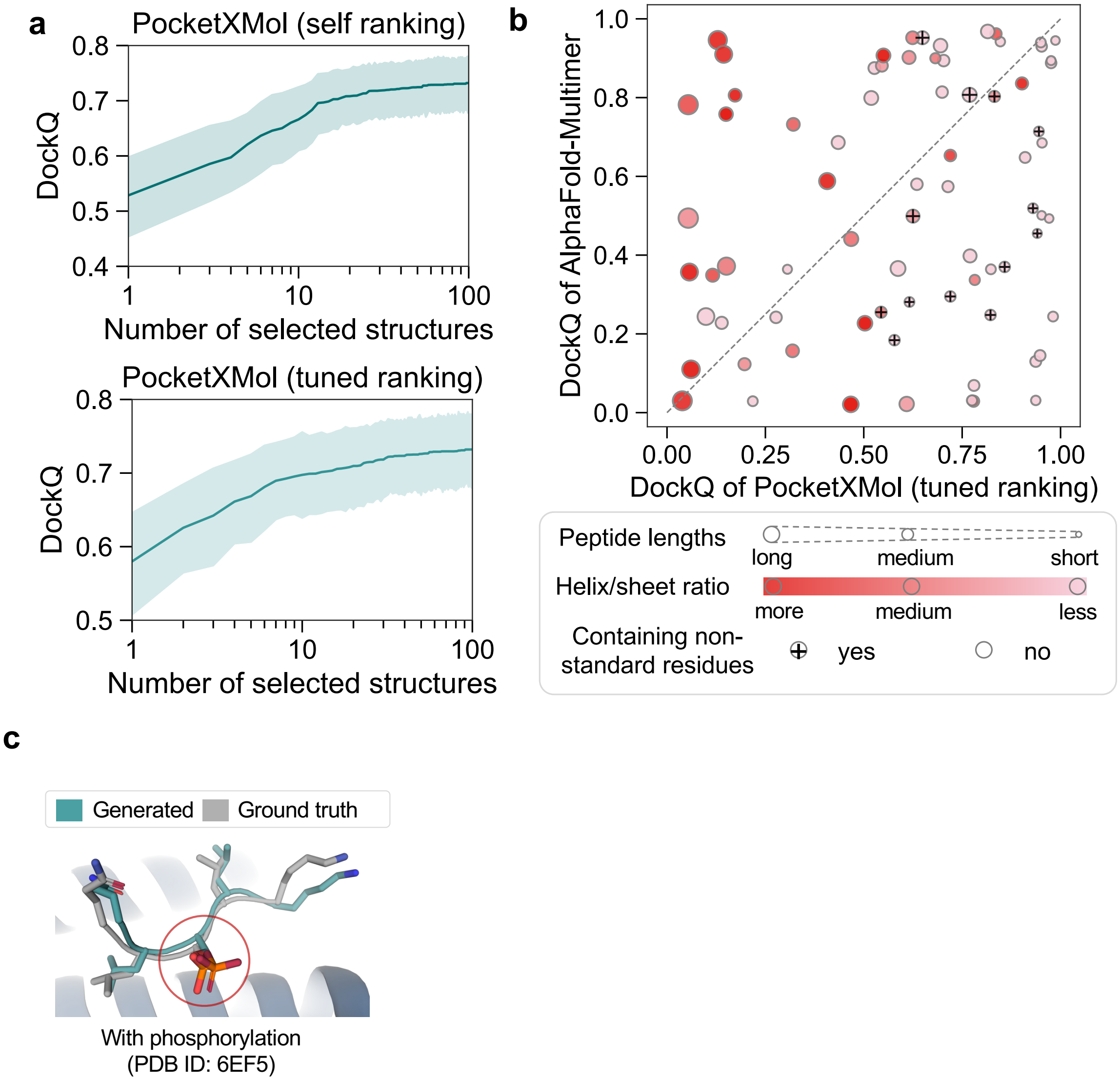

**Figure S40. Extended results for peptide docking on the PepBDB set. a**, Docking performance of selecting different numbers of peptides from the 100 generated structures per sample. **b**, DockQ for the protein-peptides docked by AlphaFold-Multimer and PocketXMol. The size, color, and marker represent the peptide length, the ratio of different secondary structures, and whether the peptides contain non-standard amino acids, respectively. **c**, an example of docked peptide with phosphorylation

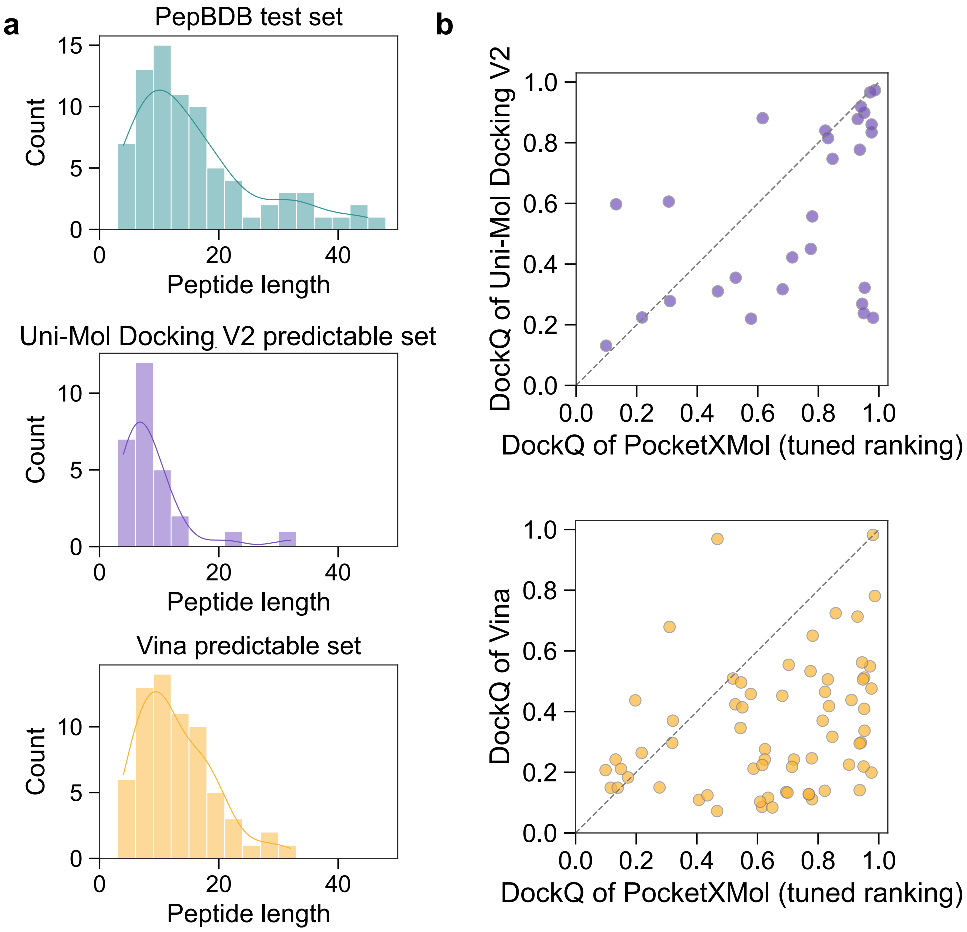

**Figure S41. Peptide docking performance (on the PepBDB set) of small-molecule docking tools.** **a**, The distributions of peptide lengths in the peptide docking test set, the Uni-Mol Docking V2 predictable set, and the Vina predictable set. The predictable peptides are those that the tools can output the docking results. Some large peptides may cause errors or get stuck for these tools originally designed for small molecules and thus are not predictable for them. **b**, Comparison of the peptide docking performance between PocketXMol and the small-molecular docking tools.

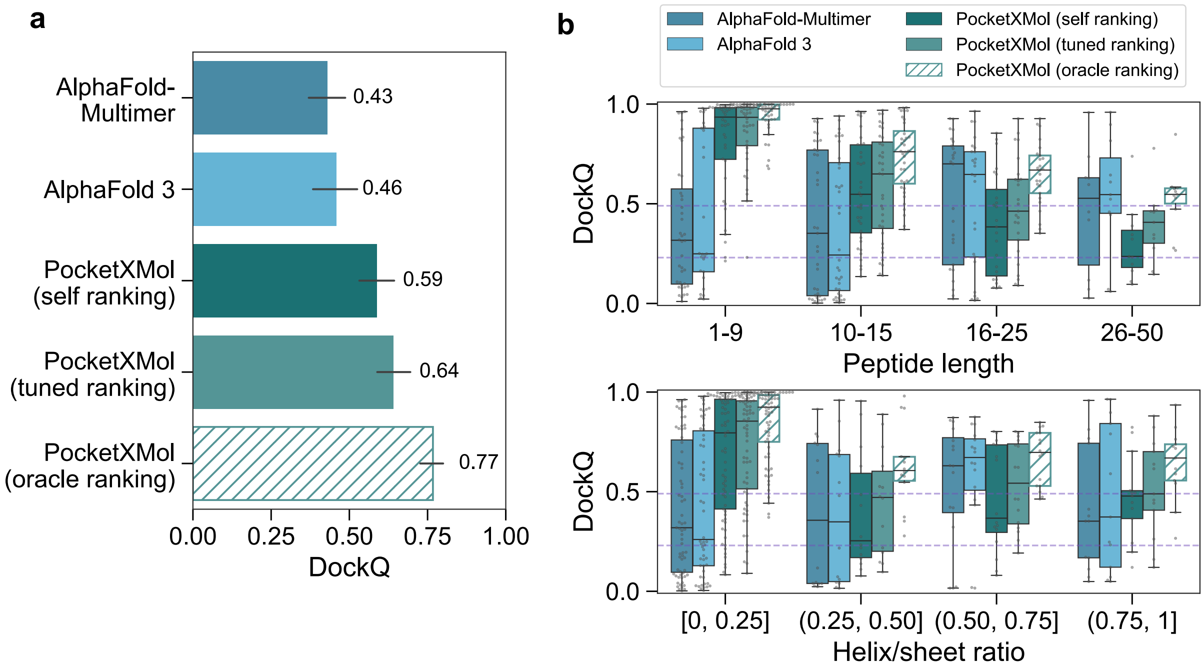

**Figure S42. Docking performance on the BioLiP peptide docking test set for AlphFold-Multimer, AlphaFold 3, and PocketXMol. a**, DockQ of protein-peptide docking by different models. Here we only considered the best baselines AlphaFold-Multimer and AlphaFold 3. Error bars represent 95% confidence intervals ($n = 112$). **b**, DockQ for peptide docking with respect to the peptide length (top) and ratio of secondary structures helix and sheet (bottom). The box plot outlines quartiles of the metrics with whiskers spanning up to 1.5 times the interquartile range ($n = 112$).

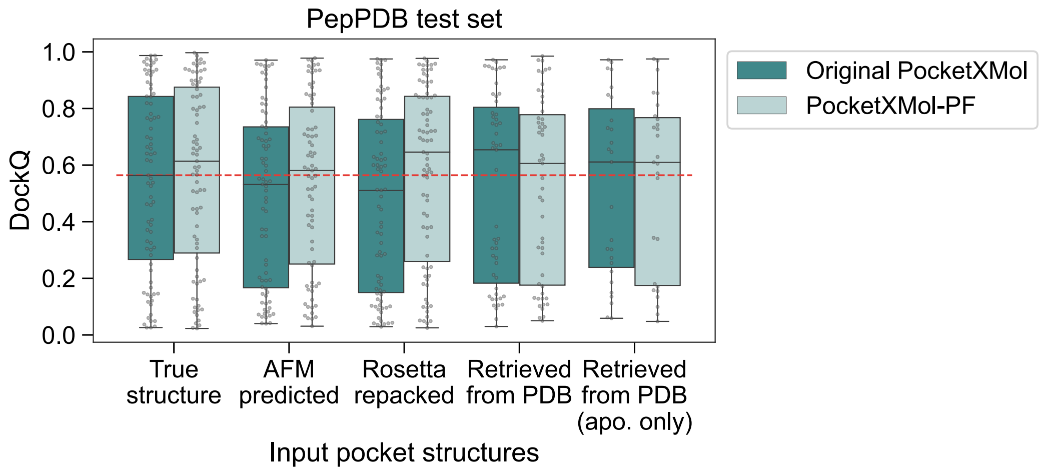

**Figure S43. Peptide docking on PepBDB set for different PocketXMol and different input pocket structures.** DockQ for different PocketXMol (original and fine-tuned ones) and different input pocket structures. The pocket structures included the true structures of the data, structures predicted by AlphaFold-Multimer, structures with side-chains repacked by Rosetta, all structures in RCSB PDB that had the same sequences as the data, and apo structures in RCSB PDB that had the same sequences as the data.

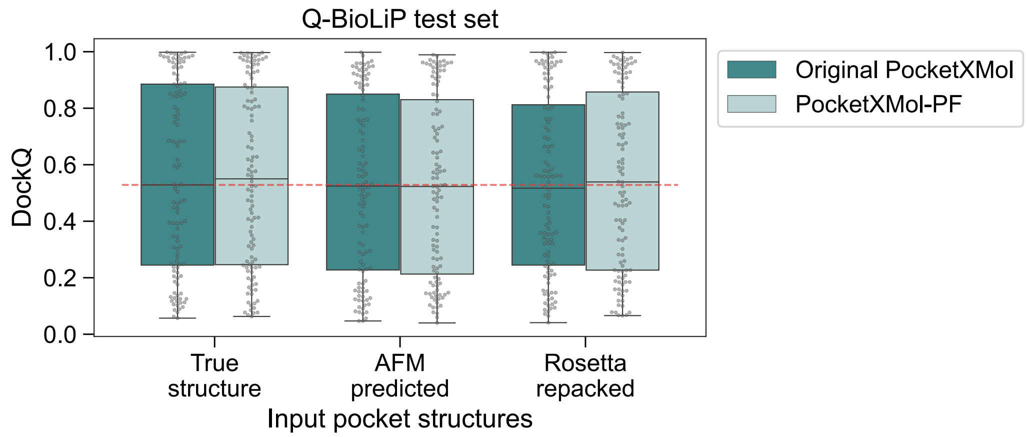

**Figure S44. Peptide docking on the Q-BioLiP set for different PocketXMol and different input pocket structures. a**, DockQ for different PocketXMol (original and fine-tuned ones) and different input pocket structures. The pocket structures included the true structures of the data, structures predicted by AlphaFold-Multimer, and structures with side-chains repacked by Rosetta. For the Q-BioLiP test set, we did not show the performance for structures retrieved from RCSB PDB because the proteins in this test set are mostly novel sequences that had few resolved structures in RCSB PDB.

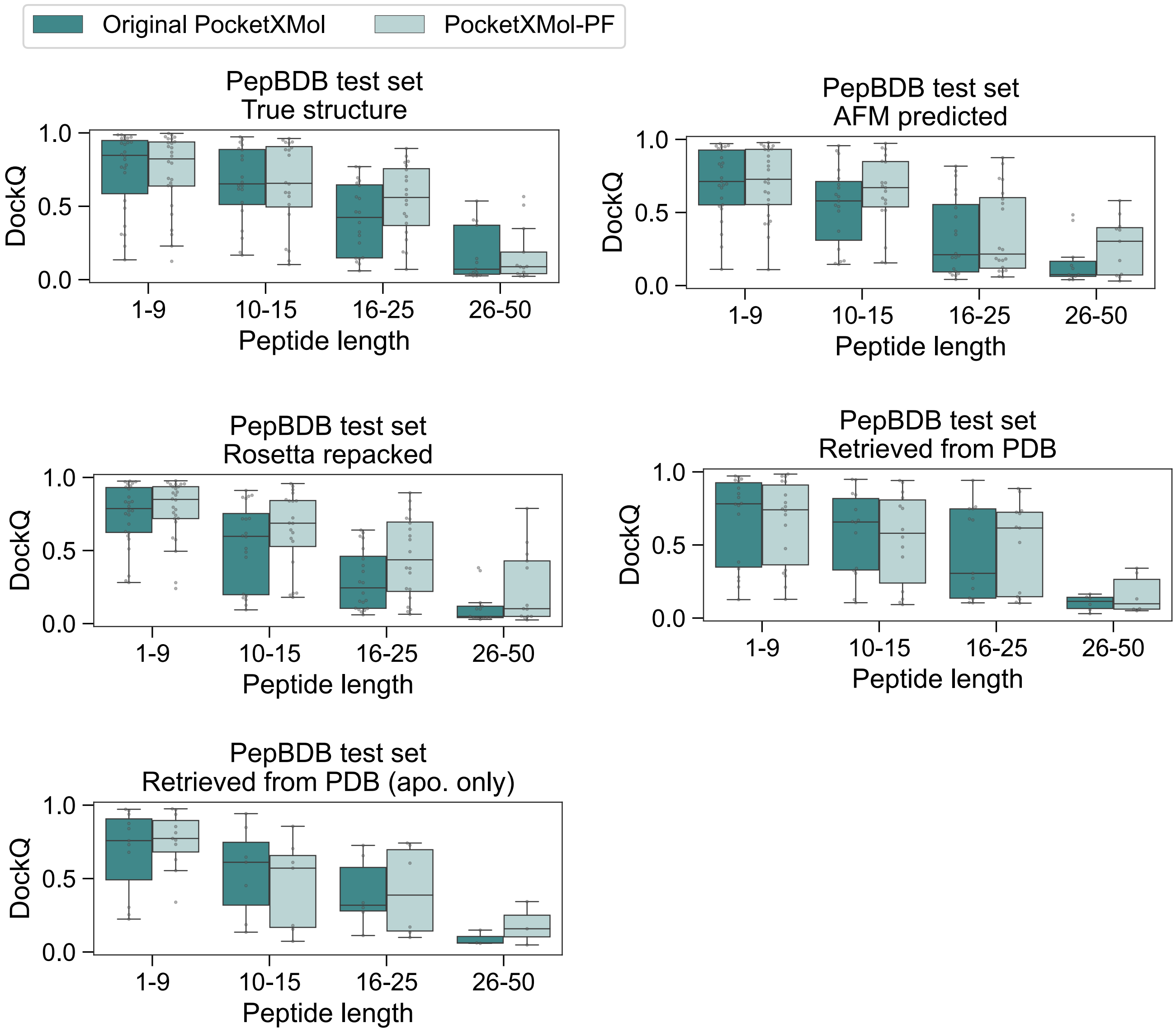

**Figure S45.** Relationship between DockQ of peptide docking (on the PepBDB set) and the peptide length in cases of different PocketXMol and input pocket structures.

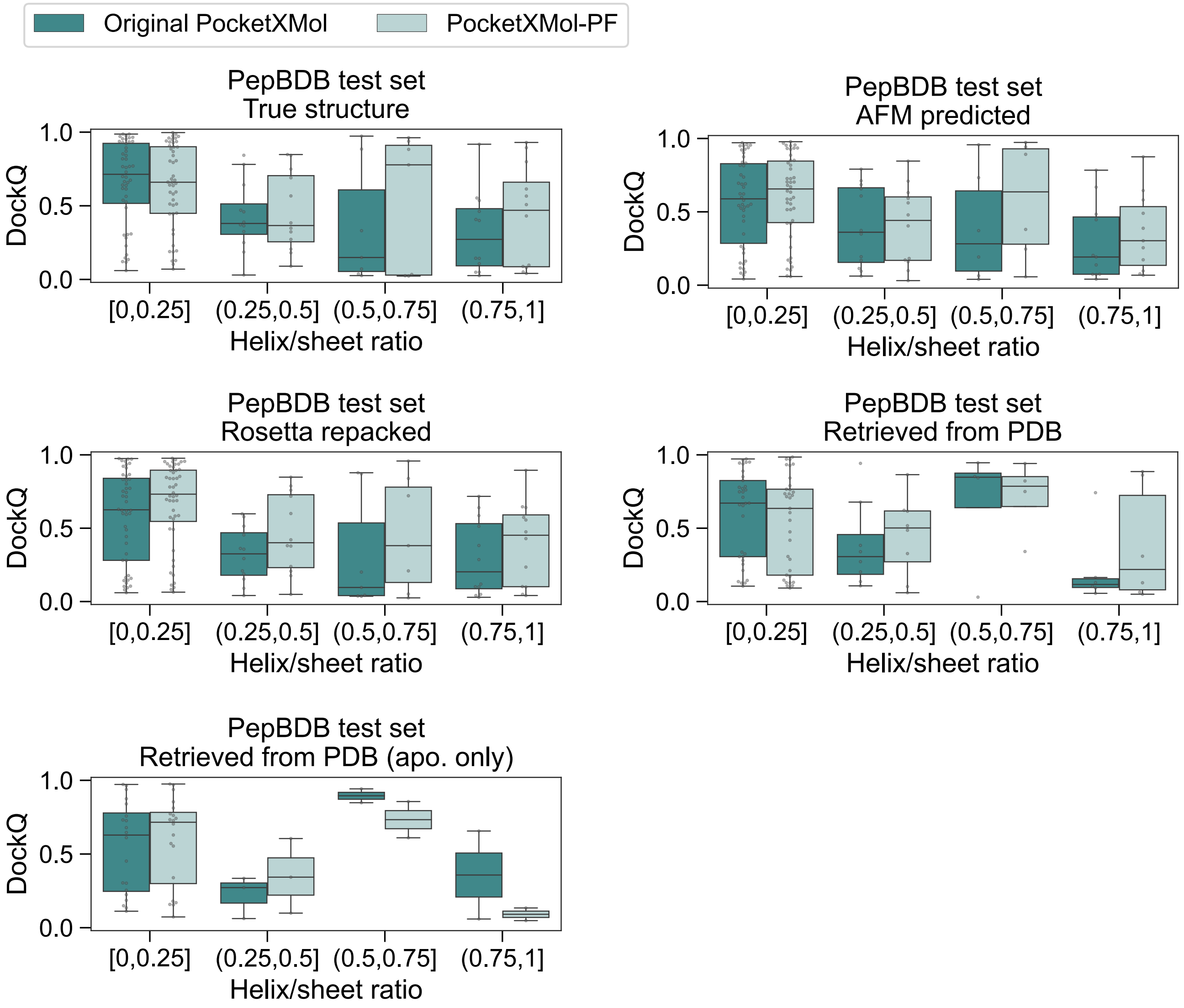

**Figure S46.** Relationship between DockQ of peptide docking (on the PepBDB set) and the ratio of secondary structures helix and sheet in cases of different PocketXMol and input pocket structures.

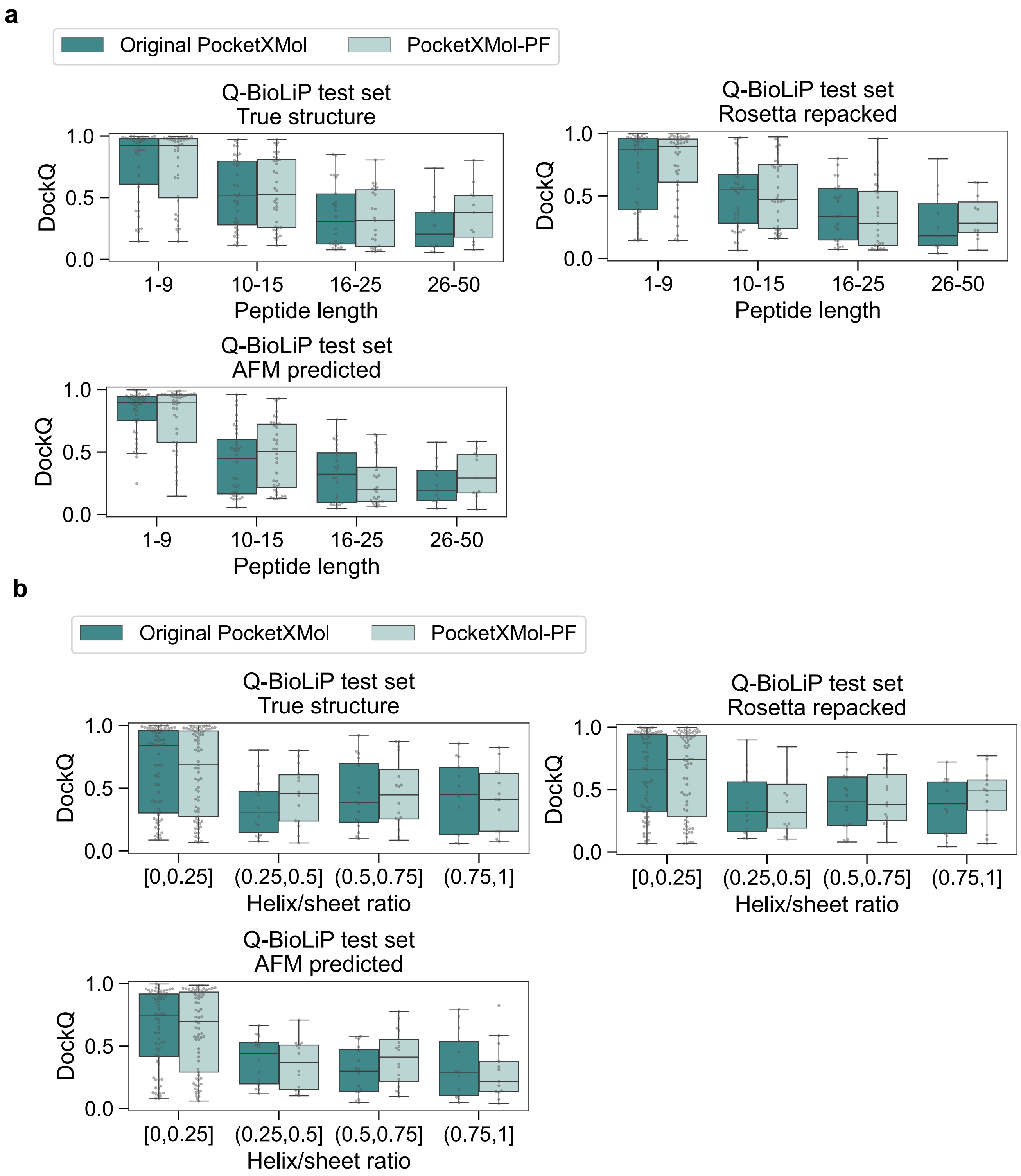

**Figure S47. Relationship between DockQ of peptide docking (on the Q-BioLiP set) and the peptide properties in cases of different PocketXMol and input pocket structures. a**, Relationship between DockQ and the peptide length. **b**, Relationship between DockQ and the ratio of secondary structures helix and sheet.

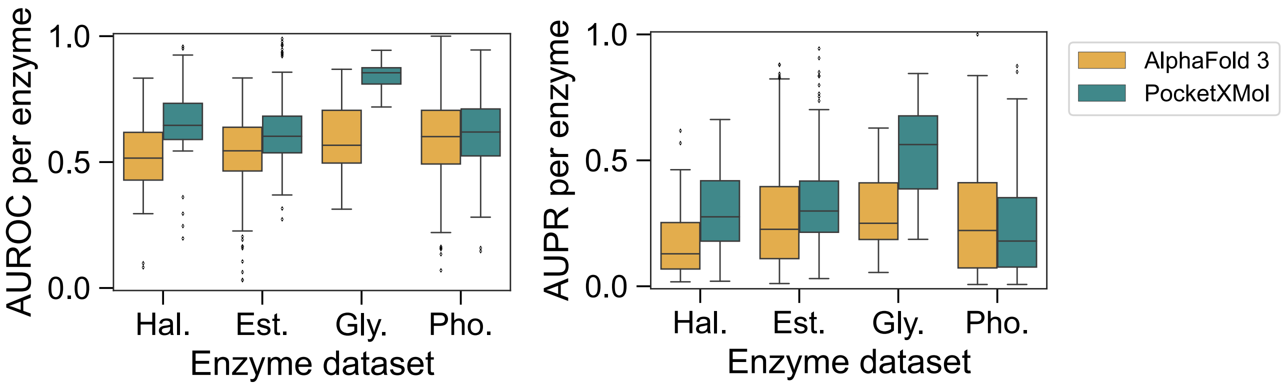

**Figure S48. Performance of substrate activity prediction by model confidence scores.** For each enzyme-substrate pair, we used the PocketXMol or AlphaFold 3 to predict the complex structures. Then, the confidence scores of the models were used as the metrics to distinguish whether the pair was active or not. We calculated (**a**) the AUROC (the area under the receiver operating curve) scores and (**b**) the AUPR (area under precision-recall curve) scores for individual enzyme proteins. The x-axis is different enzyme datasets and represent Halogenase, Glycosyltransferase, Esterase, and Phosphatase, from left to right. $P$ values of one-sided $t$-test are $2.18 \times 10^{-5}$, $8.41 \times 10^{-6}$, $1.70 \times 10^{-16}$, and 0.099 for AUC scores, and were $7.62 \times 10^{-8}$, $4.58 \times 10^{-5}$, $9.85 \times 10^{-15}$, and 0.9986 for AUPR scores from left to right, respectively.

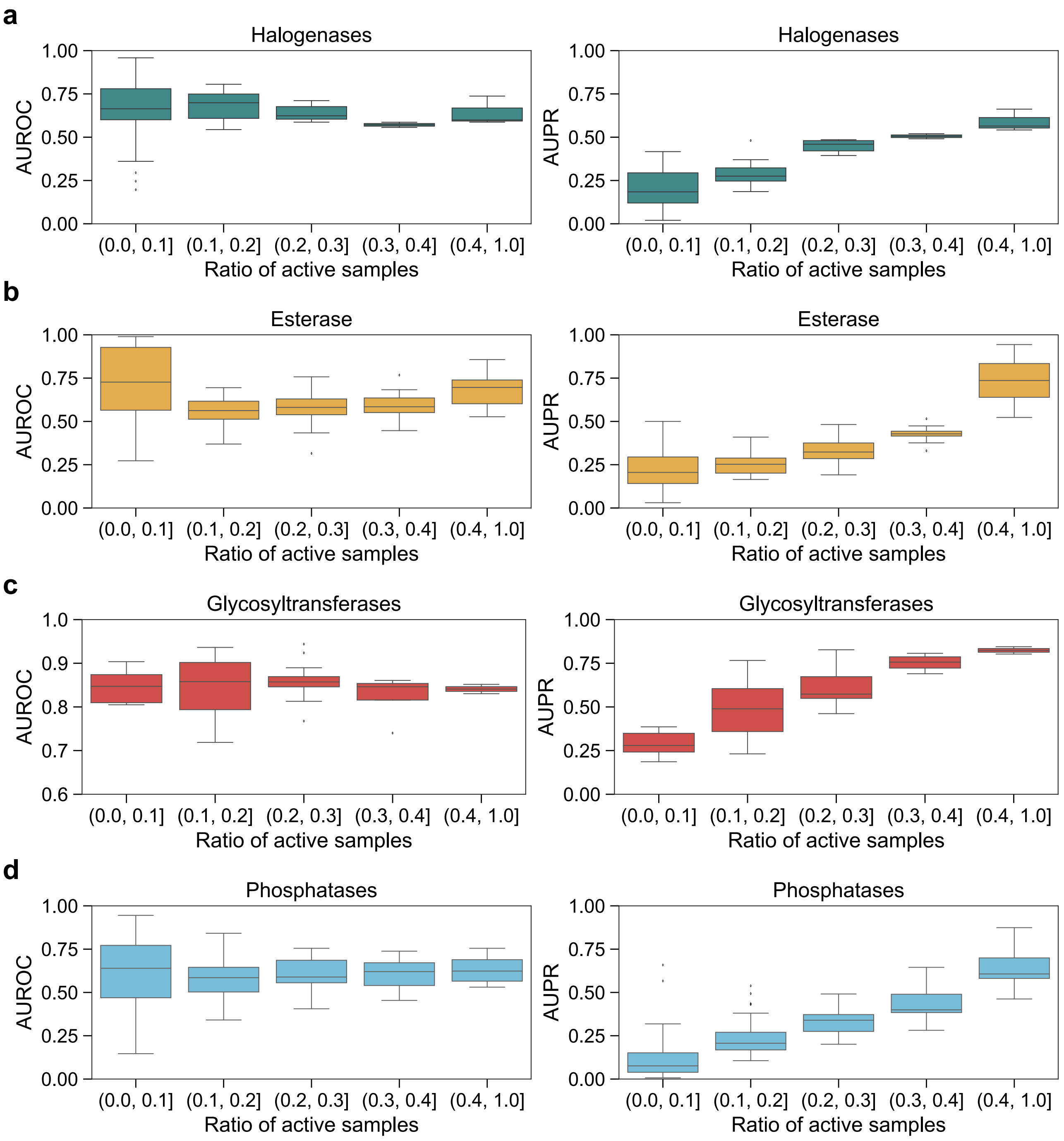

**Figure S49. Relationship between the enzyme-substrate activity prediction metric and ratio of active samples of the enzyme proteins.** AUROC and AUPR scores were calculated for each protein in enzyme datasets, including (**a**) Halogenase, (**b**) Glycosyltransferase, (**c**)Esterase, and (**d**) Phosphatase.

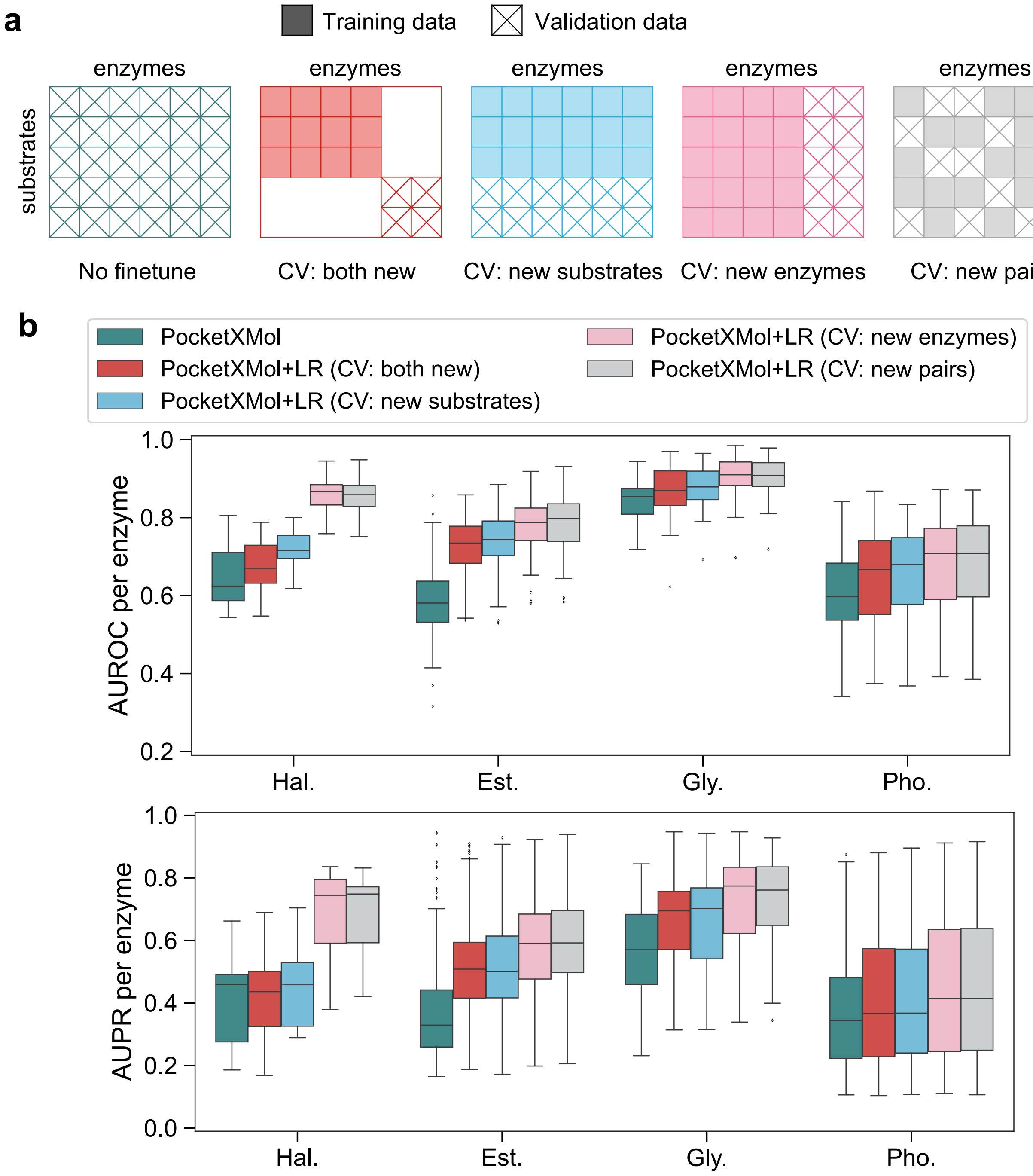

**Figure S50.** **Performance on enzyme activity prediction using logistic regression models and PocketXMol representation vectors.** **a**, Schematic illustration of various training–validation split strategies used in cross-validation. **b**, AUROC and AUPR scores for each enzyme under different split settings, using logistic regression models with PocketXMol-derived representation vectors as input features. These vectors were extracted from PocketXMol by encoding the enzyme–substrate complex with a docking prompt and collecting the intermediate feature representations prior to the final prediction layers. All cross-validation experiments were performed using ten folds, except for the "both-new" setting, which employed a nine-fold (3×3) scheme to ensure that every data point served as a validation sample in one fold. The x-axis is different enzyme datasets and represent Halogenase, Glycosyltransferase, Esterase, and Phosphatase, from left to right.

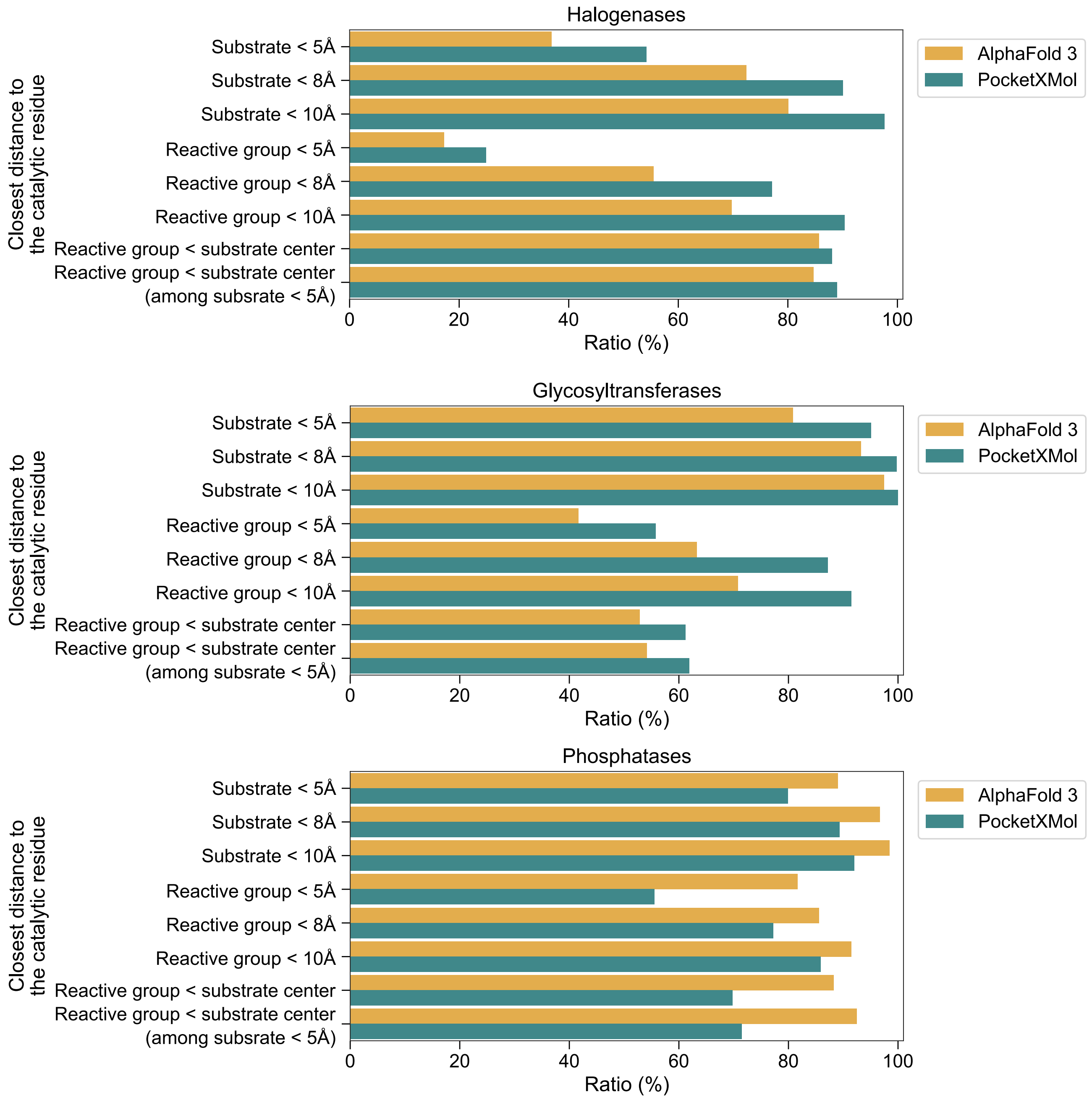

**Figure S51. Comparison of predicted enzyme–substrate structure characteristics between PocketXMol and AlphaFold 3.** Distances were computed between the side chains of catalytic residues in the enzyme and various atomic regions of the substrate. The following distance-based metrics were evaluated as the proportion of samples with distances below specific thresholds: (1) Substrate distance: the minimum distance between any atom of the substrate and the catalytic residues. (2) Reactive group distance: the minimum distance between any atom in the reactive group of the substrate and the catalytic residues. (3) Substrate center distance: the average distance from all substrate atoms to the catalytic residues.
